# Mechanistic Gene Networks Inferred from Single-Cell Data are Better Predictors than Neural Networks

**DOI:** 10.1101/2021.05.12.443819

**Authors:** Jung Min Han, Sudheesha Perera, Zeba Wunderlich, Vipul Periwal

## Abstract

With advances in single-cell techniques, measuring gene dynamics at cellular resolution has become practicable. In contrast, the increased complexity of data has made it more challenging computationally to unravel underlying biological mechanisms. Thus, it is critical to develop novel computational methods capable of dealing with such complexity and of providing predictive deductions from such data. Many methods have been developed to address such challenges, each with its own advantages and limitations. We present an iterative regression algorithm for inferring a mechanistic gene network from single-cell data. Using this regression, we infer a developmental model for the gene dynamics in *Drosophila melanogaster* blastoderm embryo. Our results show that the predictive power of the inferred model is higher than that of other models inferred with least squares and ridge regressions.

Furthermore, model predictions of the gene dynamics are more accurate than predictions made with neural networks. This holds true even in the limit of small sample sizes. We compare predictions for various gene knockouts with published experimental results, finding substantial agreement. We also make predictions for gene dynamics under various gene network perturbations, impossible in non-mechanistic models.

## 1 Introduction

Gene regulatory networks govern the dynamics of gene expression levels, which, ultimately, control cellular function. Deciphering the blueprint of regulatory networks is a long-standing challenge in systems biology. On account of advances in single-cell technology, there is an increasing interest in reverse engineering or inferring gene regulatory networks from single-cell data. Many algorithms have been developed to model gene regulatory networks, including regression approaches [21, 22], Bayesian modeling [9, 17], and mutual information [7, 23]. Inference methods have different domains of applicability, as suggested by many review papers that discuss and compare different inference methods based on their underlying biological data and theoretical approaches [2, 5, 10, 11].

Here we present a regression method for the inference of a mechanistic gene network that drives the discrete gene dynamics observed in *Drosophila melanogaster* blastoderm embryo. The data, published in Fowlkes et al. [4], were obtained through single-cell fluorescence microscopy, and contain gene expression of a set of genes at cellular spatiotemporal resolution. Instead of counting the total number of reads as in single-cell RNA-sequencing (scRNA-seq), which involves the isolation of individual cells and their RNA, reverse transcription, amplification, library generation, and sequencing, fluorescence microscopy involves *in situ* hybridization of probes with a fluorescence tag to target genes and quantifying the amount of fluorescence with imaging. One of the advantages of this data set is that it contains cellular resolution spatial information of the measured gene expressions, which allows for the analysis of local gene expression variation. However, imgaing can distinguish only a limited number of different fluorophores simultaneously, putting a restriction on obtaining comprehensive gene profiles of a given sample [3, 16]. We give details about the data we use in Section 2.1.

For network inference, we introduce a stable variant of least absolute deviation (LAD) regression that inherits its robustness to outliers, while rectifying its instability in parameter optimization. The efficiency of the regression is rated by how well the inferred gene network model captures the observed discrete gene dynamics. Our results show that the gene expression predictions from our model are more accurate than those from models inferred with other regressions and predictions obtained using neural networks. More importantly, in the limit of small sample size, the models from our regression method with *L*_2_ regularization perform better than standard ridge regression or neural networks. We then address the universality of the inferred gene network and discuss how cell-to-cell variability in gene dynamics affects the model inference. We compare our model predictions with published experimental results on the development of the blastoderm under various knockout conditions, finding substantial agreement. Finally, we perform a series of model predictions by introducing small perturbations to individual edges in the gene network and simulating the effects on target genes one time-step later.

## 2 Methods

### 2.1 Data

Animal transcription networks comprise many regulators and target genes. To obtain a comprehensive view of how the dynamics of transcription factors affect their downstream targets, it is necessary to concurrently measure as many gene expression levels as possible. However, with conventional fluorescence microscopy, we can only record the expression of a few gene products at a time. To overcome this experimental limitation, Fowlkes et al. [4] used a computational method developed by Rübel el at. [16], which uses a registration technique to compile image-based data from hundreds of *Drosophila melanogaster* blastoderm embryos into a common spatiotemporal atlas, so that the average expressions of many gene products can be captured.

The method consists of two key steps: spatial and temporal registration of embryo nuclei. A total of 1822 embryos were fluorescently stained to label the mRNA expression patterns of two genes, one of which was either *eve* or *ftz* as a reference gene, and nuclear DNA. In total, 95 mRNA and four protein expressions were labeled. The embryos were grouped into six cohorts based on their developmental stages, which spanned 50 minutes prior to the onset of gastrulation. Within each temporal cohort, the reference gene profiles, the average nuclear distribution, and the mean embryo shape were used to build a morphological template with a fixed number of nuclei that represented the average embryo. Then, correspondences between nuclei across the cohorts were established. As a result, a total of 6078 nuclei were registered spatially and temporally between six cohorts. Simply put, the results are analogous to having a set of the expression of gene products measured in or around 6078 nuclei at six discrete timepoints that are approximately 10 minutes apart.

### 2.2 Missing value imputation

About 37% of the genes in the dataset were missing data. Out of 99 variables (mRNAs and/or proteins), 27 of them had complete temporal profiles, i.e., recorded at all 6 time-points, while the remaining 72 variables were missing some time-point measurements. From here on, we refer to the 27 genes with complete information as the complete genes and the other 72 genes as the incomplete genes. To interpolate the missing data, we used neural networks. Disregarding the time-point aspect for the time being, we can think of the dataset as a set of *N* = 36468 (6078 nuclei × 6 timepoints) samples with 99 features. To impute missing data of an incomplete gene *i*, we followed the steps below:

1. Identify the samples (nuclei) where the gene *i* was measured.
2. Using the 27 complete genes in the samples from Step 1 as the predictors and the gene *i* measurements in the corresponding samples as the targets, train and validate a neural network. The neural network consists of three layers: an input layer with 27 nodes, one hidden layer with 13 nodes, and an output layer with 1 node.
3. With the trained and validated neural network, make a prediction for the value of gene *i* in the samples where it was not measured, using the 27 complete genes in the corresponding samples as the predictors.

Fig. 1 illustrates the schematics of the imputation. We trained and validated 72 independent neural networks to interpolate the missing data for 72 incomplete genes.

**Figure 1:**
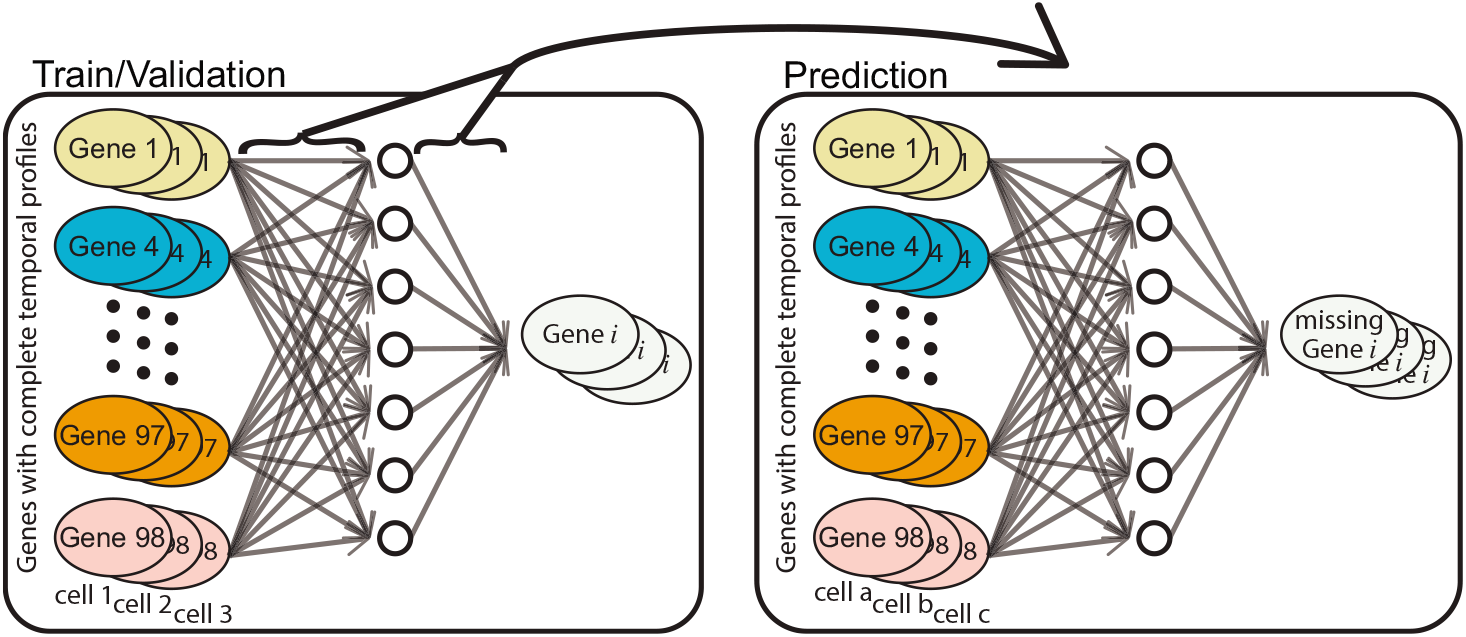
A schematic diagram of the imputation method. The missing expression levels for each incomplete gene were imputed with a neural network that was trained with the complete gene expression levels in the samples where the incomplete gene was measured as predictors and the measured incomplete gene as targets.

We then investigated the performance of our imputation method. From the 27 complete genes, we randomly selected 3 genes and removed some of their time-point measurements and treated those as missing data. Using the remaining 24 complete genes, we followed the steps described above to impute the removed data. We performed 500 trials of this process. Each of the 27 complete genes was selected for 55 trials on average. We then computed the correlation coefficients between the observed data that were treated as missing data and their imputed data. Fig. 2 shows the mean correlation coefficient and the standard deviation for each of the complete genes.

**Figure 2:**
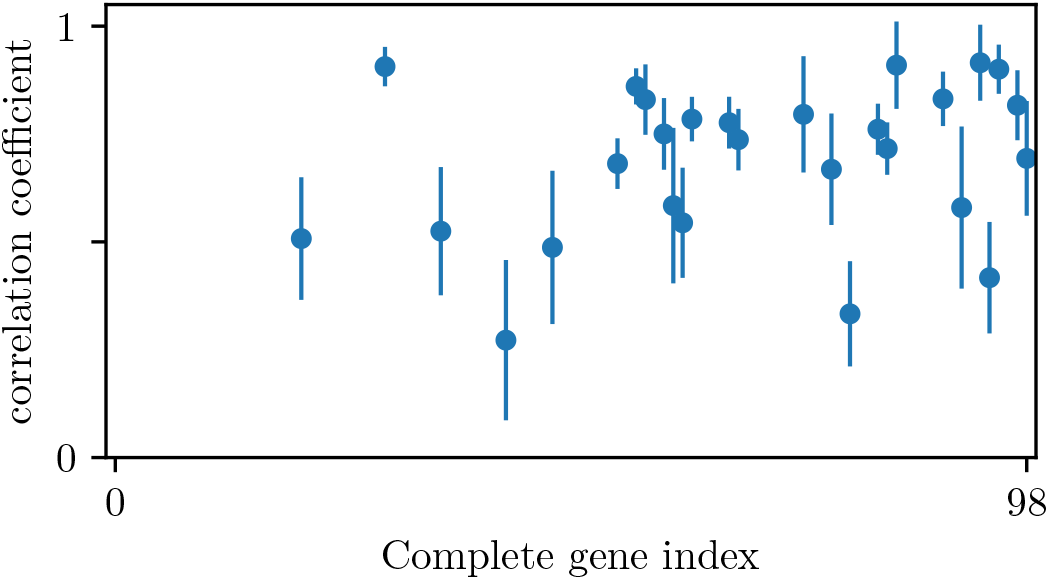
Validation of our imputation method. The plot shows the mean correlation coefficient and the standard deviation between the true and the imputed values for each of the complete genes over approximately 55 independent trials each.

To find the optimal number of nodes in the hidden layer of the neural network, we tried a few different configurations. Initially, we set the default number of nodes on the hidden layer to be 13, approximately half of input variables. We then increased the nodes to 27 and 54 and decreased it to 7 as well. For each configuration of neural networks, we followed the method described above and imputed 3 simulated incomplete genes using 24 complete genes, over 500 trials. We compared the correlation coefficients between the imputed and observed data across all configurations of neural networks, and concluded that the performance of the neural networks is approximately independent of their complexity. The correlation coefficients are shown in the Supporting Materials A (Fig. S1).

### 2.3 Model inference

The primary aim is to introduce a regression algorithm for the inference of a gene network model in *Drosophila melanogaster* blastoderm embryo, referred to as *W*, and gene-specific biases, that describe the observed discrete gene dynamics, and hence satisfy the following linear system:

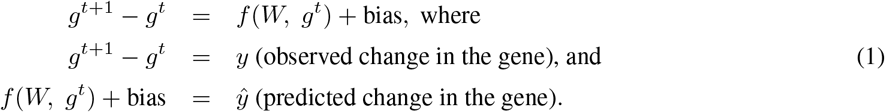

*f* (·) is some function of the gene expressions. *y* is a 30390-by-99 matrix with five vertically stacked sub-matrices of gene expression differences between two consecutive timepoints. To solve the system, we developed a robust and stable variation of least absolute deviation (LAD) regression that reduces the error in prediction during its iterative process. The method will be referred to as erf-weighted LAD from here on. The details of the method are elaborated in the next subsection, Section 2.3.1.

In order to evaluate the performance of the model inferred with erf-weighted LAD, we used two other methods to infer models, least squares regression (with or without the *L*_2_ regularization, depending on the sample size) and neural networks, and compared the predictive power of the models. For least squares and ridge regressions, we implemented the linear_model module from Scikit-learn [14]. To build a predictive model using neural networks we trained and validated multilayer networks of neurons using a widely used API, TensorFlow.Keras [1].

#### 2.3.1 erf-weighted LAD

We want to use LAD because we want to suppress the effects of outliers in our analysis. Usual LAD regression suffers from the well-known instability of minimizing a cost that is a sum of absolute values. This instability is related, obviously, to the susceptibility of the median to noise in the data. It is this instability that we will try to eliminate in our modification of LAD.

Recall that one implementation of LAD regression is as an iterative weighted least squares, as follows. We want to minimize

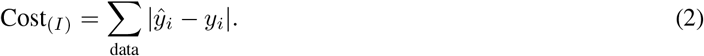

To make this computation amenable to simple linear algebra, we write this cost function as a weighted minimization within an iteration:

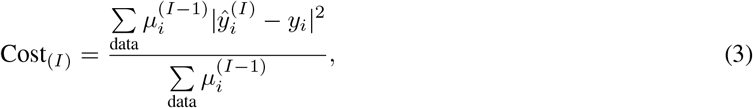

with 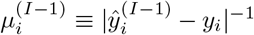. This defines the cost at iteration *I* that we minimize using weighted least squares. As is evident, the weights *μ_i_* can become singular if model predictions approach observed values, so the susceptibility of LAD to noise is not ameliorated by this iterative formulation.

Consider an ensemble of sets of measurements *y_i_*. Model predictions will not vary because the model is independent of measurement noise. However, the discrepancies between model and measurements will be distributed with ensembles of random variables drawn from the same distribution. If we try to define more robust weights that apply to all ensembles, we end up evaluating an expectation value for the discrepancies over this distribution. This suggests that we could factorize and write

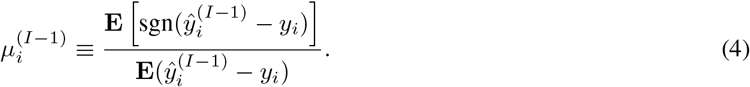

These expectations are over our imagined ensemble of sets of measurements. Evaluated over a normal distribution with mean equal to the actual observed discrepancy, we get a formula for the mean weight involving the error function Erf:

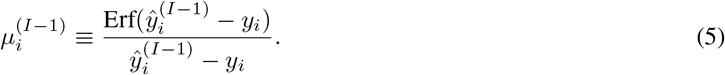

Weights defined thus are positive, non-zero and tend to the constant 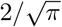 as the model prediction approaches the data measurement. Indeed, for small discrepancies, this weighted regression behaves exactly as a least squares regression would but at large discrepancies it behaves as an LAD regression should.

#### 2.3.2 Regression and 10-fold cross-validation

Following a typical machine learning scheme, a portion of samples were set aside as a test set, which was used to evaluate the performance of the inferred models. We randomly selected 608 nuclei and set aside all of their timepoint measurements as a test set. This amounts to 10% of the total data set. For consistency and fair evaluation, this test set remained fixed and universal across all inferences that we have performed throughout the study.

When inferring a model with erf-weighted LAD regression, we performed 10-fold cross-validation to avoid over-fitting. The training sets were used to train a model, while the validation sets played the role of a stopping criterion. For each combination of train/validation, we used the gene profiles of randomly selected 4923 nuclei (6078 nuclei × 0.9 × 0.9) as a training set and the rest as a validation set. As a result, we obtained 10 different models, i.e., 10 sets of *W*’s and biases. We then averaged the *W*’s and biases as the final step of the inference and used them to make gene profile predictions on the test set. Fig. 3 illustrates the process. The inference algorithm does not have any constraints on the predictions. Hence, it is possible to obtain negative predictions. Since the predictions represent gene expressions, we rectified negative predictions to 0’s. As a way of quantifying the prediction power of the models inferred with erf-weighted LAD, we defined the fractional error of a gene within the test set as follows:

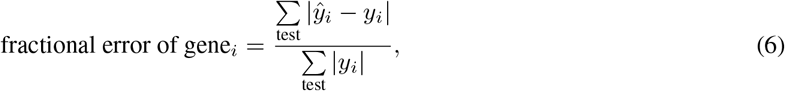

where *ŷ_i_* and *y_i_* represent the predictions and the observations of gene *i*, respectively.

**Figure 3:**
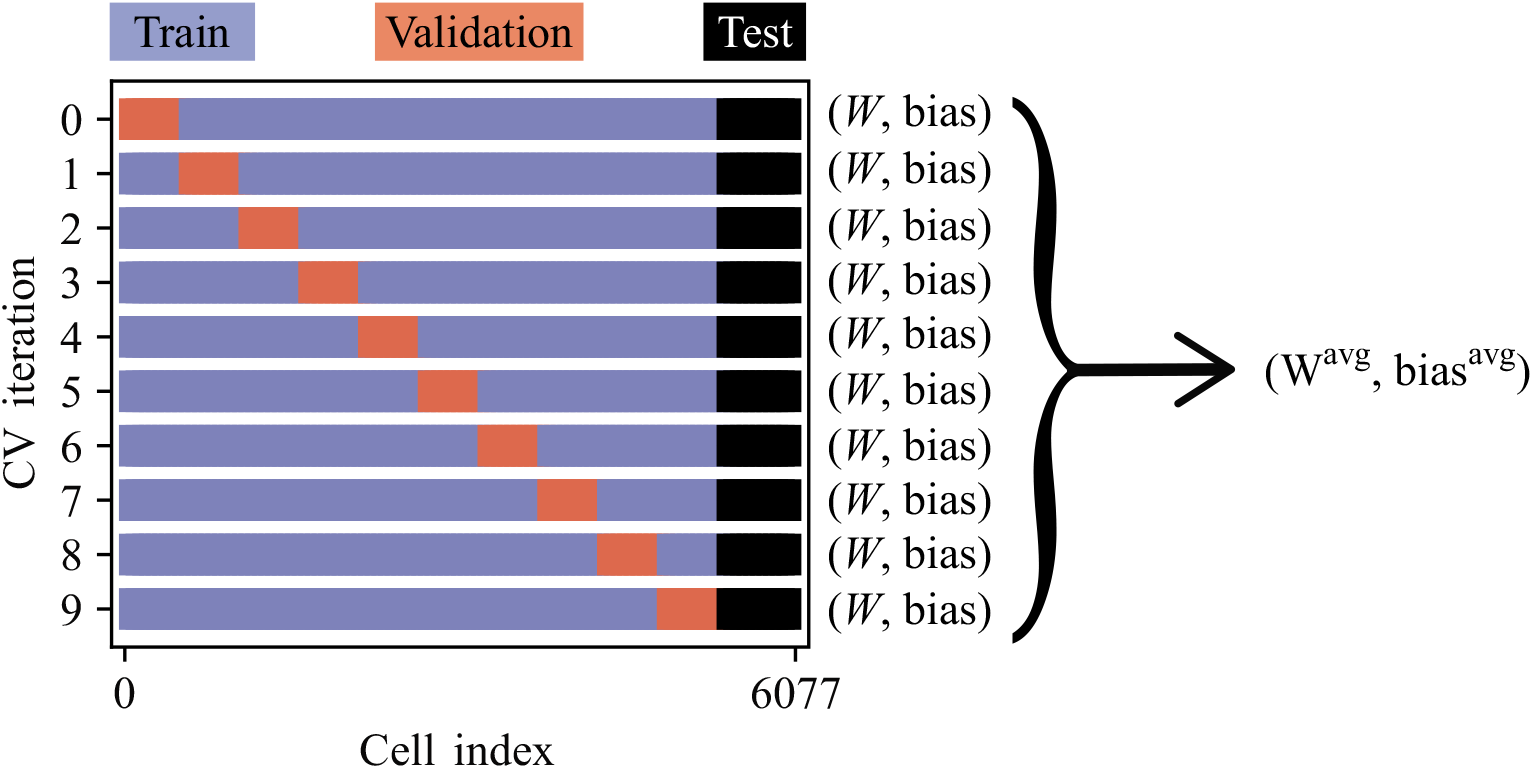
10-fold cross-validation for erf-weighted LAD and least squares regressions. For every inference instance, we held out the same set of nuclei, which was later used to evaluate the model accuracy. The remaining nuclei were shuffled and split into 10 groups. At each cross-validation iteration, a model was trained on 9 groups, while the remaining group was a validation set. Each group took turns as a validation set. 10 iterations of cross-validation resulted in 10 *W*’s and bias’, which were averaged into *W*^avg^ and bias^avg^.

We took the same 10-fold cross-validation approach when inferring a model with least squares regression. However, since least squares regression minimizes a different cost function than erf-weighted LAD, we rescaled the fractional errors of the gene profiles predicted with the models inferred with least squares to be:

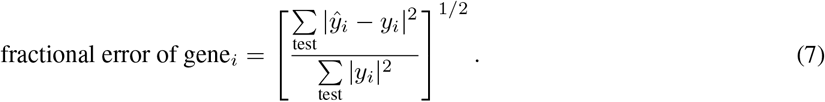

Moreover, least squares regression is not an iterative algorithm, and uses an analytic solution. Thus, it was not necessary to have a validation set when inferring a model with least squares regression. However, in order to conduct a fair comparison between the methods, the training sets that were used to infer models with erf-weighted LAD were also used for least squares regression.

When training neural networks, they were optimized to minimize the mean absolute error, using the *MeanAbsoluteError* class. The training also maintained the 10-fold cross-validation technique. This yielded 10 trained neural networks. However, there is no established way to obtain an average representation from a set of trained neural networks. Instead, for each round of train/validation, we saved the best performing model by utilizing the *EarlyStopping* and the *ModelCheckpoint* callback features in Keras, then made predictions on the test set with the best model. As the final step, we averaged the set of 10 predictions to obtain a mean predicted *ŷ*.

### 2.4 Repository

Complete source code with documentation is available at https://github.com/nihcompmed/erf-LAD.

## 3 Results

The primary aim of this study is to infer a mechanistic model that can describe the observed discrete gene dynamics in *Drosophila melanogaster* blastoderm embryos and, ultimately, make predictions on the effects of perturbed gene regulatory networks. In this section, we compared the accuracy of the predictions made from the model inferred with erf-weighted LAD and those of the predictions from other algorithms. To be expansive and stringent on the model evaluation, we varied the size of training samples from 10% to 80% of the total sample size, and analyzed the subsequent predictions. As for the other algorithms, we used least squares regression, with and without regularization, depending on the training sample size, and neural networks. Finally, we made model predictions that are experimentally verifiable.

### 3.1 Inference: Linear predictors vs. linear and quadratic predictors

Using erf-weighted LAD, we inferred gene networks in *Drosophila melanogaster* blastoderm embryo that would predict gene profiles at one time step ahead, given two different tiers of the current time step’s gene profiles as predictors. Firstly, the linear profiles of 99 genes were used as predictors. Then, we added 4851 quadratic products of all 99 genes as predictors, totaling 4950 predictors. An advantage of using the quadratic profiles is that they provide some information about collective effects of two genes on a third gene.

With each inferred gene network model, we predicted gene profiles at *t* = 1, 2, 3, 4, and 5 in the test set, given the corresponding set of predictors. The negative predictions were rectified to 0’s. When we compared the negative predictions from both models, the model inferred with linear and quadratic predictors had a smaller median in magnitude. The difference in the medians of the negative predictions was statistically significant. The figures showing the distributions of the negative predictions are included in the Supporting Materials A, Fig. S2.

We evaluated the performance of the inferred models by computing and comparing fractional errors of the predicted gene profiles, Eq. 6. Fig. 4 shows the fractional errors of the predictions from the gene network models. The fractional errors computed from the model inferred with the linear and quadratic gene profiles are sorted from the largest to the smallest. The resulting gene index was applied to plot the errors from the model inferred with linear gene profiles. As shown in the figure, the model inferred from the linear profiles and the quadratic products from all 99 genes perform better, yielding smaller fractional errors than the other model that was inferred with the linear gene profiles as predictors alone. Therefore, all the simulations and predictions presented in this paper are based on the combination of linear and quadratic gene profiles as predictors.

**Figure 4:**
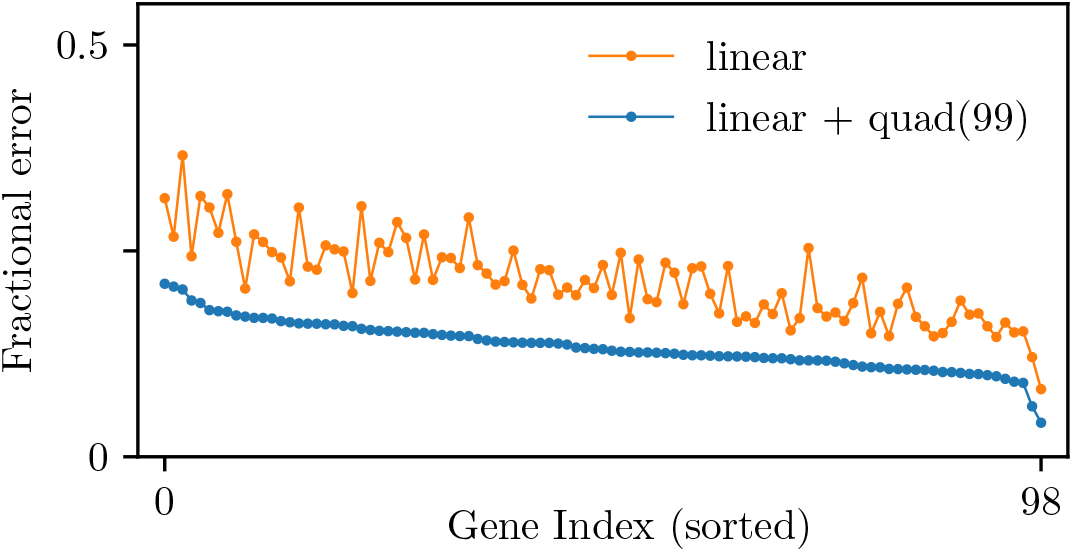
Fractional errors of the predicted gene fluorescence intensity. Using different tiers of the gene profiles specified in the legend at *t* = 0, 1, 2, 3, and 4 as predictors and the linear gene profiles at *t* = 1, 2, 3, 4, and 5 as targets, we inferred the gene network model W and bias using erf-weghted LAD. The models were used to predict the linear gene profiles at *t* = 1, 2, 3, 4, and 5 in the test set. The markers show the fractional error of the predictions for each gene, as defined in Eq. (6).

### 3.2 erf-weghted LAD vs. ridge regression vs. neural networks in the limit of small sample size

A common problem in the field of single-cell analyses is small sample sizes. Thus, it is critical to develop theoretical methods that can extract interpretable information from limited data. With this motivation, we reduced the size of training samples to further test the performance of our inference method in the limit of small sample sizes. The training and validation set of samples was titrated to 10%, 20%, 30%, 40%, 50%, 60%, 70%, and 80% of the total sample size. As we decreased the number of training samples, we noticed that the maximum coefficient of *W* became larger and larger. This was due to a well-known problem where a regression with a smaller sample size is more likely to have a degenerate covariance matrix. In order to avoid this problem, we imposed *L*_2_ regularization on the regression. By adding a small parameter to the diagonals, this regularization allows a singular covariance matrix to be inverted. The regularization rate was fixed at 0.0001 for all titration levels. With the small regularization parameter, we were able to maintain the efficiency of erf-weighted LAD, while constraining the *W* coefficients from uncontrollably becoming large.

For each titration level, 11 different sets of samples were used to infer a set of 11 gene network models. As a metric of the performance, we computed the medians of the *L*_1_ norms of the errors and the total fractional errors in the test set, across the 11 models. Using the same sets of training/validation samples, we inferred another set of models using ridge regression and trained neural networks with one hidden layer and 10000 nodes.

The medians of the ||Error||_1_ and the total fractional errors computed under these methods were compared to those of our inference method. As shown in Fig. 5(A) and 5(C) in the limit of small sample size, erf-weighted LAD performs better than ridge regression and neural networks, and yields gene network models that better predict the gene dynamics, indicated by the smaller medians in both metrics.

**Figure 5:**
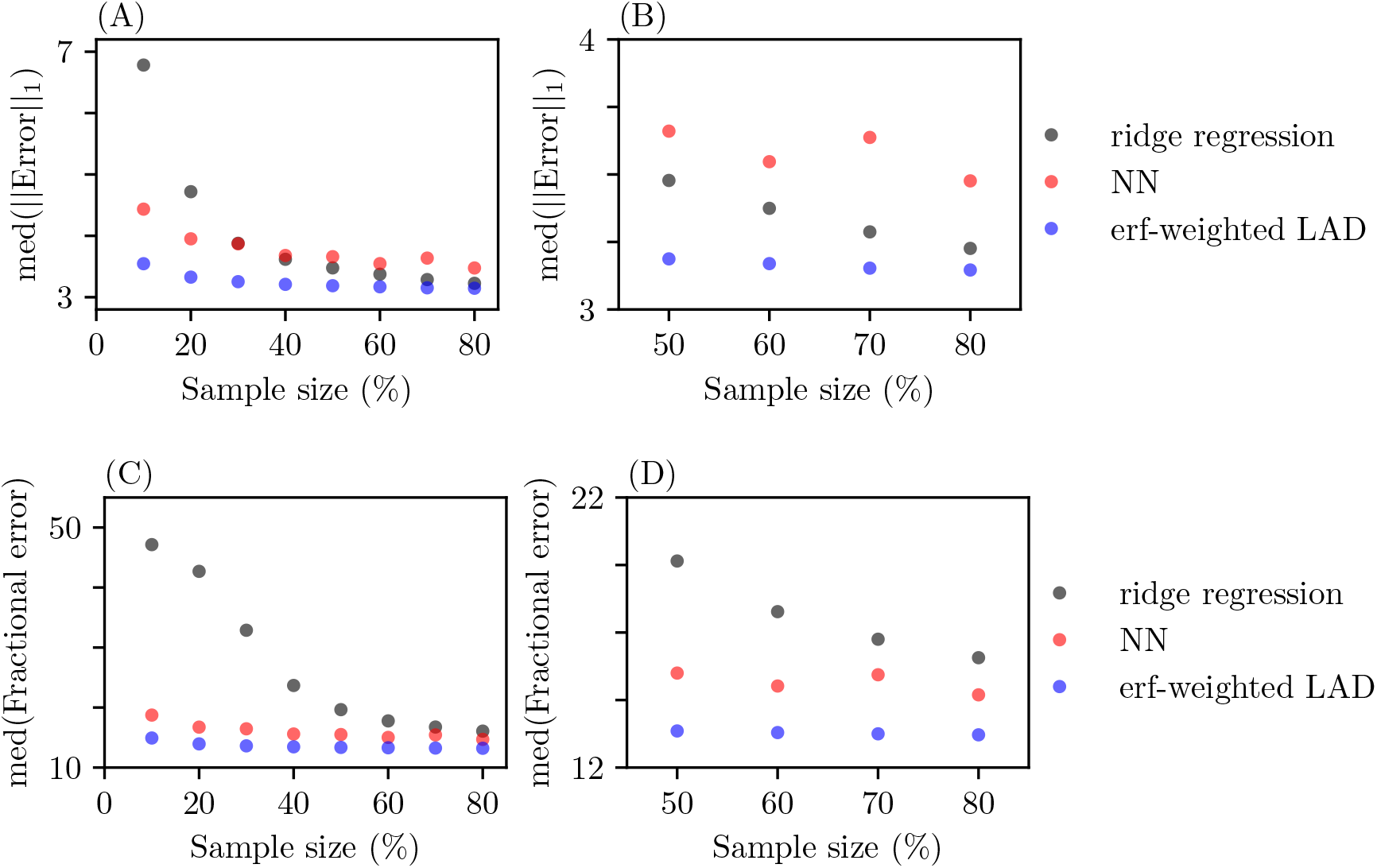
Comparing the predictive power of the neural networks and the gene network models inferred with erf-weghted LAD and ridge regression. The number of the training samples were decreased to 10%, 20%, 30%, 40%, 50%, 60%, 70%, and 80% of the total samples. The panels on the first column are showing all the proportion levels, while the panels on the second column focus on the larger levels with a smaller y-axis window to clearly show the differences. The panels on the top and the bottom rows show the median *L*_1_ norms of the errors and the median fractional errors in the test set, respectively.

As the titration level decreases, the medians increase for all three methods. This is not surprising as the models were trained with smaller sample sizes. Interestingly, the medians from ridge regression increase at a faster rate than those from erf-weighted LAD. Furthermore, at all titration level, erf-weighted LAD performs better than the neural networks, indicated by the smaller medians shown in Fig. 5(A) – (D). It is important to note that the medians from neural networks are not strictly decreasing as the titration level increases whereas the medians from erf-weighted LAD decrease steadily.

We then compared the distributions of prediction errors in the test set, in the limit of small sample sizes. Fig. 6 shows the distributions from erf-weighted LAD and ridge regression under two different metrics. The distributions are plotted on a logarithmic scale for better visualization. Firstly, the *L*_1_ norm of the errors were compared, shown in Fig. 6(A). Comparing two methods under this metric is advantageous for erf-weighted LAD, since it is exactly what the method is trying to minimize. The distributions clearly show a big difference. For more scrutiny, we performed the Mann-Whitney *U* test. With a *p*-value of 0, the test rejects the null hypothesis H_0_: the distributions of *L*_1_ norm of the errors are equal. The median *L*_1_ norm of the errors from erf-weighted LAD is 3.55, while that from the ridge regression is 6.78.

**Figure 6:**
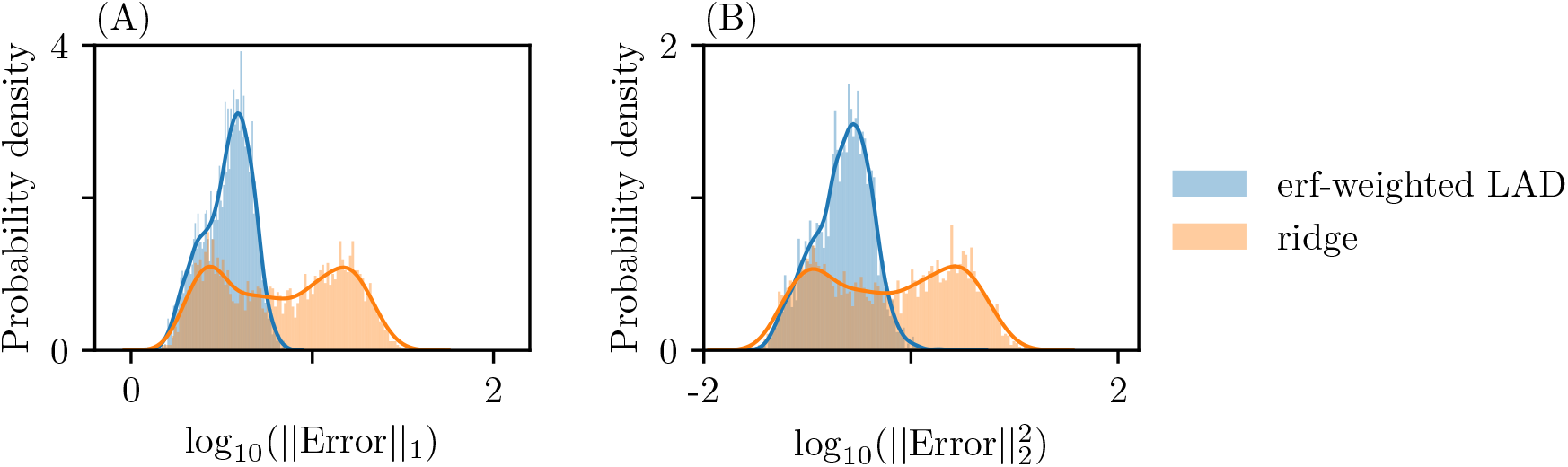
Comparison between the prediction errors of the gene network models. The models inferred from erf-weghted LAD and ridge regression with 10% of the total samples were used to forecast the gene profiles in the test set. The errors between the predictions and the observed gene profiles were measured. Panel (A) shows the distributions of the *L*_1_ norms of the errors, while panel (B) shows the distributions of 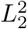 norms. In both panels, the blue and orange distributions are from the erf-weghted LAD and ridge regression predictions, respectively.

Since comparing the *L*_1_ norm of the prediction errors favors erf-weighted LAD, we then compared the 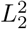 norm of the errors. Under this metric, ridge regression has the upper hand. Again, the distributions show a significant difference; see Fig. 6(B). When we performed the Mann-Whitney *U* test, it rejected the null hypothesis H_0_: the distributions of 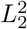 norm of the errors are equal, with a *p*-value of 0. The median 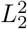 norm of the errors from erf-weighted LAD is 0.23, while that from ridge regression is 0.62.

From Fig.s 5(A) and 5(C), it is clear that the gene network model inferred with erf-weight LAD has better predictive power than the trained neural networks, in the limit of small sample sizes. To further examine the differences, we compared the error distributions of the models that were trained with only 10% of the full data set. Fig. 7 shows the *L*_1_ norm of the prediction errors, on a logarithm scale. Based on the figure, it is clear that the median of the erf-weighted LAD’s error distribution is smaller than that of the neural networks’ error distribution. This is confirmed through the Mann-Whitney *U* test that gave a 0 *p*-value. The median of the neural networks’ error distribution is 4.43.

**Figure 7:**
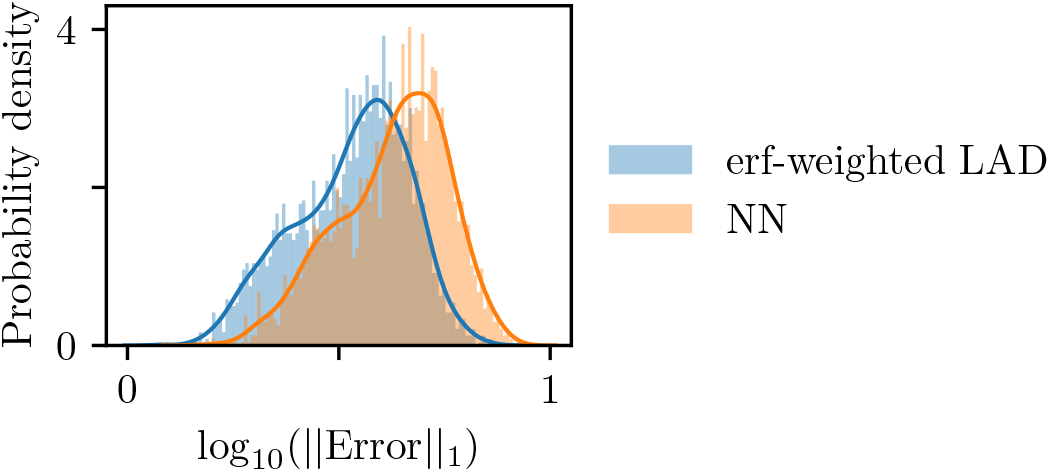
Comparison between the prediction errors of the gene network models. The models inferred from erf-weghted LAD and the neural networks trained with 10% of the total samples were used to forecast the gene profiles in the test set. The errors between the predictions and the observed gene profiles were measured. The distributions of the L1 norms of the errors are shown in a logarithm scale. The blue and orange distributions are from the erf-weghted LAD and the neural networks predictions, respectively.

Overall, we conclude that, in the limit of small samples sizes, the gene network model inferred with erf-weighted LAD and *L*_2_ regularization gives better predictions for the gene dynamics in the majority of nuclei.

### 3.3 erf-weghted LAD vs. neural networks in the limit of full sample size

To further attest to the practicality of erf-weighted LAD, we compared the prediction errors of the model trained with erf-weighted LAD with those of a neural network in the limit of full sample size. We did not impose the *L*_2_ regularization this time, as the prediction power between the model inferred with the regularization and the one inferred without the regularization did not show a significant difference. Fig. 8(A) – (D) show the distributions of the *L*_1_ norms of prediction errors from the trained neural networks with one hidden layer and 50, 100, 1000, and 10000 nodes, respectively, plotted with the error distributions from erf-weighted LAD. From the figures, it is clear that the center of the distribution from erf-weighted LAD is to the left of that of the distribution from the neural networks, in all four cases. This is statistically confirmed through the Mann-Whitney *U* tests with a p-value of 0 in all four comparisons.

**Figure 8:**
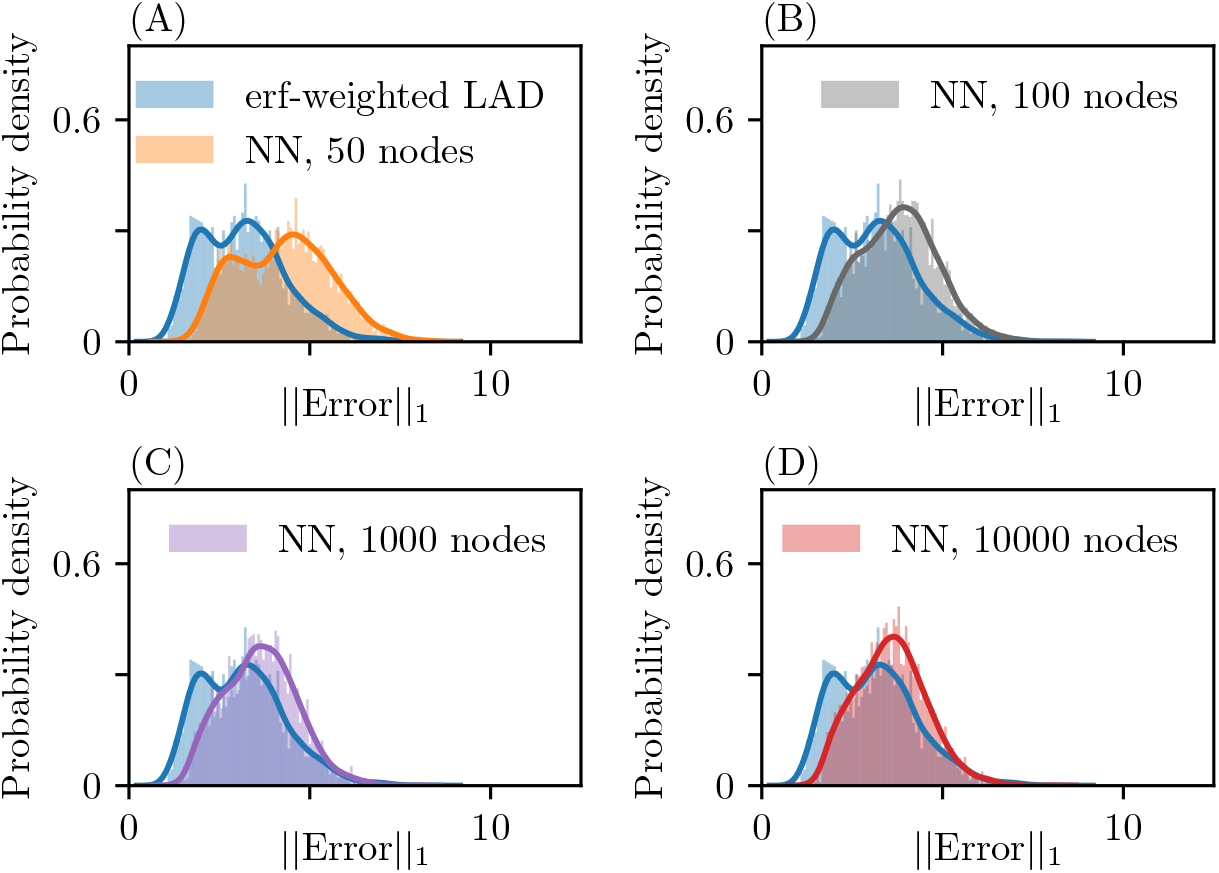
Comparison between the prediction errors of the models inferred from erf-weghted LAD and neural networks (NN) with one hidden layer. The trained neural networks with one hidden layer of varying numbers of nodes were used to predict the gene profiles in the test set. The blue distributions in all panels show the *L*_1_ norms of the prediction errors from the gene network model inferred with erf-weghted LAD. The other distributions in Panels (A) – (D) show the distributions of *L*_1_ norms of the prediction errors from the neural networks with 50, 100, 1000, and 10000 hidden nodes, respectively.

We then trained neural networks with five hidden layers. The purpose of this training was to investigate whether increasing the complexity of neural networks affects prediction power. Using the full sample size, we trained four neural networks with five hidden layers. The number of nodes on the layers of each neural network were 50, 100, 1000, and 10000. We compared the distributions of *L*_1_ norm of prediction errors from the neural networks and that from erf-weighted LAD. The distributions are shown in Fig. 9(A) – (D). We performed the Mann-Whitney *U* test on each pair of distributions and obtained *p*-values of 0 for all pairs. From these results, we concluded that the neural networks, even with added complexity, do not give as accurate predictions as the inferred mechanistic model.

**Figure 9:**
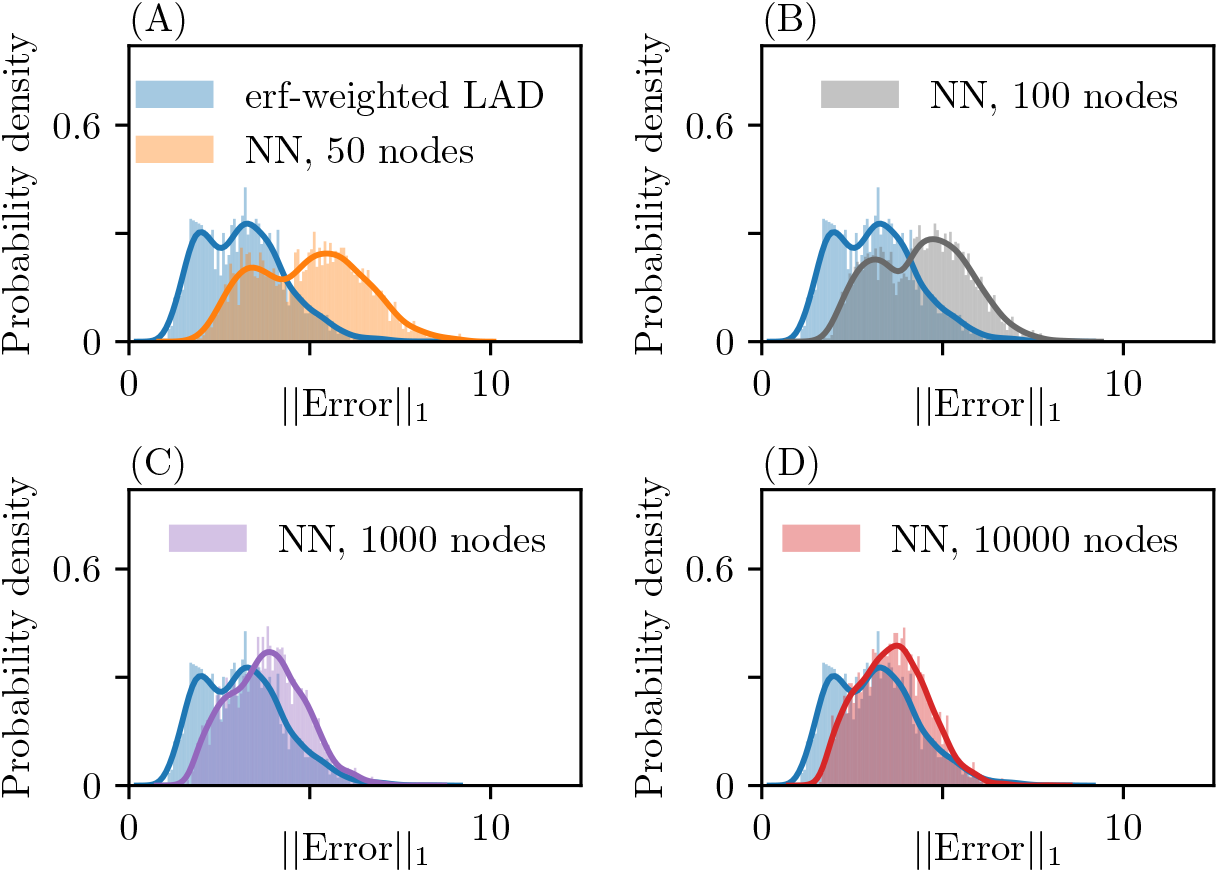
Comparison between the prediction errors of the models inferred from erf-weghted LAD and neural networks with five hidden layers. The trained neural networks with five hidden layers with varying numbers of nodes were used to predict the gene profiles in the test set. The blue distributions in all panels show the *L*_1_ norms of the prediction errors from the gene network model inferred with erf-weghted LAD. The other distributions in Panels (A) – (D) show the distributions of *L*_1_ norms of the prediction errors from the neural networks with 50, 100, 1000, and 10000 hidden nodes, respectively.

### 3.4 Universality of the gene network model

We next addressed whether the locations of the train/validation samples affect the model inference. In other words, if we were to infer a model using a subset of nuclei that are located on the anterior of the embryo, would it have comparable prediction power to a model inferred with the same number of nuclei that are dispersed across the embryo?

Nuclei in close proximity on the embryo exhibit consonant gene dynamics [19]. Accordingly, we studied the variation of the gene dynamics across the whole embryo. Using t-distributed stochastic neighbor embedding (TSNE), a statistical dimension reduction technique for visualizing high-dimensional data [20], we obtained a two-dimensional representation of the gene expressions in 6078 nuclei. Fig. 10 shows the TSNE plot of 6078 nuclei with a color scheme based on their positions on the anterior-posterior axis. The plot suggests that nuclei in close proximity share similar gene expression patterns, and hence, show a limited variation in gene dynamics. Considering this, we postulated that a gene network inferred from closely positioned nuclei is less accurate across the entire test set than one inferred from scattered nuclei. The rationale is simply that by inferring gene network parameters from more variable data, parameters are better tuned to predict in a wider range of initial conditions whereas a network obtained from a less varied set of initial conditions will struggle with predictions for more varied data.

**Figure 10:**
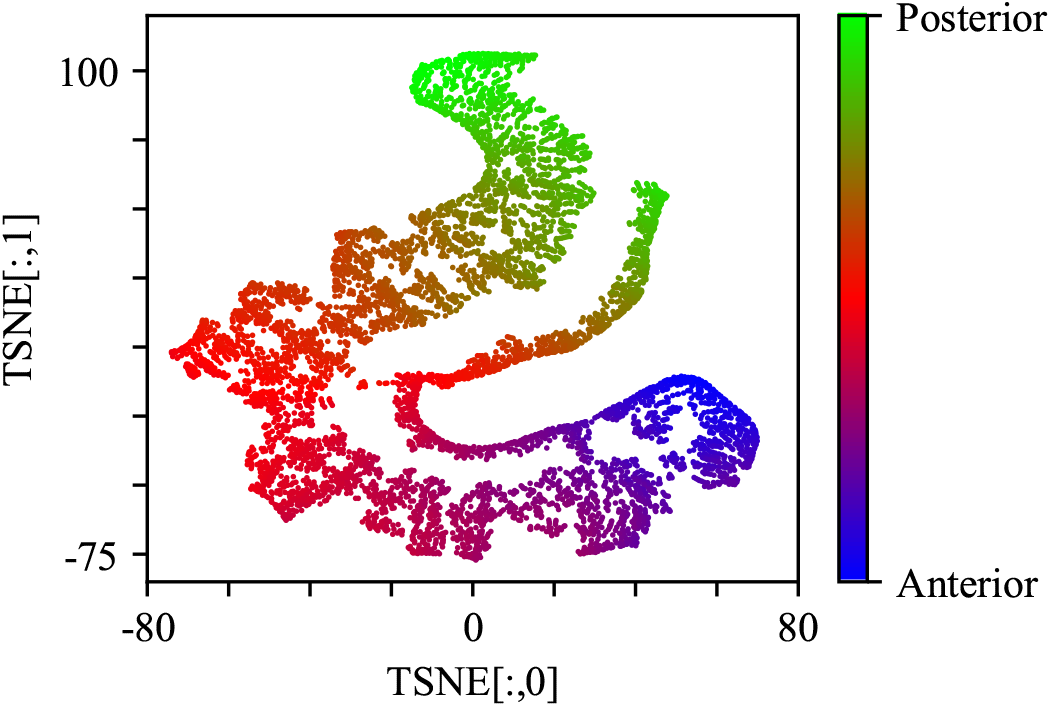
A TSNE plot of the data. The scatter plot shows a two-dimensional representation of the gene expressions in 6078 nuclei.

The same set of nuclei that was previously used as a test set was set aside for later use. Starting from the anterior, the embryo was incrementally partitioned so that each partition contains approximately 30% of the total number of nuclei, see Fig. 11. Using erf-weighed LAD and 10-fold cross-validation, we inferred three different gene network models from three patches on the embryo. As we did in the titration study, we introduced the *L*_2_ regularization to the regression to constraint the magnitudes of *W*. Interestingly, when we used 0.0001 as a regularization parameter, the inferred model had much lower prediction power than the one inferred with randomly selected nuclei. We had to impose a stronger regularization with 1 as the regularization parameter, in order to improve the prediction power. This is because the covariation between nuclei in a specific section of the embryo is much less than the covariation over the whole embryo for the same number of training nuclei.

**Figure 11:**
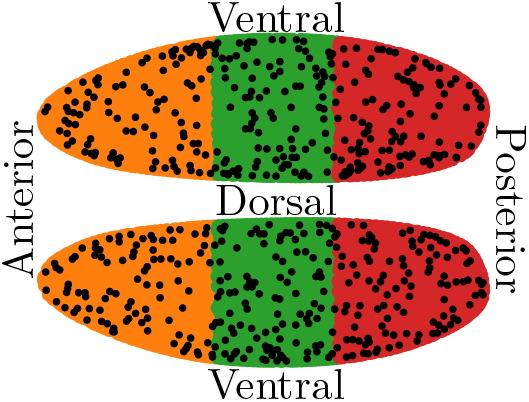
Sectioned embryo. The number of nuclei on each section is approximately 30% of the total number of nuclei. The test set of nuclei are marked in black.

The inferred gene network models were used to predict the gene dynamics in the test set, and the accuracies of the models were evaluated with the fractional errors defined in Eq. 6 and the *L*_1_ norm of the errors. Fig. 12 shows the fractional errors, and Fig. 13 shows the error distributions. As a reference point, the figures also include the errors from 11 different models inferred at 30% titration level, where the training/validation samples were randomly selected, previously presented in Section 3.2. The gene indexes were rearranged so that the errors from the models inferred with random selections of nuclei were sorted by their magnitude. This sorting system is used throughout the section. The models inferred with nuclei in patches of the embryo did not fare as well on predicting gene dynamics compared to the ones inferred with randomly scattered nuclei. This implies that a model that governs the gene dynamics in a cluster of nuclei is only partially shared between different locations in the embryo, at least when only this small subset of 99 gene products is used, and may not be as accurate in predicting gene dynamics in nuclei located in other parts of the embryo.

**Figure 12:**
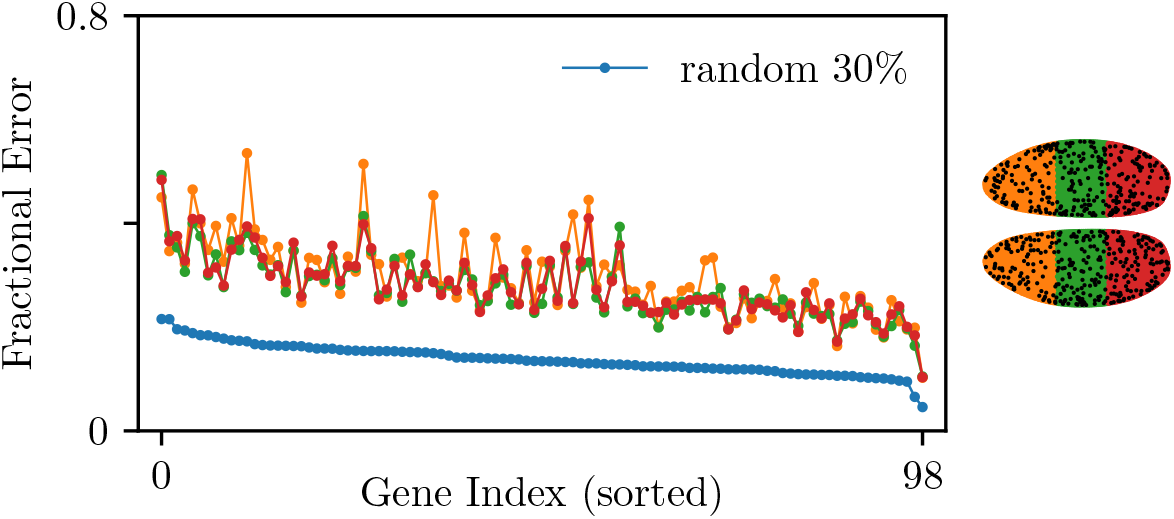
Fractional errors of the gene predictions from the gene network models inferred with different sections of the embryo. Each line represents the fractional errors of the predictions from each inferred gene network model. The line colors are matched to the color scheme of the embryo shown on the right. The blue line shows the fractional errors from the gene network models inferred with randomly selected nuclei that amounts to 30% of the total nuclei.

**Figure 13:**
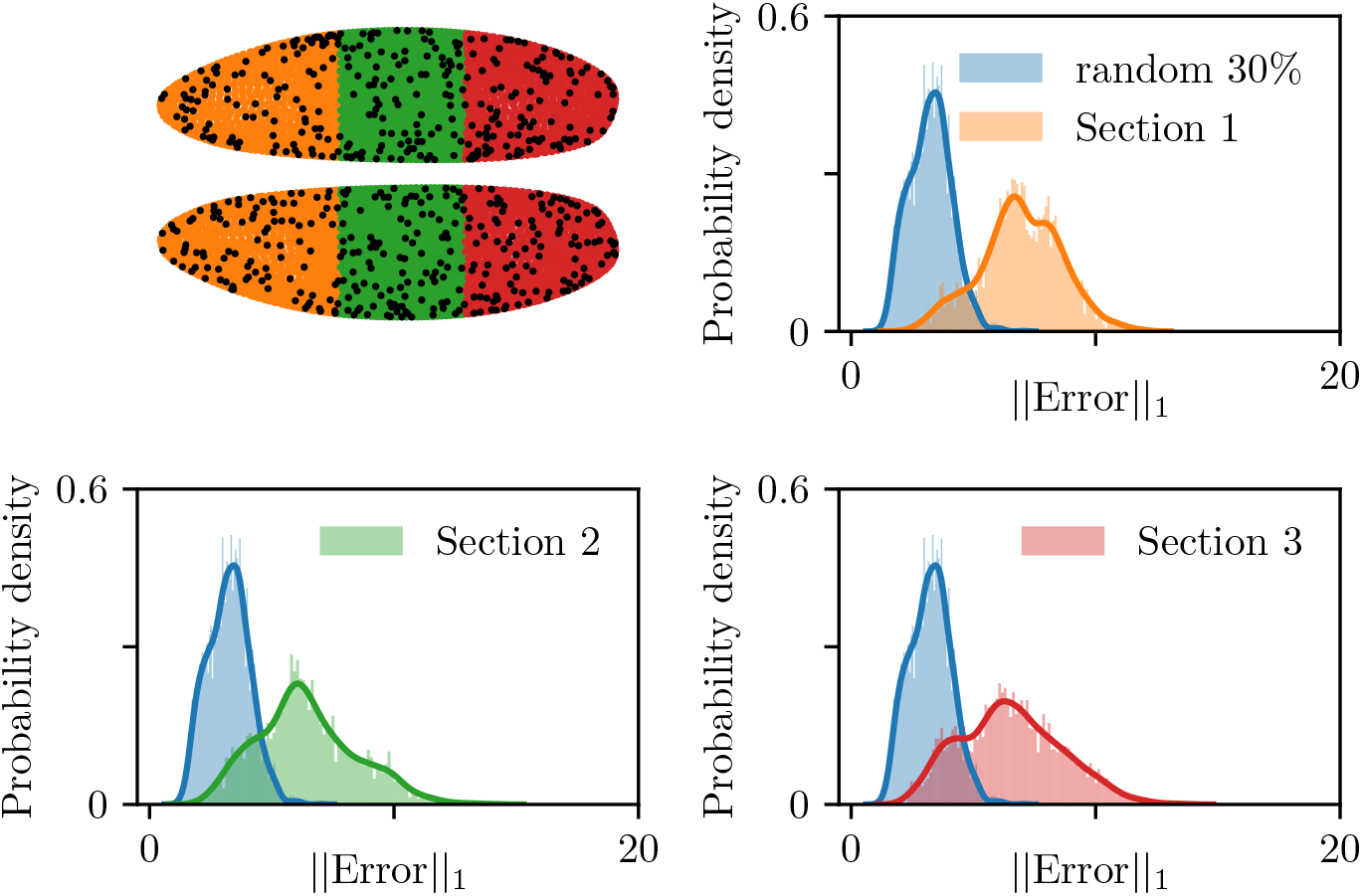
*L*_1_ norm distributions of the errors of the gene predictions from the models inferred with different sections of the embryo. The distributions in blue show the *L*_1_ norm of the predictions errors from the model inferred with random 30% of the total nuclei. The other distribution on each panel shows the *L*_1_ norm of the prediction errors from a model inferred with each partition of the embryo shown on the upper left.

The sections were expanded so that each section contains approximately 45% of the total number of nuclei. The gene network models inferred from the other two sections of the embryo resulted in a similar magnitude of fractional errors, as shown in Fig. 14. However, they were not as small as the fractional errors from the model inferred with randomly selected nuclei. We performed Mann-Whitney *U* to check whether there is any statistically significant difference between two sets of the fractional errors. With a *p*-value of 0.13, there was no significant difference between the sets. Fig. 15 shows the distributions of the *L*_1_ norm of the errors. From the figure, it is clear that the distribution from the model inferred with random nuclei has a smaller median than those from the model inferred with either section of the embryo. When the distributions from the sections of embryo were compared, the median of the anterior’s error distribution was smaller than that of the posterior’s error distribution. The Mann Whitney *U* test gave a *p*-value of 0.016.

**Figure 14:**
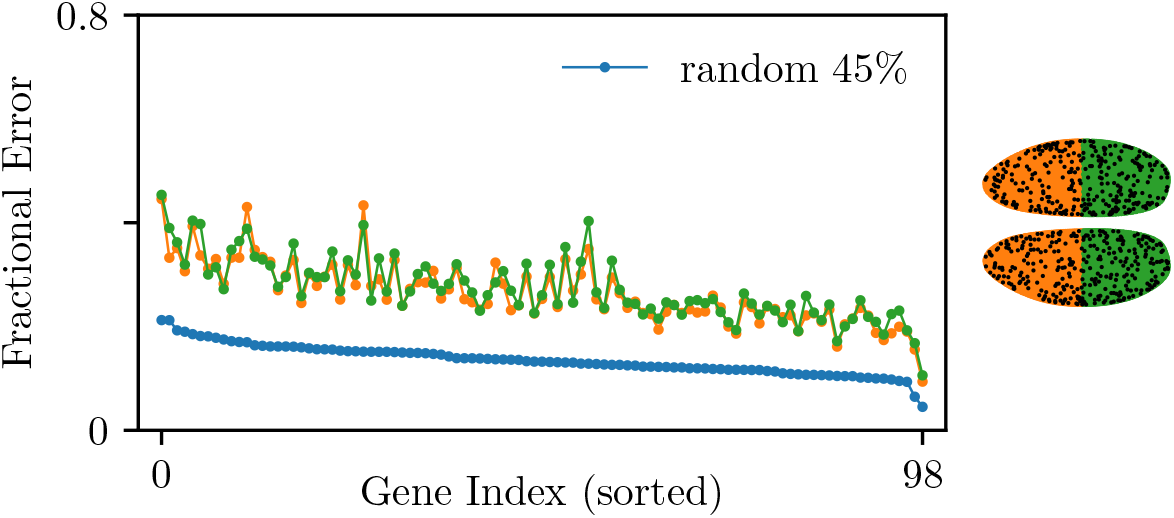
Fractional errors of the gene predictions from the gene network models inferred with different sections of the embryo. Each line shows the fractional errors of the gene expression predictions from each inferred gene network model. The line colors are matched to the color scheme of the embryo on the right. The blue line shows the fractional errors of the predictions from the gene network models inferred with random 45% of the total nuclei.

**Figure 15:**
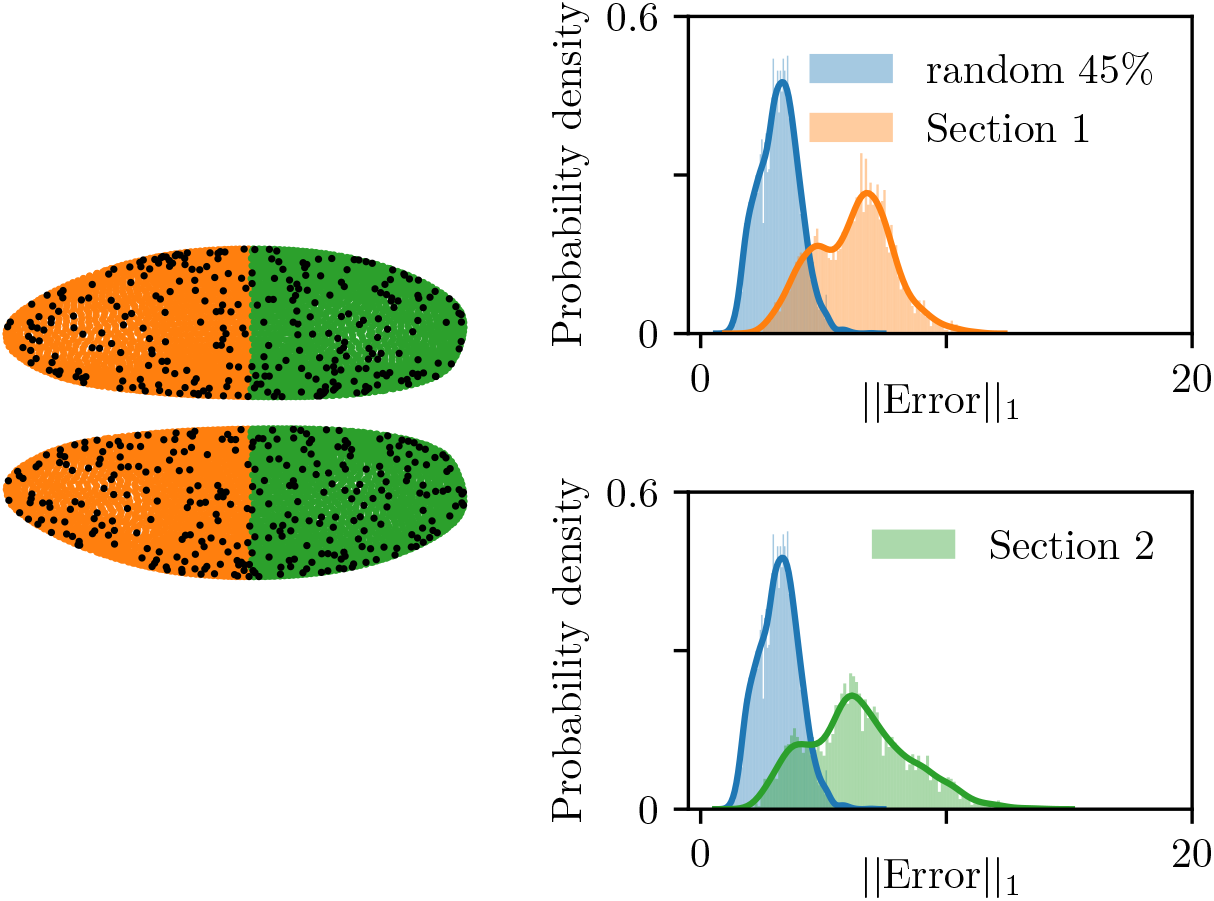
*L*_1_ norm distributions of the errors of the gene predictions from the models inferred with different sections of the embryo. The distributions in blue show the *L*_1_ norm of the predictions errors from the model inferred with random 45% of the total nuclei. The other distribution on each panel shows the *L*_1_ norm of the prediction errors from a model inferred with each partition of the embryo shown on the left.

Lastly, the embryo was sectioned along the dorsal-ventral axis, shown in the right panels of Fig. 16. Following the same schemes as before, the embryo was sectioned into three, then into two sections, so that the sections would include 30% and 45% of the total nuclei, respectively. The panels in Fig.16 indicate that the gene network models inferred with the sections of the embryo do not have as good predictive power as the models inferred with the same number of randomly selected nuclei. The same conclusion can be drawn from the panels of Fig. 17. When we compared the error distributions of the ventral and the dorsal section, there was no significant difference between the medians.

**Figure 16:**
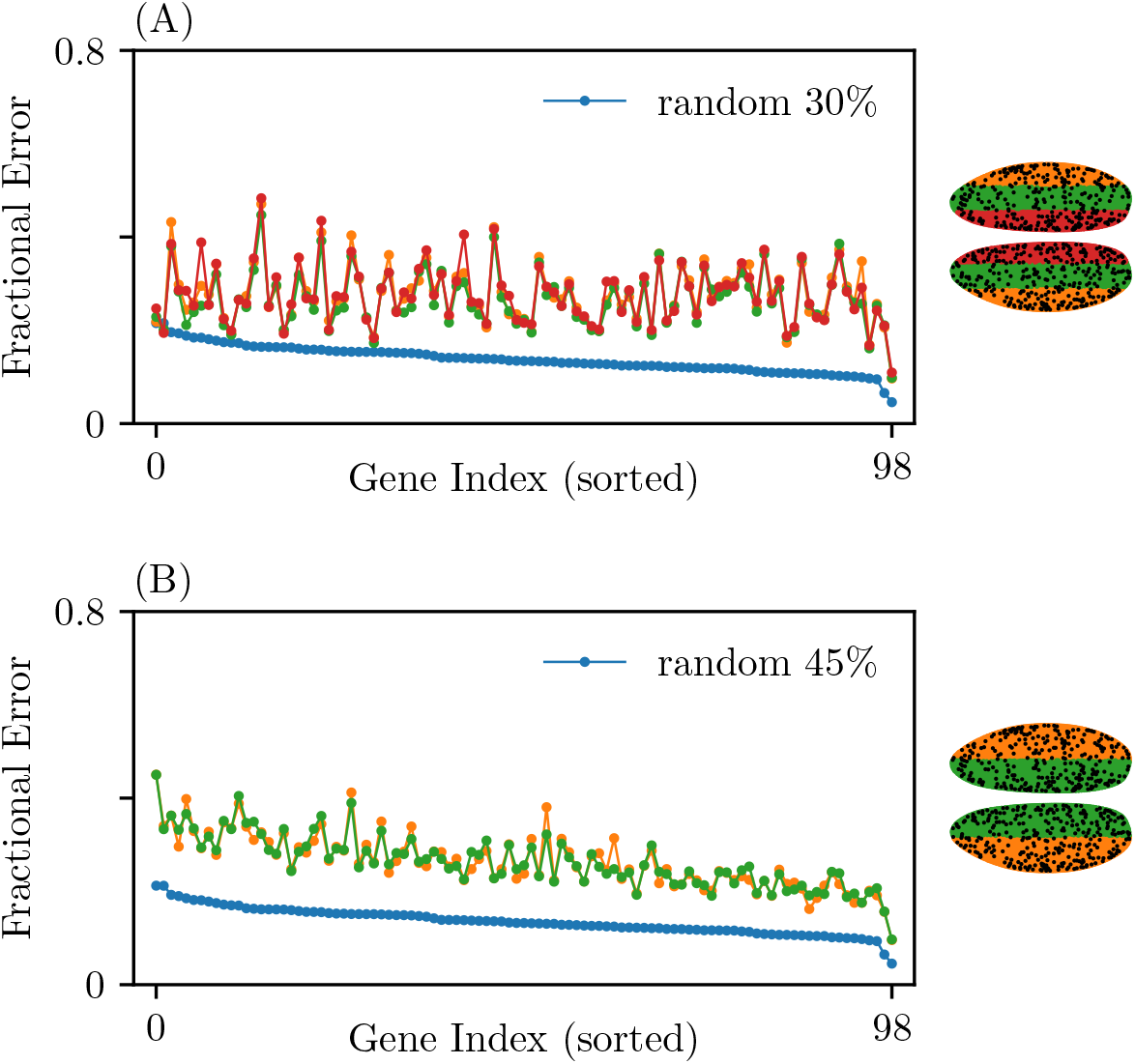
Fractional errors of the gene predictions from the models inferred with different sections of the embryo. The panels show the fractional errors of the gene expression predictions from the inferred gene network model that were trained with (A) 30% and (B) 45% of the total nuclei. The line colors are matched to the color scheme of the embryo on the right. The blue line shows the fractional errors of the predictions from the gene network models inferred with random selections of the total nuclei that amounts to the equal portion of the nuclei for a fair comparison.

**Figure 17:**
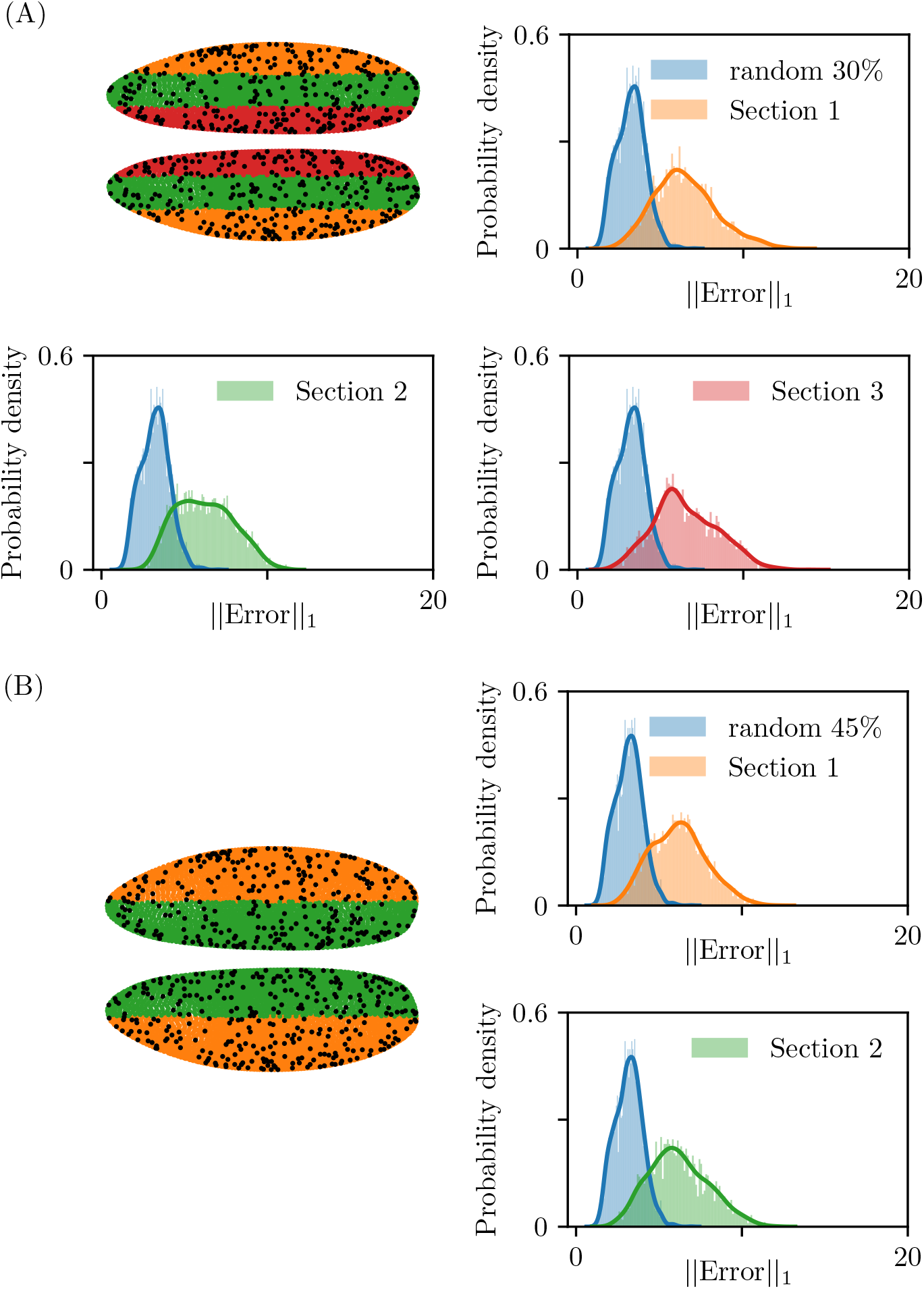
*L*_1_ norm distributions of the errors of the gene predictions from the models inferred with different sections of the embryo. The distributions in blue show the *L*_1_ norm of the predictions errors from the model inferred with random (A) 30% and (B) 45% of the total nuclei. The other distribution on each panel shows the *L*_1_ norm of the prediction errors from a model inferred with each partition of the embryo shown on the left.

To give more insight into the obtained results, we compared prediction errors of the test nuclei that are located within individual sections of the embryo and those of all the test nuclei scattered across the entire embryo. Fig. 18 shows the corresponding comparison. A gene network inferred from a section of embryo predicts the gene dynamics of the test nuclei in that particular section better than it predicts those of all the test nuclei that are scattered across the embryo. The results suggest that, to predict gene dynamics in a group of nuclei that are concentrated on a part of the embryo, it is more efficient to infer a gene network from neighboring nuclei.

**Figure 18:**
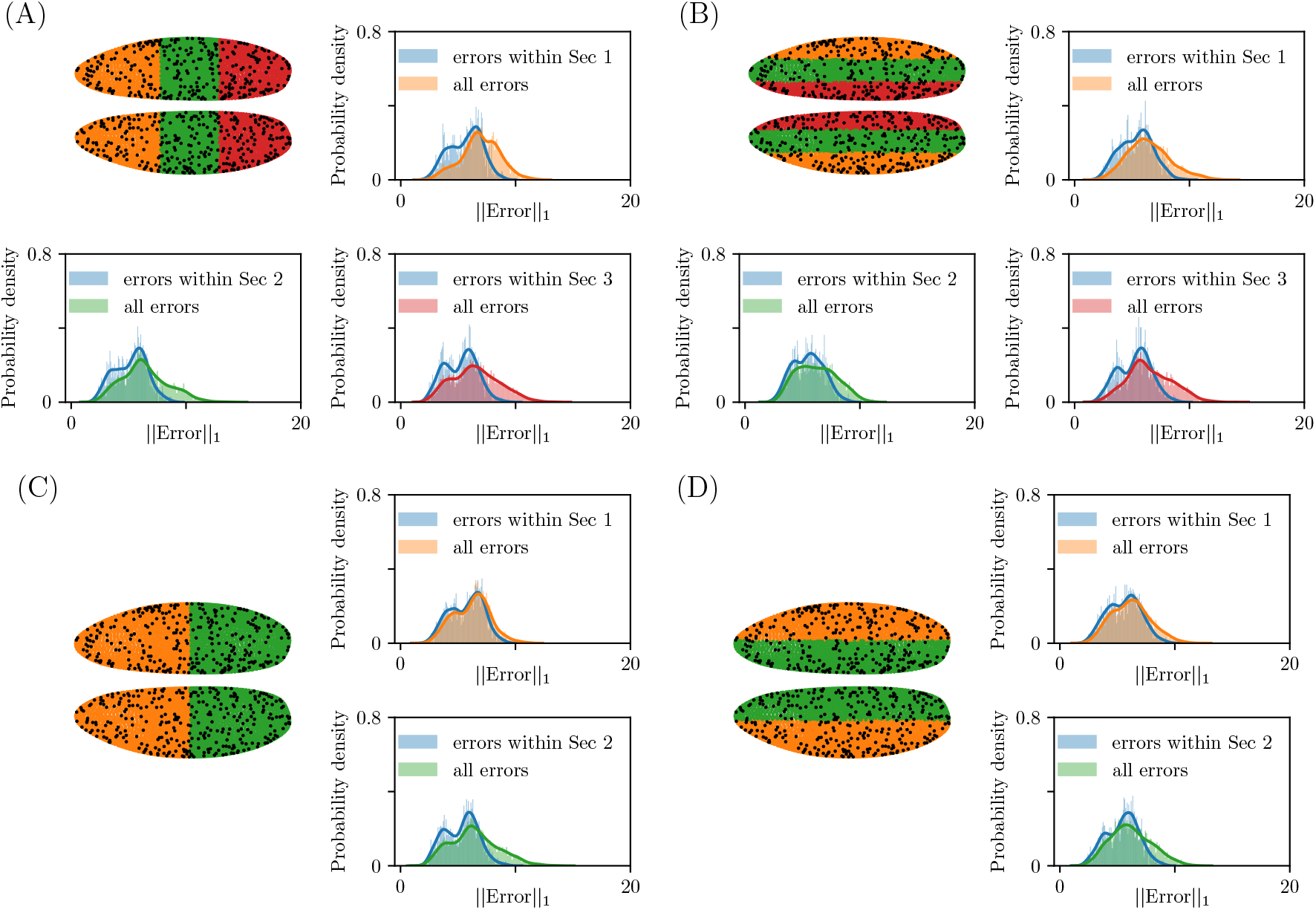
On each panel, orange, green, or red distribution shows *L*_1_ norm of the prediction errors of all the test nuclei scattered across the entire embryo. The predictions are made with models inferred from different sections of embryo. The blue distributions show *L*_1_ norms of the predictions errors of the test nuclei located in that particular section. (A) and (C): 30% and 45%, sectioned along the anterior-posterior axis, (B) and (D): 30% and 45%, sectioned along the dorsal-ventral axis.

### 3.5 Mutant Predictions and Validation

Schroeder et al. [18] studied the regulatory network of *Drosophila* segmentation genes. The study was particularly focused on the pair-rule genes, *hairy (h), even skipped (eve), runt (run), fushi tarazu (ftz), odd skipped (odd), paired (prd), and sloppy paired (slp)*. These genes are expressed in seven evenly spaced stripes across the embryo and play a pivotal role in the segment formation and development of the larva [12]. As part of the study, the effects of pair-rule gene mutants on pair-rule expression were examined. The effects were evaluated at a time when the stripe patterns are clearly visible in the wild type. For the mutants, null conditions were assayed for all genes of interest. As a model validation, we compared their empirical observations to our model predictions on the effects of pair-rule gene mutants. We omitted simulations of *run* mutants, as the data set does not include its expression.

To simulate the mutants, we computed the gene expression time series with the inferred gene network, *W*, and mutant conditions. In order to determine the appropriate time steps for the mutant simulations, we examined the pair-rule gene expression patterns in our data set and concluded that, at *t* = 2, the strip patterns are comparable to those reported in Schroeder et al. [18]. It should be noted that the stripe patterns of *prd* and *slp1* are not as prominent as the others. We tried to accommodate this by simulating the gene expressions by another time step forward. As a result, the stripe patterns in the observed and the simulated wild types were comparable to that of Schroeder et al. [18]. However, the mutant simulations were somewhat erratic and, therefore, inadequate to draw consistent comparisons.

A genetic null allele can be an RNA null, one that produces no RNA transcript and/or a protein null, which produces no protein. For our mutant simulations, we assume that a null allele is both an RNA null and a protein null. The simulations were started from modified initial conditions where the mutated gene expressions were set to 0. Then, we modified the inferred gene regulatory network, *W*, in such a way where the weights of the edges directed towards the mutated gene and the edges branching out from the mutated gene, are set to 0. It is important to note that the edge weights from the quadratic terms that involve the mutated gene are also set to 0. These conditions ensure that, at every forward time simulation, the mutated gene measurements stay at 0. Fig. 19 shows the mutant simulations along with observed stripe patterns (first row) and wild type simulations (second row) for comparison. The list below gives a point-by-point comparison between the results of Schroeder et al., in italic, and our model prediction. For a clear interpretation, the stripes are numbered 1 to 7, counting from anterior to posterior.

• *The mutants exhibit irregularities in the intensity, width, and spacing of pair-rule gene stripes.*
→ Partial agreement. Generally, the simulated mutants showed changes in stripe intensity, but not in the width or spacing.
• *In h mutants, stripe intensity and width were primarily affected.*
→ Partial agreement. In simulated *h* mutant (third row), the stripe intensity of all the other pair-rule genes was affected. The intensity of *eve* and *prd* stripes was weakened, while *ftz* and *odd* showed an increase in their stripe intensity.
• *In eve mutants, all genes showed impaired stripe spacing and intensity.*
→ Agreement. In simulated *eve* mutant, the stripe spacing and intensity seem to be impaired in all the other pair-rule genes, when compared with the simulated wild type.
• In *eve* mutants, stripe 1 and 2 of *slp1* are fused. As a result, stripe 1 of *ftz and odd*, and stripe 2 of *h* were repressed.
→ Partial agreement. In simulated *eve* mutant, stripe 1 expanded towards stripe 2. Stripe 1 of *ftz* was impaired, but the stripes of *odd* and *h* were unaffected. We suspect that this is because, in our data, the position of *slp1* stripe 1 does not overlap with the positions of *odd* stripe 1 and *h* stripe 2.
• *In ftz mutants, odd stripes showed a weakening and loss of regularity.*
→ Agreement. In simulated *ftz* mutant, *odd* stripes were severely impaired.
• *In odd mutants, the width of eve stripes was increased.*
→ Partial agreement. In simulated *odd* mutant (sixth row), the stripe intensity of *eve* was decreased, and the stripe boundaries became slightly fuzzy, which could possibly be interpreted as broadening.
• *In prd mutants, the intensity of stripe 1 and 2 of* h *was weakened.*
→ Agreement. When compared with the simulated wild type, stripes 1 and 2 of *h* were noticeably weakened.
• *In slp mutants, strip 1 of other pair-rule genes were shifted anteriorly.*
→ Disagreement. The simulated *slp* mutant did not display any visible changes in the stripe positions.

**Figure 19:**
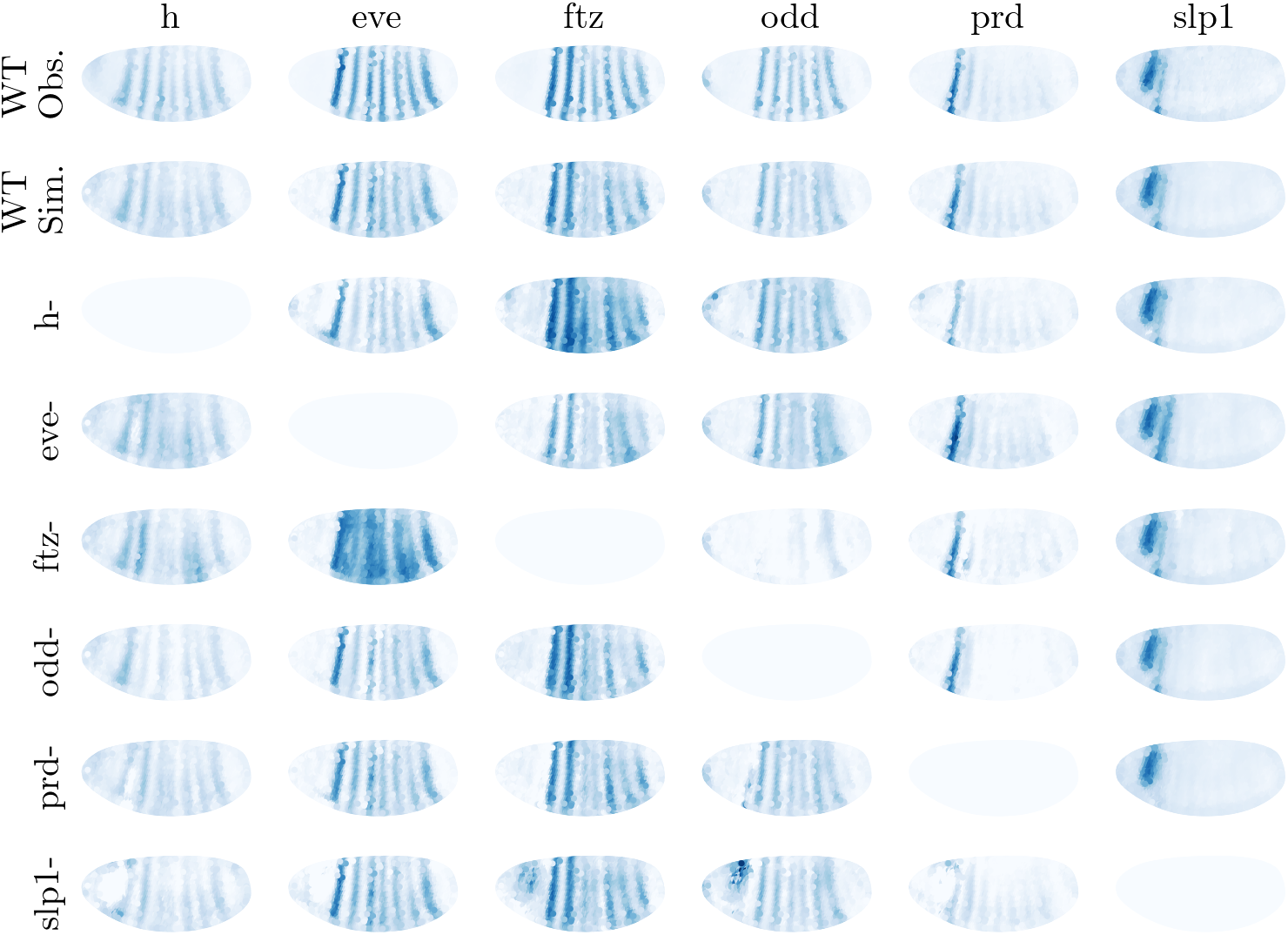
Mutant simulations: pair-rule genes. The model was used to simulate the effects of nullifying each pair-rule gene on transcript patterns of other pair-rule genes at *t* = 2. Genotypes are arranged by row, transcript patterns by column. The first and second rows show the observed and simulated wild type transcript patterns at *t* = 2, respectively. The gene expressions are normalized with respect to the corresponding wild type expressions.

Overall, we found that the association between the null condition of pair-rule genes and the stripe intensities and regularities was reproduced in our mutant simulations. However, the changes in stripe location and/or stripe width observed in the mutants were not captured in the simulations. We suspect that this is because we do not have information on the true initial conditions of the mutants. This is discussed more in detail in Section 4.

Additionally, we made pair-rule gene mutant predictions from the trained neural networks with 1000 nodes on one hidden layer. The initial expressions of a mutated gene were set to 0. Unlike our effective gene network model, where we can suppress pertinent effects by setting corresponding edge weights to 0’s, we cannot emulate this in neural networks. Thus, when simulating mutants with the trained neural networks, the predicted expressions of mutated genes were manually rectified to 0’s, after each time step forward. The results are included in the Supporting Materials A (Fig. S3), which show the averaged expressions predicted from 10 neural networks that we obtained through 10-fold cross-validation.

We then expanded the scope of mutant simulations to the maternal-effect genes, *bicoid (bcd)* and *caudal (cad)*, and the gap genes, *hunchback (hb), giant (gt), Krüppel (Kr), knirps (kni), tailless(tll), and huckebein (hkb)*. The maternal-effect genes are critical for anterior-posterior pattern formation, while the gap genes are involved in the development of the segmented embryo [15]. These segmentation genes are closely related and form a hierarchical regulatory network. The maternal-effect genes control the expressions of the gap genes, which in turn regulate the pair-rule gene expressions. The mutant simulations are shown in Figure 20. There are some genes whose proteins were also measured, *bcd, hb, gt*, and *Kr*. For their mutant simulations, we set the initial expressions of the proteins and the weights of all the edges going in and out of the corresponding linear and quadratic terms to 0’s. Here are some prominent results of the mutant simulations:

- *bcd* mutant: The expression of other genes was disrupted in the anterior region of the embryo.
- *cad* mutant: The stripe intensities of *h, ftz* and *odd* were weakened.
- *hb* mutant: All maternal-effect, gap, and pair-rule genes showed significantly disrupted gene expression patterns.
- *gt* mutant: There was an up-regulation of *hb* in the anterior region of the embryo. The regularity of stripe intensity was significantly impaired in all pair-rule genes.
- *Kr* mutant: All maternal-effect, gap, and pair-rule genes showed disrupted expression patterns in the central region where a *Kr* band is exhibited in the observed and simulated wild type.
- *kni* mutant: *cad, eve, ftz*, and *odd* were notably up-regulated in the anterior.
- *tll* mutant: The posterior of the embryo showed perturbed gene expressions of *cad, hb, gt, kni, h, eve, ftz* and *slp1*.
- *hkb* mutant: All maternal-effect, gap, and pair-rule genes showed up-regulation or down-regulation in the most anterior and posterior regions of the embryo.
- *slp1* mutant: *cad, gt, kni,* and *till* showed changes in the gene expression patterns, mainly in the anterior region.

**Figure 20:**
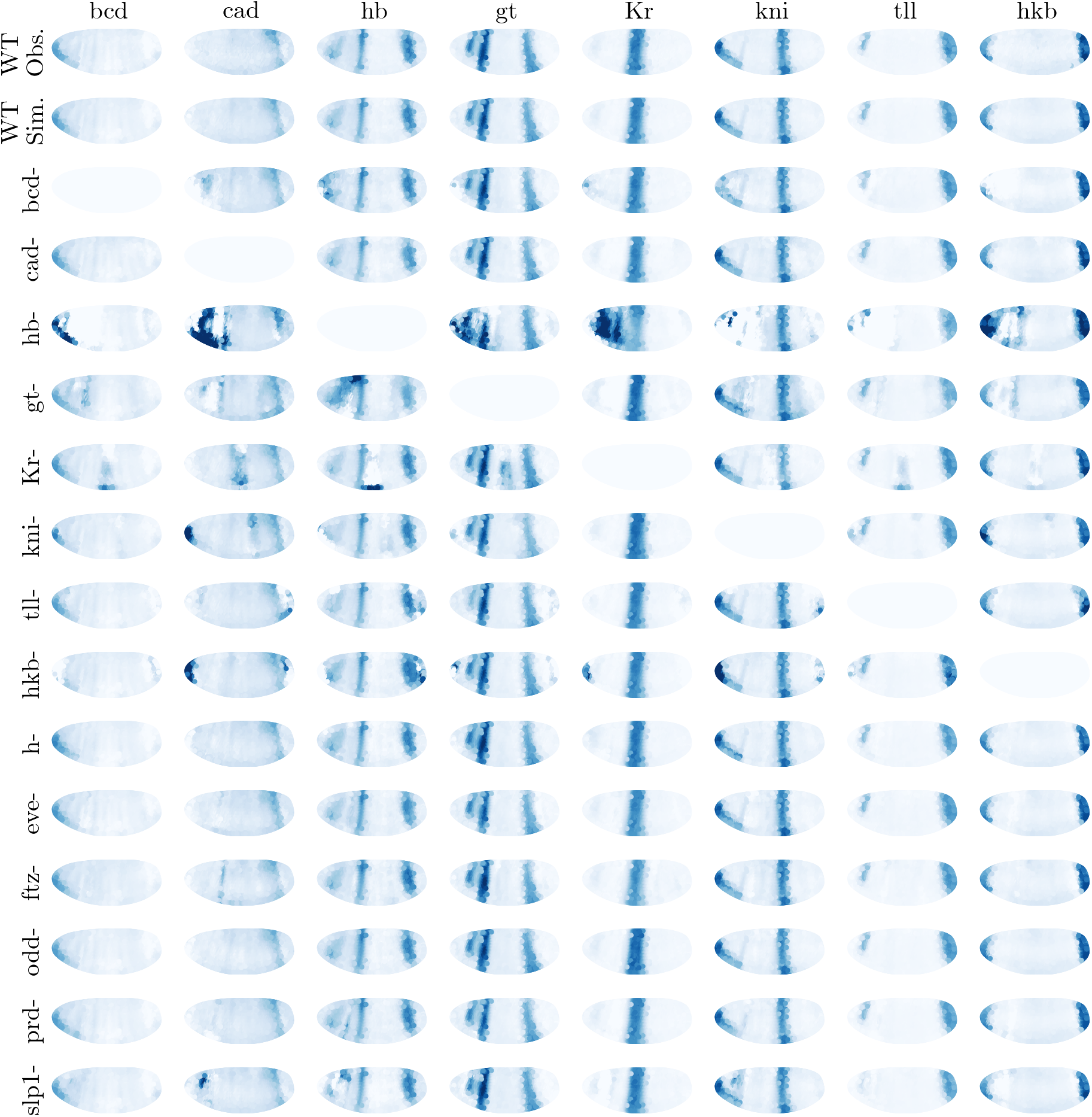
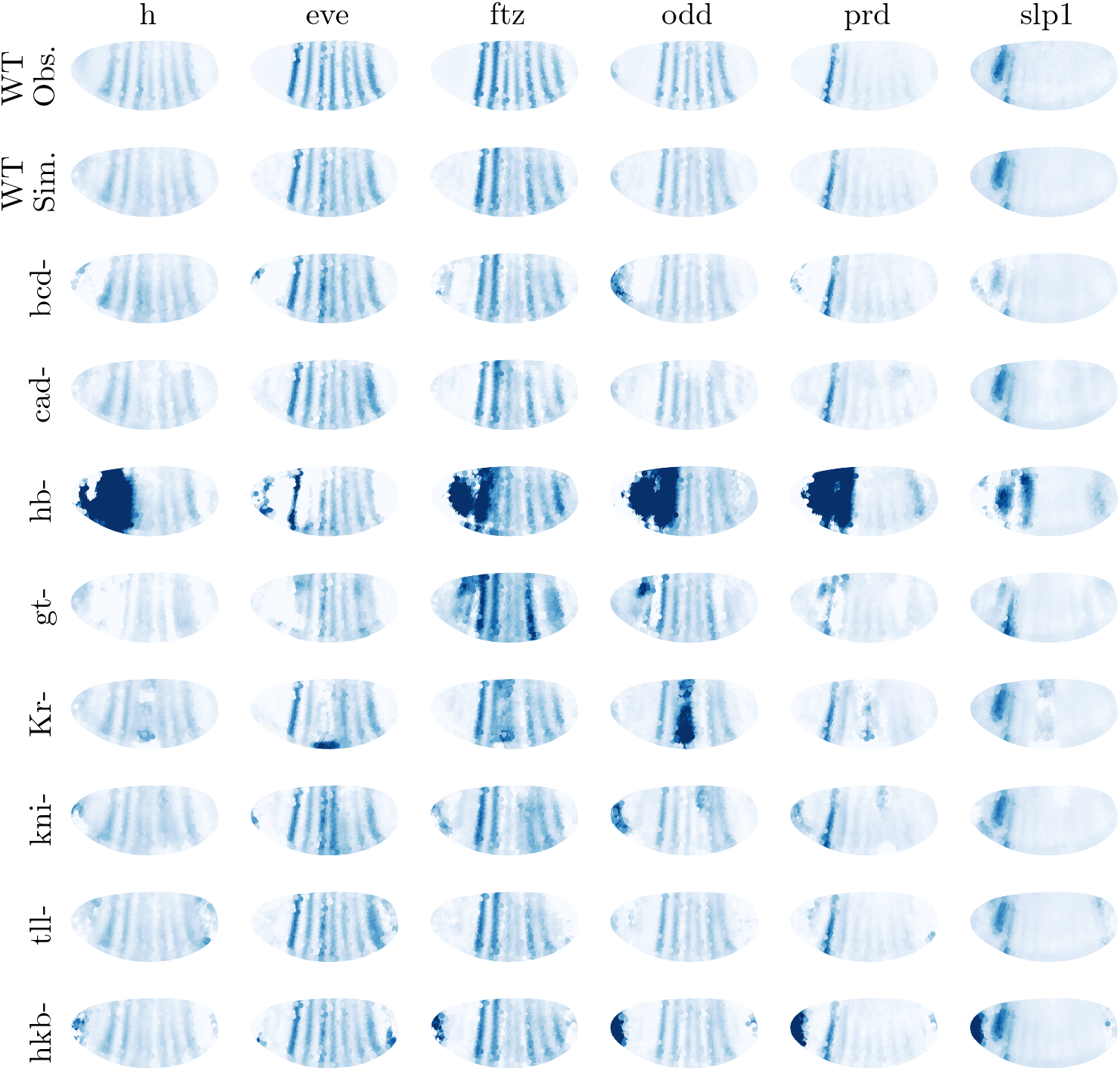
Mutant simulations: maternal genes, gap genes, and pair-rule genes. The model was used to simulate the effects of nullifying the segmentation genes (maternal-effect, gap, and pair-rule) on transcription patterns of the maternal-effect genes and the gap genes. Genotypes are arranged by row, transcript patterns by column. The first and second rows show the observed and simulated wild type transcription patterns at *t* = 2, respectively. The gene expressions are normalized with respect to the corresponding wild type expressions.

Interestingly, in most of the simulated pair-rule gene mutants, there was no major change in the maternal-effect and the gap gene expression patterns. This observation supports the hierarchical regulatory relationship between the maternal-effect genes and the pair-rule genes. The effects of each genetic null allele on the measured protein (*bcd, hb, gt*, and *Kr*) are included in the Supporting Materials A (Fig. S4). For completeness, we also simulated the effects of nullifying individual genes and proteins. The results are included as a separate supporting document, Supporting Materials B.

### 3.6 Visualizing the gene network model and making predictions for the effects of network perturbations

As the mutant predictions showed, the major advantage of mechanistic modeling, here performed with erf-weighted LAD, is that we can access the inferred gene network model, *W*, which is not possible by training black-box neural networks. Fig. 21 shows a partial network representing the interactions between any two genes. The quadratic effects are not shown in the figure. With the network in hand, we can answer the following question: What are the effects of introducing a small perturbation to the network? Some of these predictions are wishful thinking from an experimental point of view as an understanding of transcriptional regulation at the required fine level of control is lacking. Nevertheless, we present them as aspirational.

**Figure 21:**
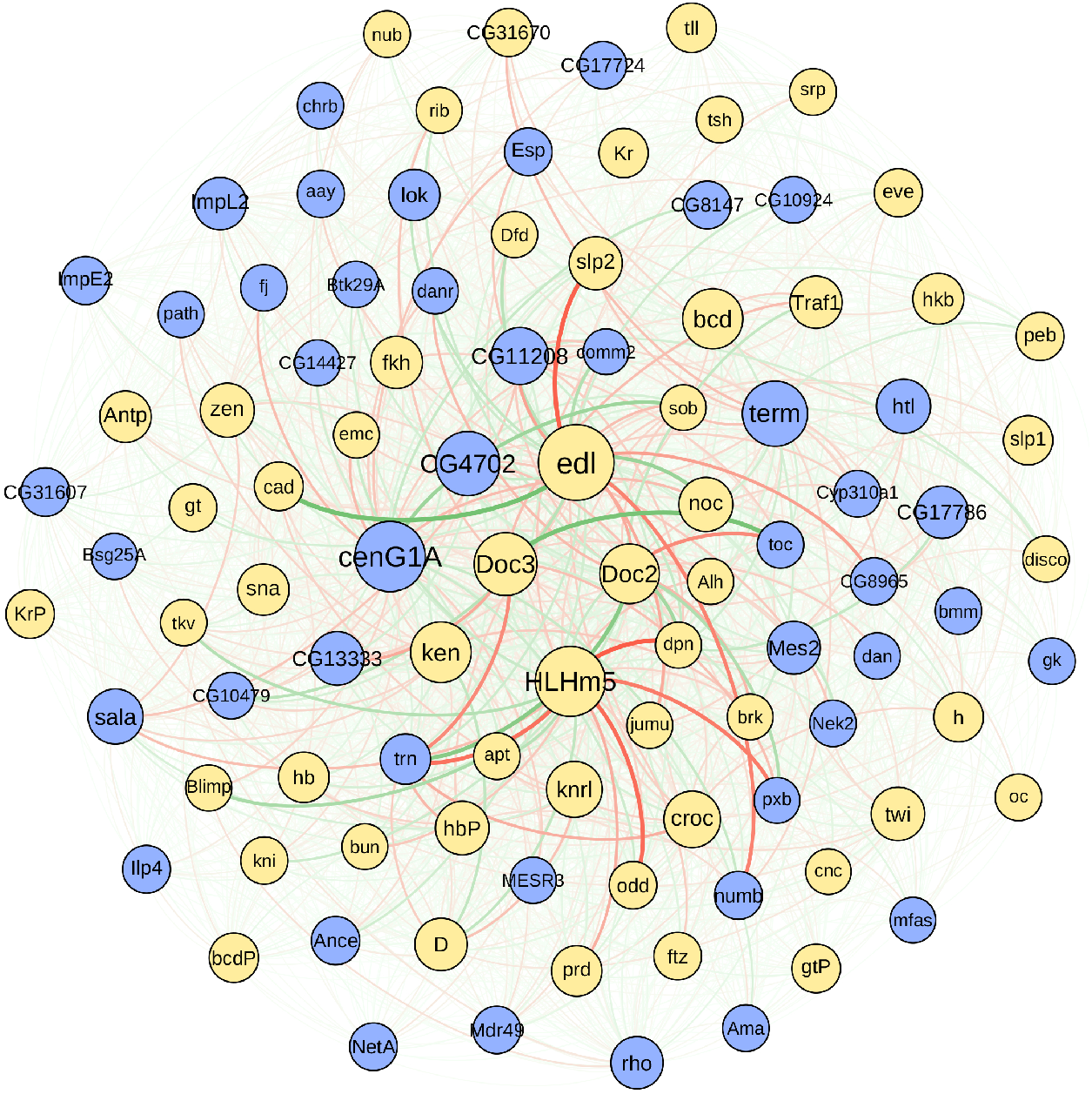
A network graph showing the linear effects between the genes. The genes in the yellow circles are transcription factors and the genes in the blue circles are non-transcription factors. The size of the each circle is proportional to the total effect of the corresponding gene to all the other genes. The pink and green edges represent the positive and negative effects, respectively, while the edge transparency and thickness are proportional to the magnitude of the effect. The directionality of the edges is clockwise.

#### Perturbing the effect of Dorsocross2 (*Doc2*) on tartan (*trn*)

*Doc2* encodes one of the three tissue-specific T-box transcription factors and is crucial for numerous cell activities including completion of differentiation and proliferation arrest [8]. *trn* is another protein coding gene that encodes a transmembrane leucine-rich repeat protein [8]. *Doc2*, located at the center of the gene network in Fig. 21, has a positive effect on *trn*, located at the center of the lower left quadrant of the gene network. We enhanced this effect by 15% and simulated the changes induced in *trn* at *t* = 1, given the measured gene expressions at *t* = 0. Needless to say, in some nuclei where *Doc2* expression level is zero at *t* = 0, the perturbation would not cause any changes in the expression level of *trn* at the next time-point. On the same note, it is expected that the perturbation would induced a relatively bigger change in *trn* in nuclei with a high expression of *Doc2* at *t* = 0, compared to those with a low expression of *Doc2*. Since *Doc2* expression level is not homogeneous across the entire embryo, as shown in Fig. 22(A), we partitioned the embryo into four groups of nuclei: (1) nuclei with a high expression level of *Doc2*, denoted as cell_high_, (2) nuclei with a mid expression level of *Doc2*, cell_mid_, (3) nuclei with a low expression, cell_low_, and (4) nuclei with zero expression level, cell_zero_; see Fig. 22(B).

**Figure 22:**
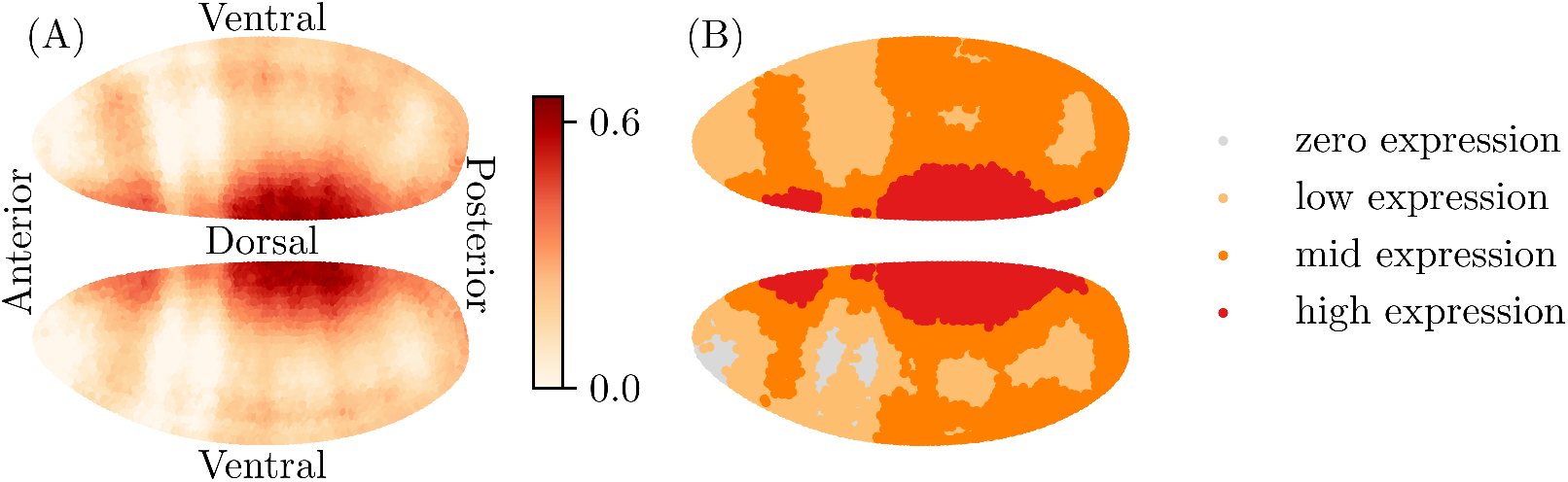
Expressions of *Doc2* at *t* = 0. (A) The expressions of *Doc2* in the nuclei are displayed on a continuous color spectrum. (B) The nuclei are color-coded based on their *Doc2* expression levels.

From the initial conditions of the genes and the perturbed gene network, we simulated the gene profiles after one time step. The resulting profiles of *trn*, *trn_W_P__*, were then compared to those simulated without the perturbation, *trn_W_*. The differences were rescaled by the mean of the differences in cell_high_:

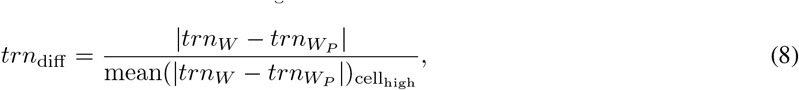

so that we can compare the normalized differences in all nuclei. Fig. 23(A) shows the distributions of the normalized differences in the three groups of nuclei, grouped based on the expression level of *Doc2* at *t* = 0. As we expected, the changes in *trn* expression level induced from the perturbation were predominantly larger in cell_high_, compared to the changes in cell_mid_ and cell_low_.

**Figure 23:**
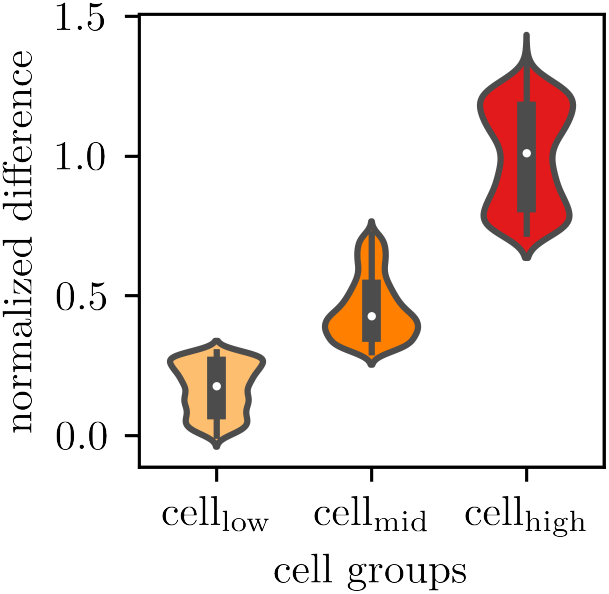
Violin plots showing the distributions of rescaled differences between *trn* simulated with a perturbed gene network and those simulated with the un-perturbed gene network. The effect of *Doc2* on *trn* was (A) increased by 15% and (B) decreased by 10%. We used Eq. 8 to compute and compare the rescaled differences. The color-coding is the same as in Fig. 22(B). The normalized differences in cell_zero_ are not shown, as they are all 0’s.

We then suppressed the effect of *Doc2* on *trn* by 10% and evaluated the effects of the perturbation. Fig. 23(B) shows the distributions of rescaled differences between trn profiles simulated with and without the perturbation in the network. As expected, the simulation showed that the perturbation would induce larger changes in *trn* at *t* = 1 in cell_high_, compared to the changes in other nuclei.

#### Perturbing the effect of Centaurin gamma 1A (*cenGlA*) on sister of odd and bowl (*sob*)

Here is another example of simulating a perturbation in the network. *cenG1A* and *sob* are protein encoding genes. While *cenGlA* encodes a GTPase and is involved in positive regulation of GTPase activity, there are only speculations about the role of *sob* [8]. *cenG1A* has a positive effect on sob. We perturbed this effect by increasing it by 15% and simulated the gene trajectories for one time step. Following the same steps as before, the nuclei were partitioned into three groups based on their *cenGlA* expressions at *t* = 0, cell_high_, cell_mid_, and cell_low_. The color-coded sections are shown in Fig. 24. We then calculated *sob*’s rescaled differences induced from the perturbation:

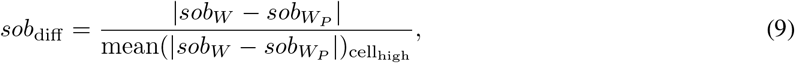

where *sob_W_* and *sob_W_P__* are simulated profiles of *sob* at *t* = 1, with and without the perturbation in the network. The distributions of normalized differences are shown in Fig. 25. Again, we compared the distributions in three groups of nuclei, which showed that the majority of cell_high_ group, the ones with high expressions of *cenG1A* at *t* = 0, has the largest changes in *sob* at *t* = 1. We conducted a series of perturbations for both positive and negative effects, and they are included in the Supporting Materials A (Figs. S5 - S14).

**Figure 24:**
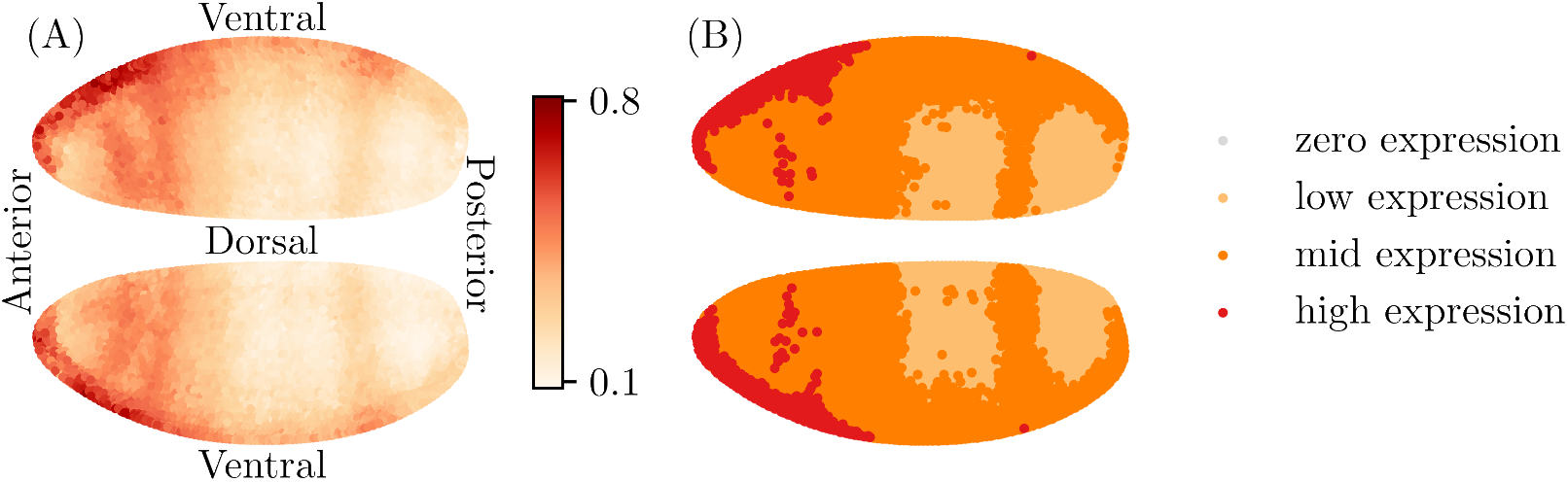
Expressions of *cenG1A* at *t* = 0. (A) The expressions of *cenG1A* in the nuclei are displayed on a continuous color spectrum. (B) The nuclei are partitioned and color-coded based on their *cenG1A* expression levels.

**Figure 25:**
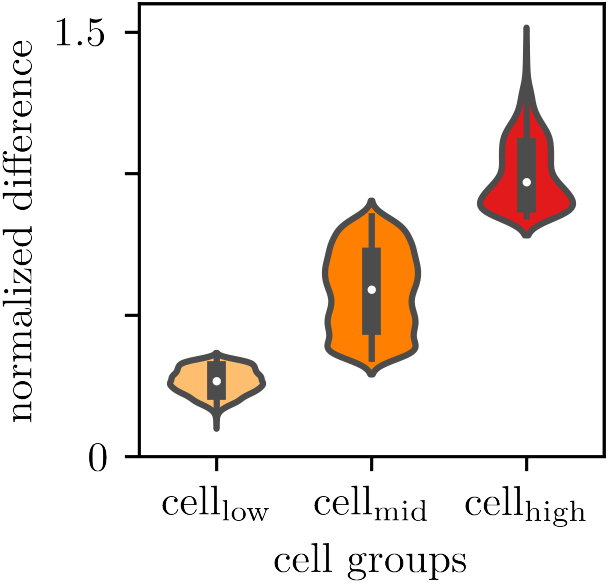
Violin plots showing the distributions of rescaled differences between *sob* simulated with a perturbed gene network and those simulated with the un-perturbed gene network. The effect of *cenG1A* on *sob* was increased by 15%. We used Eq. 9 to compute and compare the rescaled differences. The color-coding is the same as in Fig. 24(B)

## 4 Discussion

In this paper, we introduced a stable variation of LAD regression that sidesteps the singularity problem of LAD regression when the prediction errors are small while maintaining its robustness to the effects of outliers. We used this regression to infer a mechanistic gene network model that governs the discrete gene dynamics observed in *Drosophila melanogaster* blastoderm embryo. The model not only takes the linear gene profiles into account but also incorporates the quadratic products of gene pairs.

The performance of erf-weighted LAD regression was evaluated by comparing the predictive power of the inferred model with those of models inferred with least squares and ridge regressions and trained neural networks. To be comprehensive, we inferred models at various sample sizes. As we decreased the sample size, we implemented ridge regularization in our regression to remediate having a degenerate covariance matrix. Our results show that at various sample sizes, the models inferred with our methods predict more accurately than neural networks. Our method significantly outperforms ridge regression, especially in the limit of small sample size.

By inferring gene network models from different sections of the embryo, we addressed the global applicability of the models across the entire embryo. The results suggest that due to the limited cell-to-cell variability in gene dynamics, the models inferred with subsets of nuclei in close proximity are not as apt to predict the gene dynamics in nuclei on other parts of the embryo.

With the inferred mechanistic gene network, we made predictions on the effects of gene knockouts and introducing a small perturbation to a gene-to-gene interaction. Obtaining an effective gene network and making such predictions is not feasible through a black-box neural network modeling approach.

The mechanistic gene network presented in this paper is inferred from spatiotemporal gene expression data. Despite the comprehensiveness of the data set, it was incomplete. The data set had some missing time-points for some genes. We trained neural networks to impute the missing data. While we cannot be certain that our imputation method is optimal for the given data, we thoroughly examined the neural network imputation with randomized trials of imputing when simulating missing gene expressions.

When comparing model simulations of pair-rule gene mutants to the empirical studies in Schroeder et al. [18], we noticed a common phenomenon. The stripe intensities and regularities affected by the pair-rule gene mutations were well captured in the model simulations. On the other hand, if the mutations led to a change in the stripe position, i.e., shifting anteriorly, this change was not reproduced in the model simulations. We believe this is due to the lack of information on the true initial gene expressions of the mutants. In our mutant simulations, the initial conditions are the same as the wild type, except for the mutated gene expression, which was set to 0. However, it is not unfounded to expect significantly different initial conditions in mutants, and those may involve a shift in the stripe positions.

We emphasize that the inferred gene network is a universal and comprehensive model of gene dynamics in the nuclei that cover the entire embryo. This is different modeling approach from the one presented by Fowlkes et al. [4], where, for a given target gene, a small set of regulators that best describes that gene’s spatiotemporal dynamics was selected and separate domains of target pattern (i.e., individual stripes) were regressed independently. Similarly, one could infer multiple gene networks for different clusters of nuclei to better understand the local gene dynamics. Then, the challenge would be how to stitch the networks together to obtain a comprehensive common network. However, we do not know how to do this in a principled way, and we suspect that a more useful approach would be to work with larger sets of gene measurements instead.

As our future work, we aim to deepen the understanding of gene regulatory network in *Drosophila melanogaster* embryo by applying the inference algorithm to the extensive single-cell data presented by Karaiskos et a. [6] and Nitzan et al. [13].

## Acknowledgement

Research reported in this publication was supported by the Intramural Research Program of the National Institute of Diabetes and Digestive and Kidney Diseases, NIH (to JMH, SP, and VP), NIH NICHD K99/R00 HD073191 (to ZW). This work utilized the computational resources of the NIH HPC Biowulf cluster (http://hpc.nih.gov). We are grateful to Dr. Allon Klein at Harvard Medical School for helpful comments and questions.

