## Supporting Materials A for "Mechanistic Gene Networks Inferred from Single-Cell Data are Better Predictors than Neural Networks"

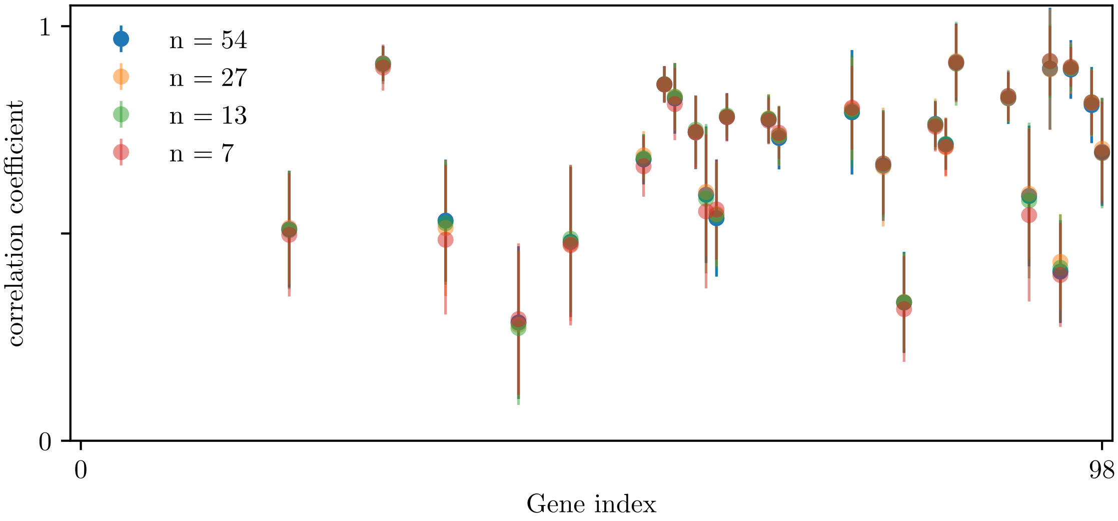


Figure S1: The correlation coefficients between the imputed and observed gene expressions. The number of nodes on the hidden layer was varied, $n = [7, 13, 27, 54]$.

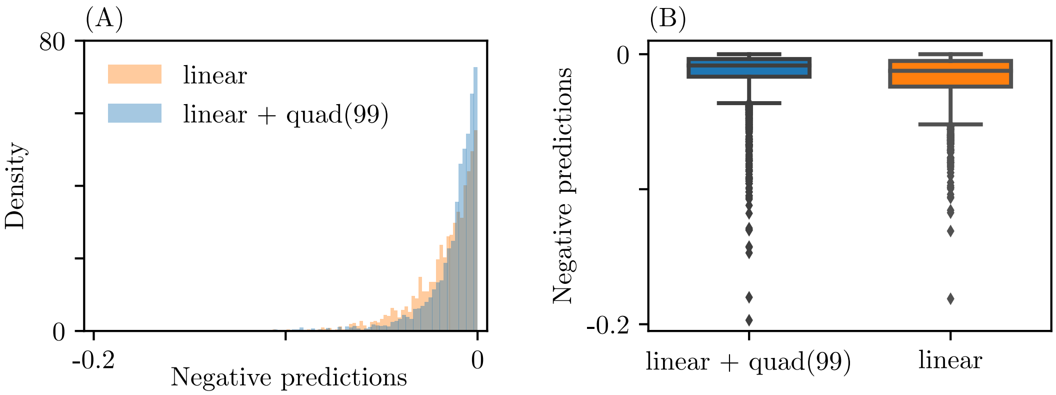


Figure S2: The negative predictions from the models inferred with linear predictors only and linear and quadratic predictors, displayed in (A) distributions and (B) boxplots. The Mann-Whitney $U$ test confirms that the two sets of negative predictions have significantly different medians with a $p$-value of 5.83 $\times 10^{-31}$.

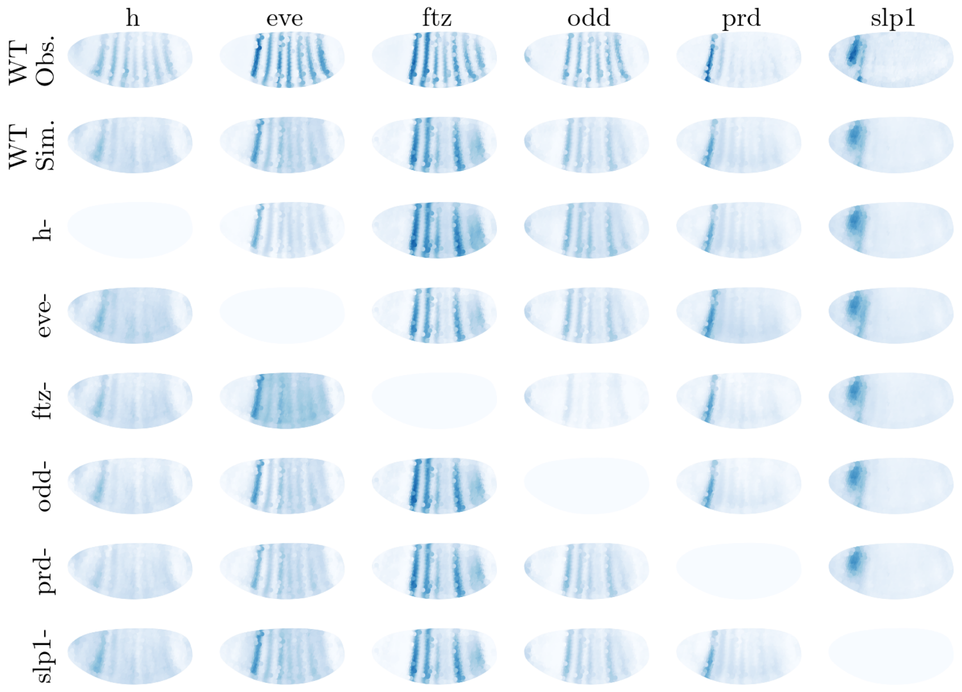


Figure S3: Mutant simulations using the trained neural networks.

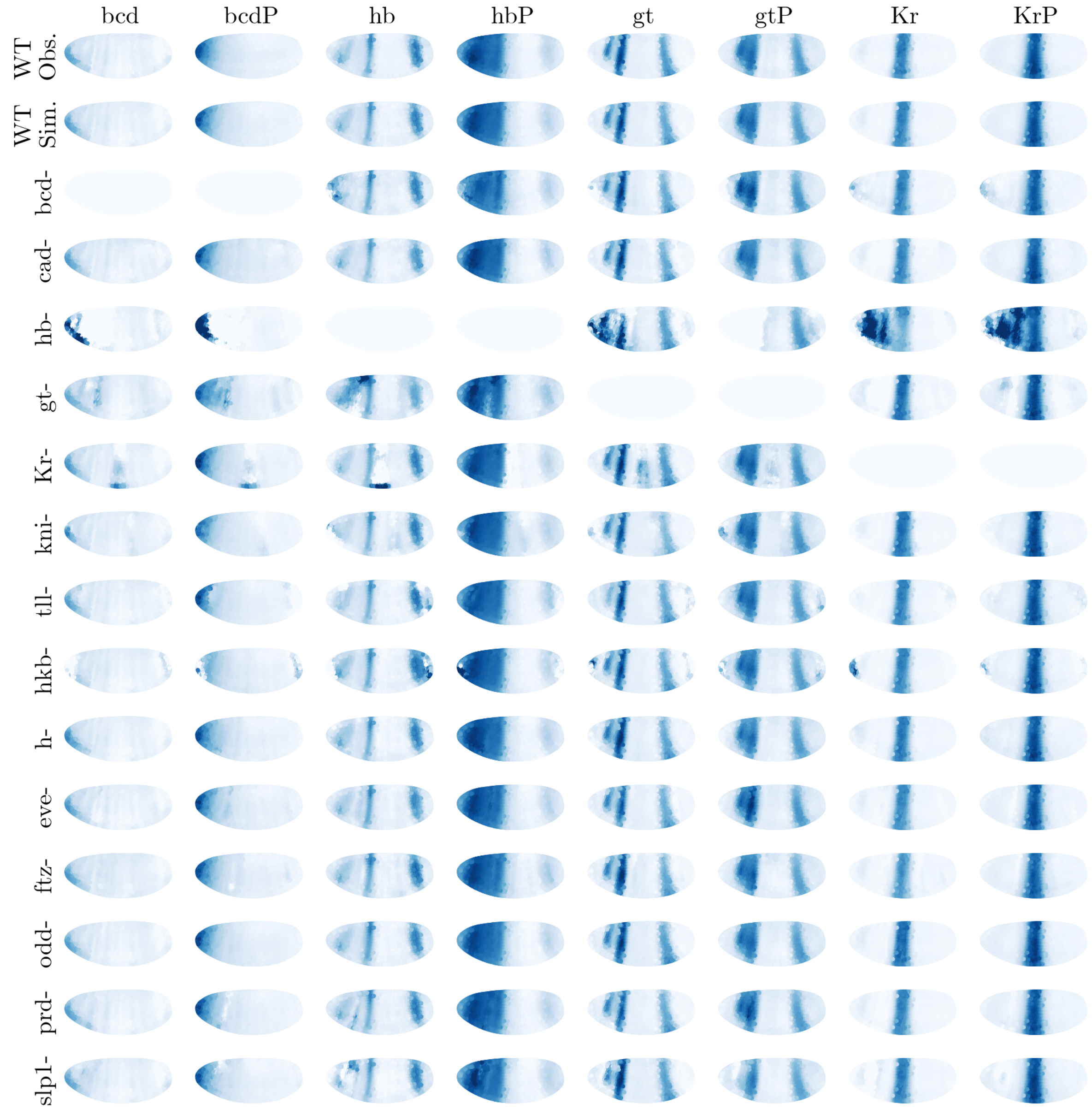


Figure S4: Mutant simulations: effects of nullifying maternal genes, gap genes, and pair-rule genes on the measured protein levels.

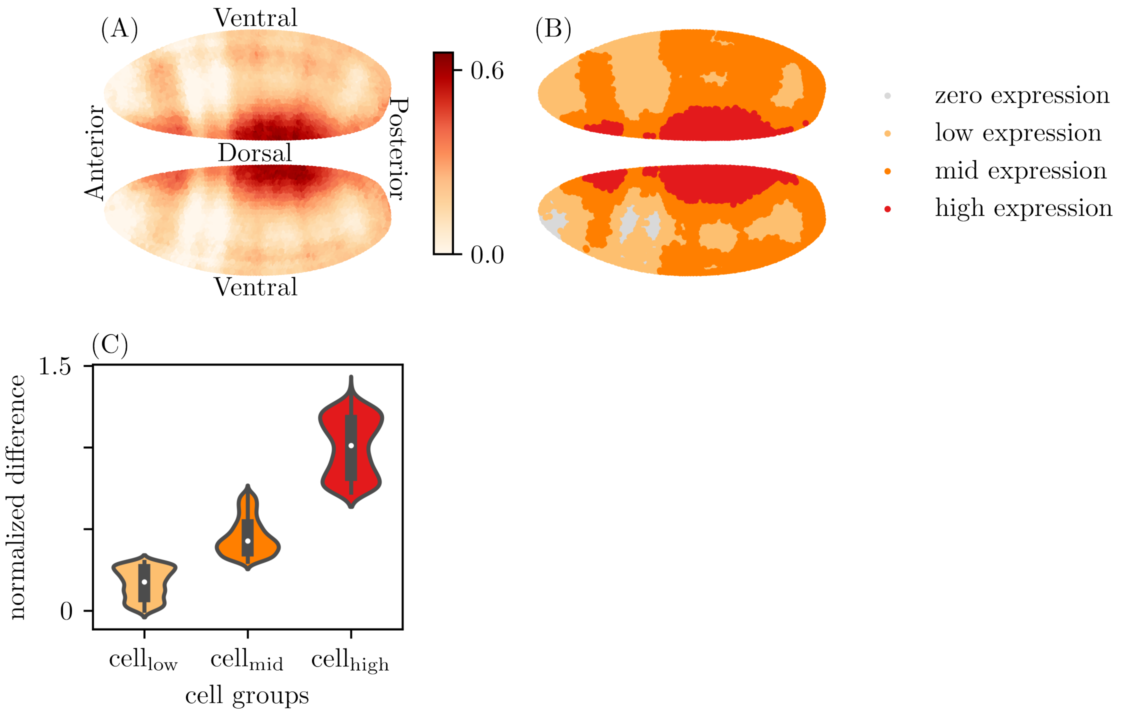


Figure S5: The model prediction of increasing the effect of *Dorsocross2* on *deadpan* by 15%.

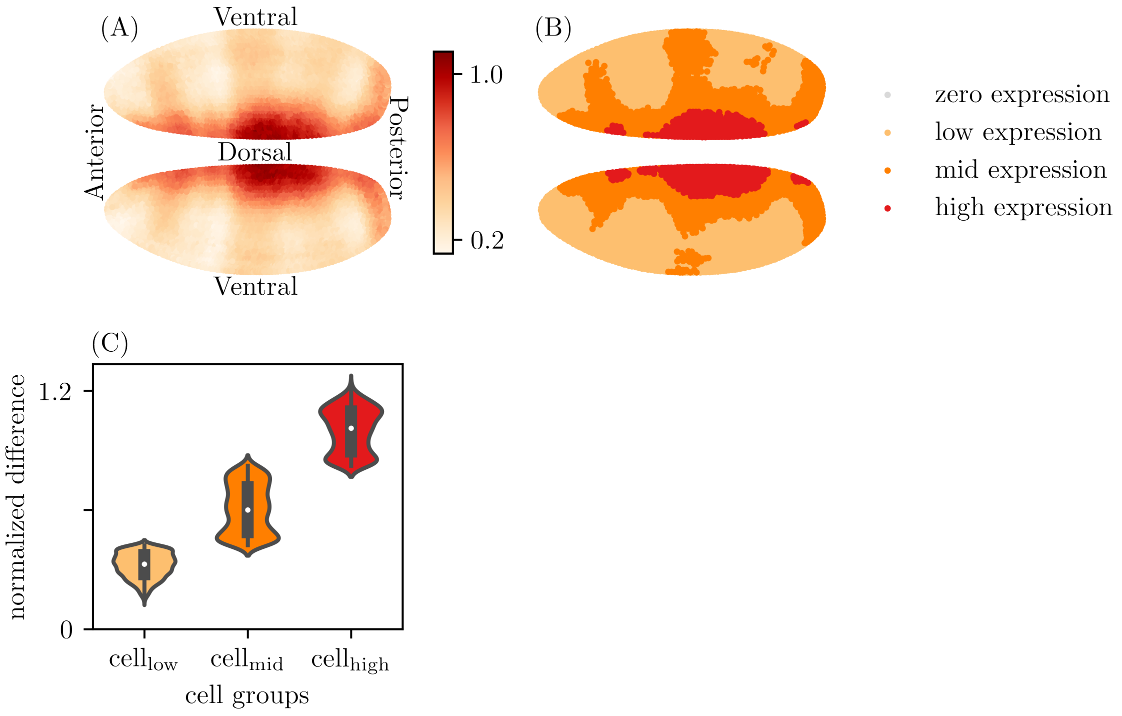


Figure S6: The model prediction of increasing the effect of *Dorsocross3* on *toucan* by 15%.

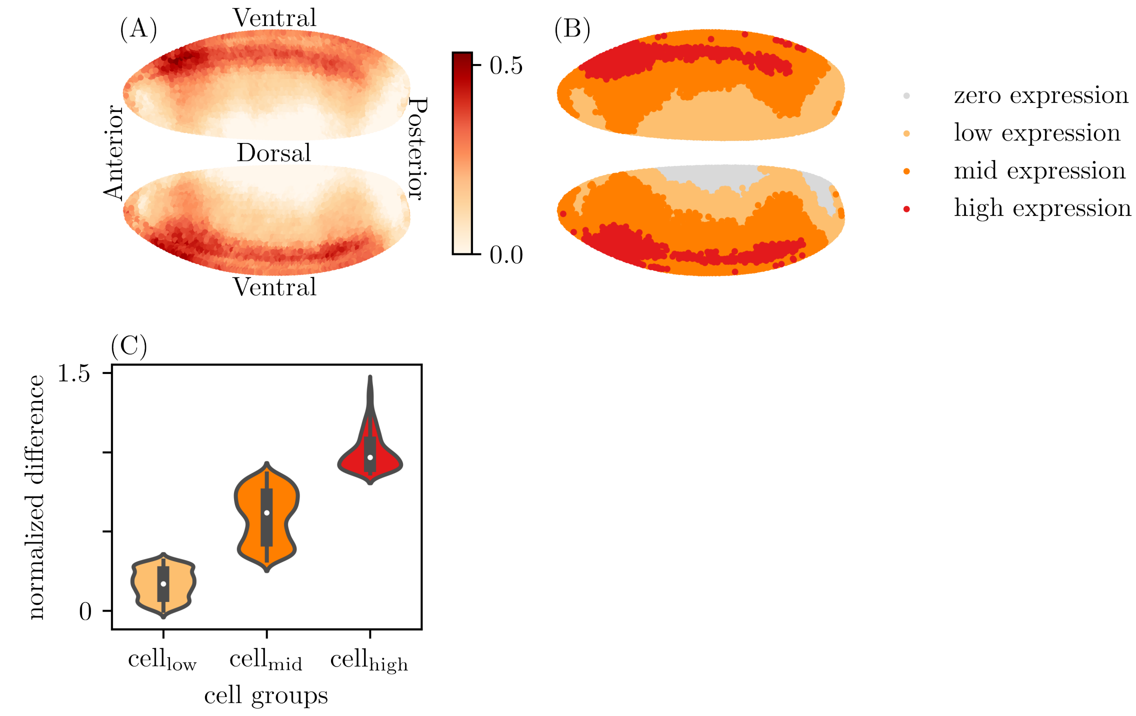


Figure S7: The model prediction of increasing the effect of *HLHm5* on *Blimp* by 15%.

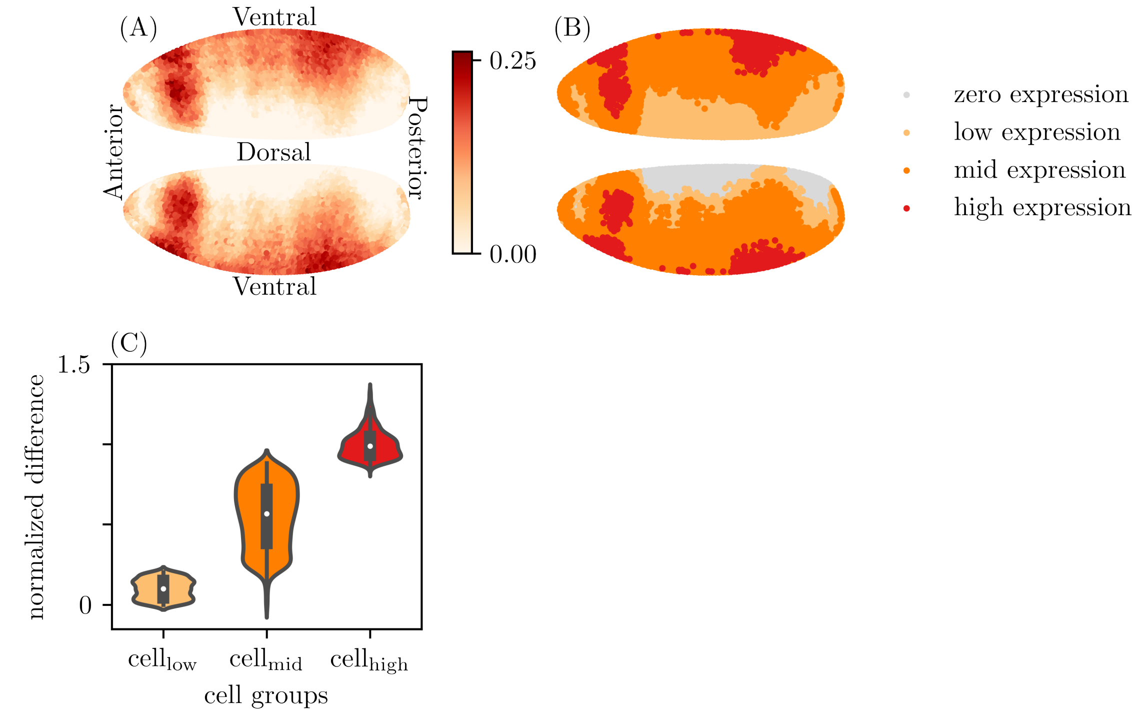


Figure S8: The model prediction of increasing the effect of *CG17786* on *deadpan* by 15%.

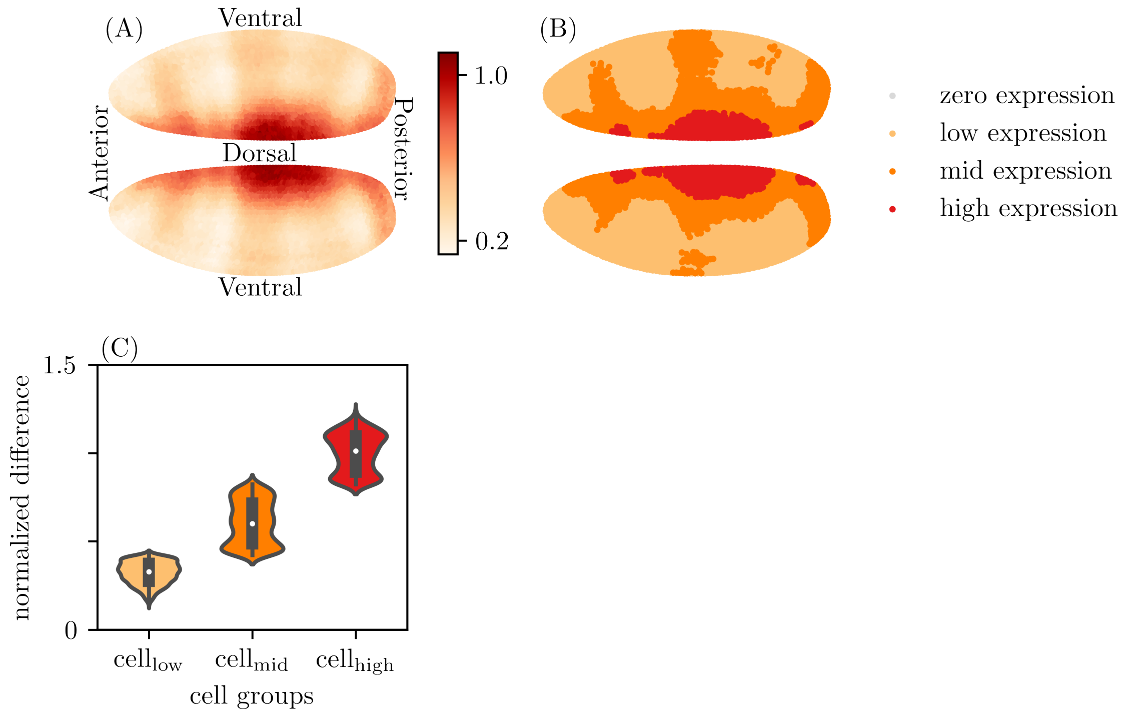


Figure S9: The model prediction of decreasing the effect of *Dorsocross3* on *tartan* by 15%.

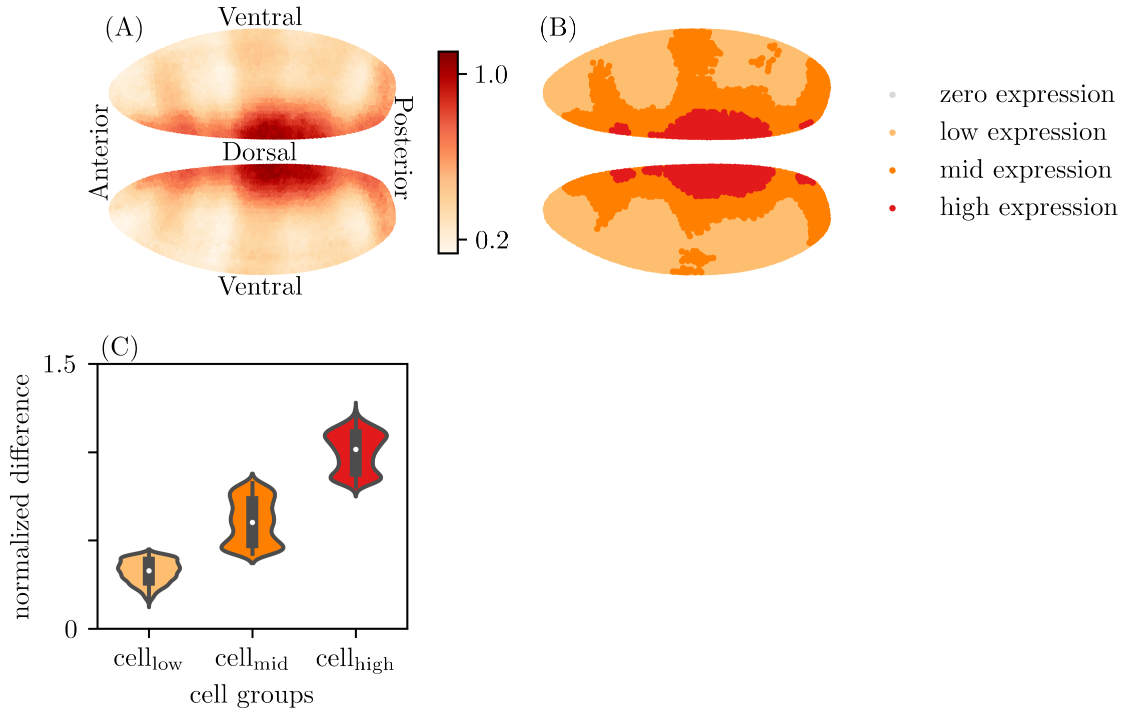


Figure S10: The model prediction of decreasing the effect of *Dorsocross3* on *CG13333* by 15%.

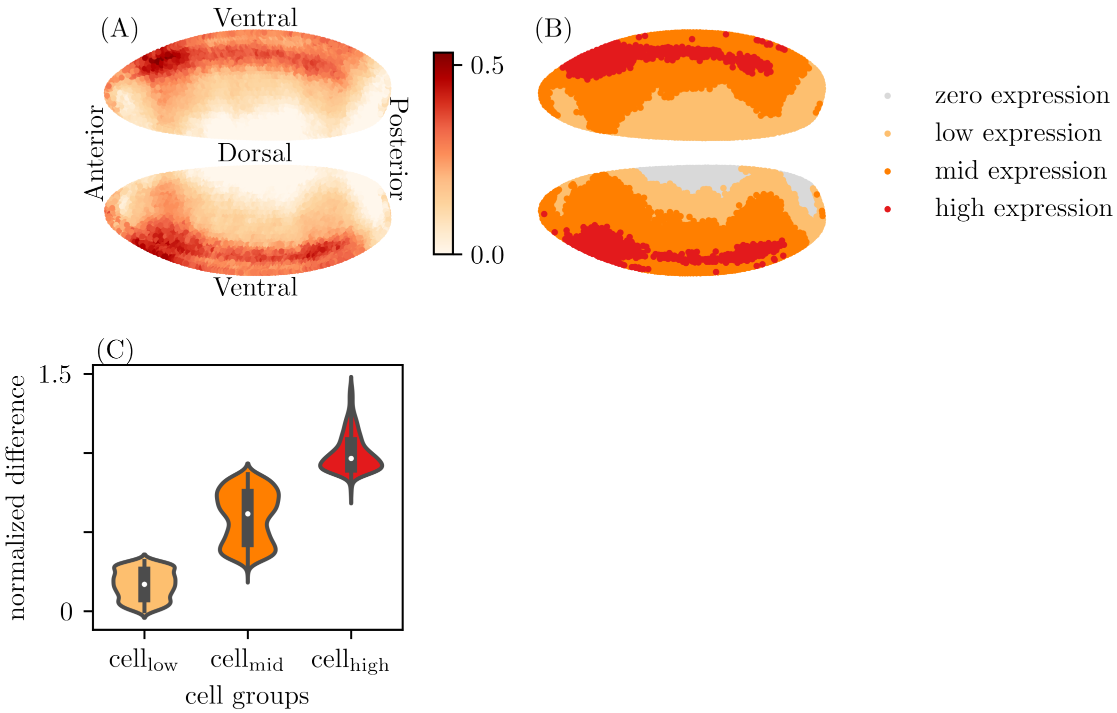


Figure S11: The model prediction of decreasing the effect of *HLHm5* on *deadpan* by 15%.

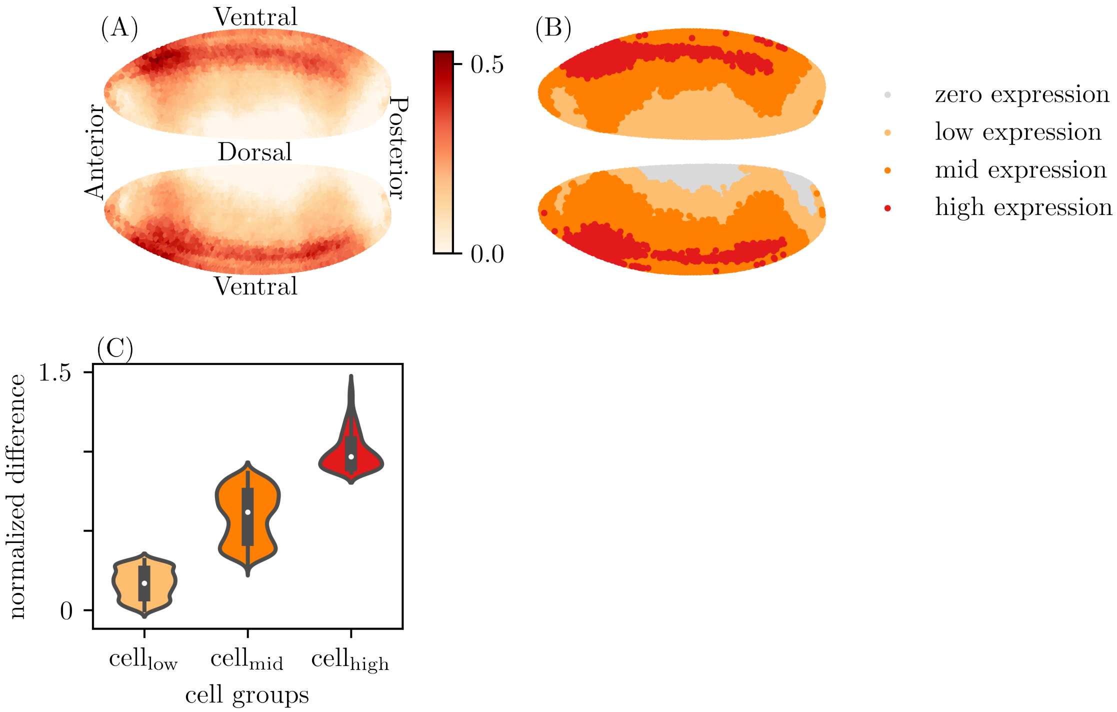


Figure S12: The model prediction of decreasing the effect of *HLHm5* on *paired* by 15%.

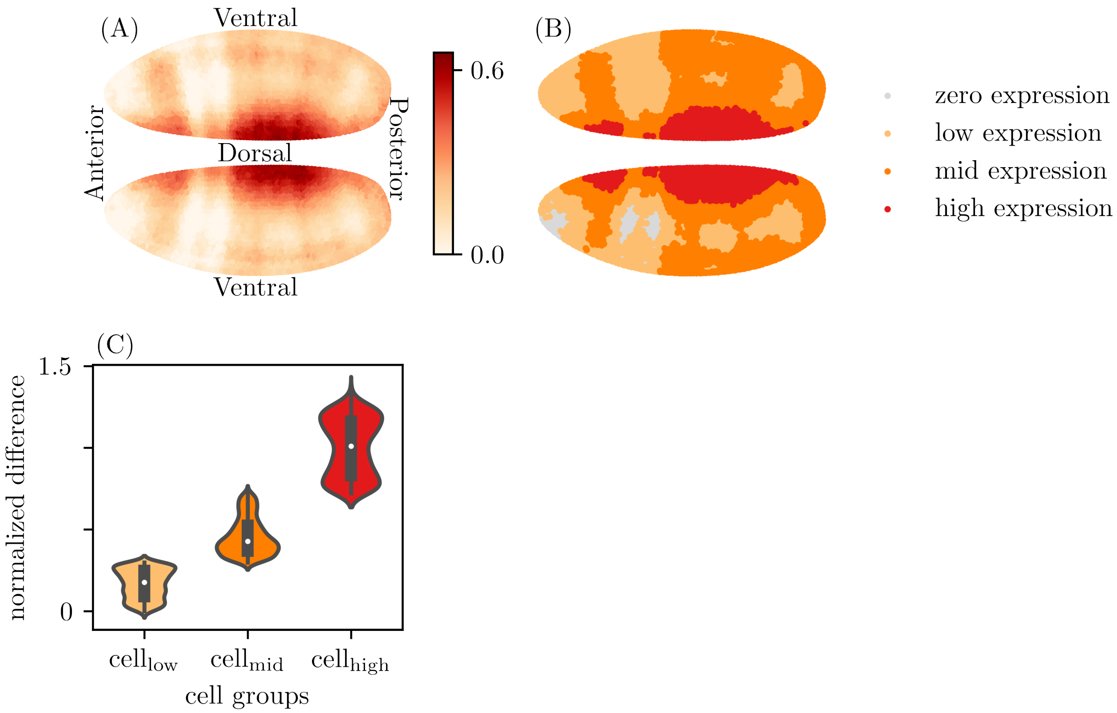


Figure S13: The model prediction of decreasing the effect of *Dorsocross2* on *toucan* by 15%.

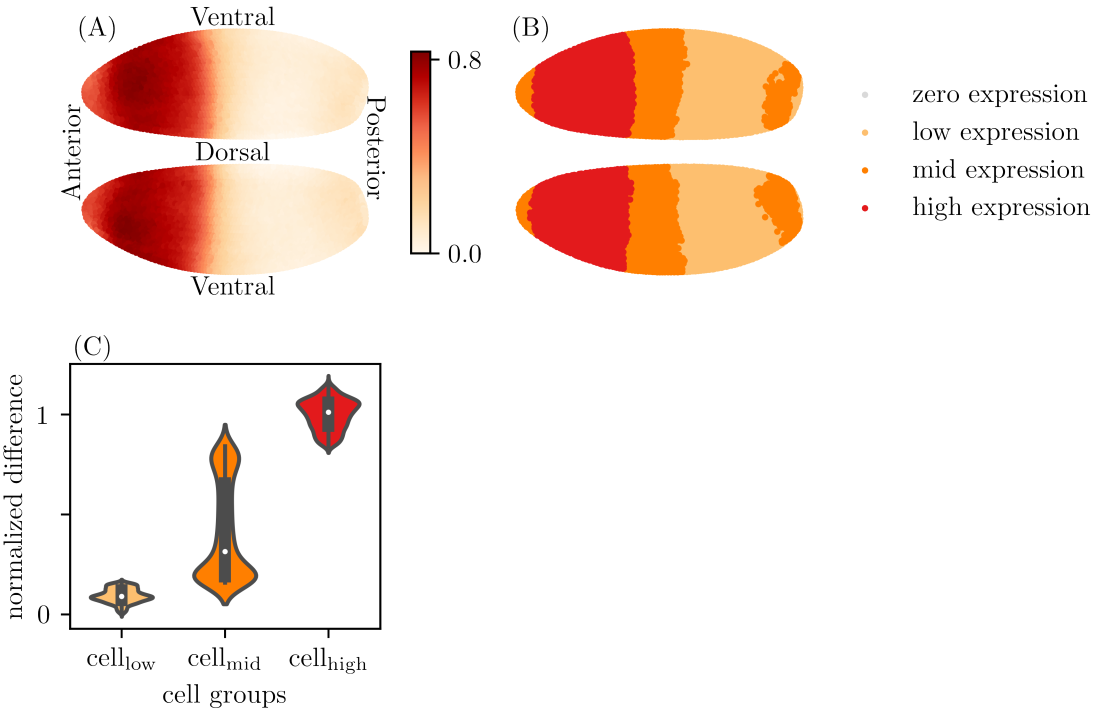


Figure S14: The model prediction of decreasing the effect of *Hunchback*(protein) on *tartan* by 15%.
