## Supporting Materials B for "Mechanistic Gene Networks Inferred from Single-Cell Data are Better Predictors than Neural Networks"

This document includes the individual mutant simulations of 95 mRNA. Genotypes are arranged by row, transcript patterns by column. On each page, the top two rows show the observed and simulated wild type gene expressions at $t = 2$, respectively.

For each simulation, we followed these steps:

1. Set the initial expressions of the mutated gene to 0's.
2. Eliminate all the effects that are correlated with the mutated gene (both linear and quadratic effects) by setting the corresponding coefficients in the model ($W$ and *bias*) to 0's.
3. Using the modified model, simulated the gene expressions at $t = 2$.

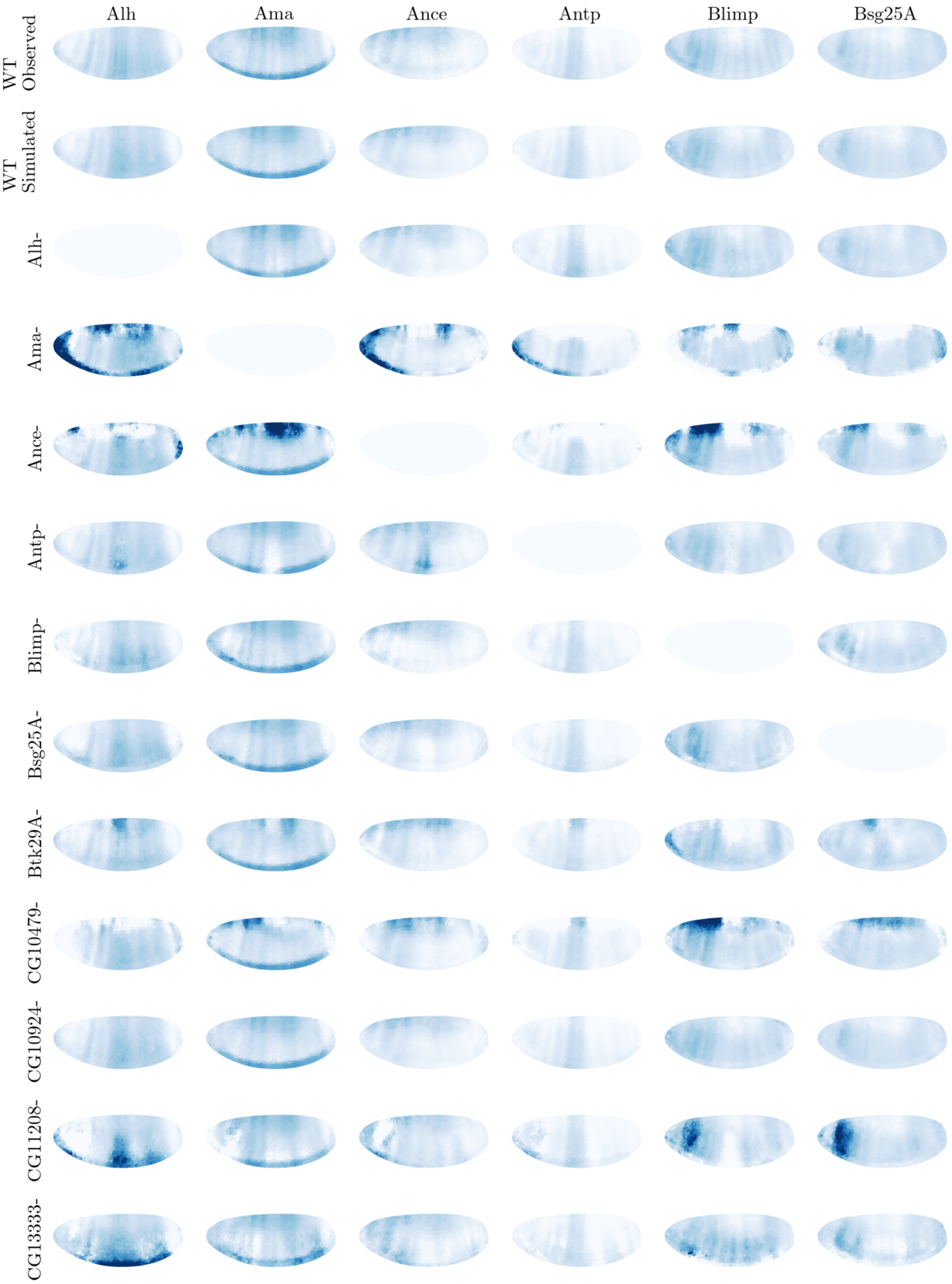
Alh
Ama
Ance
Antp
Blimp
Bsg25A
WT Observed
WT Simulated
Alh-
Ama-
Ance-
Antp-
Blimp-
Bsg25A-
Btk29A-
CG10479-
CG10924-
CG11208-
CG13333-

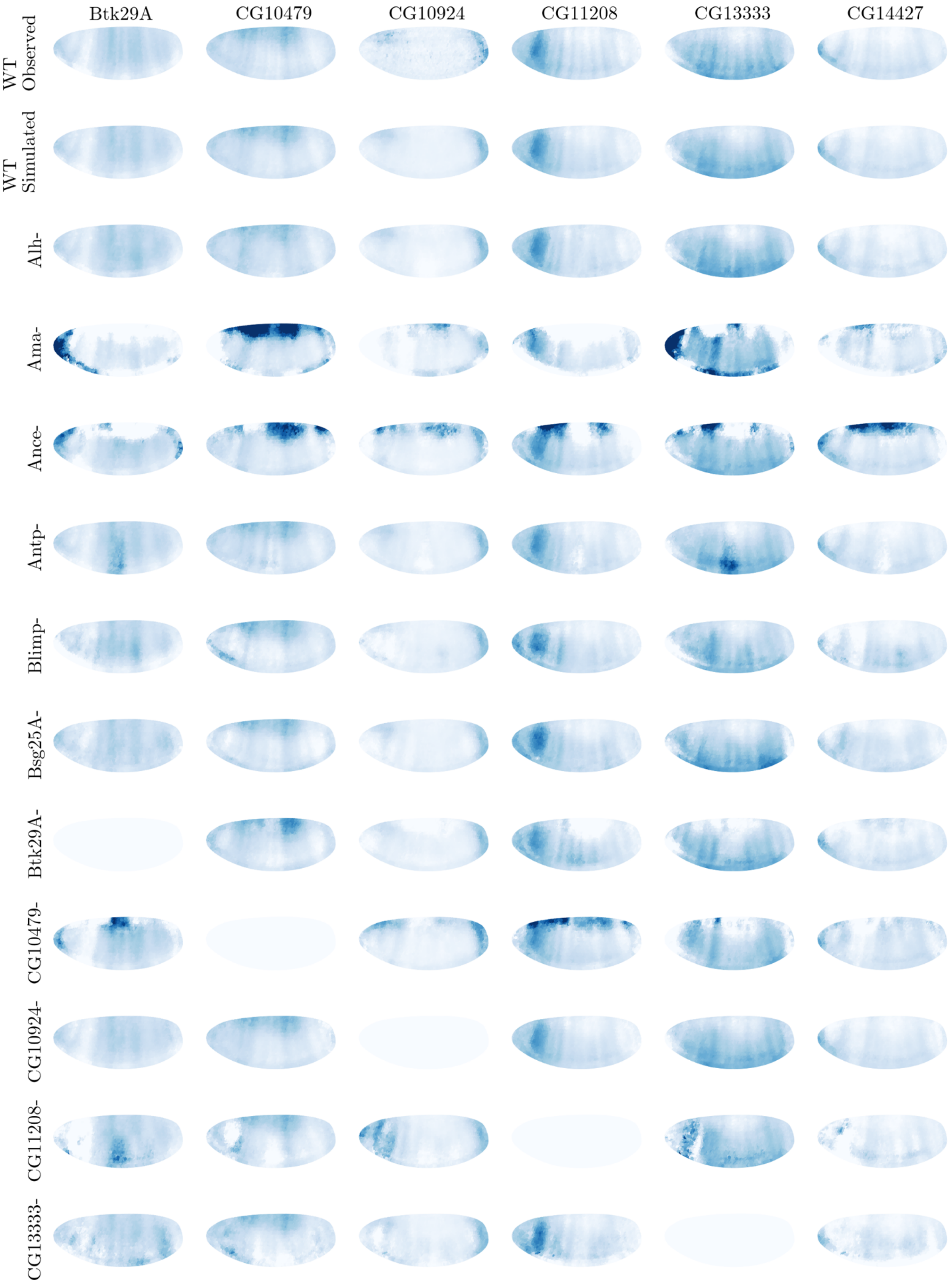
Btk29A
CG10479
CG10924
CG11208
CG13333
CG14427
WT Observed
WT Simulated
Alh-
Ama-
Ance-
Antp-
Blimp-
Bsg25A-
Btk29A-
CG10479-
CG10924-
CG11208-
CG13333-

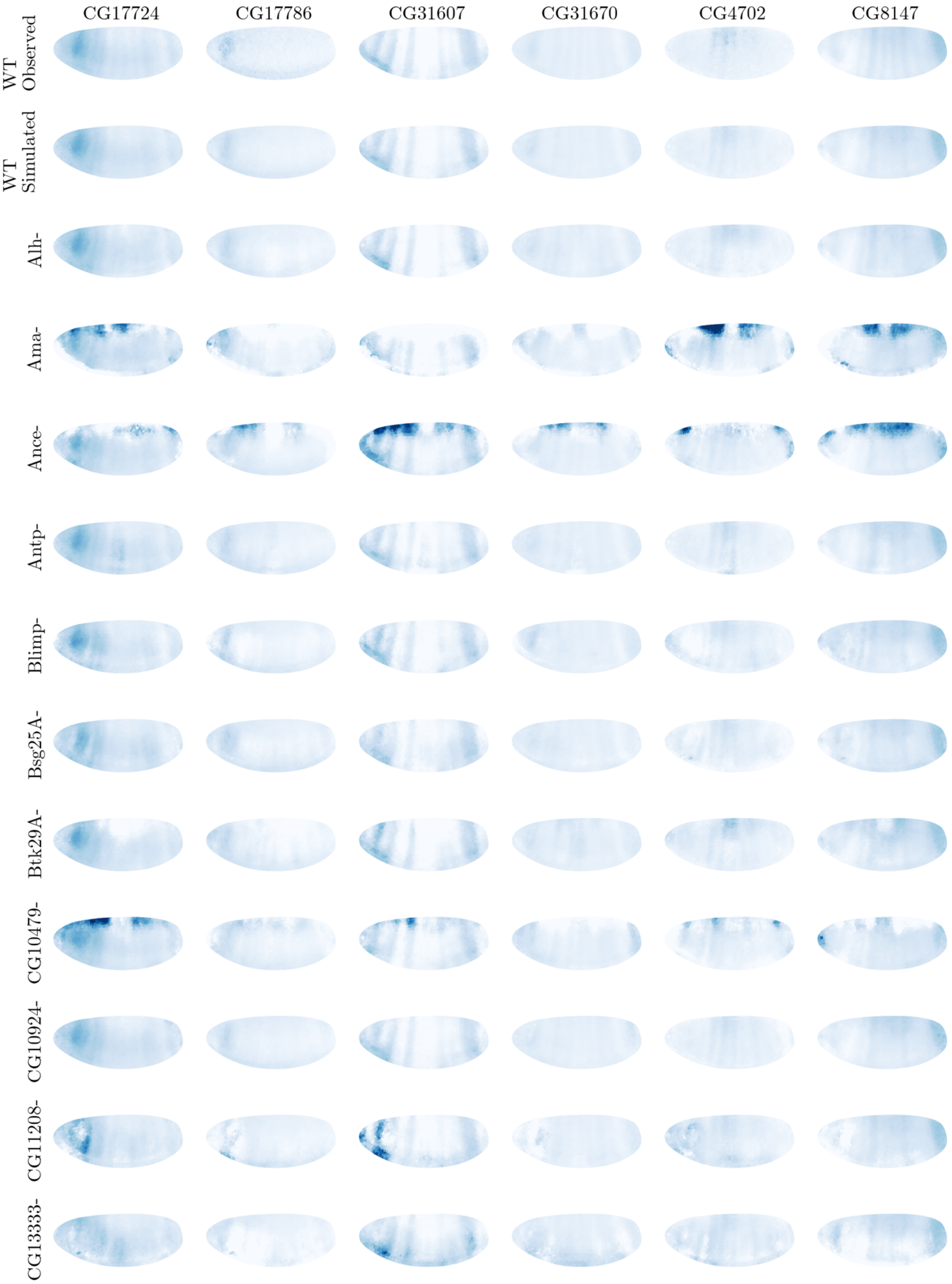
CG17724
CG17786
CG31607
CG31670
CG4702
CG8147
WT Observed
WT Simulated
Alh-
Ama-
Ance-
Antp-
Blimp-
Bsg25A-
Btk29A-
CG10479-
CG10924-
CG11208-
CG13333-

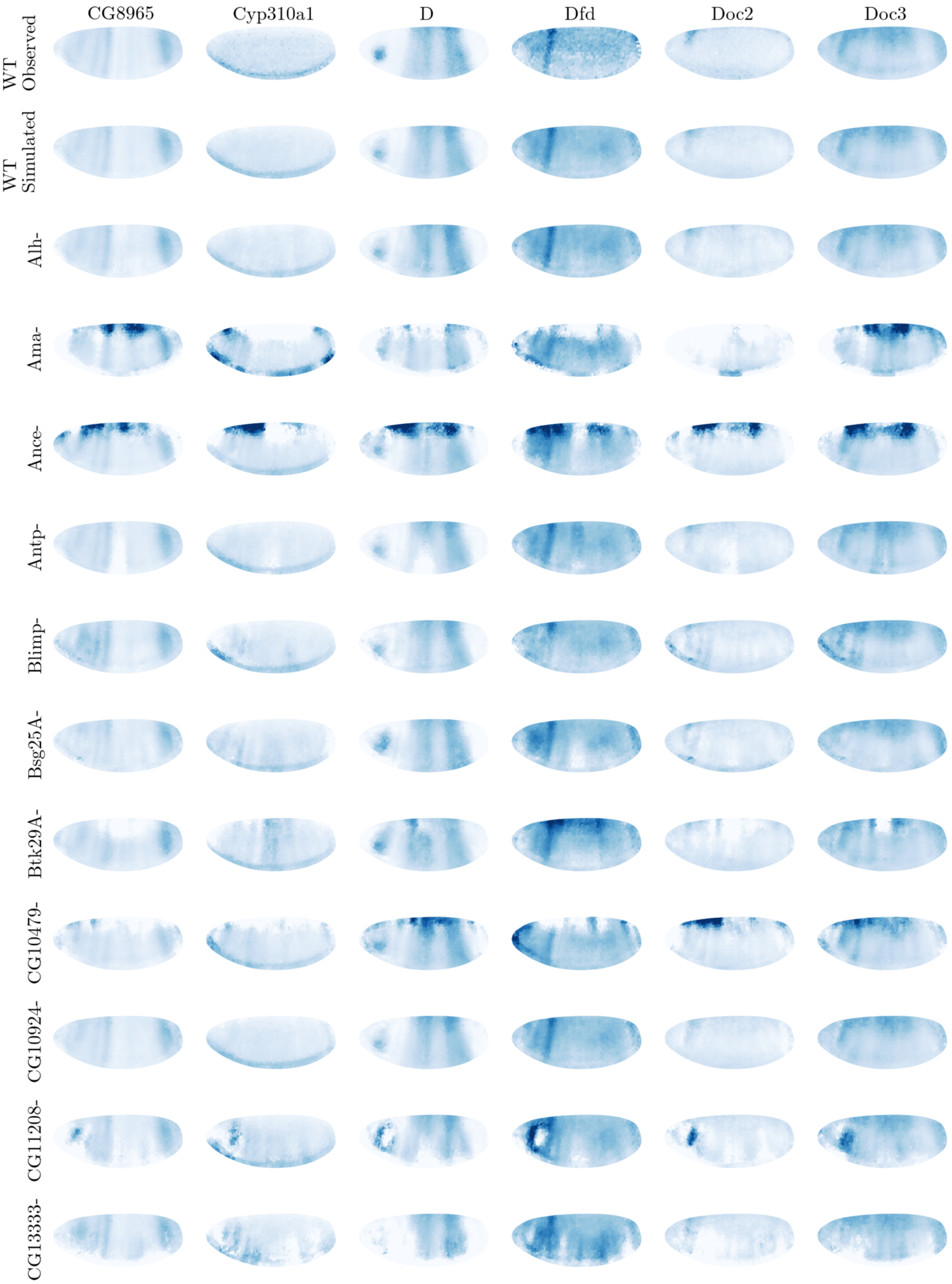
CG8965
Cyp310a1
D
Dfd
Doc2
Doc3
WT Observed
WT Simulated
Alh-
Ama-
Ance-
Antp-
Blimp-
Bsg25A-
Btk29A-
CG10479-
CG10924-
CG11208-
CG13333-

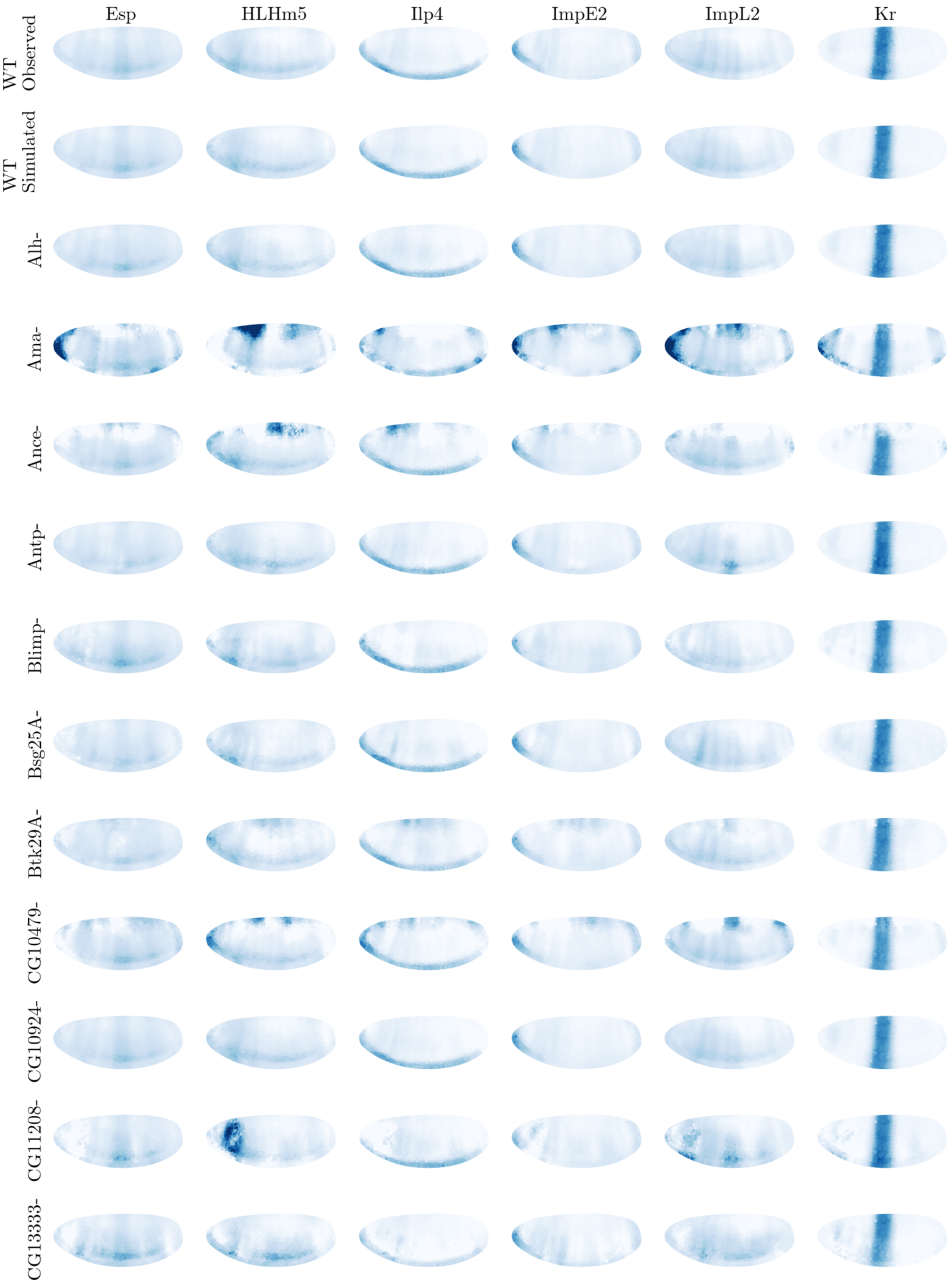
Esp
HLHm5
Ilp4
ImpE2
ImpL2
Kr
WT Observed
WT Simulated
Alh-
Ama-
Ance-
Antp-
Blimp-
Bsg25A-
Btk29A-
CG10479-
CG10924-
CG11208-
CG13333-

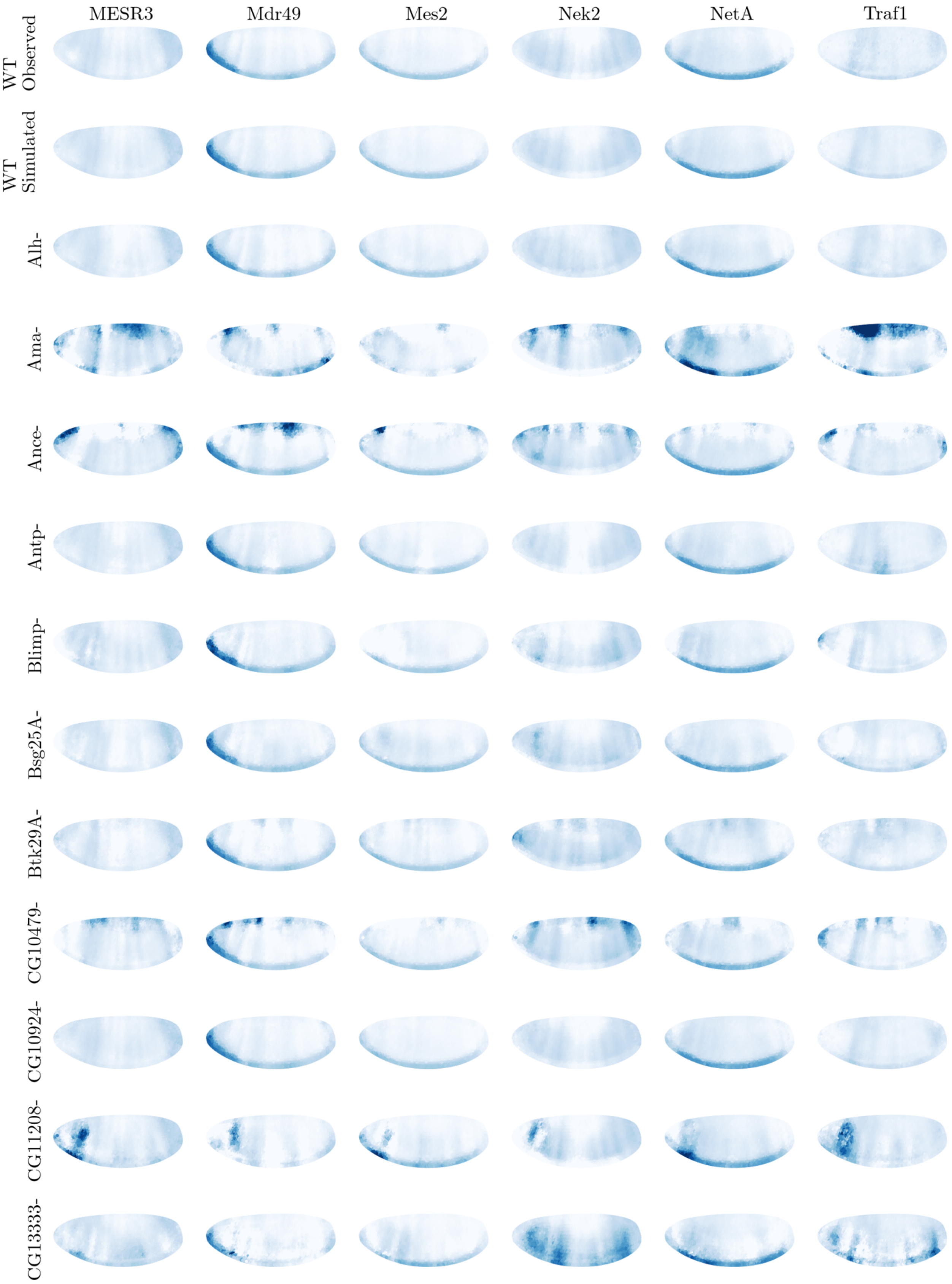
MESR3
Mdr49
Mes2
Nek2
NetA
Traf1
WT Observed
WT Simulated
Alh-
Ama-
Ance-
Antp-
Blimp-
Bsg25A-
Btk29A-
CG10479-
CG10924-
CG11208-
CG13333-

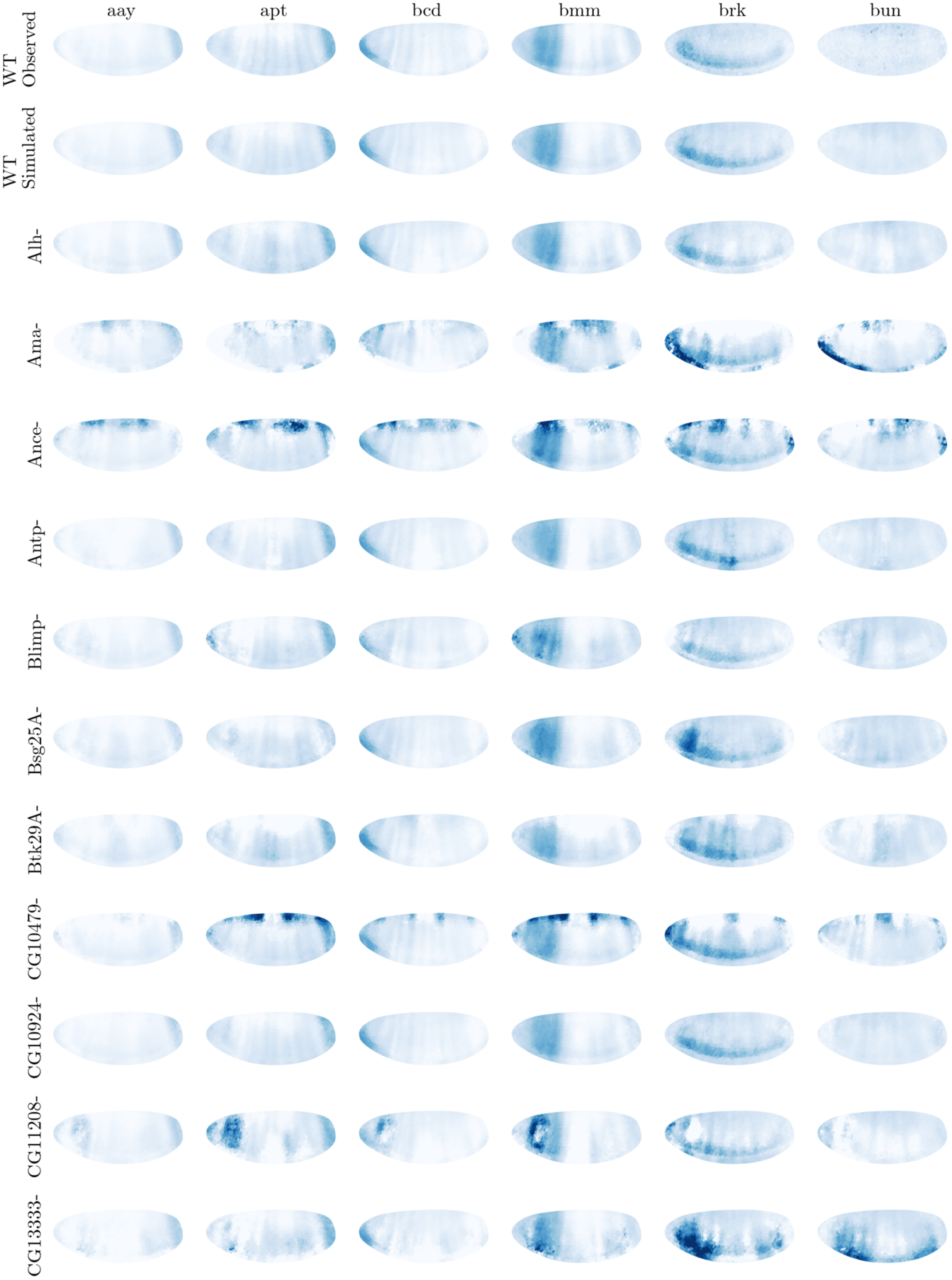
aay
apt
bcd
bmm
brk
bun
WT Observed
WT Simulated
Alh-
Ama-
Ance-
Antp-
Blimp-
Bsg25A-
Btk29A-
CG10479-
CG10924-
CG11208-
CG13333-

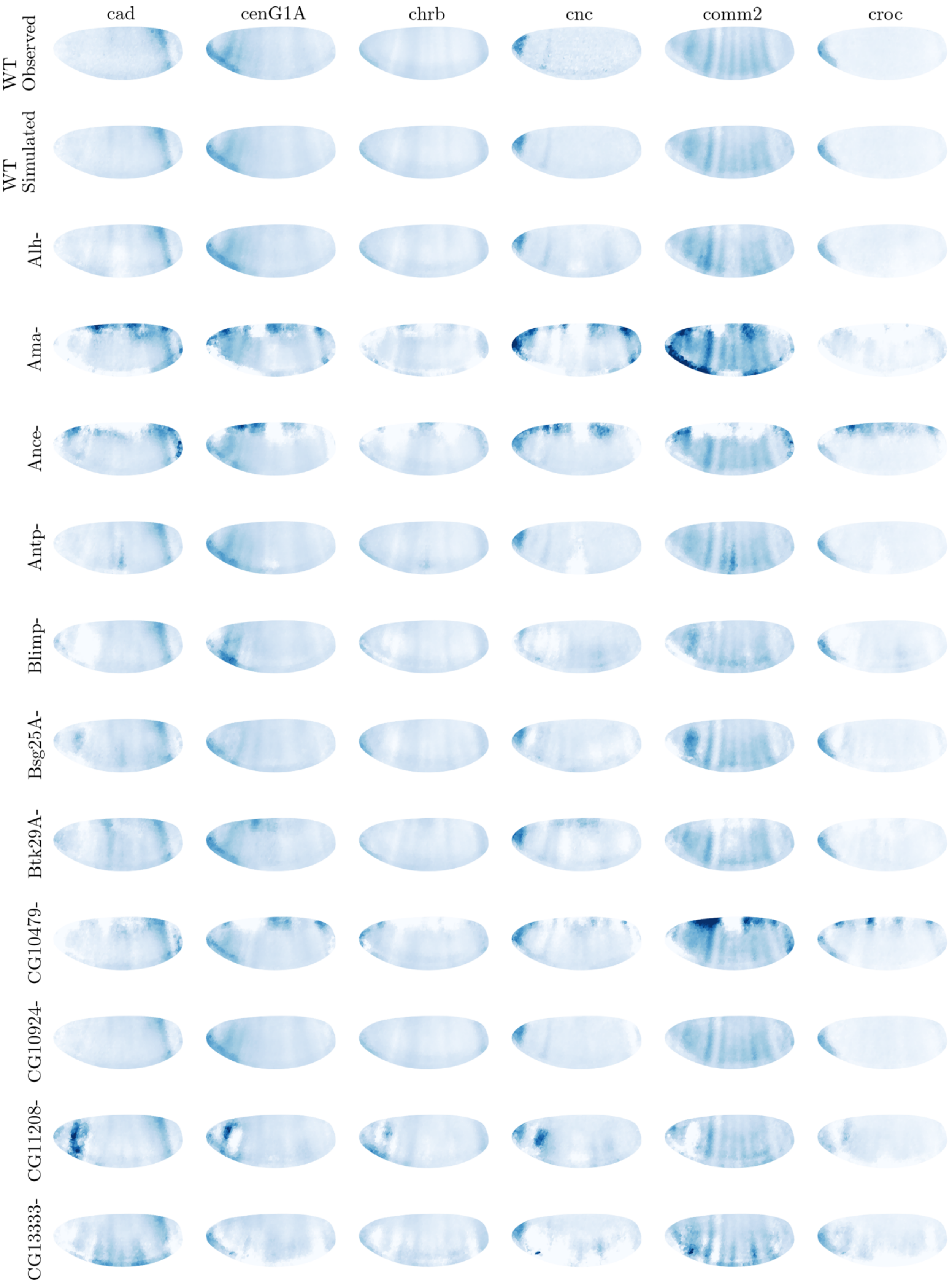

cad
cenG1A
chrb
cnc
comm2
croc
WT Observed
WT Simulated
Alh-
Ama-
Ance-
Antp-
Blimp-
Bsg25A-
Btk29A-
CG10479-
CG10924-
CG11208-
CG13333-

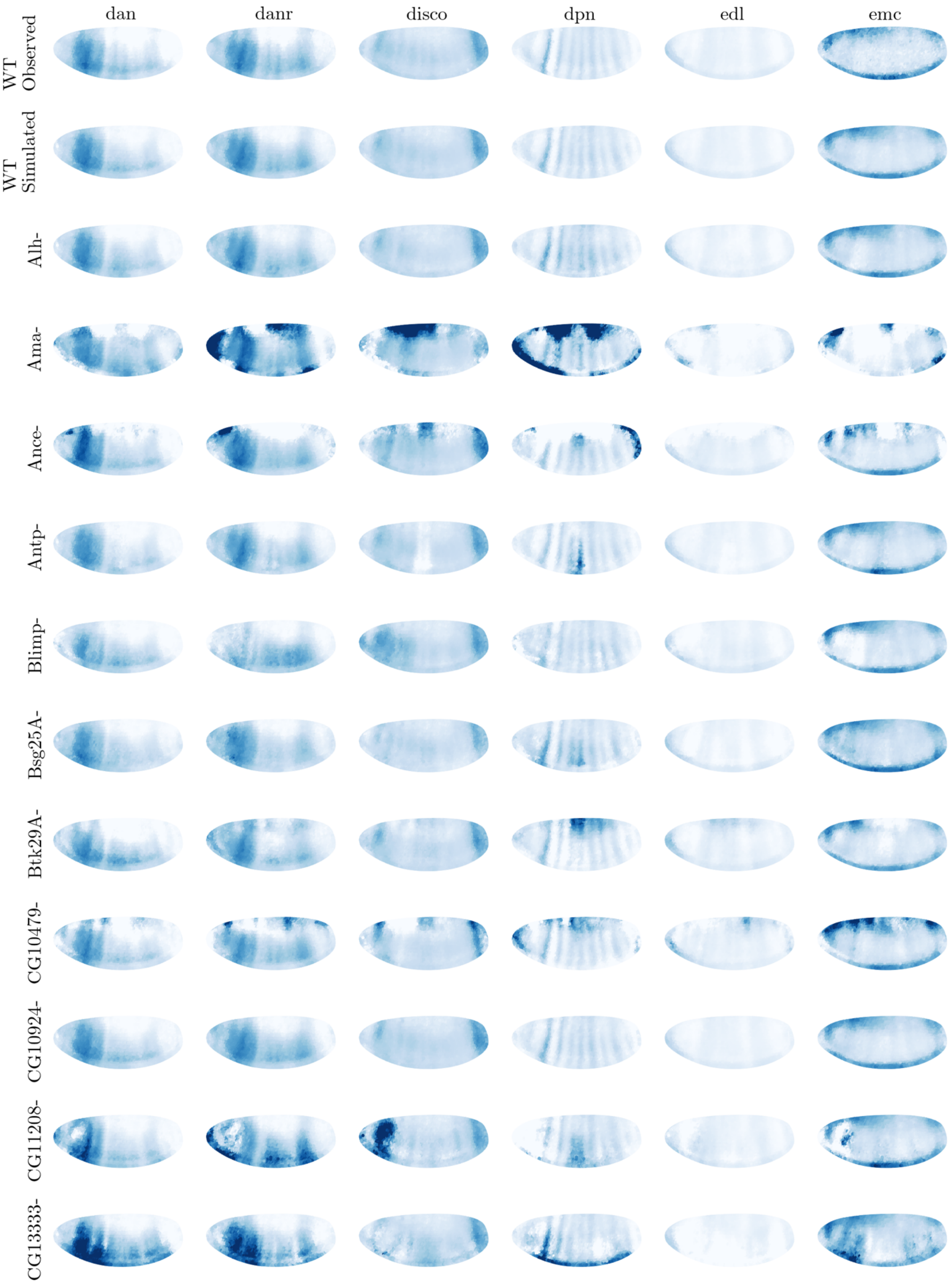
dan
danr
disco
dpn
edl
emc
WT Observed
WT Simulated
Alh-
Ama-
Ance-
Antp-
Blimp-
Bsg25A-
Btk29A-
CG10479-
CG10924-
CG11208-
CG13333-

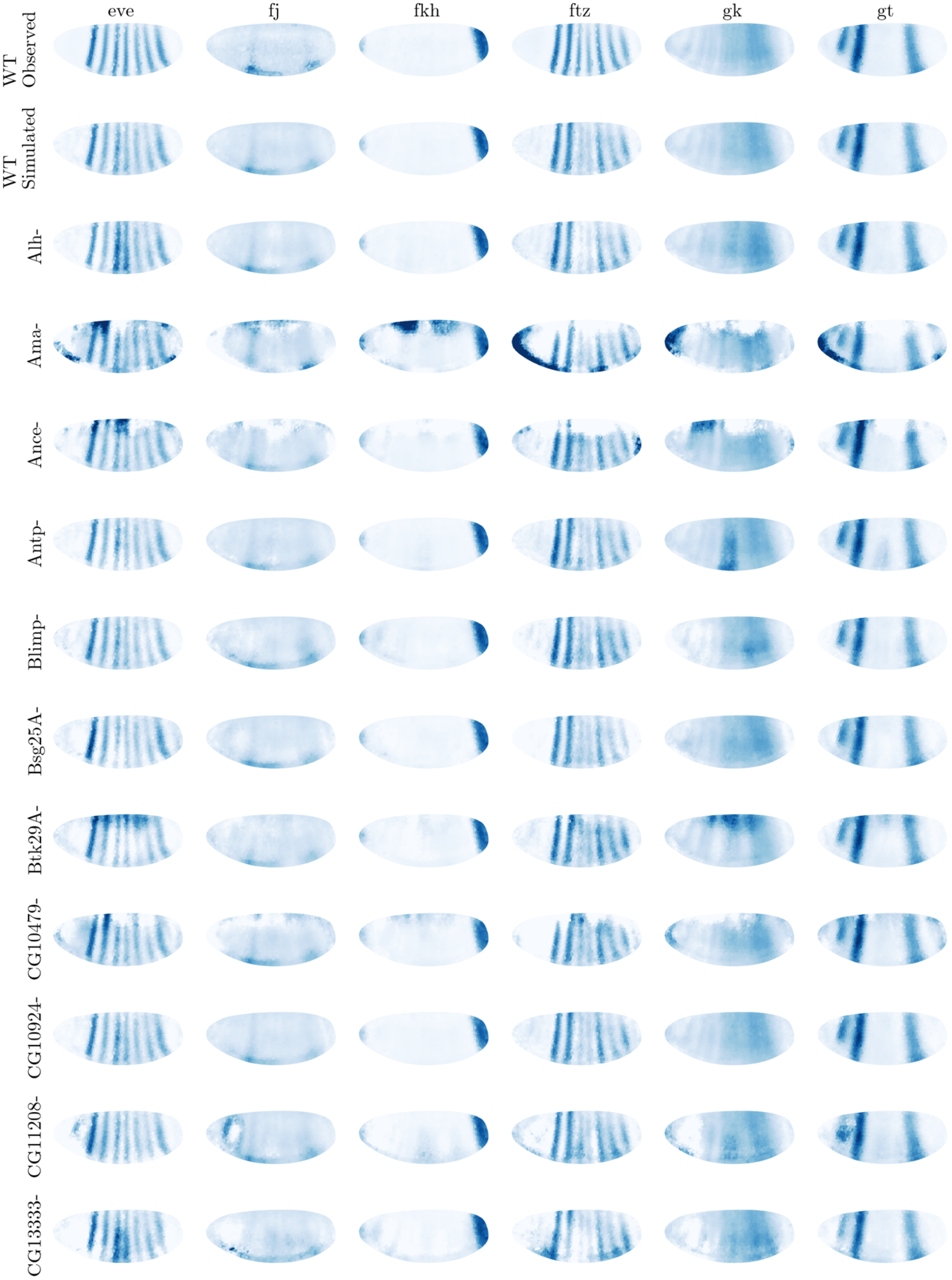
eve
fj
fkh
ftz
gk
gt
WT Observed
WT Simulated
Alh-
Ama-
Ance-
Antp-
Blimp-
Bsg25A-
Btk29A-
CG10479-
CG10924-
CG11208-
CG13333-

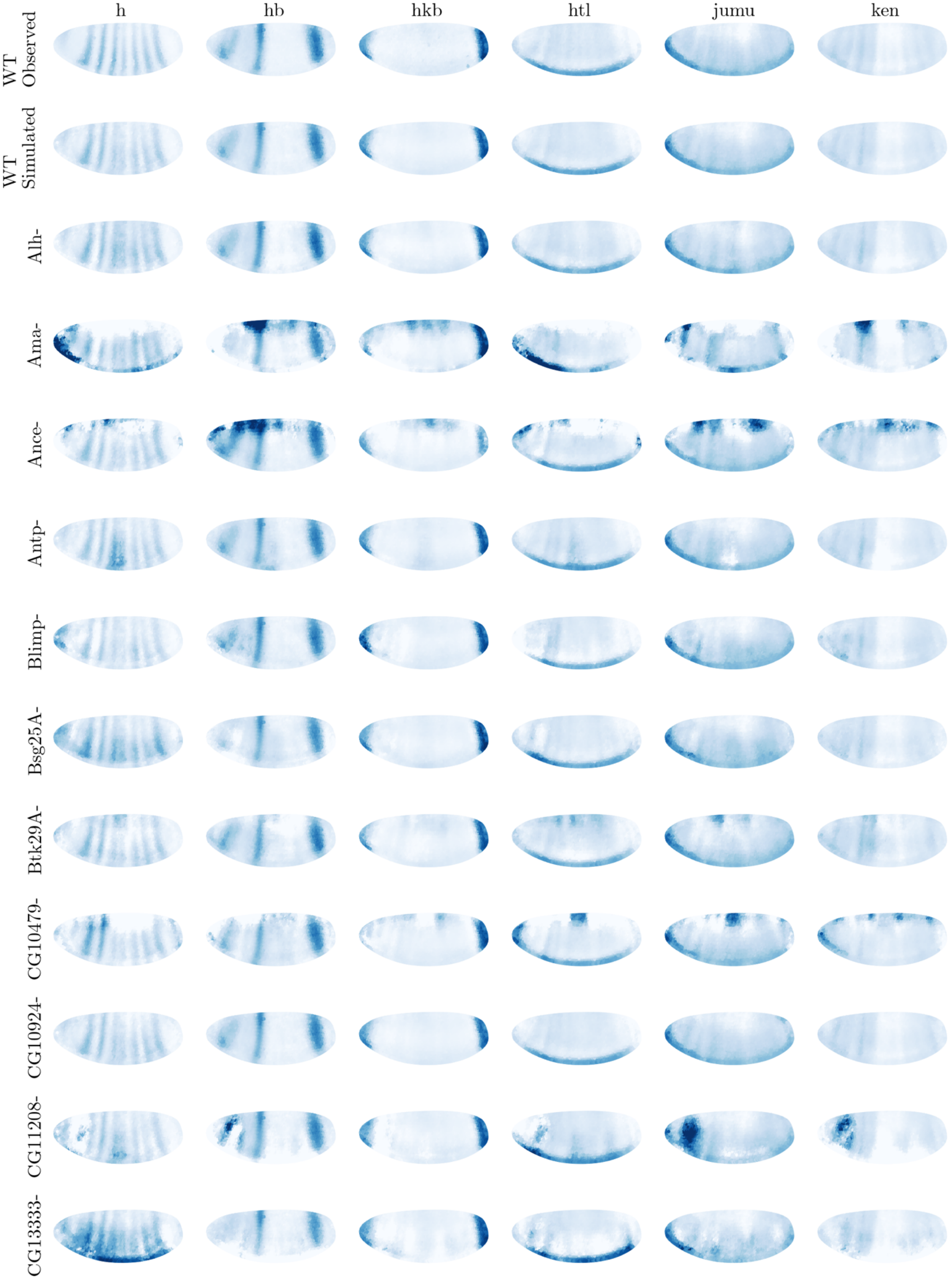

h
hb
hkb
htl
jumu
ken
WT Observed
WT Simulated
Alh-
Ama-
Ance-
Antp-
Blimp-
Bsg25A-
Btk29A-
CG10479-
CG10924-
CG11208-
CG13333-

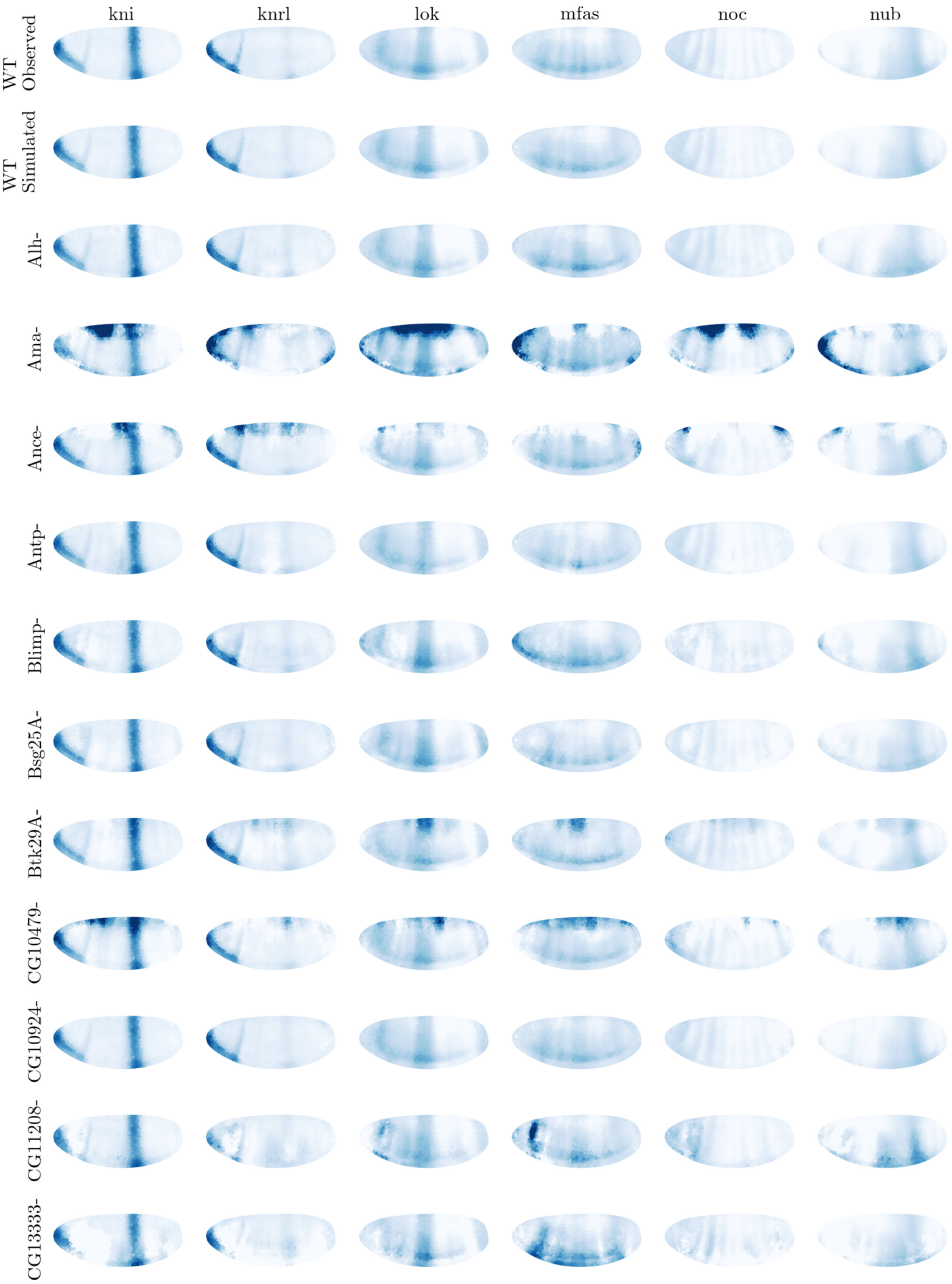
kni
knrl
lok
mfas
noc
nub
WT Observed
WT Simulated
Alh-
Ama-
Ance-
Antp-
Blimp-
Bsg25A-
Btk29A-
CG10479-
CG10924-
CG11208-
CG13333-

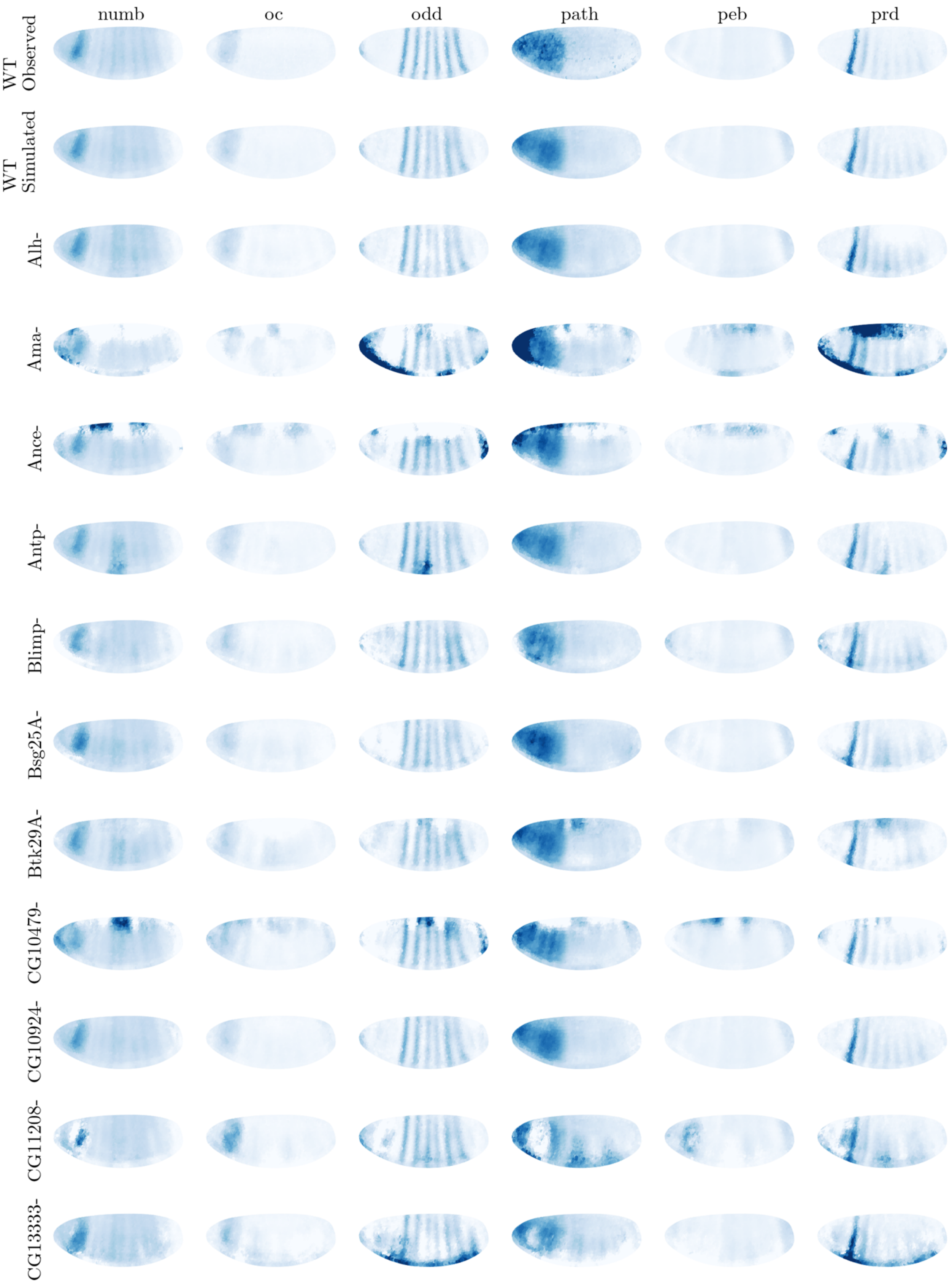

numb
oc
odd
path
peb
prd
WT Observed
WT Simulated
Alh-
Ama-
Ance-
Antp-
Blimp-
Bsg25A-
Btk29A-
CG10479-
CG10924-
CG11208-
CG13333-

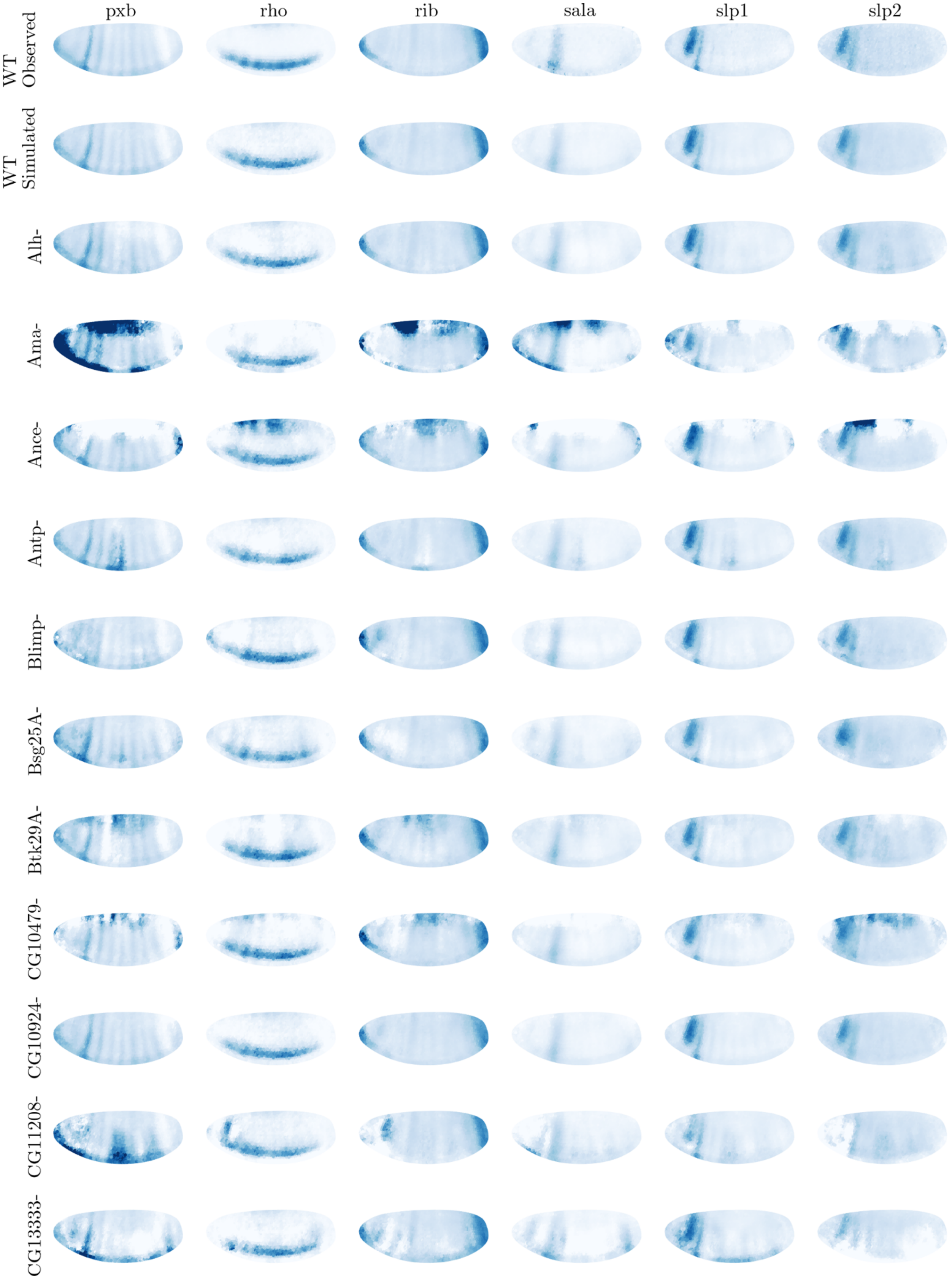
pxb
rho
rib
sala
slp1
slp2
WT Observed
WT Simulated
Alh-
Ama-
Ance-
Antp-
Blimp-
Bsg25A-
Btk29A-
CG10479-
CG10924-
CG11208-
CG13333-

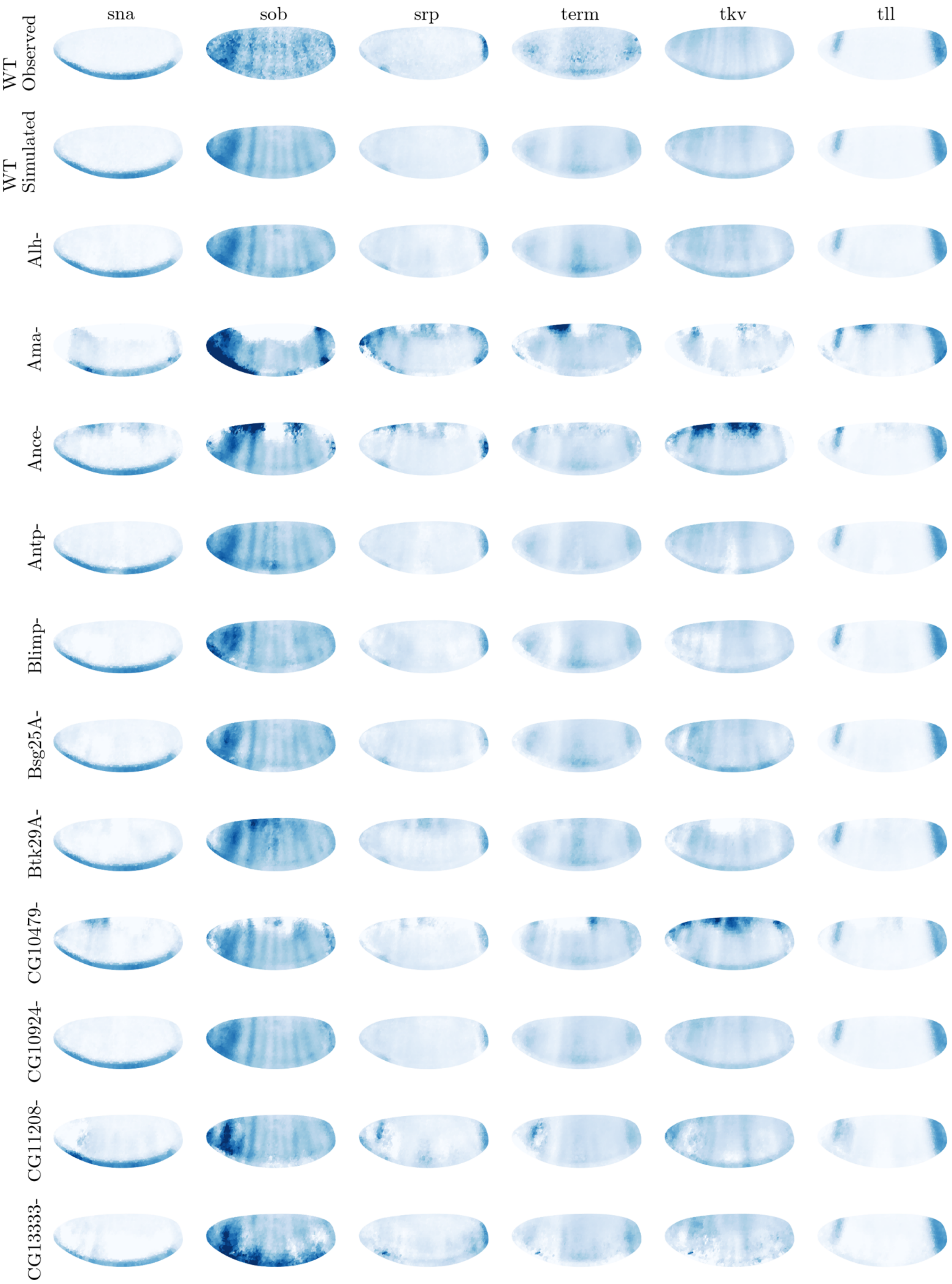
sna
sob
srp
term
tkv
tll
WT Observed
WT Simulated
Alh-
Ama-
Ance-
Antp-
Blimp-
Bsg25A-
Btk29A-
CG10479-
CG10924-
CG11208-
CG13333-

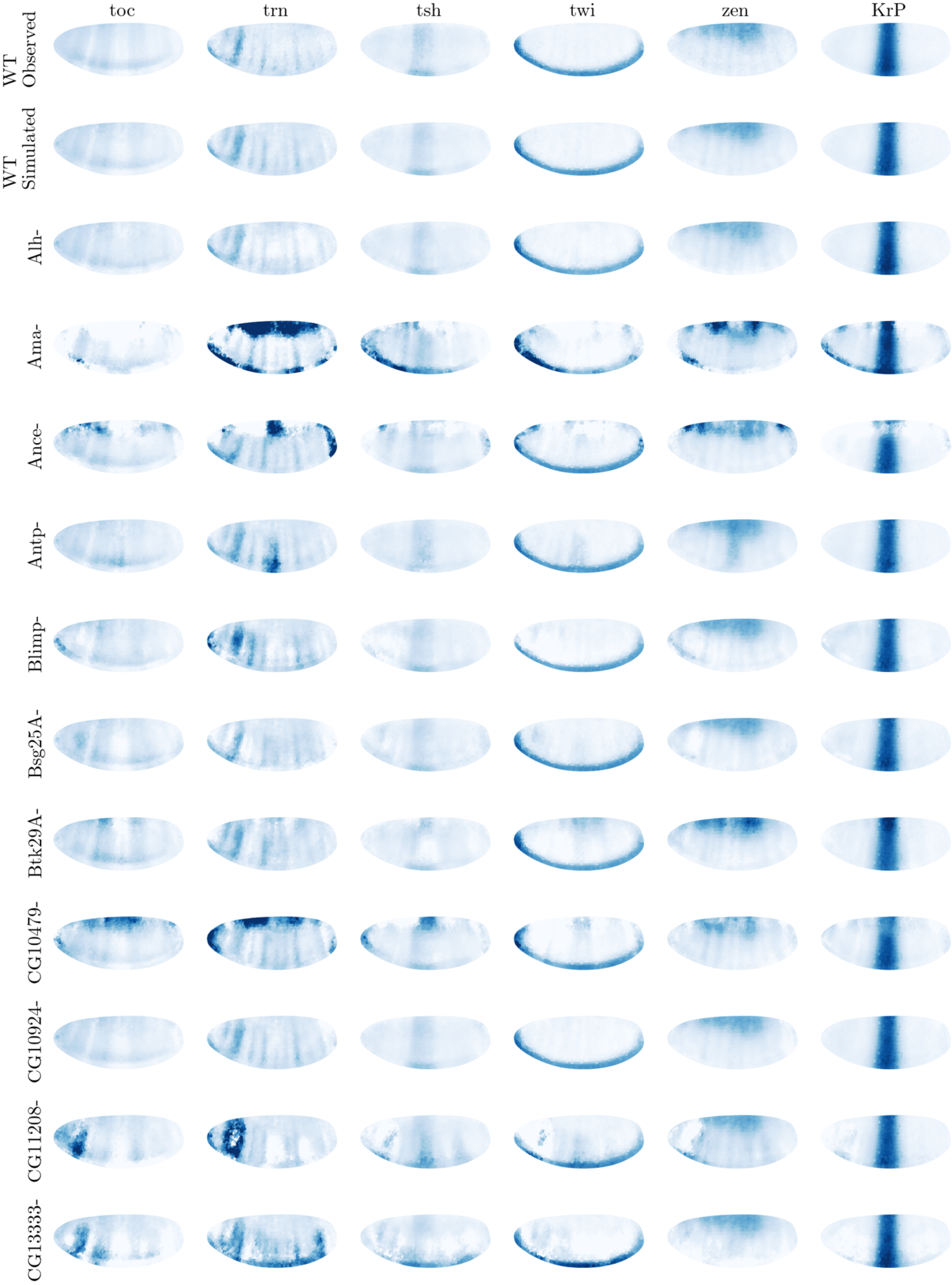

toc
trn
tsh
twi
zen
KrP
WT Observed
WT Simulated
Alh-
Ama-
Ance-
Antp-
Blimp-
Bsg25A-
Btk29A-
CG10479-
CG10924-
CG11208-
CG13333-

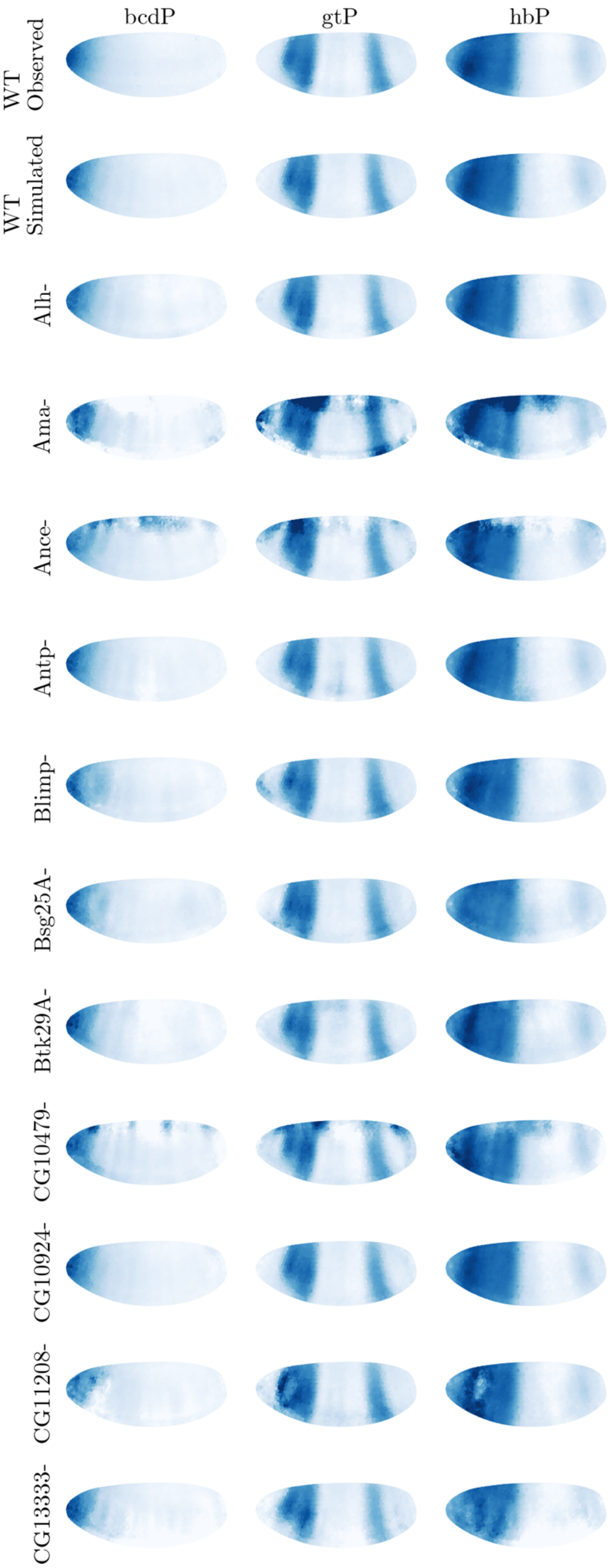
bcdP
gtP
hbP
WT Observed
WT Simulated
Alh-
Ama-
Ance-
Antp-
Blimp-
Bsg25A-
Btk29A-
CG10479-
CG10924-
CG11208-
CG13333-

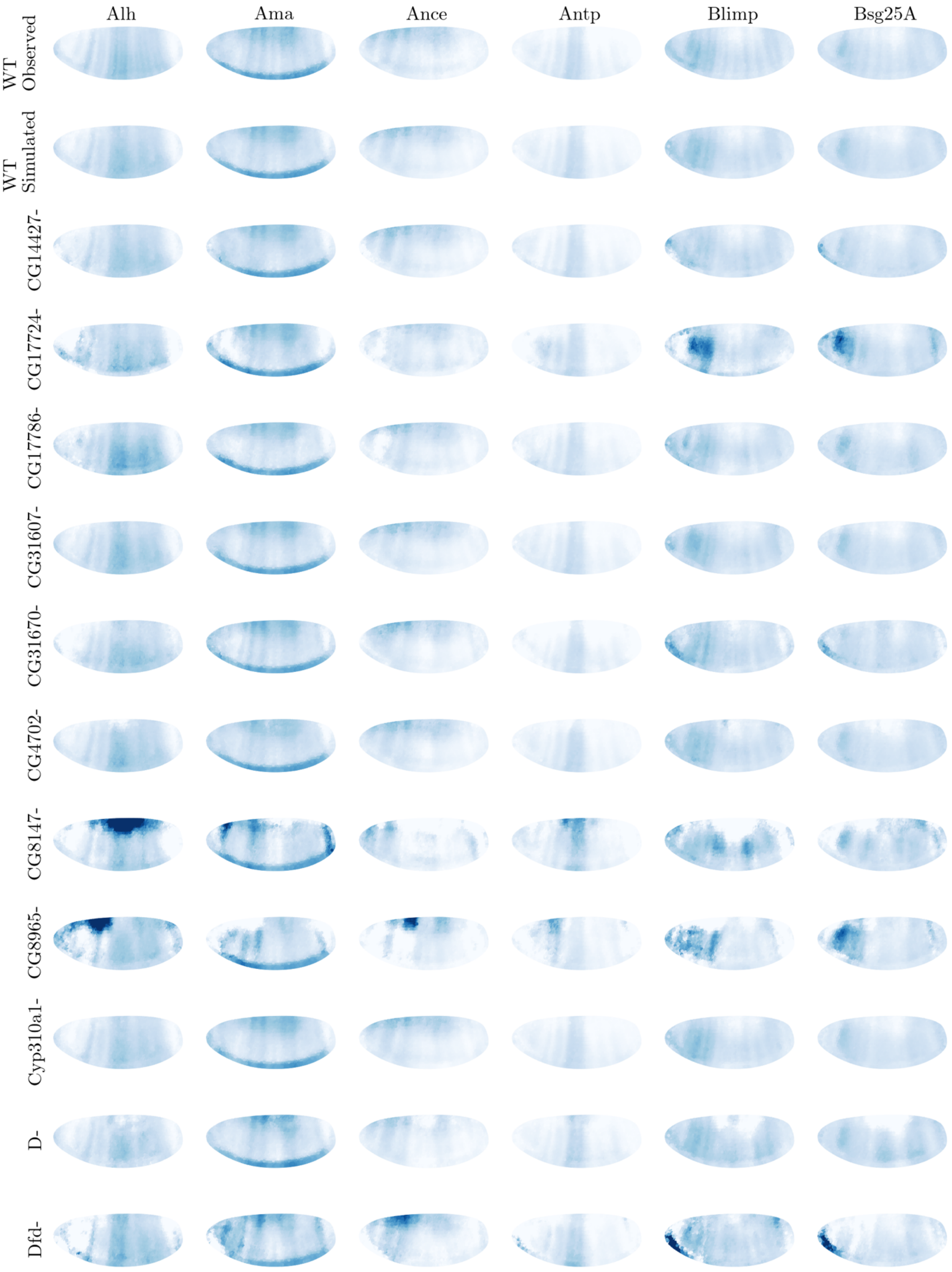
Alh
Ama
Ance
Antp
Blimp
Bsg25A
WT Observed
WT Simulated
CG14427-
CG17724-
CG17786-
CG31607-
CG31670-
CG4702-
CG8147-
CG8965-
Cyp310a1-
D-
Dfd-

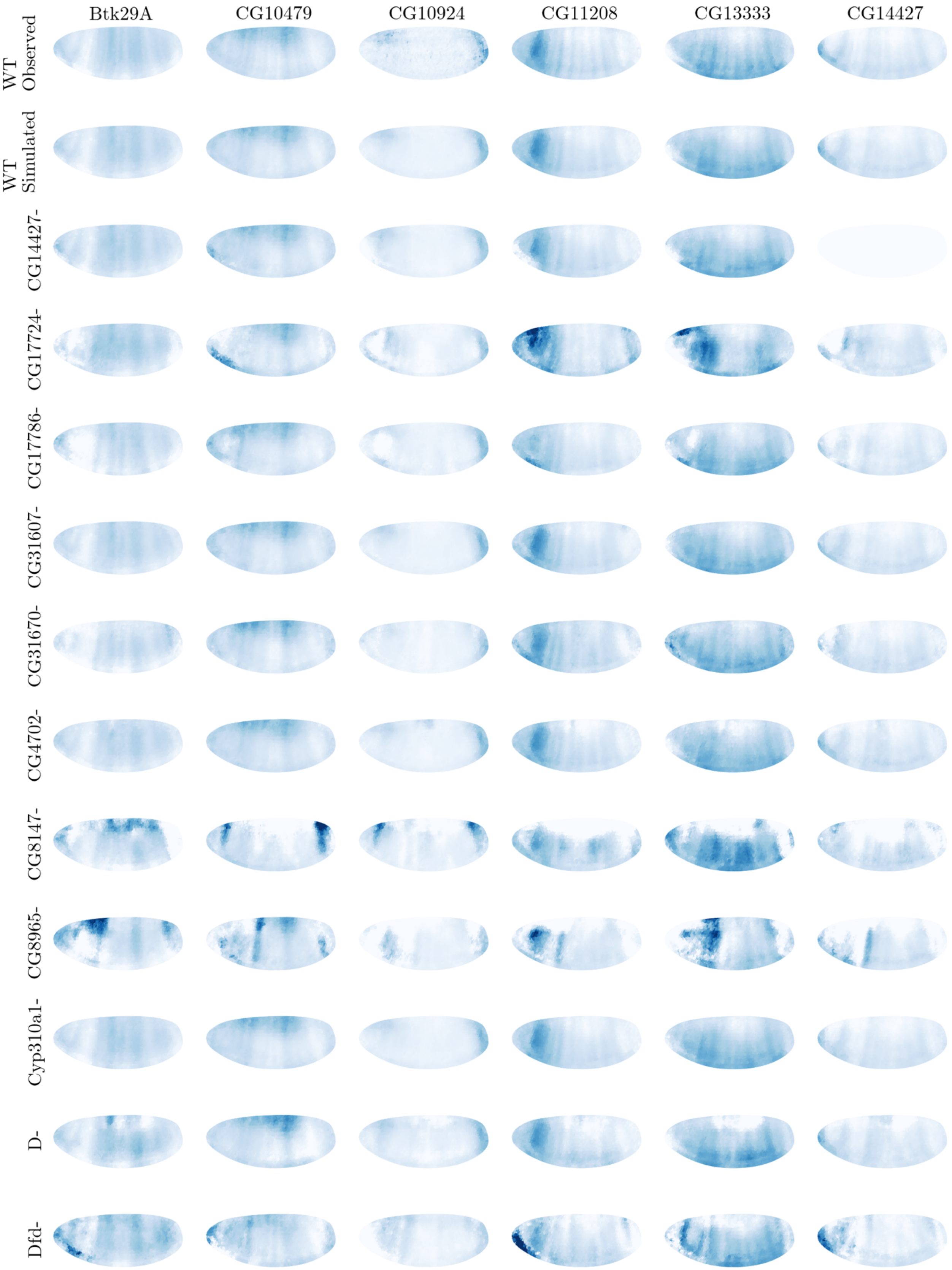
Btk29A
CG10479
CG10924
CG11208
CG13333
CG14427
WT Observed
WT Simulated
CG14427-
CG17724-
CG17786-
CG31607-
CG31670-
CG4702-
CG8147-
CG8965-
Cyp310a1-
D-
Dfd-

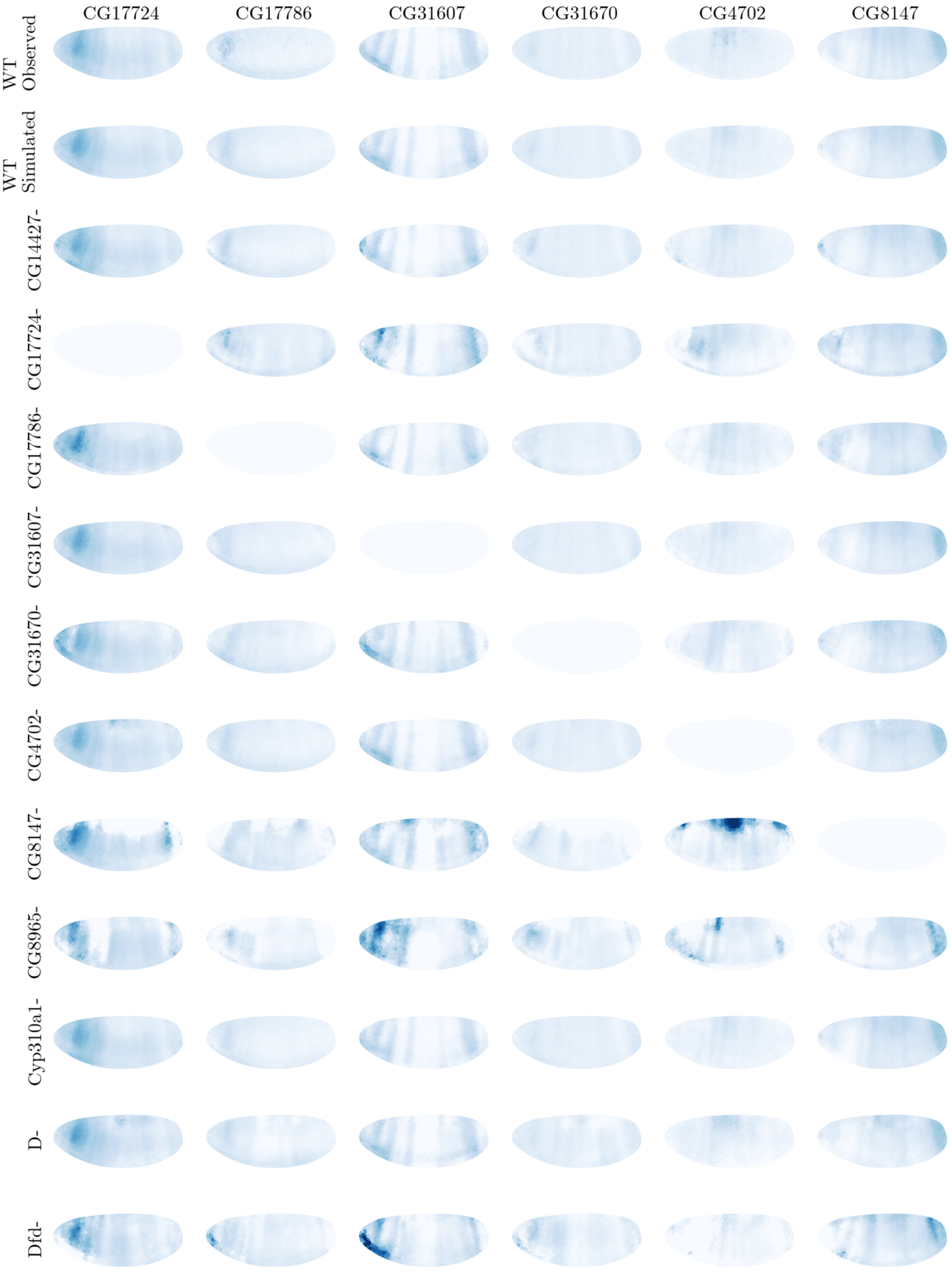
CG17724
CG17786
CG31607
CG31670
CG4702
CG8147
WT Observed
WT Simulated
CG14427-
CG17724-
CG17786-
CG31607-
CG31670-
CG4702-
CG8147-
CG8965-
Cyp310a1-
D-
Dfd-

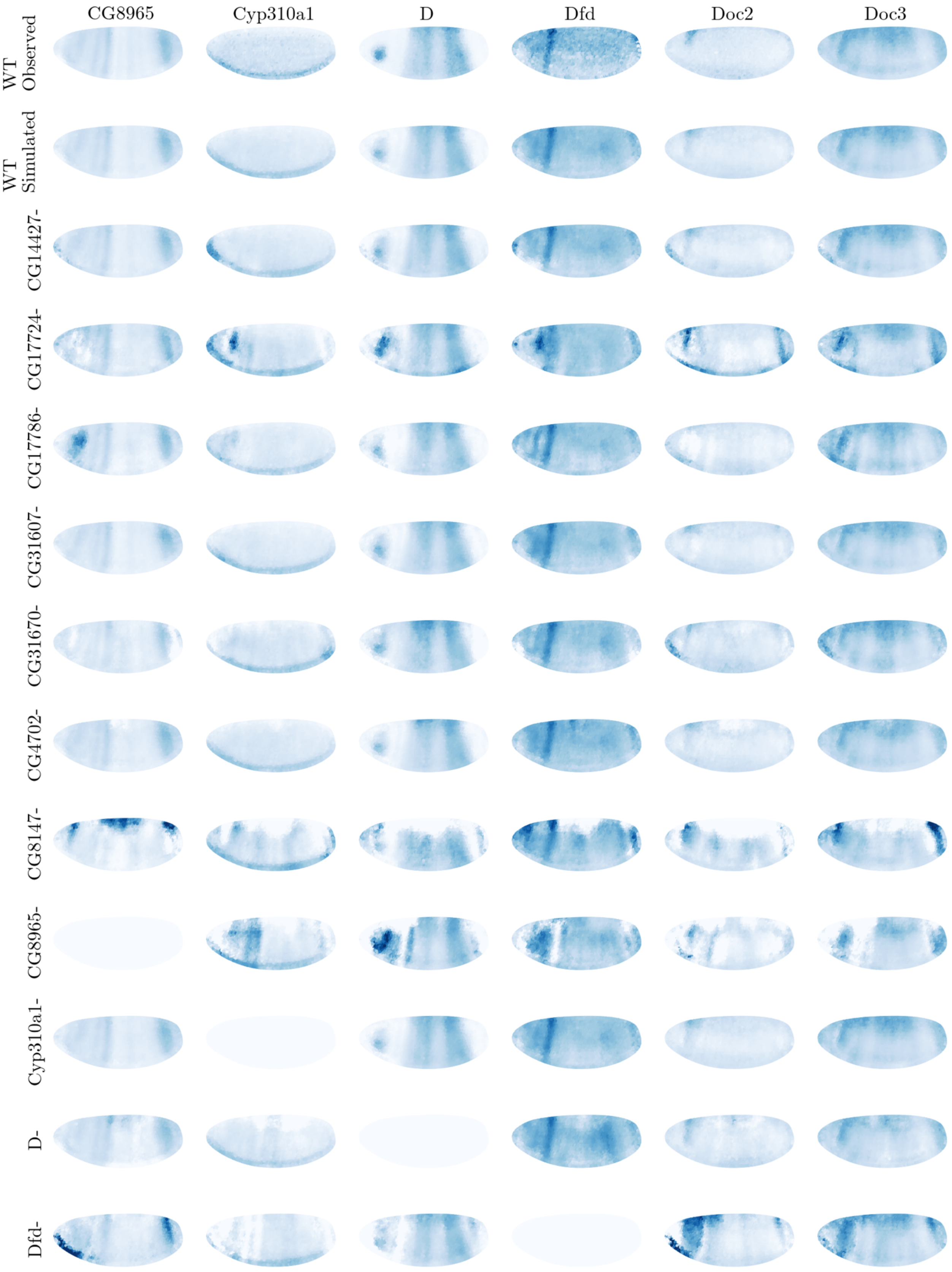

CG8965
Cyp310a1
D
Dfd
Doc2
Doc3
WT Observed
WT Simulated
CG14427-
CG17724-
CG17786-
CG31607-
CG31670-
CG4702-
CG8147-
CG8965-
Cyp310a1-
D-
Dfd-

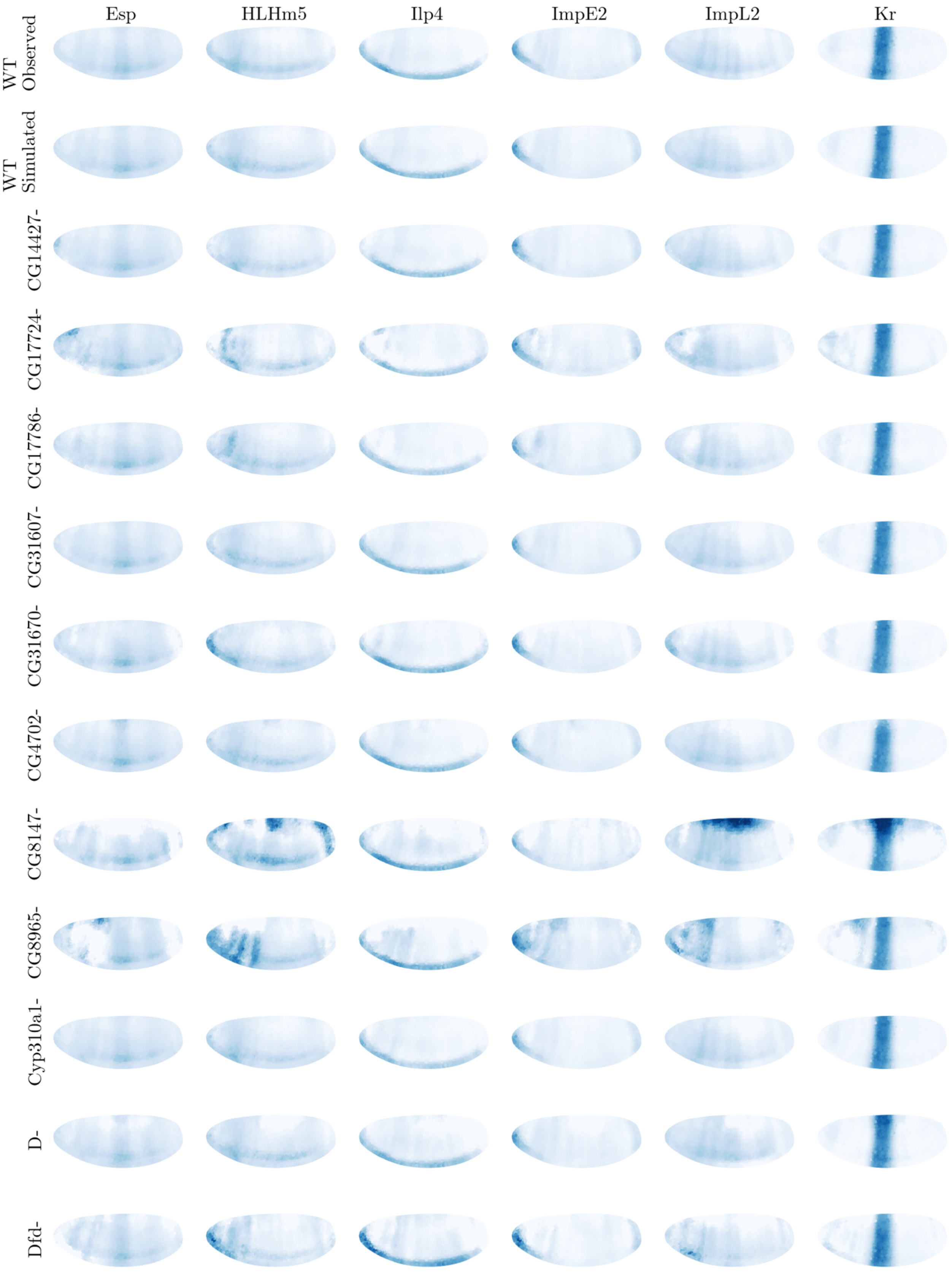
Esp
HLHm5
Ilp4
ImpE2
ImpL2
Kr
WT Observed
WT Simulated
CG14427-
CG17724-
CG17786-
CG31607-
CG31670-
CG4702-
CG8147-
CG8965-
Cyp310a1-
D-
Dfd-

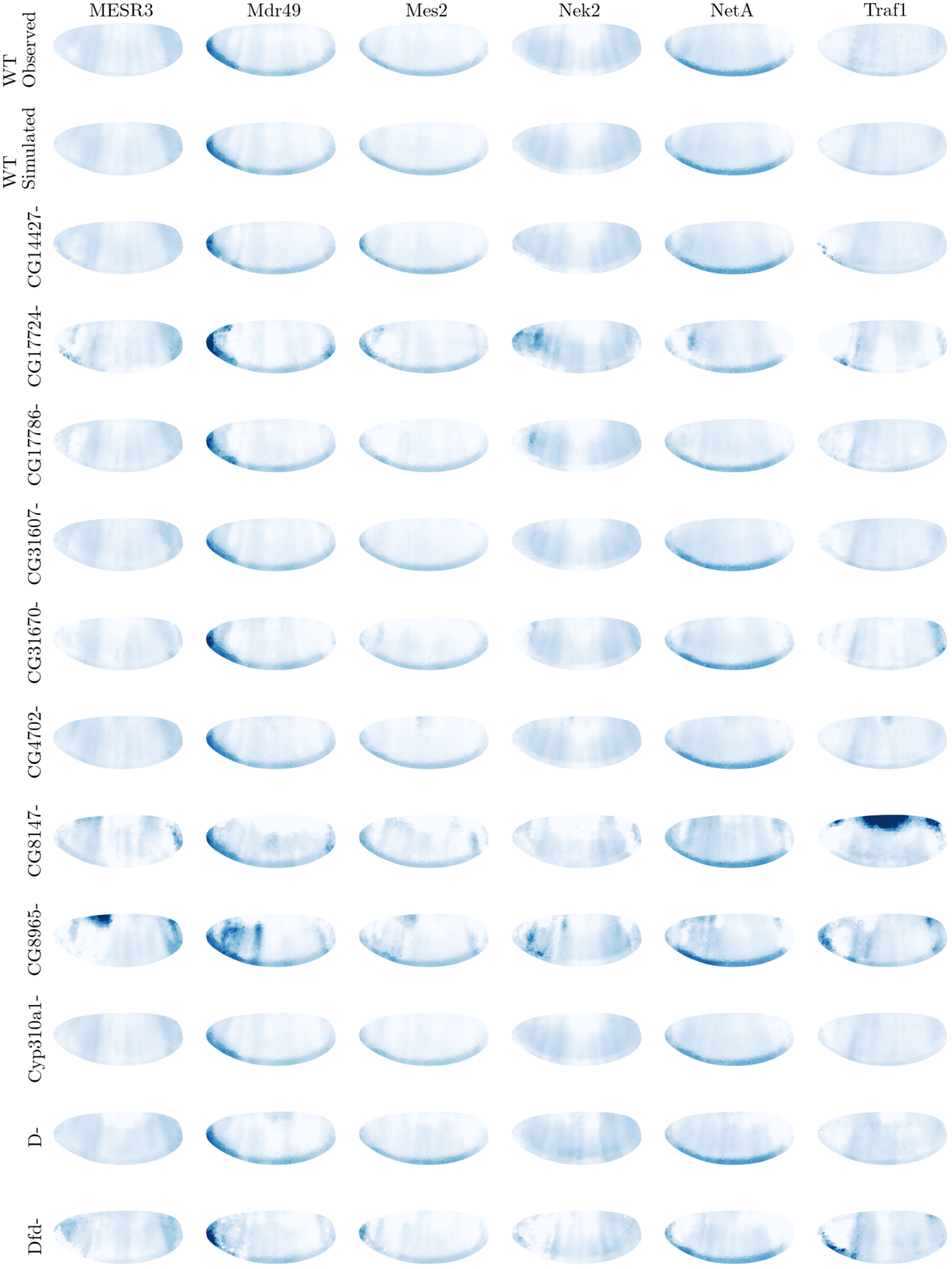
MESR3
Mdr49
Mes2
Nek2
NetA
Traf1
WT Observed
WT Simulated
CG14427-
CG17724-
CG17786-
CG31607-
CG31670-
CG4702-
CG8147-
CG8965-
Cyp310a1-
D-
Dfd-

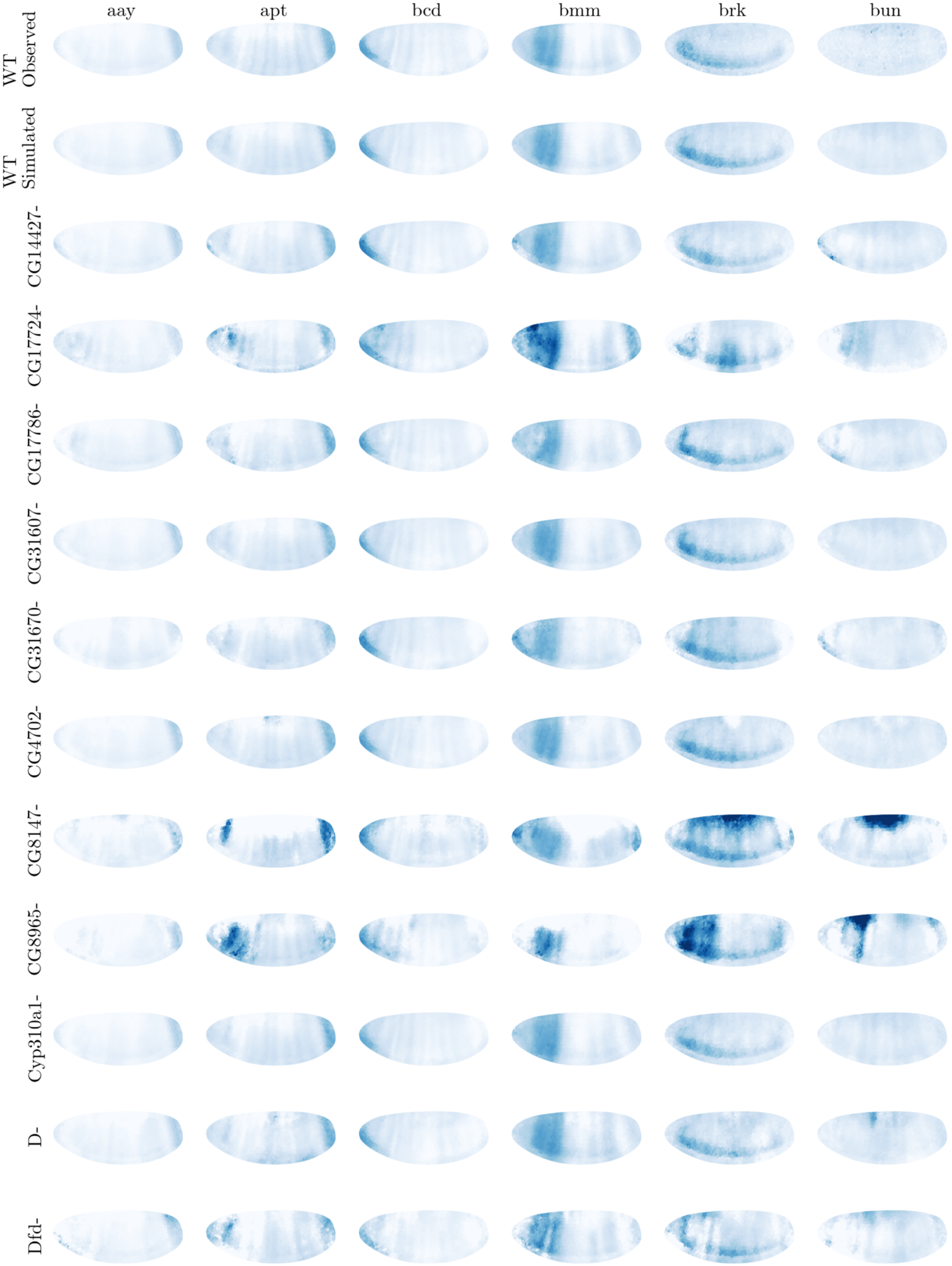
aay
apt
bcd
bmm
brk
bun
WT Observed
WT Simulated
CG14427-
CG17724-
CG17786-
CG31607-
CG31670-
CG4702-
CG8147-
CG8965-
Cyp310a1-
D-
Dfd-

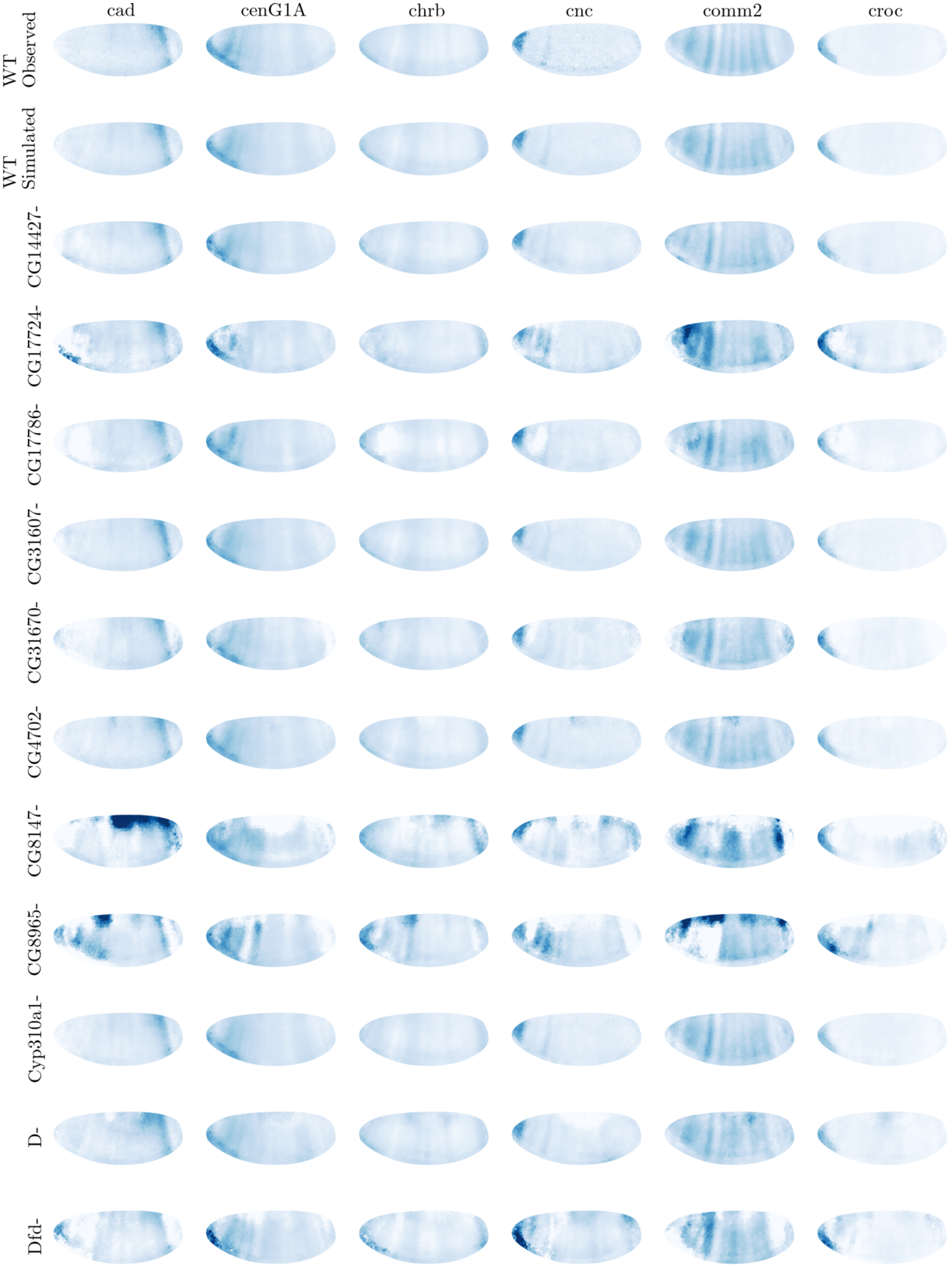
cad
cenG1A
chrb
cnc
comm2
croc
WT Observed
WT Simulated
CG14427-
CG17724-
CG17786-
CG31607-
CG31670-
CG4702-
CG8147-
CG8965-
Cyp310a1-
D-
Dfd-

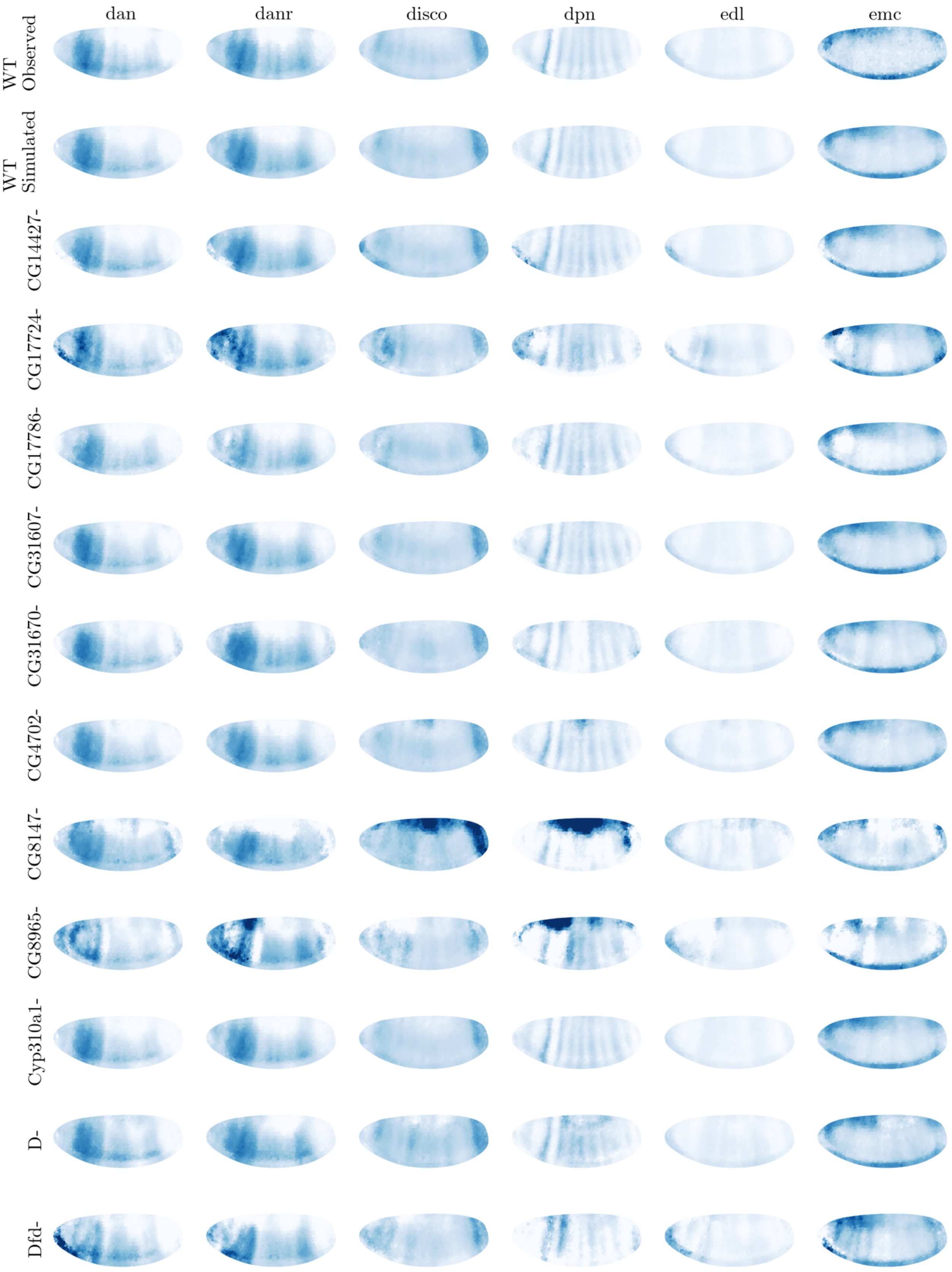
dan
danr
disco
dpn
edl
emc
WT Observed
WT Simulated
CG14427-
CG17724-
CG17786-
CG31607-
CG31670-
CG4702-
CG8147-
CG8965-
Cyp310a1-
D-
Dfd-

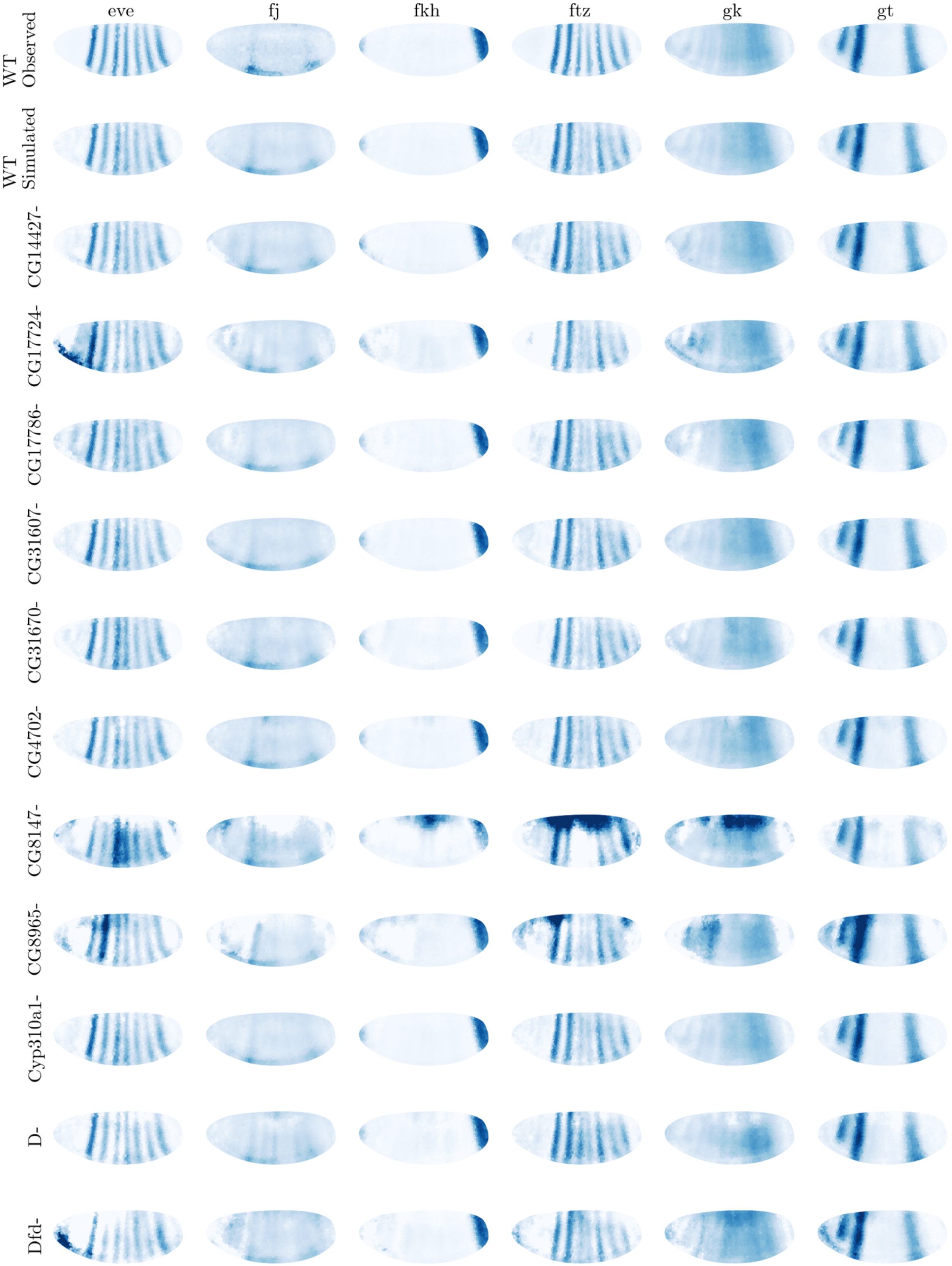
eve
fj
fkh
ftz
gk
gt
WT Observed
WT Simulated
CG14427-
CG17724-
CG17786-
CG31607-
CG31670-
CG4702-
CG8147-
CG8965-
Cyp310a1-
D-
Dfd-

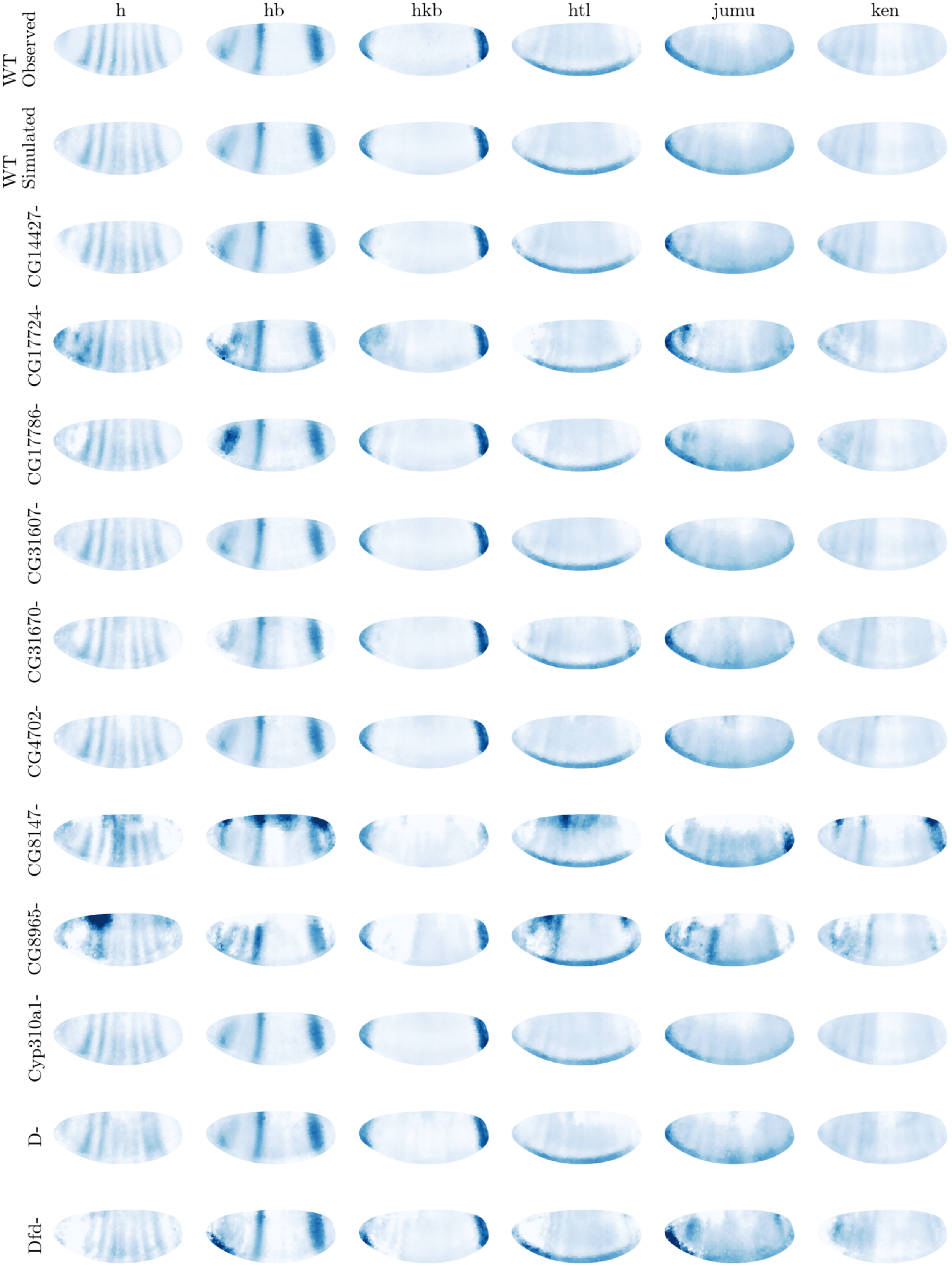
h
hb
hkb
htl
jumu
ken
WT Observed
WT Simulated
CG14427-
CG17724-
CG17786-
CG31607-
CG31670-
CG4702-
CG8147-
CG8965-
Cyp310a1-
D-
Dfd-

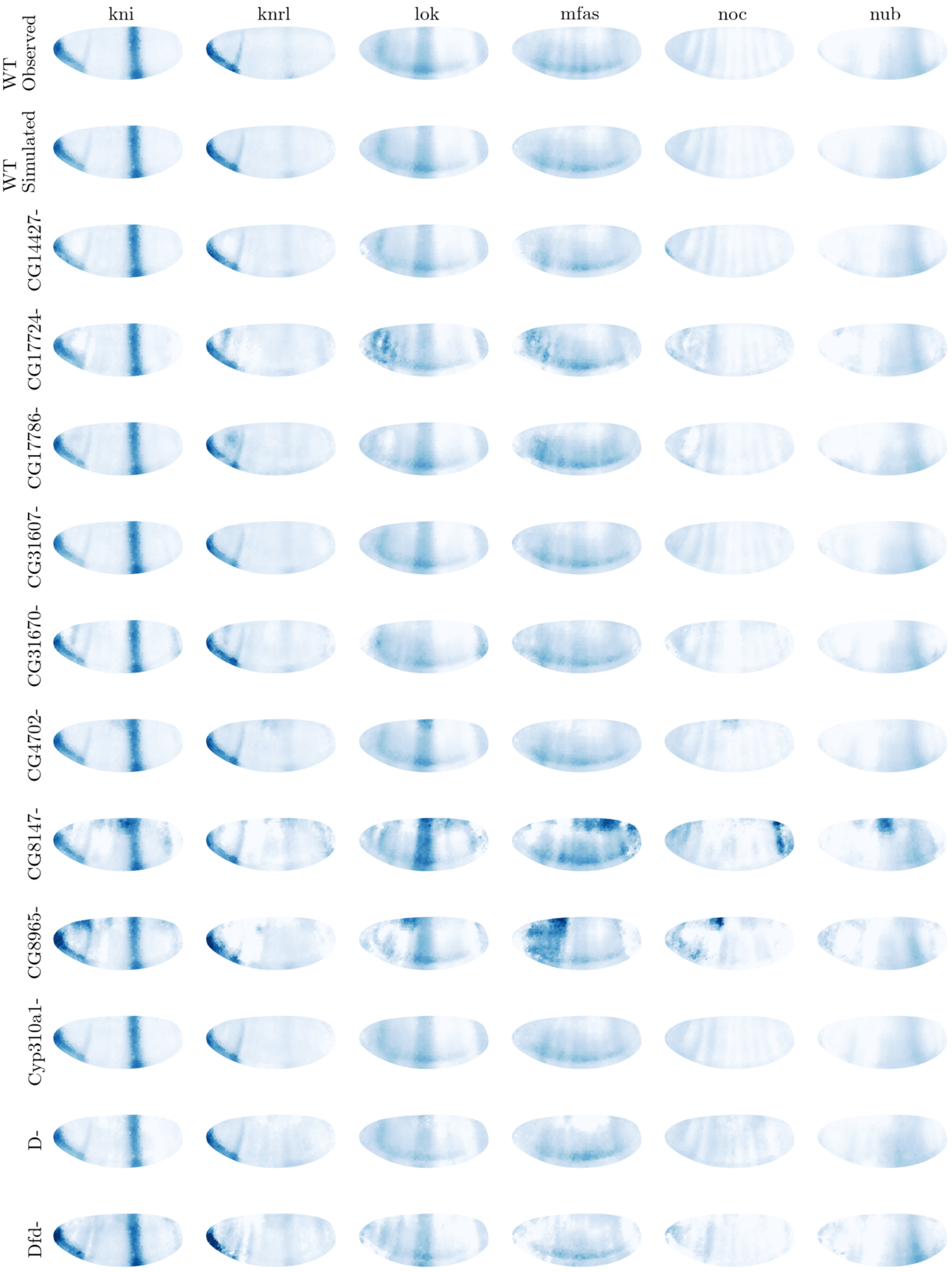
kni
knrl
lok
mfas
noc
nub
WT Observed
WT Simulated
CG14427-
CG17724-
CG17786-
CG31607-
CG31670-
CG4702-
CG8147-
CG8965-
Cyp310a1-
D-
Dfd-

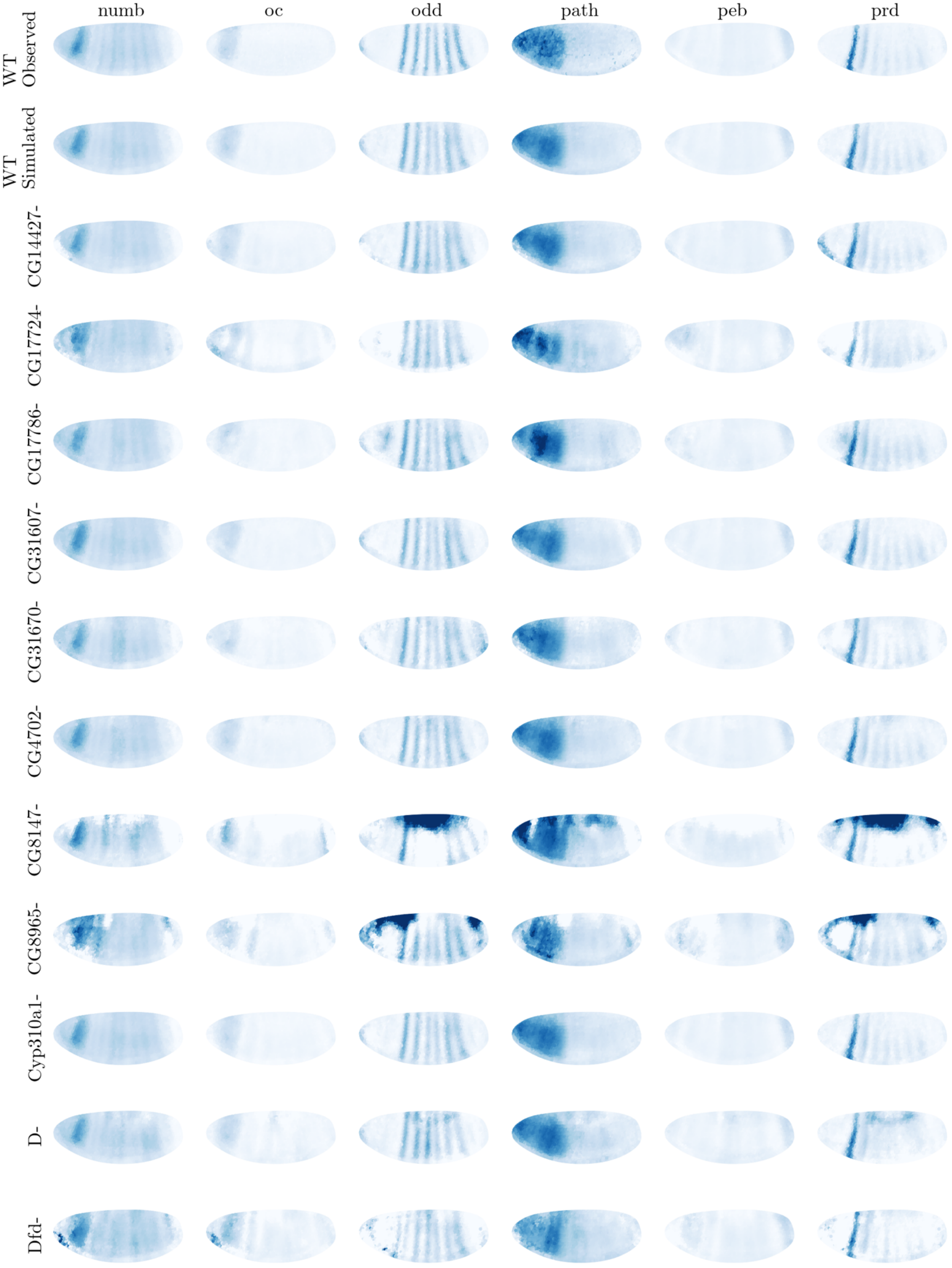
numb
oc
odd
path
peb
prd
WT Observed
WT Simulated
CG14427-
CG17724-
CG17786-
CG31607-
CG31670-
CG4702-
CG8147-
CG8965-
Cyp310a1-
D-
Dfd-

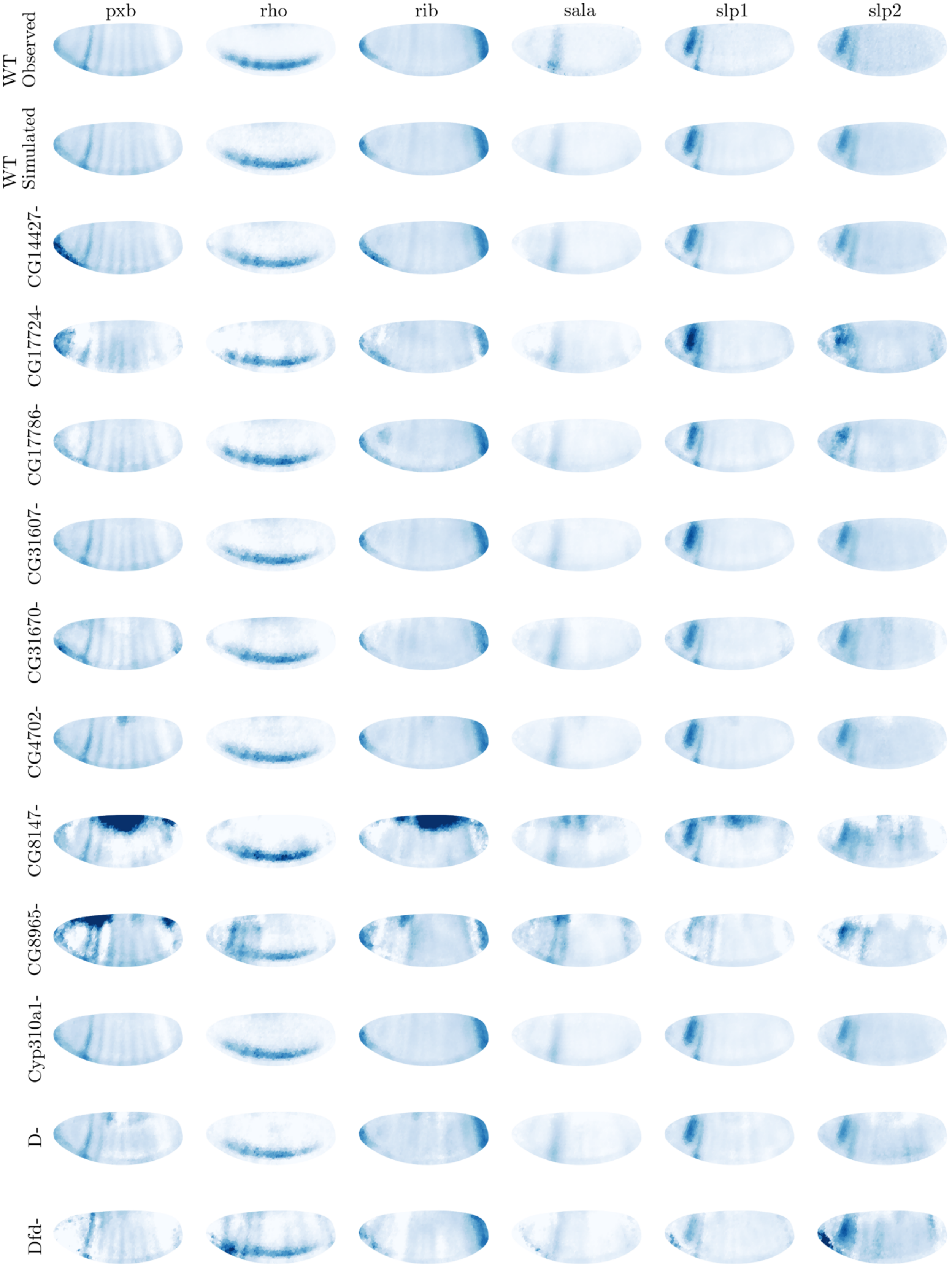
pxb
rho
rib
sala
slp1
slp2
WT Observed
WT Simulated
CG14427-
CG17724-
CG17786-
CG31607-
CG31670-
CG4702-
CG8147-
CG8965-
Cyp310a1-
D-
Dfd-

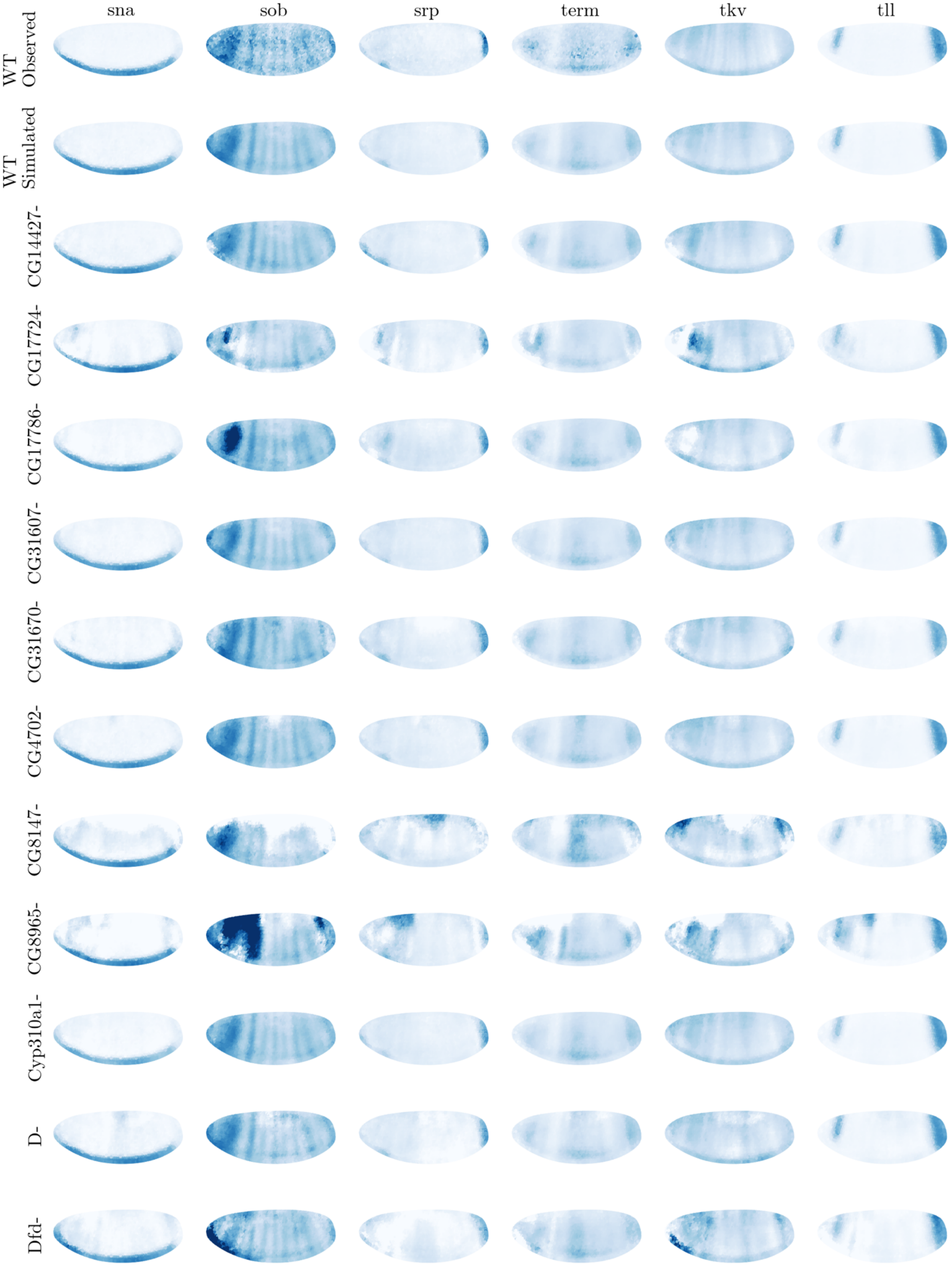
sna
sob
srp
term
tkv
tll
WT Observed
WT Simulated
CG14427-
CG17724-
CG17786-
CG31607-
CG31670-
CG4702-
CG8147-
CG8965-
Cyp310a1-
D-
Dfd-

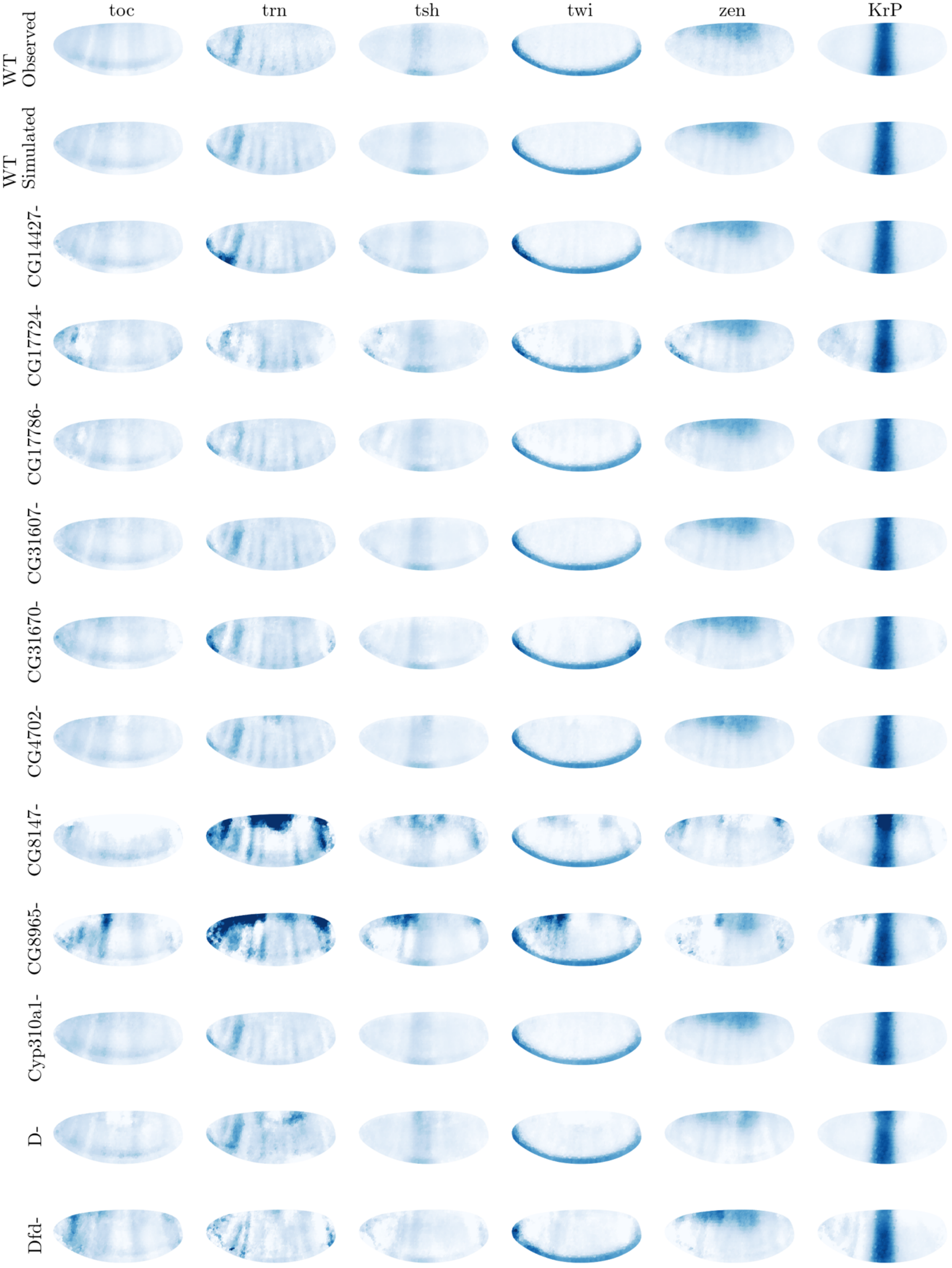
toc
trn
tsh
twi
zen
KrP
WT Observed
WT Simulated
CG14427-
CG17724-
CG17786-
CG31607-
CG31670-
CG4702-
CG8147-
CG8965-
Cyp310a1-
D-
Dfd-

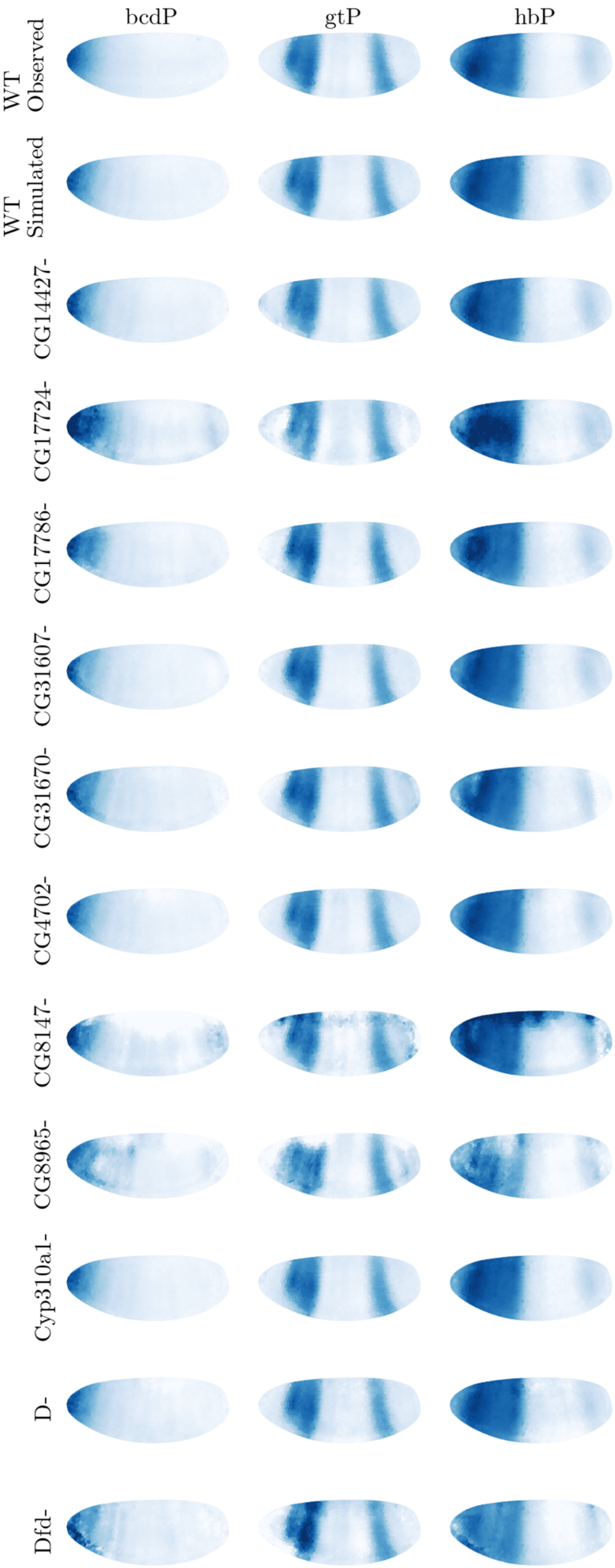
bcdP
gtP
hbP
WT Observed
WT Simulated
CG14427-
CG17724-
CG17786-
CG31607-
CG31670-
CG4702-
CG8147-
CG8965-
Cyp310a1-
D-
Dfd-

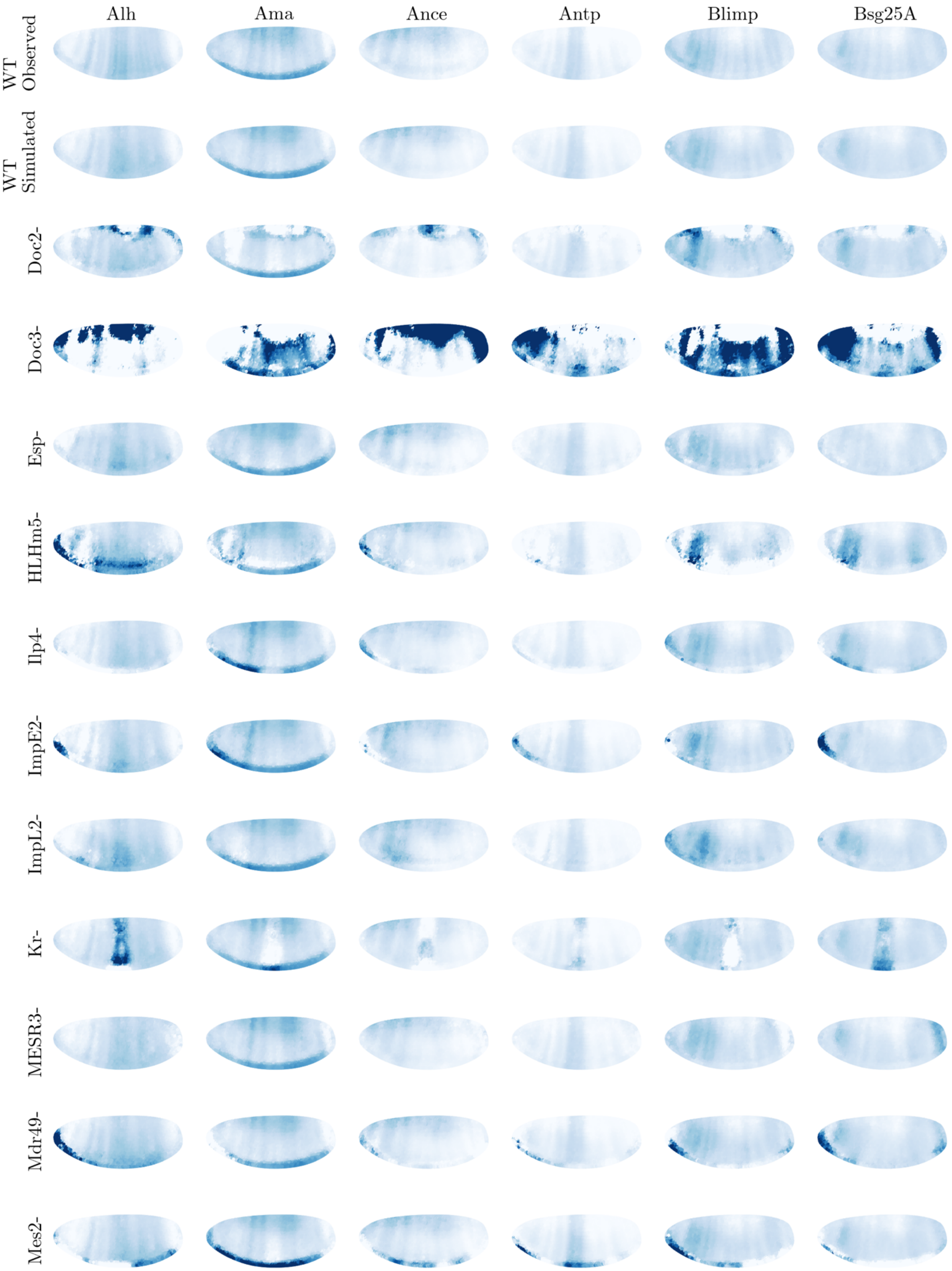
Alh
Ama
Ance
Antp
Blimp
Bsg25A
WT Observed
WT Simulated
Doc2-
Doc3-
Esp-
HLHm5-
Ilp4-
ImpE2-
ImpL2-
Kr-
MESR3-
Mdr49-
Mes2-

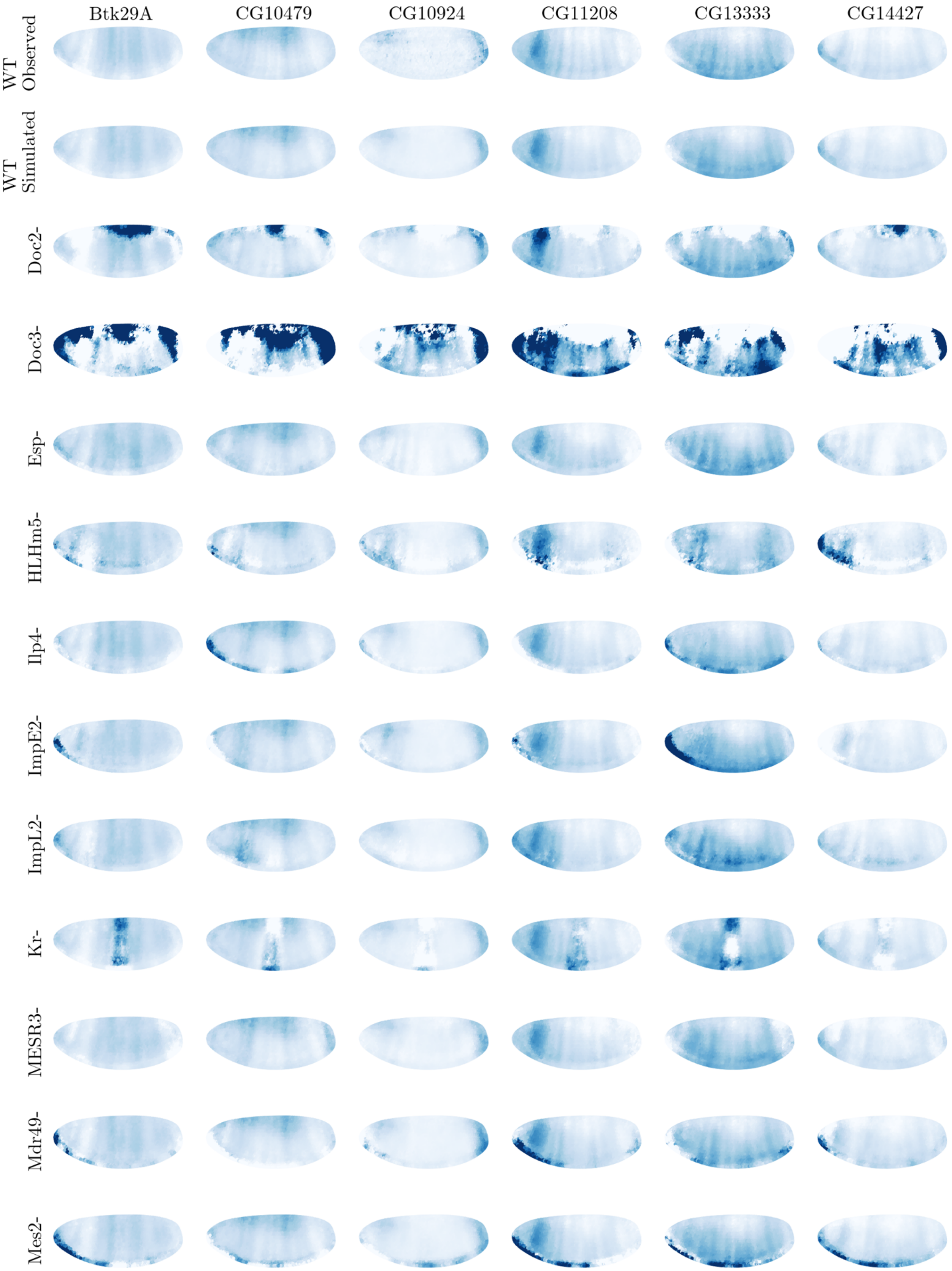
Btk29A
CG10479
CG10924
CG11208
CG13333
CG14427
WT Observed
WT Simulated
Doc2-
Doc3-
Esp-
HLHm5-
Ilp4-
ImpE2-
ImpL2-
Kr-
MESR3-
Mdr49-
Mes2-

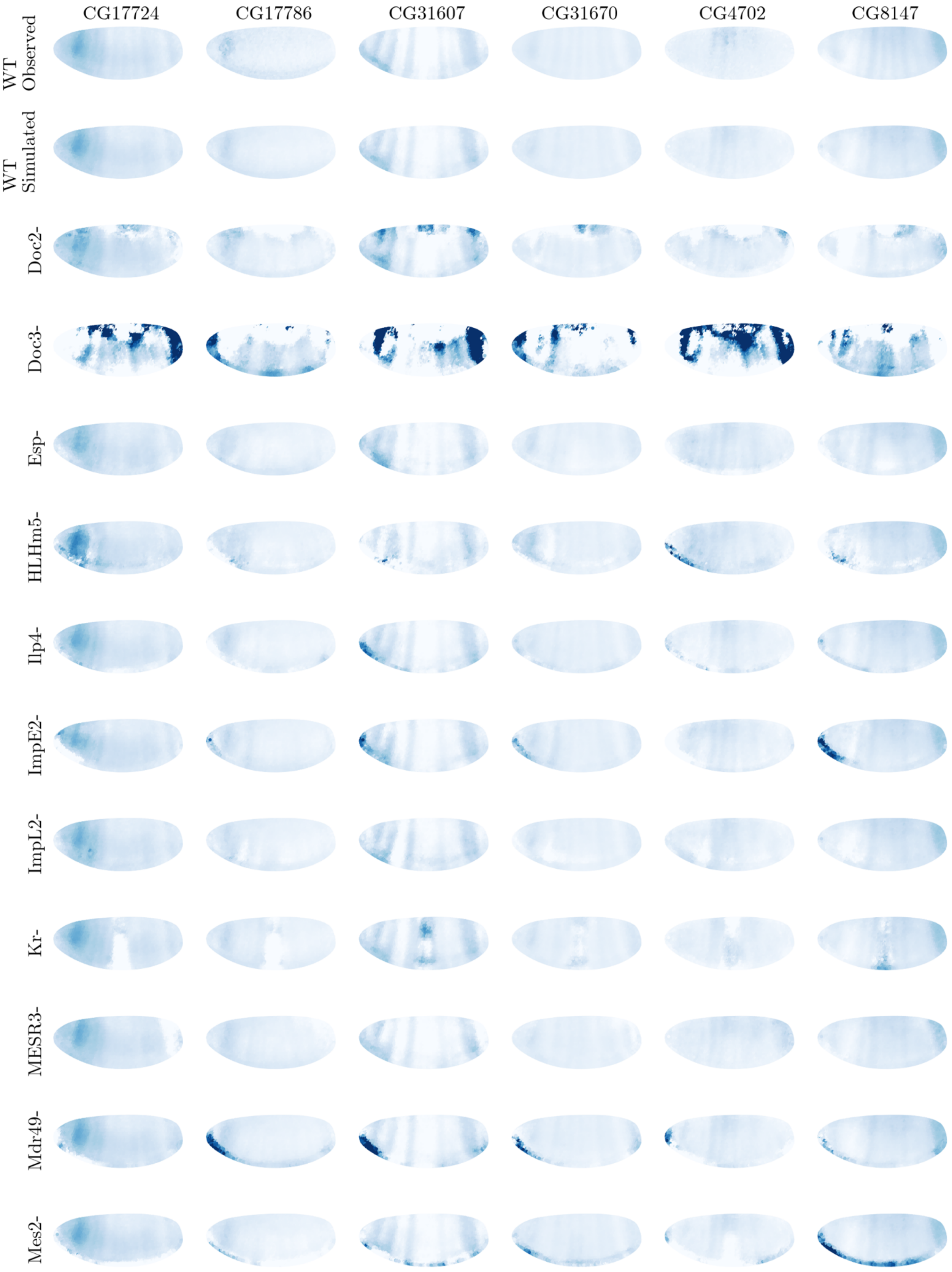
CG17724
CG17786
CG31607
CG31670
CG4702
CG8147
WT Observed
WT Simulated
Doc2-
Doc3-
Esp-
HLHm5-
Ilp4-
ImpE2-
ImpL2-
Kr-
MESR3-
Mdr49-
Mes2-

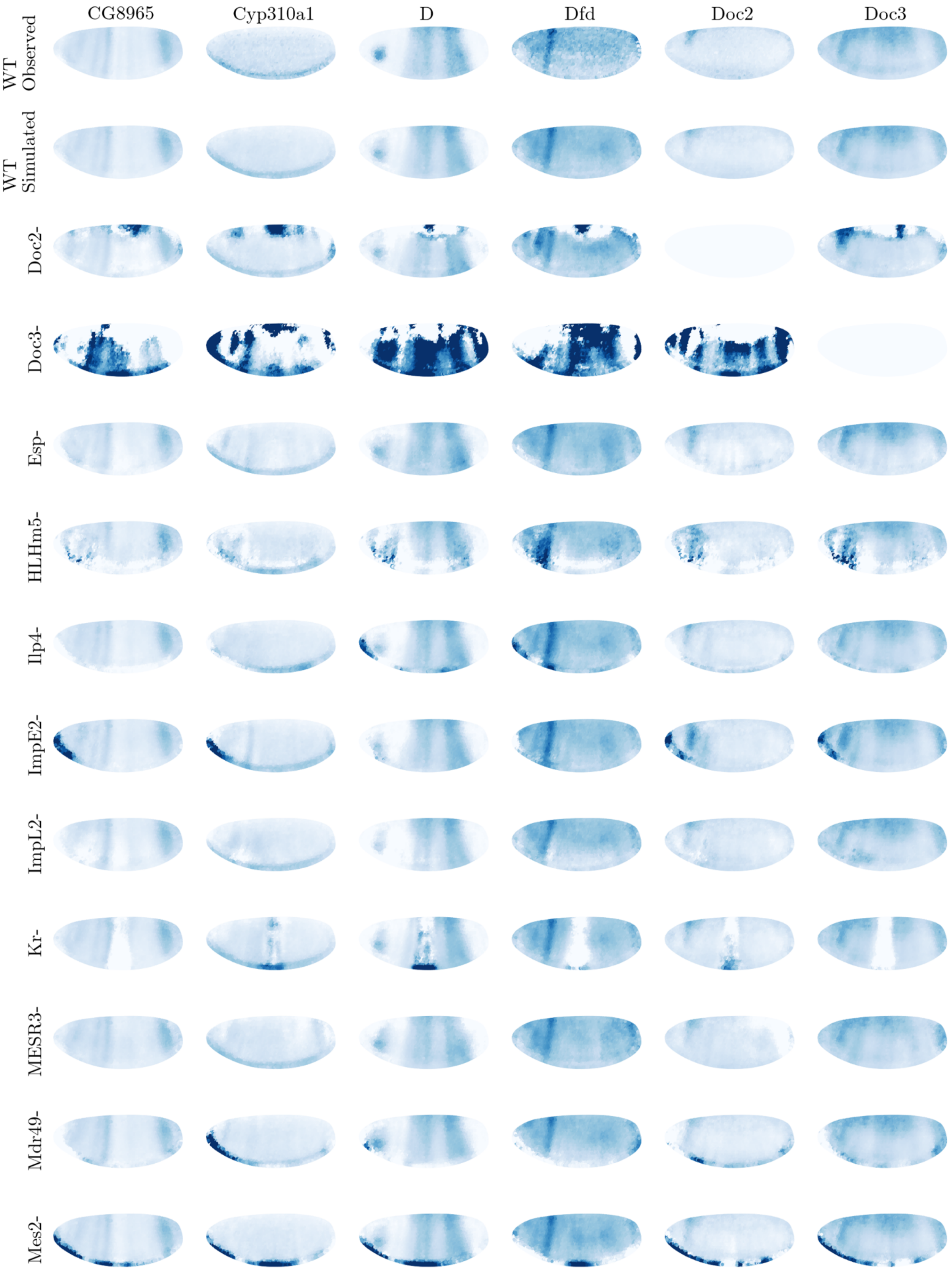
CG8965
Cyp310a1
D
Dfd
Doc2
Doc3
WT Observed
WT Simulated
Doc2-
Doc3-
Esp-
HLHm5-
Ilp4-
ImpE2-
ImpL2-
Kr-
MESR3-
Mdr49-
Mes2-

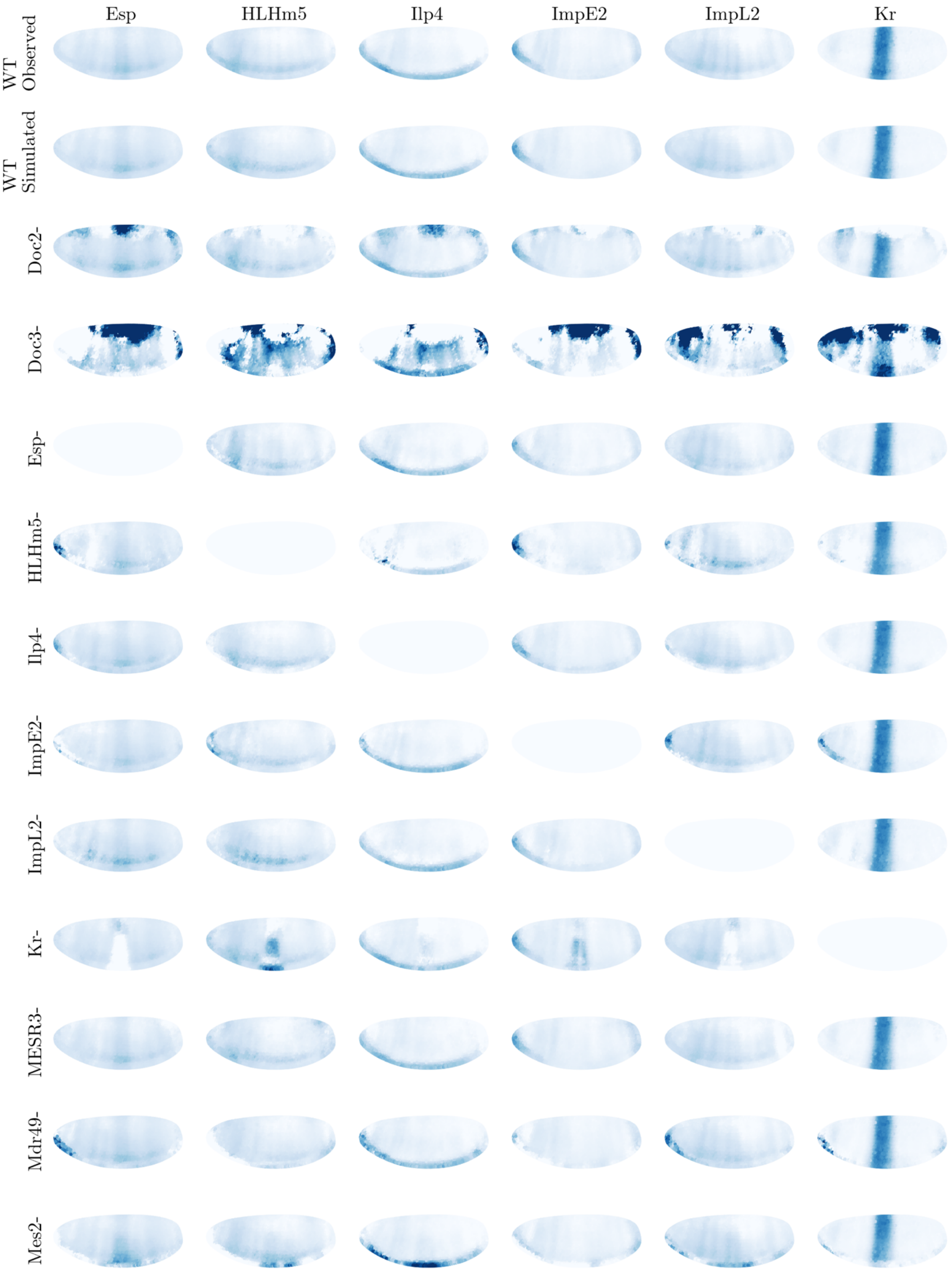

Esp
HLHm5
Ilp4
ImpE2
ImpL2
Kr
WT Observed
WT Simulated
Doc2-
Doc3-
Esp-
HLHm5-
Ilp4-
ImpE2-
ImpL2-
Kr-
MESR3-
Mdr49-
Mes2-

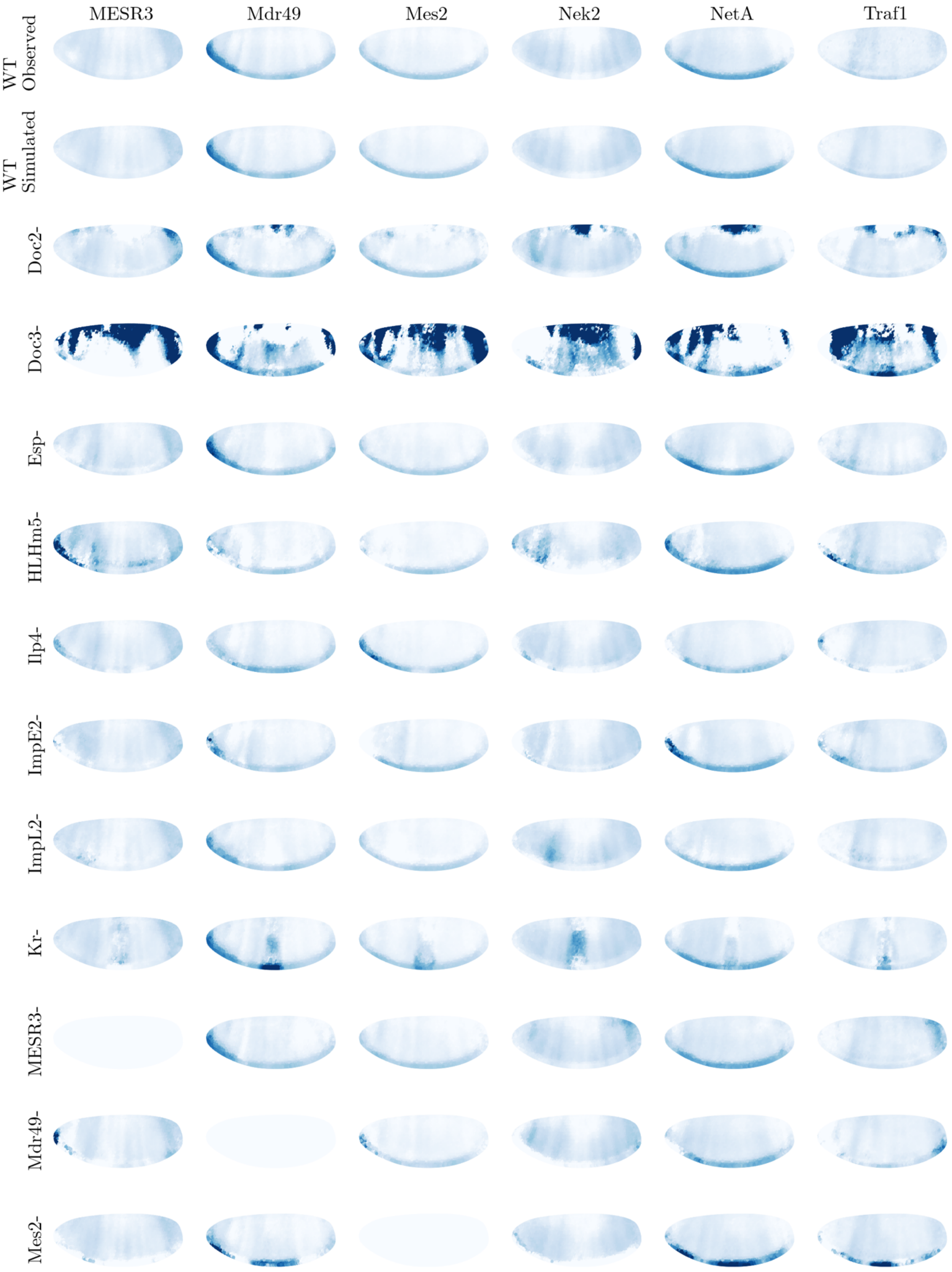

MESR3
Mdr49
Mes2
Nek2
NetA
Traf1
WT Observed
WT Simulated
Doc2-
Doc3-
Esp-
HLHm5-
Ilp4-
ImpE2-
ImpL2-
Kr-
MESR3-
Mdr49-
Mes2-

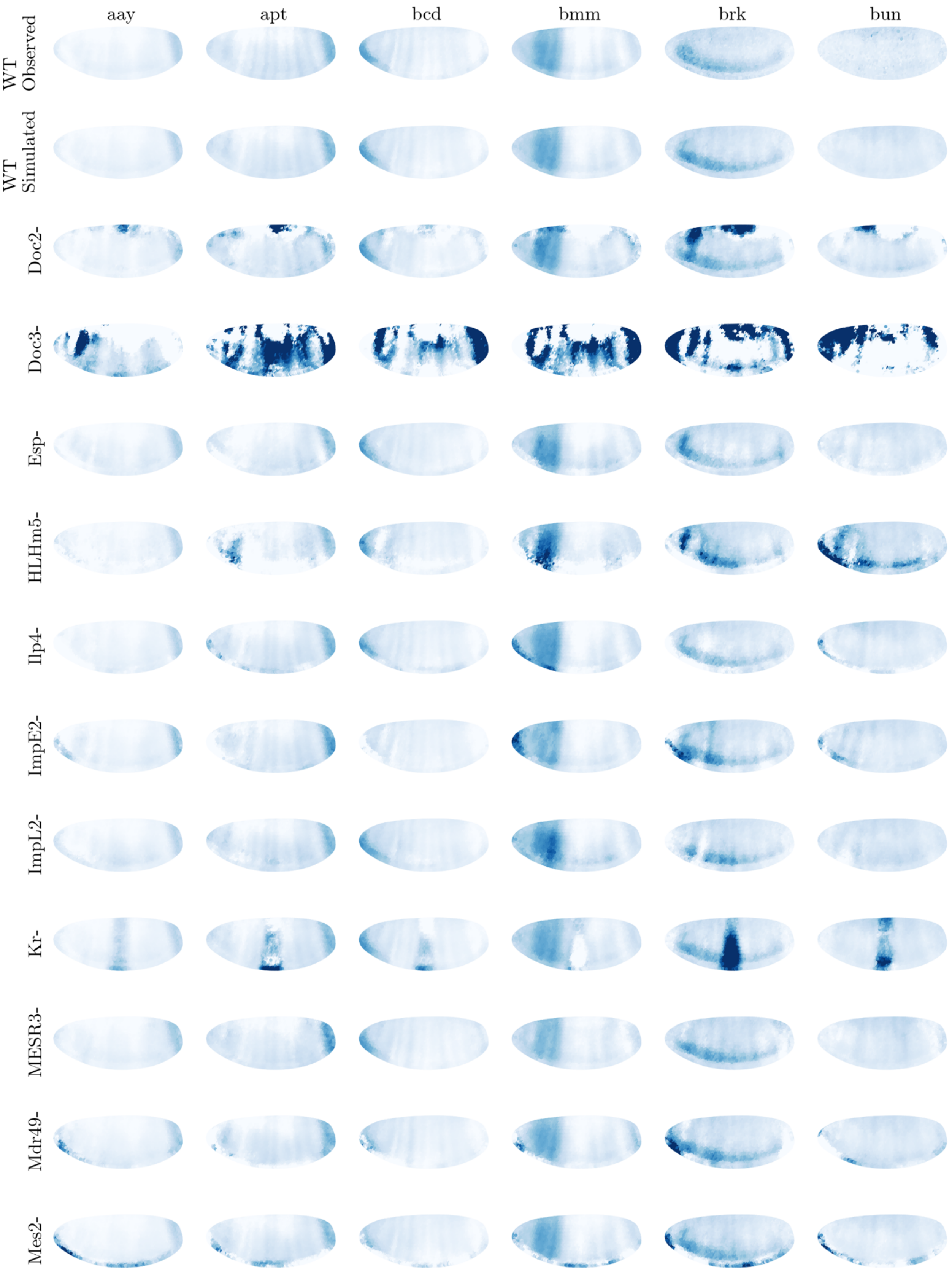
aay
apt
bcd
bmm
brk
bun
WT Observed
WT Simulated
Doc2-
Doc3-
Esp-
HLHm5-
Ilp4-
ImpE2-
ImpL2-
Kr-
MESR3-
Mdr49-
Mes2-

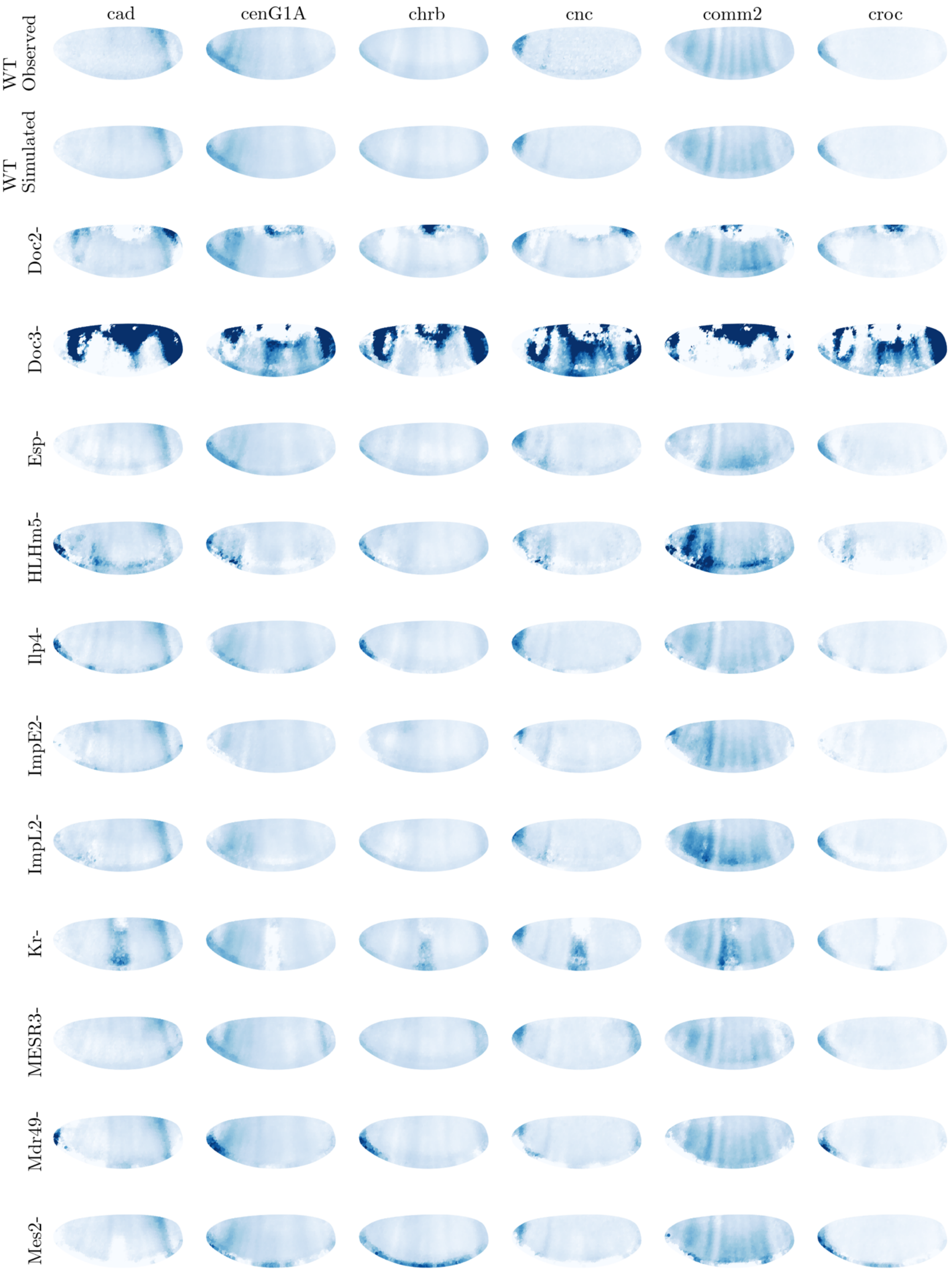
cad
cenG1A
chrb
cnc
comm2
croc
WT Observed
WT Simulated
Doc2-
Doc3-
Esp-
HLHm5-
Ilp4-
ImpE2-
ImpL2-
Kr-
MESR3-
Mdr49-
Mes2-

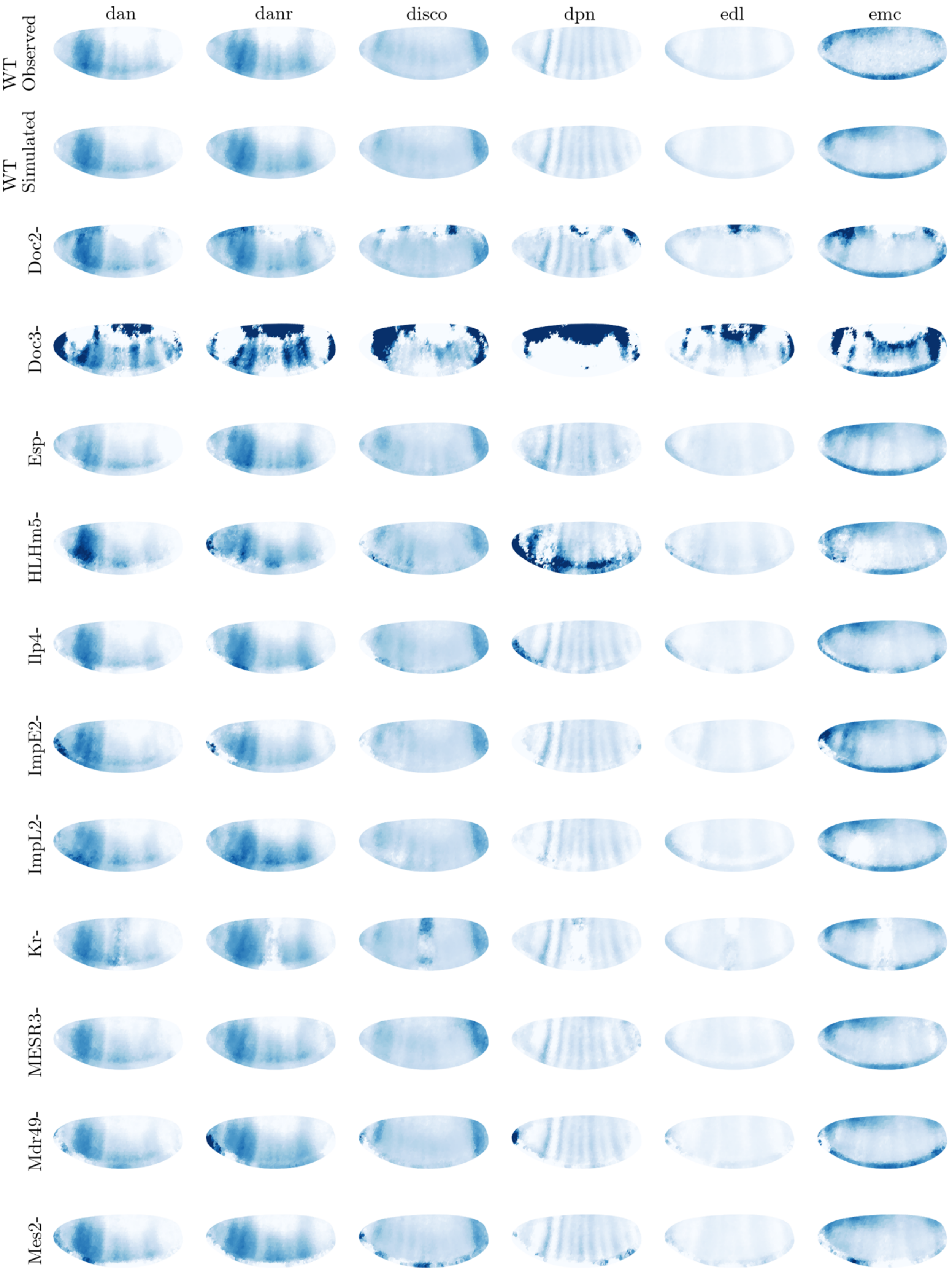
dan
danr
disco
dpn
edl
emc
WT Observed
WT Simulated
Doc2-
Doc3-
Esp-
HLHm5-
Ilp4-
ImpE2-
ImpL2-
Kr-
MESR3-
Mdr49-
Mes2-

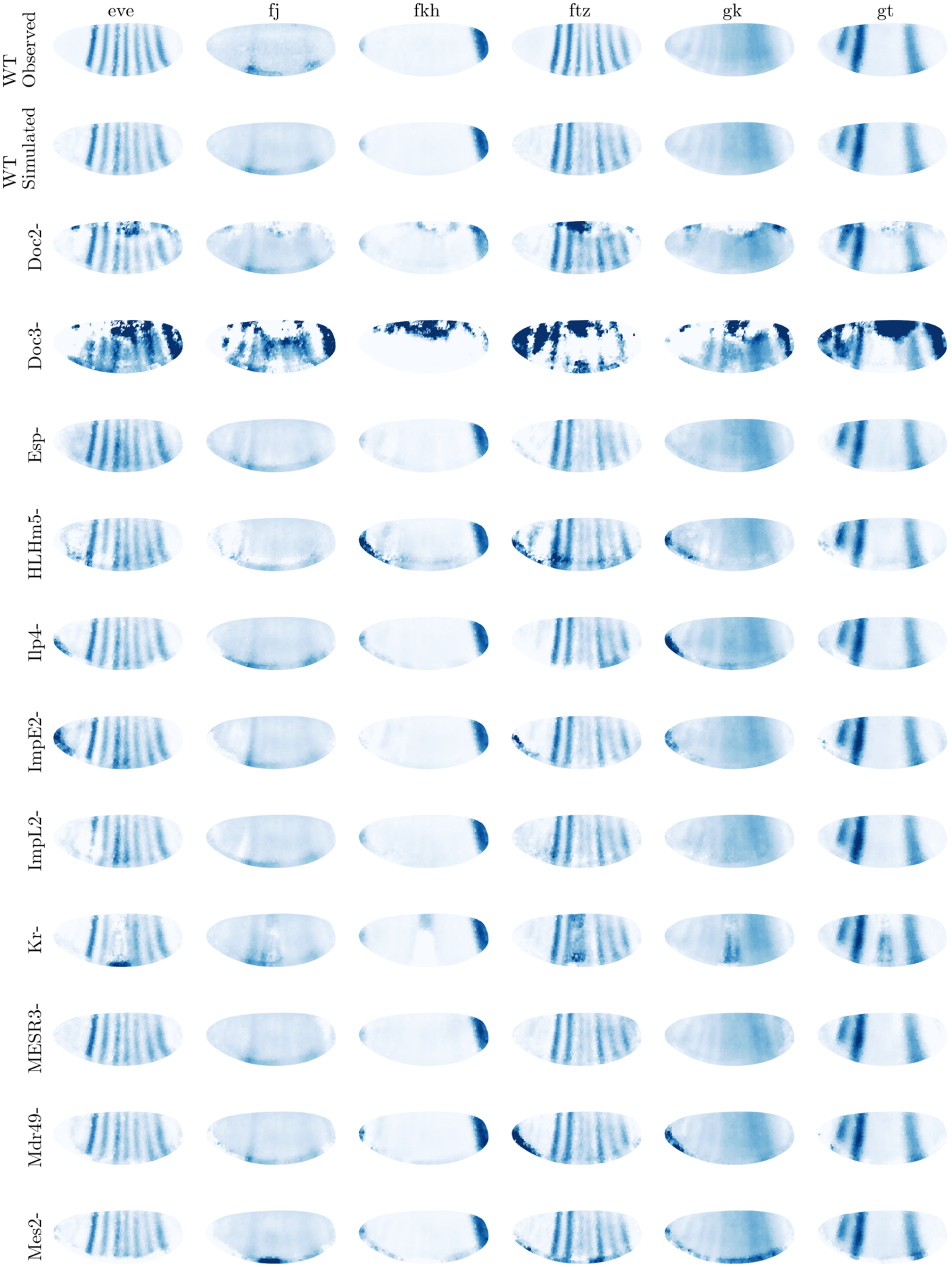

eve
fj
fkh
ftz
gk
gt
WT Observed
WT Simulated
Doc2-
Doc3-
Esp-
HLHm5-
Ilp4-
ImpE2-
ImpL2-
Kr-
MESR3-
Mdr49-
Mes2-

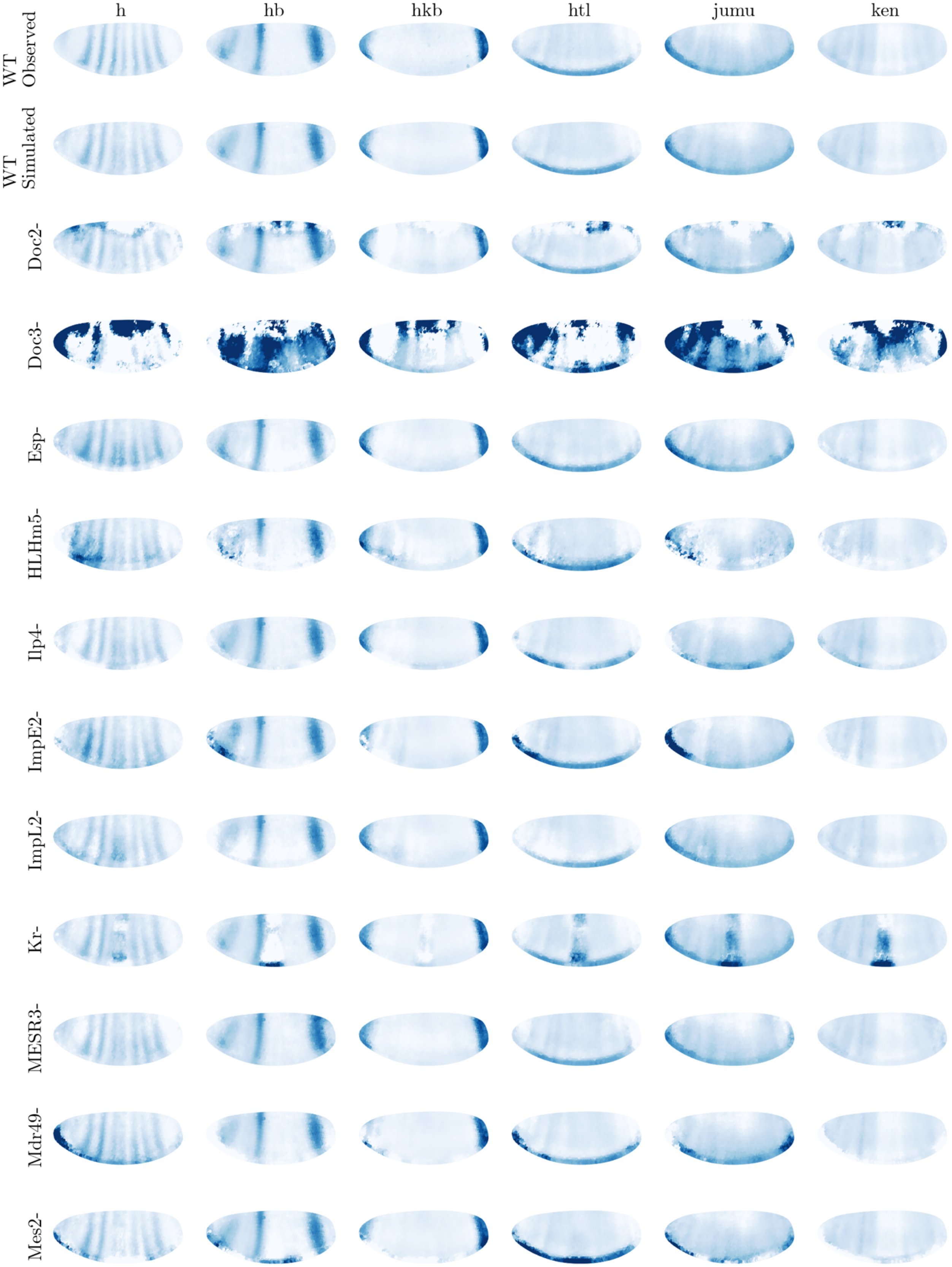
h
hb
hkb
htl
jumu
ken
WT Observed
WT Simulated
Doc2-
Doc3-
Esp-
HLHm5-
Ilp4-
ImpE2-
ImpL2-
Kr-
MESR3-
Mdr49-
Mes2-

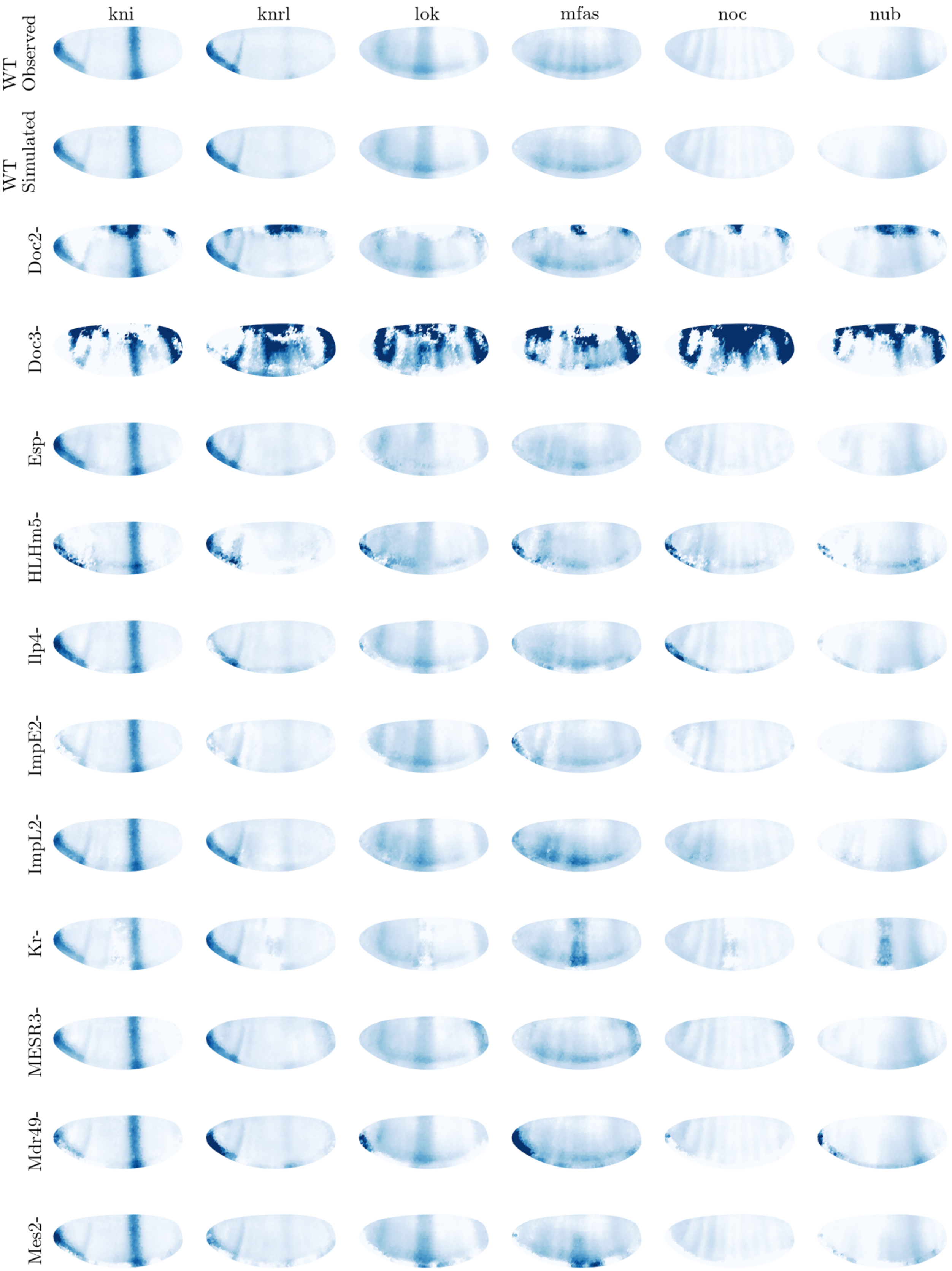

kni
knrl
lok
mfas
noc
nub
WT Observed
WT Simulated
Doc2-
Doc3-
Esp-
HLHm5-
Ilp4-
ImpE2-
ImpL2-
Kr-
MESR3-
Mdr49-
Mes2-

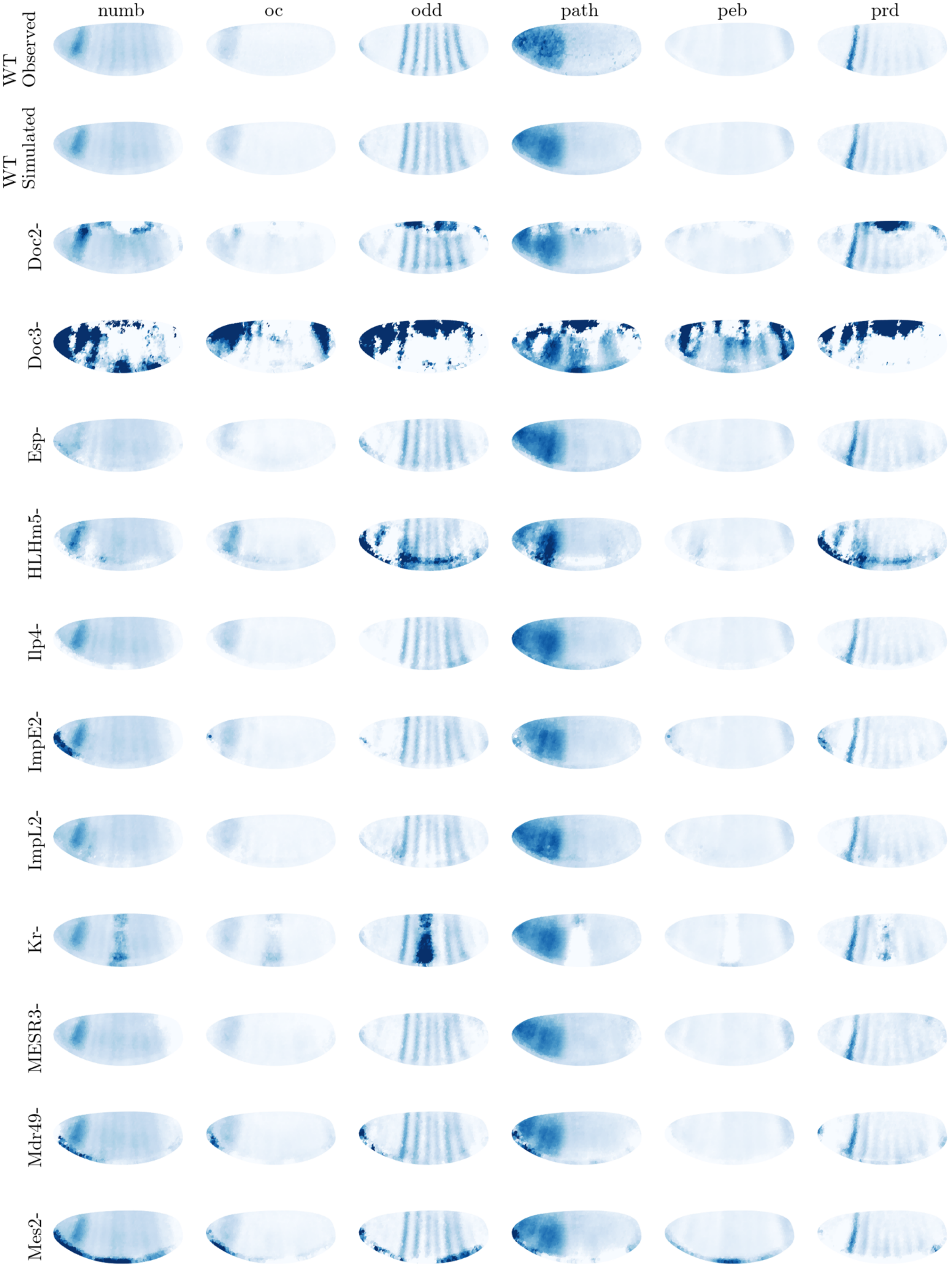

numb
oc
odd
path
peb
prd
WT Observed
WT Simulated
Doc2-
Doc3-
Esp-
HLHm5-
Ilp4-
ImpE2-
ImpL2-
Kr-
MESR3-
Mdr49-
Mes2-

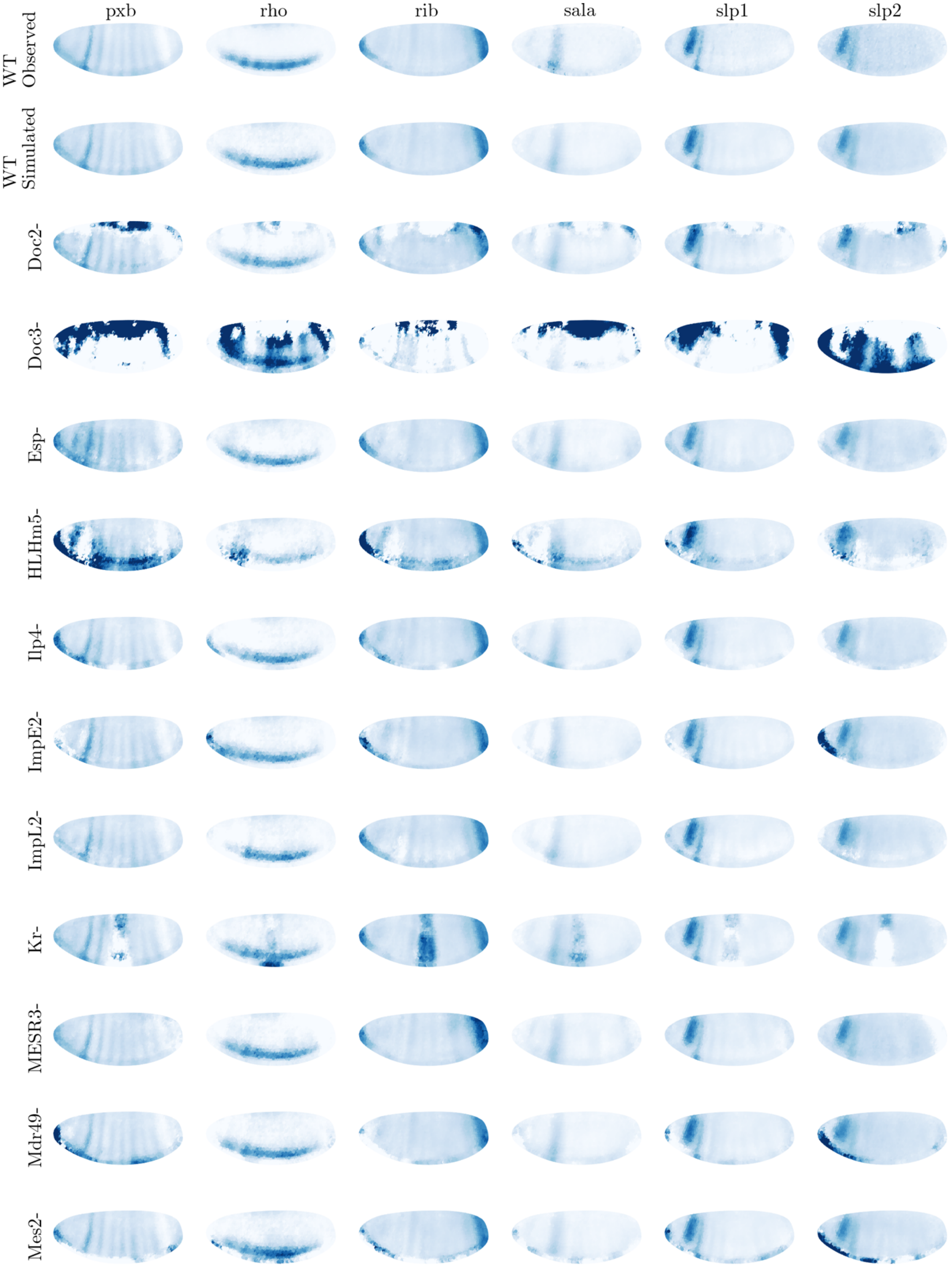
pxb
rho
rib
sala
slp1
slp2
WT Observed
WT Simulated
Doc2-
Doc3-
Esp-
HLHm5-
Ilp4-
ImpE2-
ImpL2-
Kr-
MESR3-
Mdr49-
Mes2-

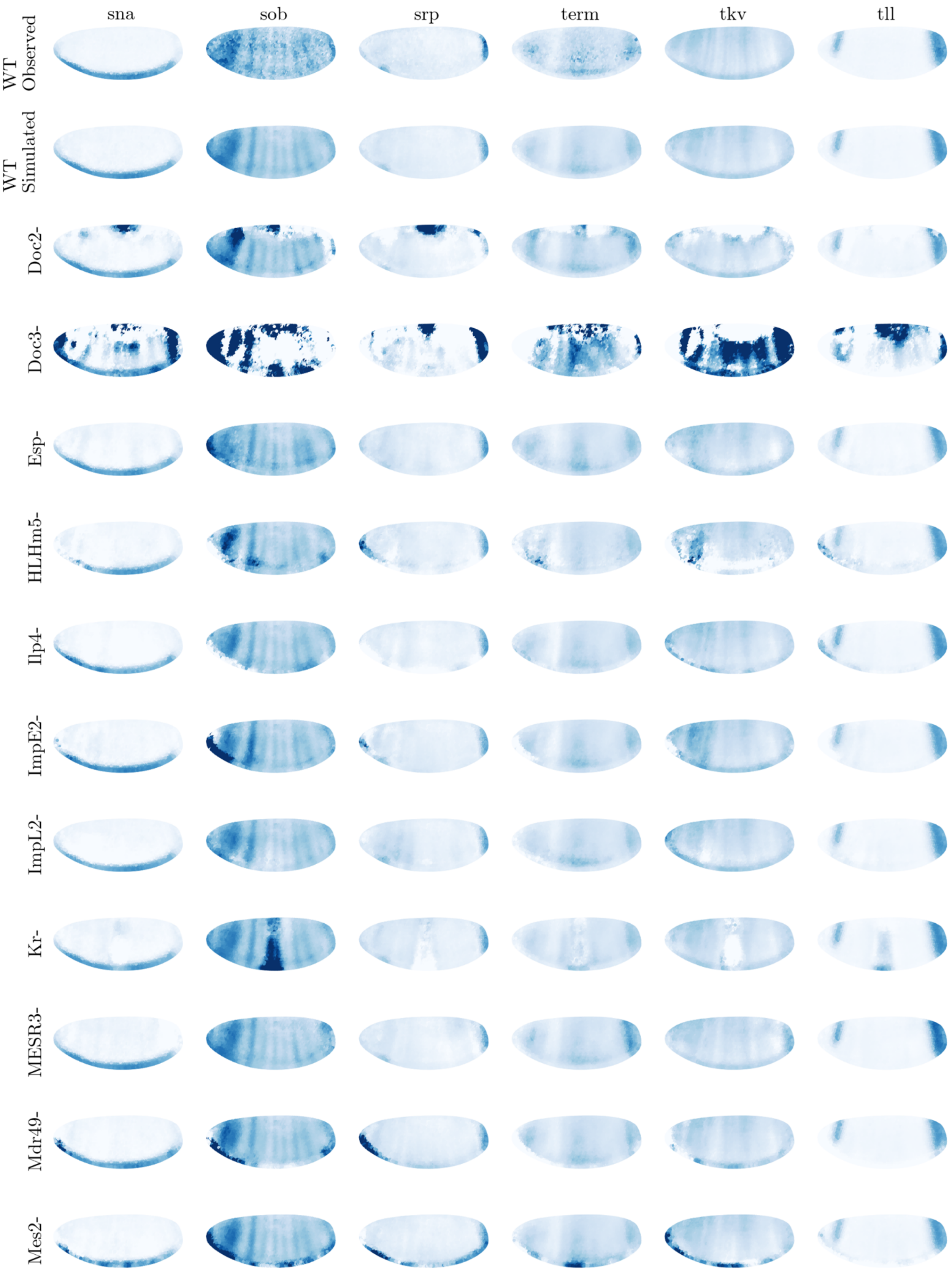
sna
sob
srp
term
tkv
tll
WT Observed
WT Simulated
Doc2-
Doc3-
Esp-
HLHm5-
Ilp4-
ImpE2-
ImpL2-
Kr-
MESR3-
Mdr49-
Mes2-

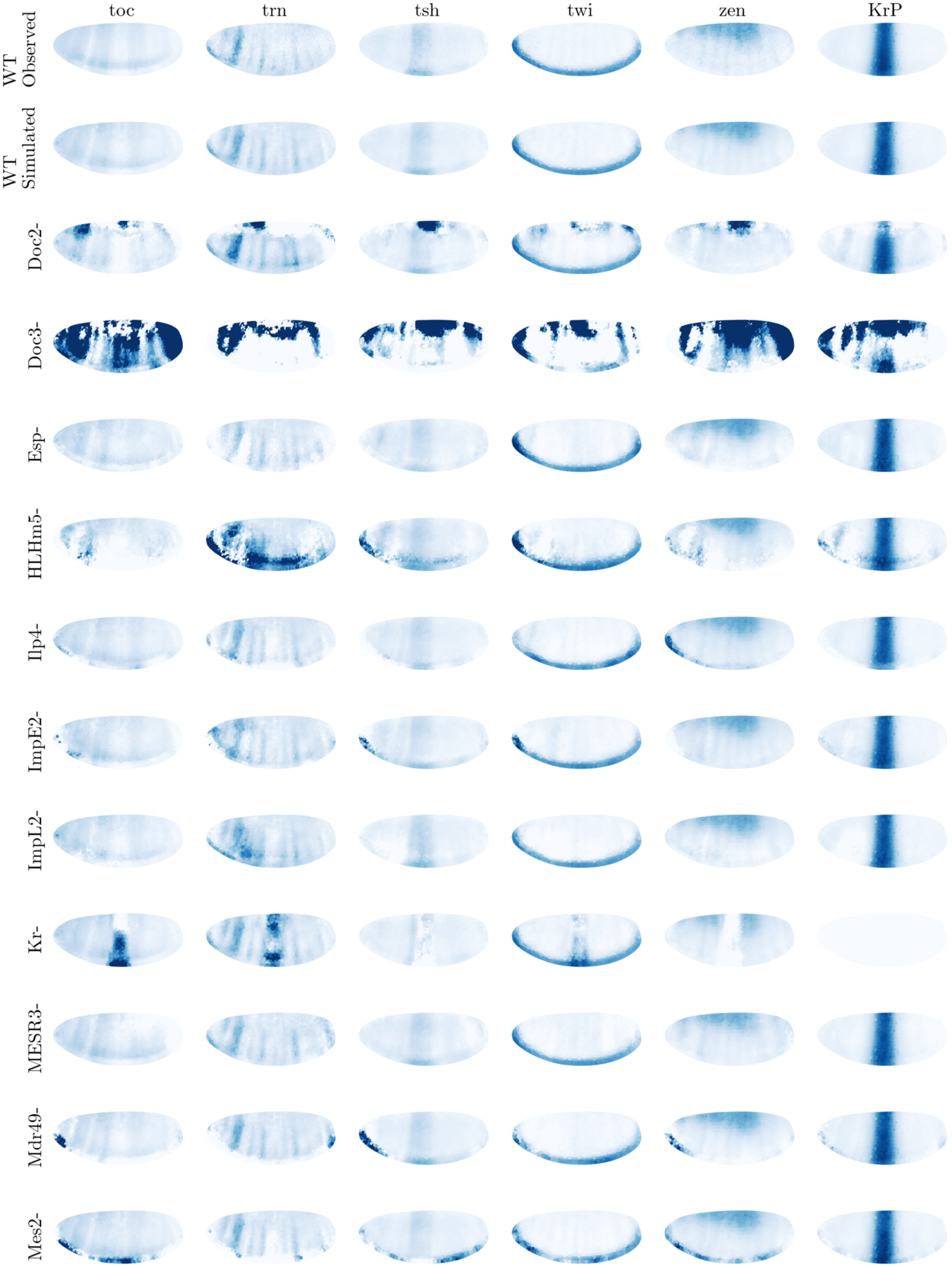

toc
trn
tsh
twi
zen
KrP
WT Observed
WT Simulated
Doc2-
Doc3-
Esp-
HLHm5-
Ilp4-
ImpE2-
ImpL2-
Kr-
MESR3-
Mdr49-
Mes2-

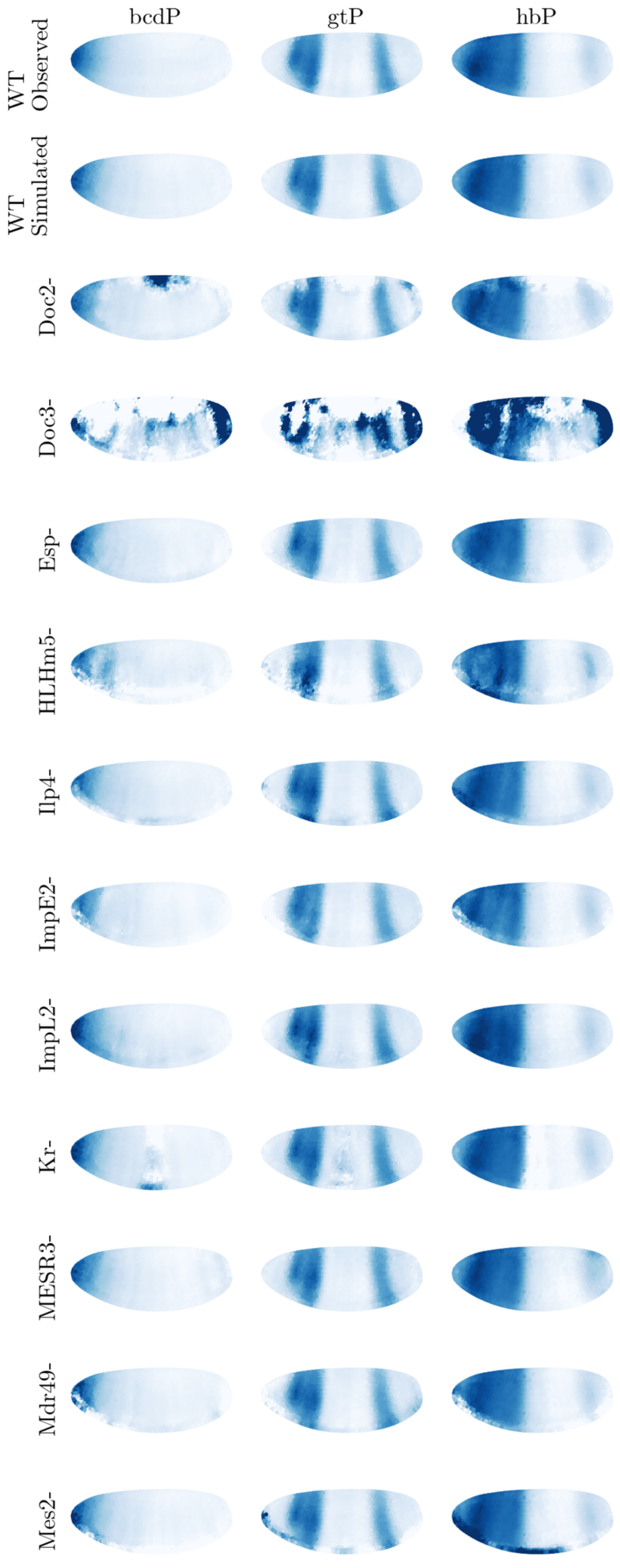
bcdP
gtP
hbP
WT Observed
WT Simulated
Doc2-
Doc3-
Esp-
HLHm5-
Ilp4-
ImpE2-
ImpL2-
Kr-
MESR3-
Mdr49-
Mes2-

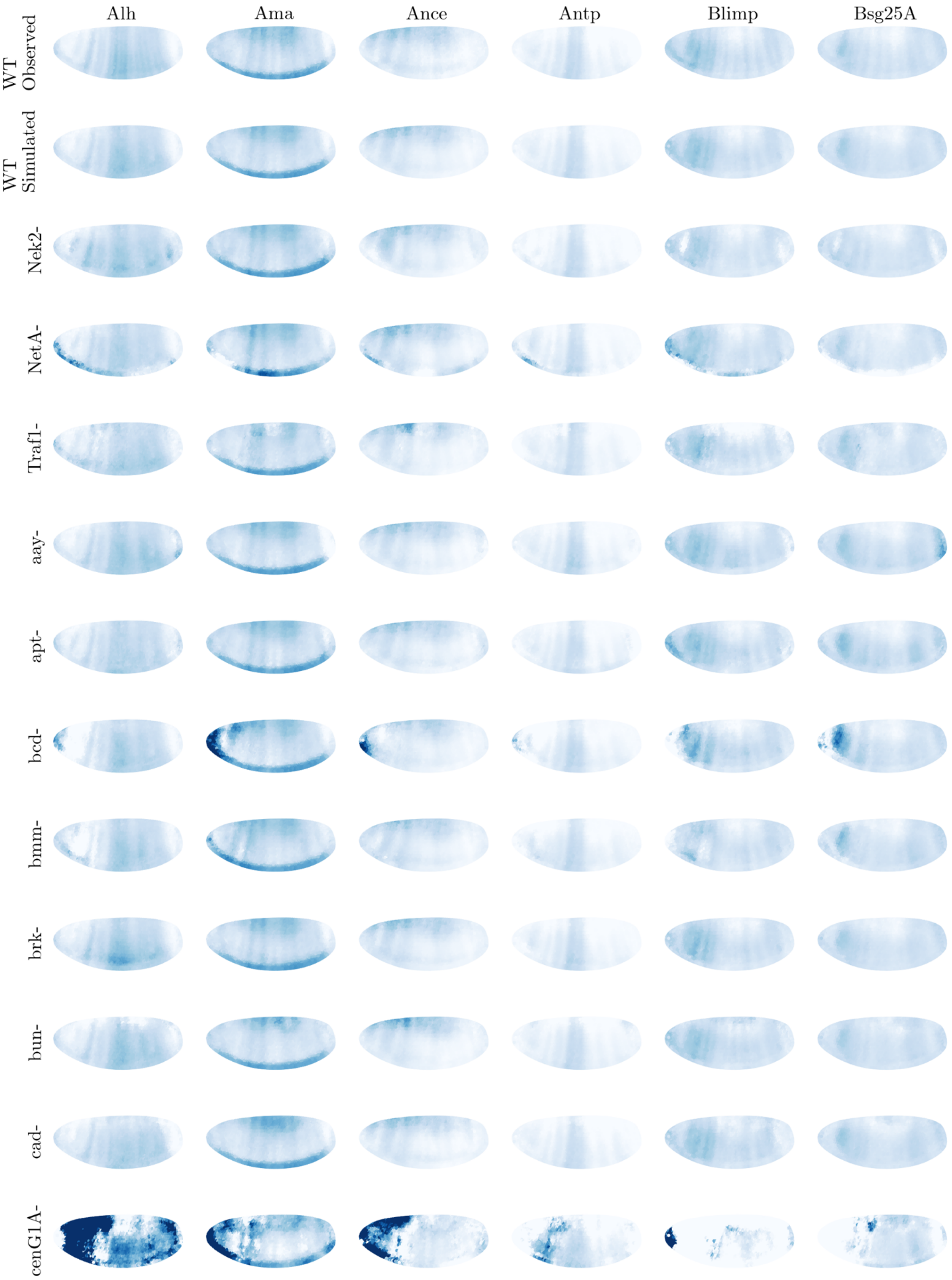
Alh
Ama
Ance
Antp
Blimp
Bsg25A
WT Observed
WT Simulated
Nek2-
NetA-
Traf1-
aay-
apt-
bcd-
bmm-
brk-
bun-
cad-
cenG1A-

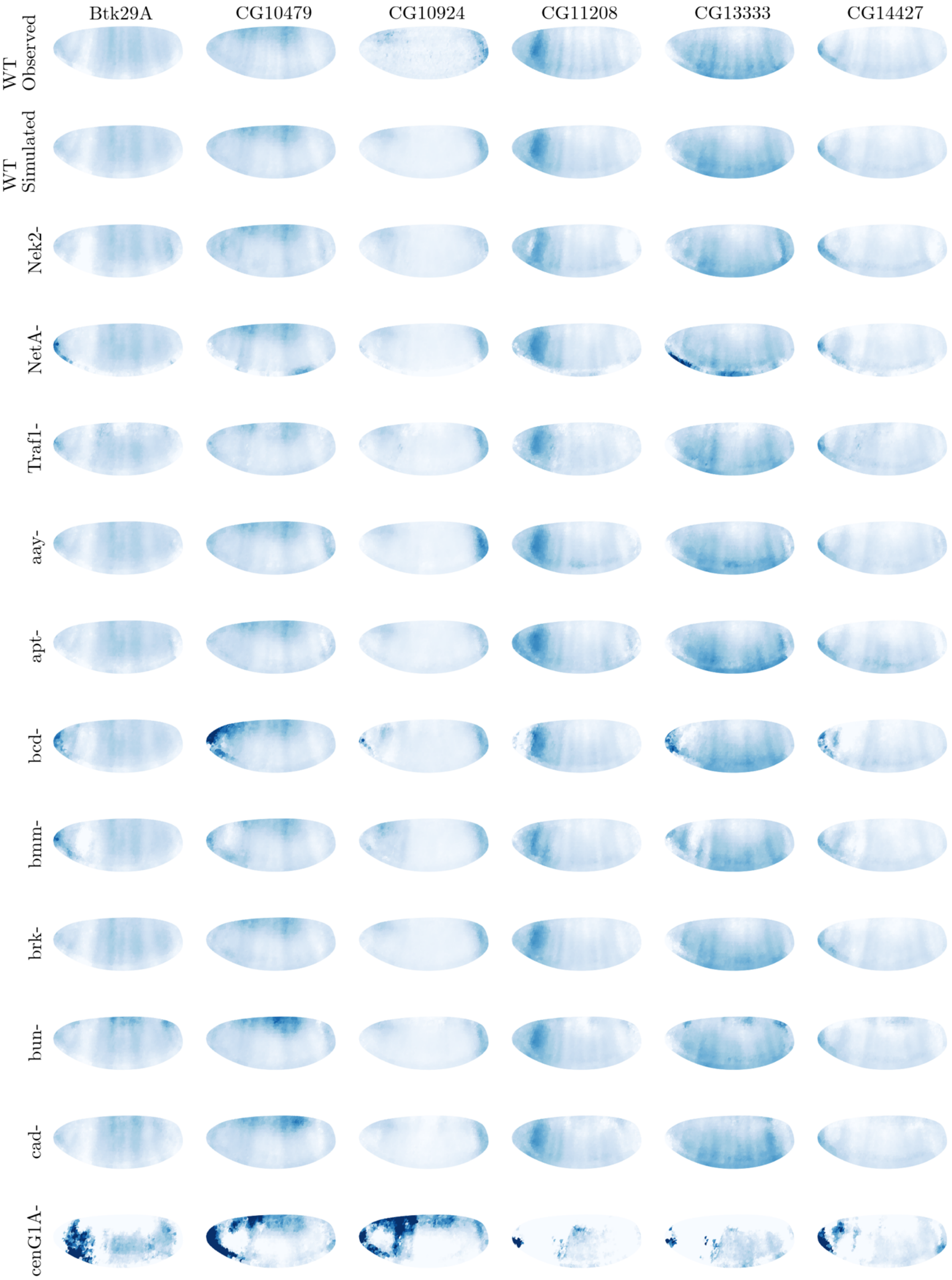
Btk29A
CG10479
CG10924
CG11208
CG13333
CG14427
WT Observed
WT Simulated
Nek2-
NetA-
Traf1-
aay-
apt-
bcd-
bmm-
brk-
bun-
cad-
cenG1A-

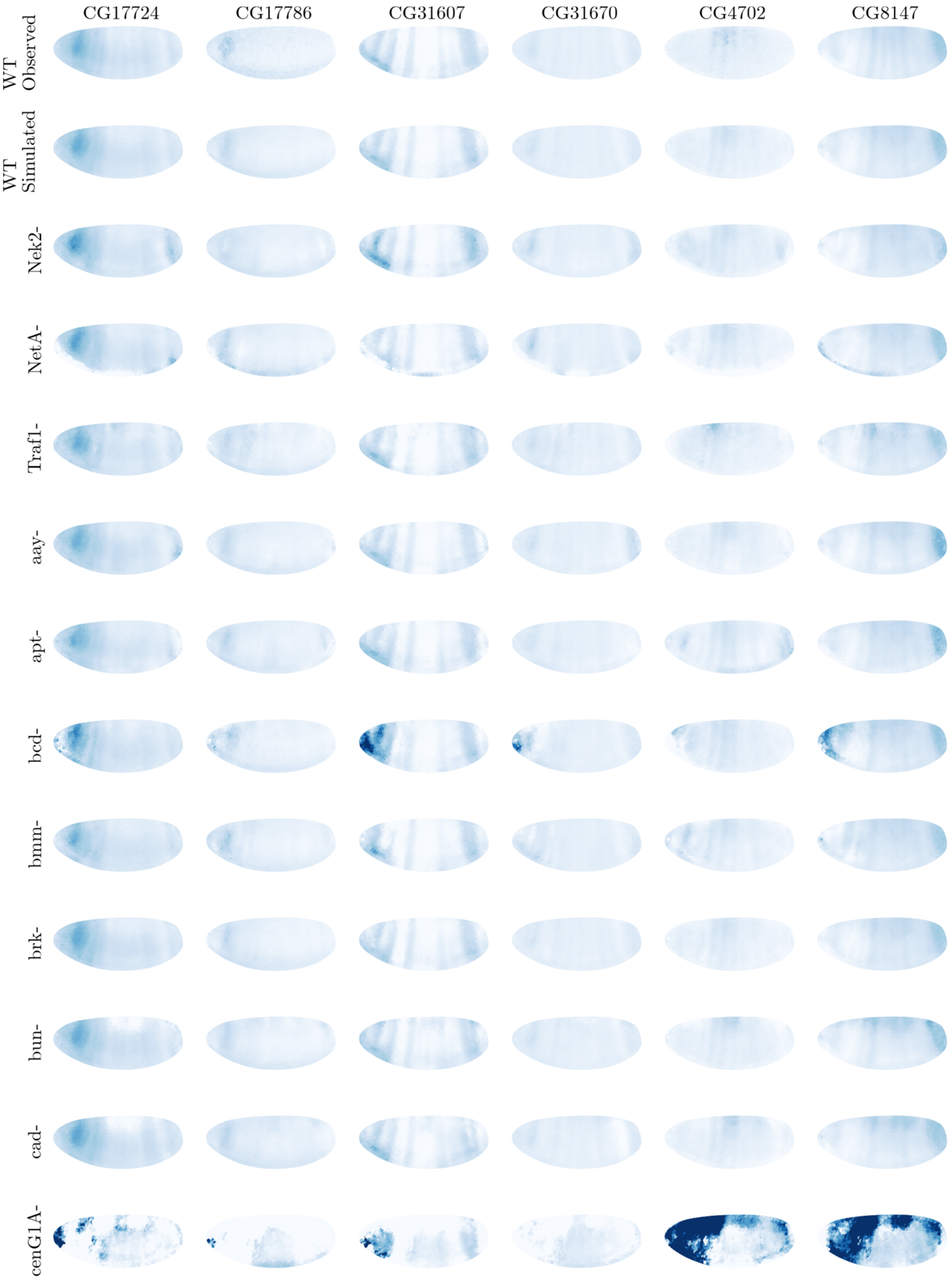
CG17724
CG17786
CG31607
CG31670
CG4702
CG8147
WT Observed
WT Simulated
Nek2-
NetA-
Traf1-
aay-
apt-
bcd-
bmm-
brk-
bun-
cad-
cenG1A-

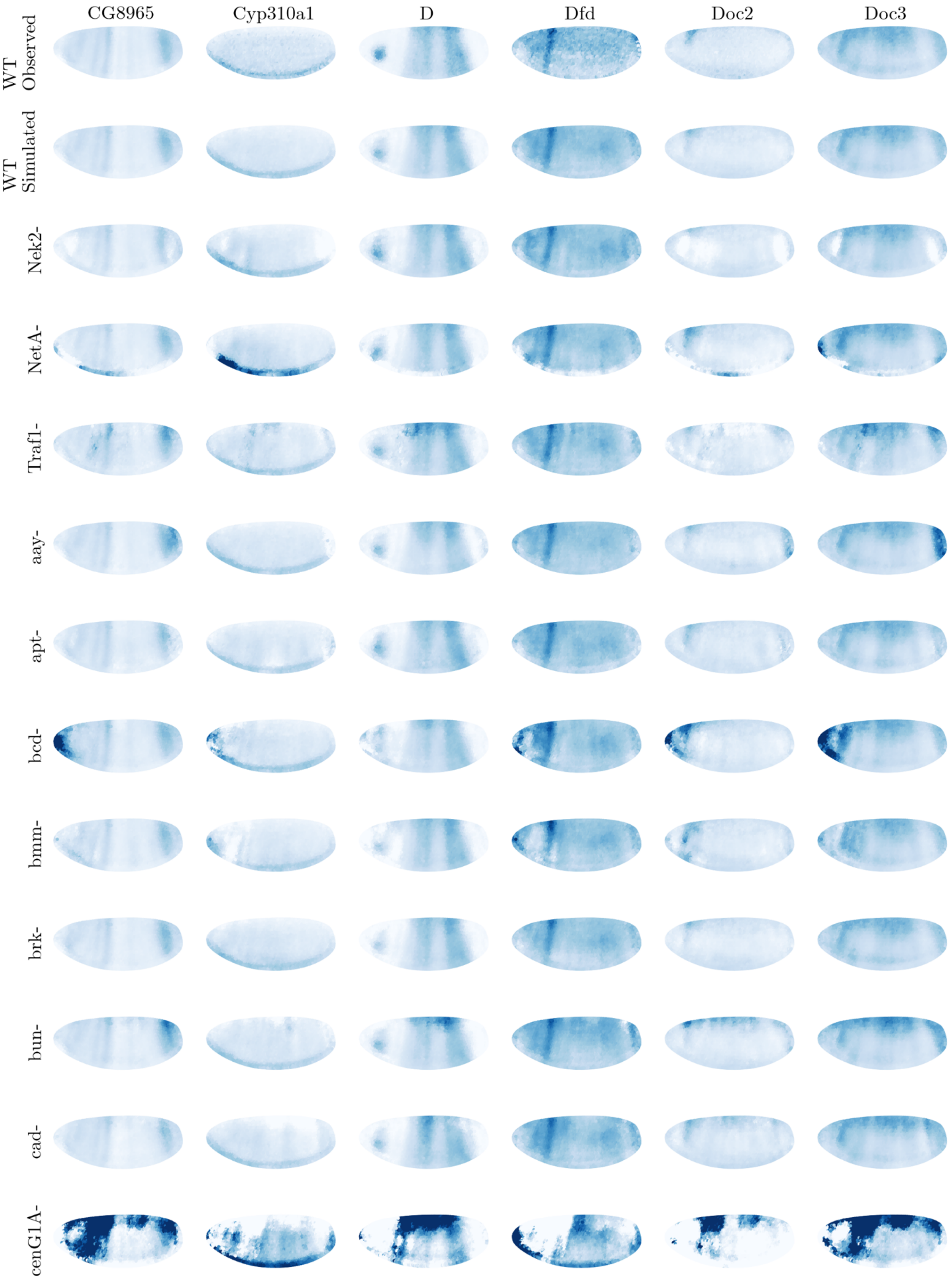
CG8965
Cyp310a1
D
Dfd
Doc2
Doc3
WT Observed
WT Simulated
Nek2-
NetA-
Traf1-
aay-
apt-
bcd-
bmm-
brk-
bun-
cad-
cenG1A-

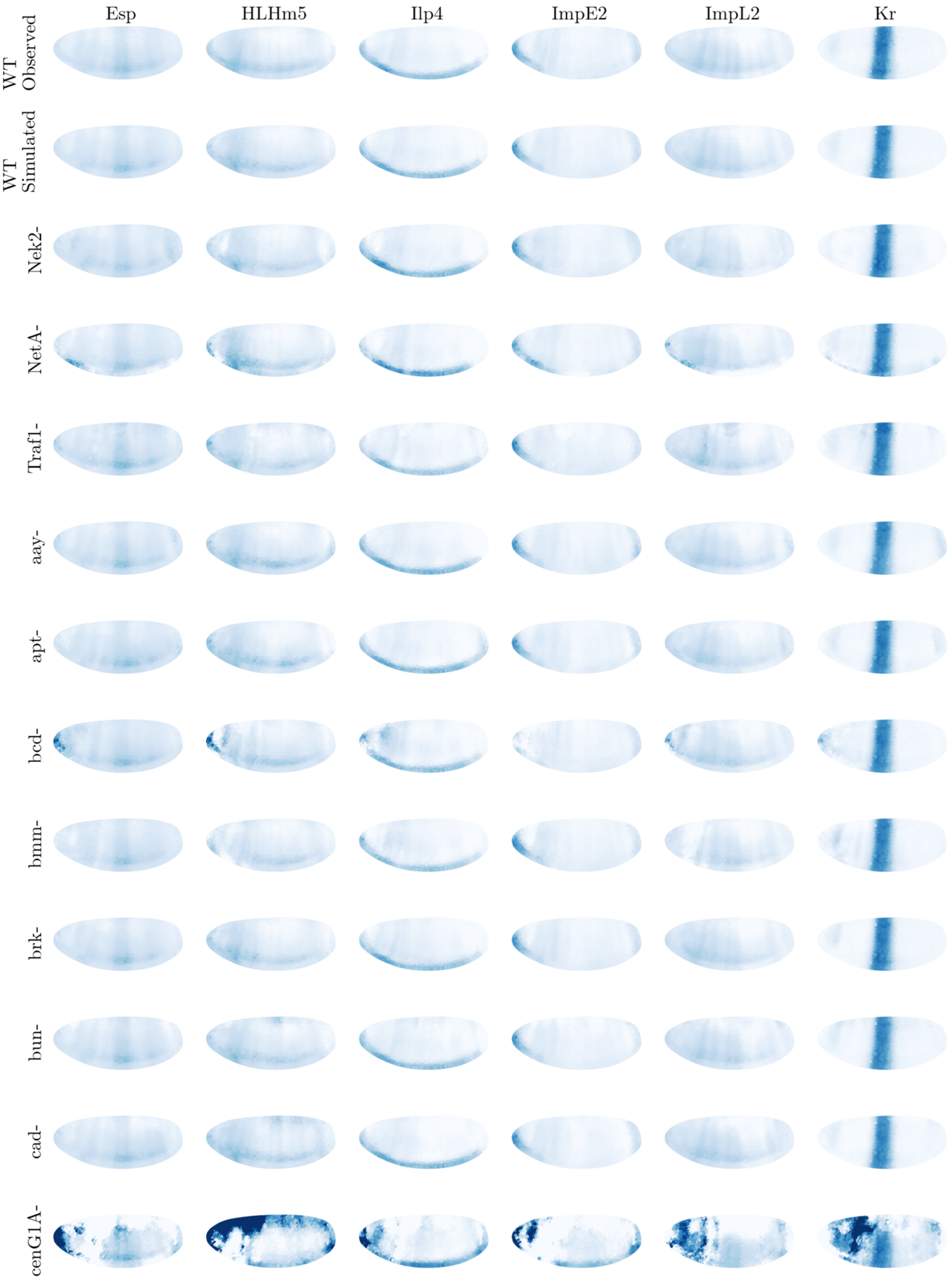
Esp
HLHm5
Ilp4
ImpE2
ImpL2
Kr
WT Observed
WT Simulated
Nek2-
NetA-
Traf1-
aay-
apt-
bcd-
bmm-
brk-
bun-
cad-
cenG1A-

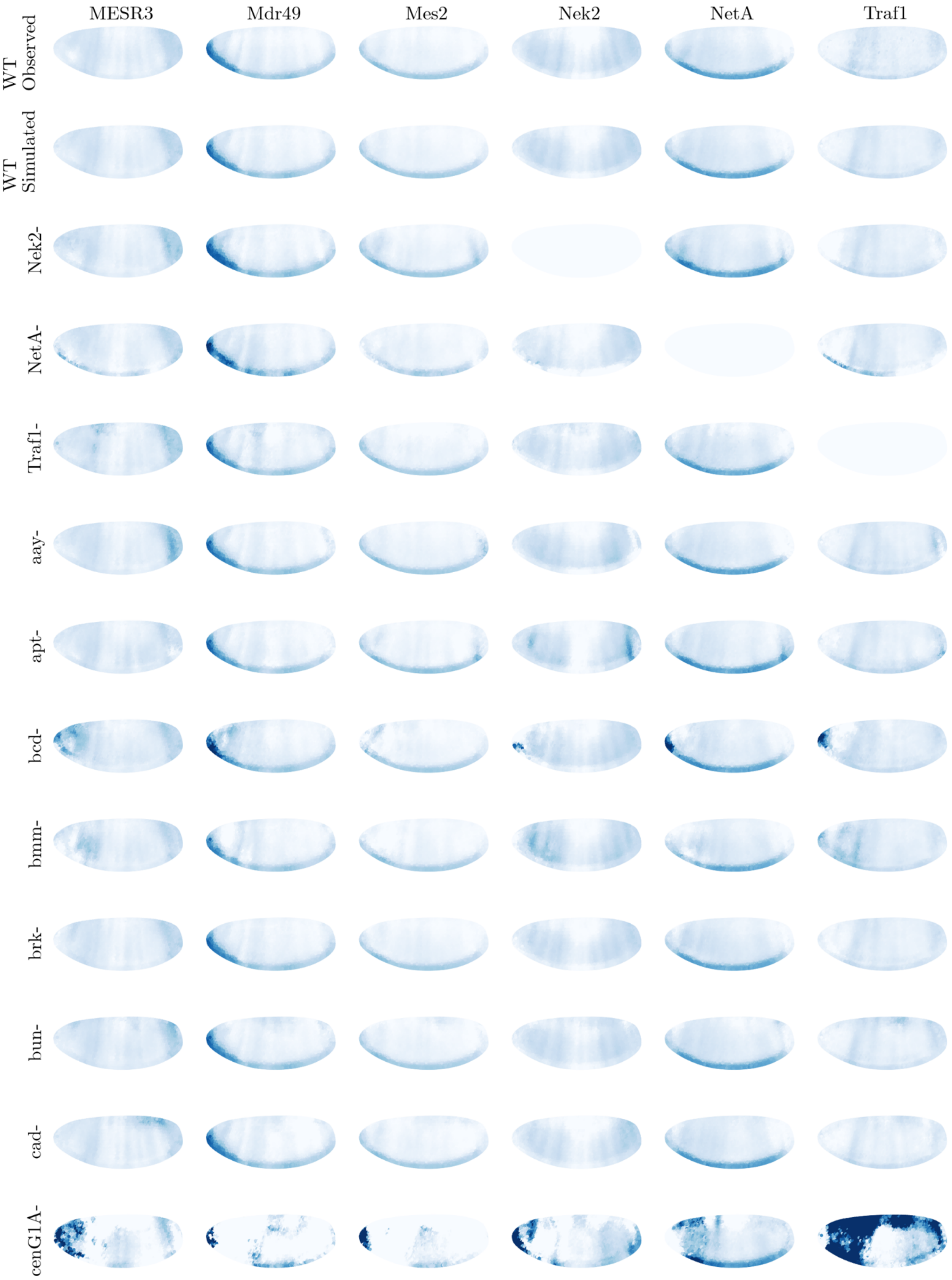
MESR3
Mdr49
Mes2
Nek2
NetA
Traf1
WT Observed
WT Simulated
Nek2-
NetA-
Traf1-
aay-
apt-
bcd-
bmm-
brk-
bun-
cad-
cenG1A-

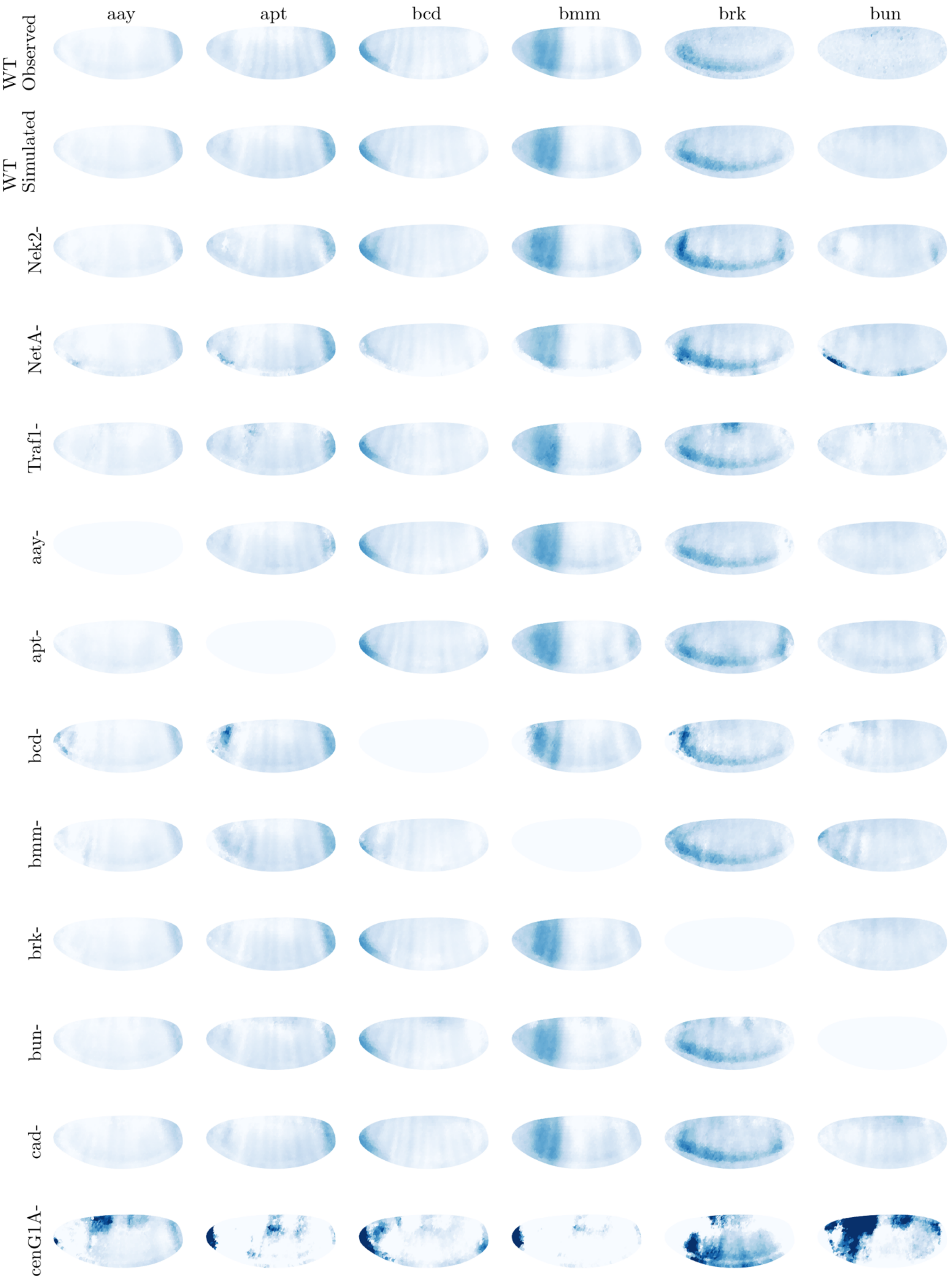
aay
apt
bcd
bmm
brk
bun
WT Observed
WT Simulated
Nek2-
NetA-
Traf1-
aay-
apt-
bcd-
bmm-
brk-
bun-
cad-
cenG1A-

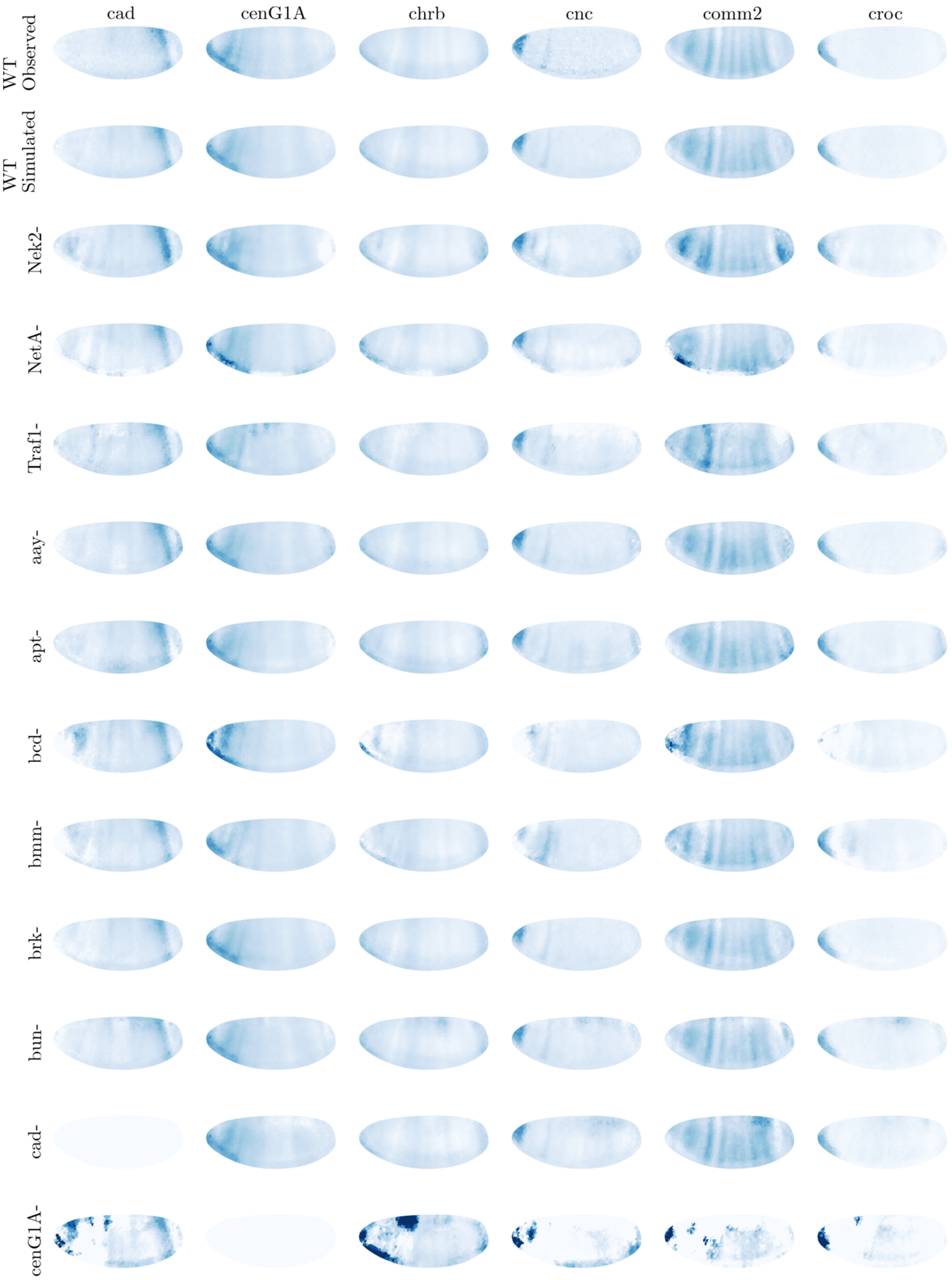
cad
cenG1A
chrb
cnc
comm2
croc
WT Observed
WT Simulated
Nek2-
NetA-
Traf1-
aay-
apt-
bcd-
bmm-
brk-
bun-
cad-
cenG1A-

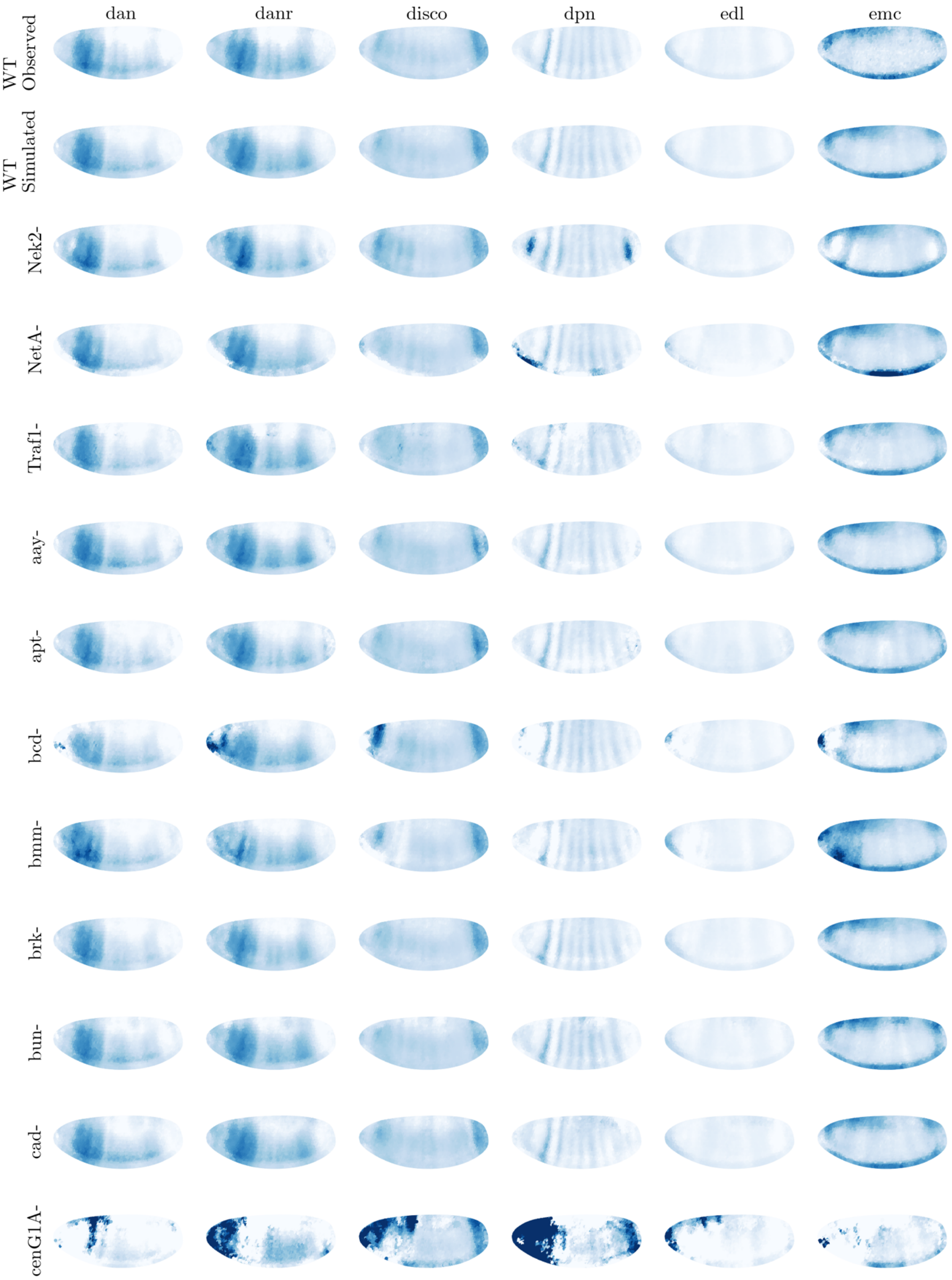
dan
danr
disco
dpn
edl
emc
WT Observed
WT Simulated
Nek2-
NetA-
Traf1-
aay-
apt-
bcd-
bmm-
brk-
bun-
cad-
cenG1A-

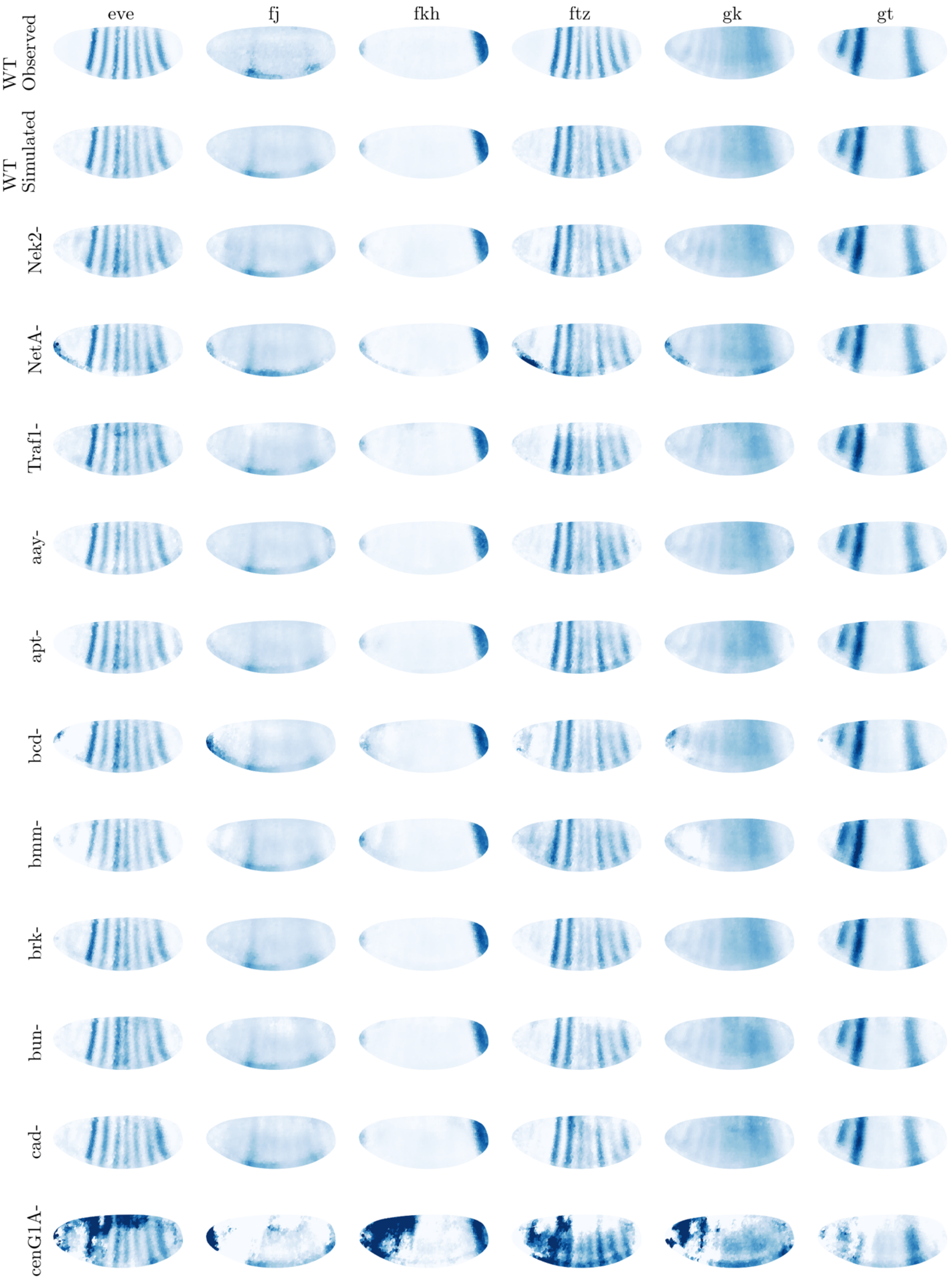
eve
fj
fkh
ftz
gk
gt
WT Observed
WT Simulated
Nek2-
NetA-
Traf1-
aay-
apt-
bcd-
bmm-
brk-
bun-
cad-
cenG1A-

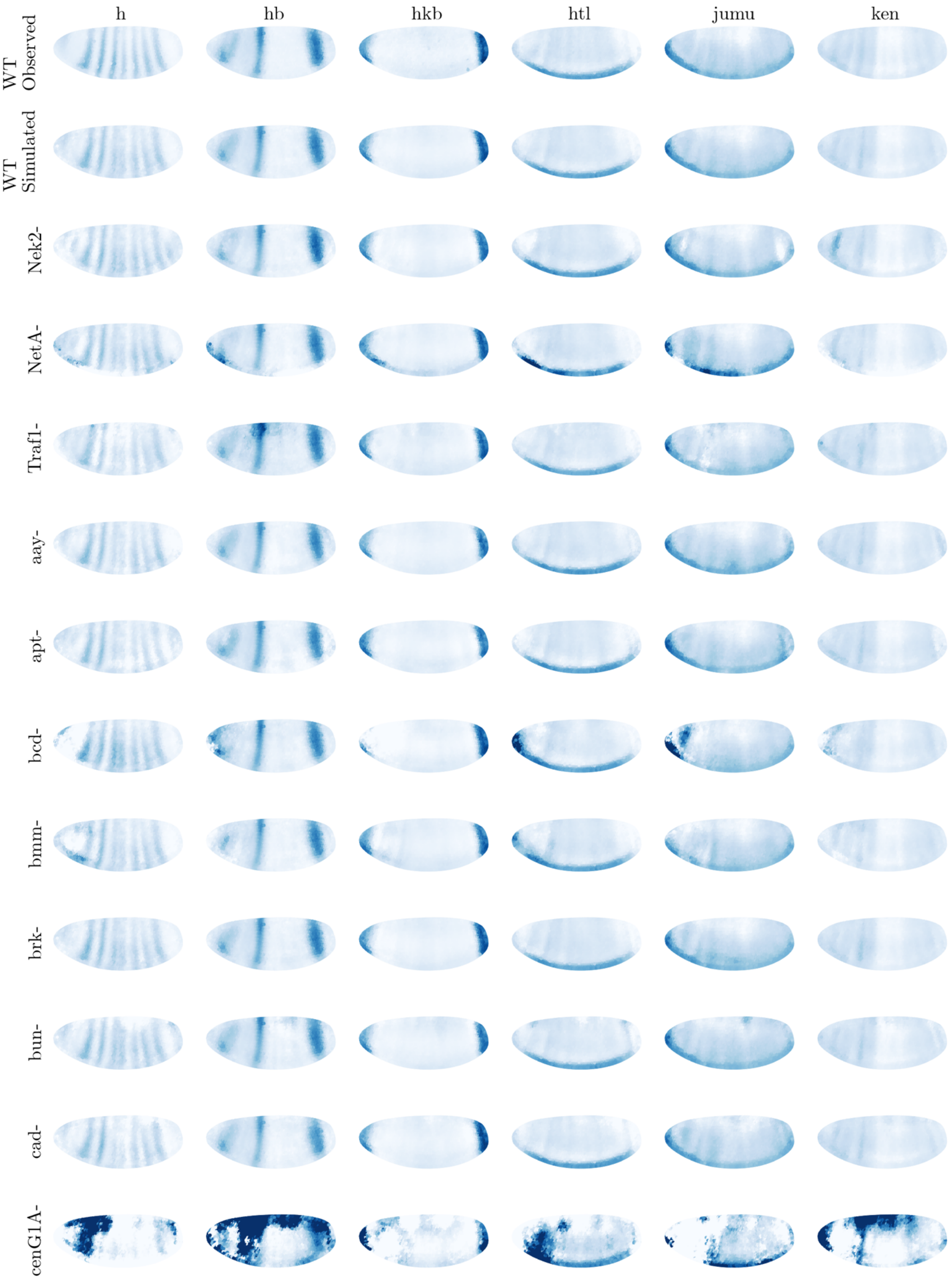
h
hb
hkb
htl
jumu
ken
WT Observed
WT Simulated
Nek2-
NetA-
Traf1-
aay-
apt-
bcd-
bmm-
brk-
bun-
cad-
cenG1A-

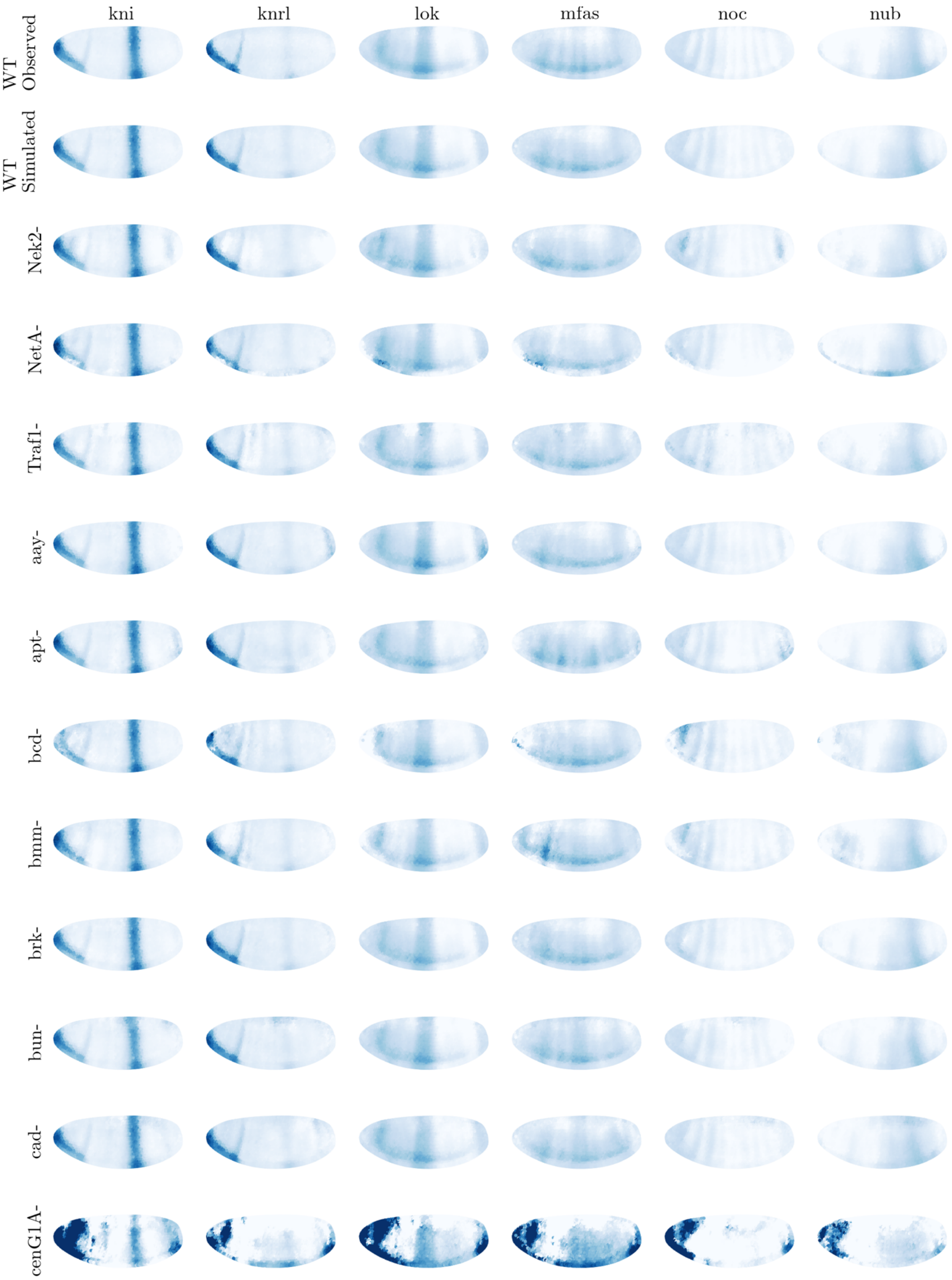
kni
knrl
lok
mfas
noc
nub
WT Observed
WT Simulated
Nek2-
NetA-
Traf1-
aay-
apt-
bcd-
bmm-
brk-
bun-
cad-
cenG1A-

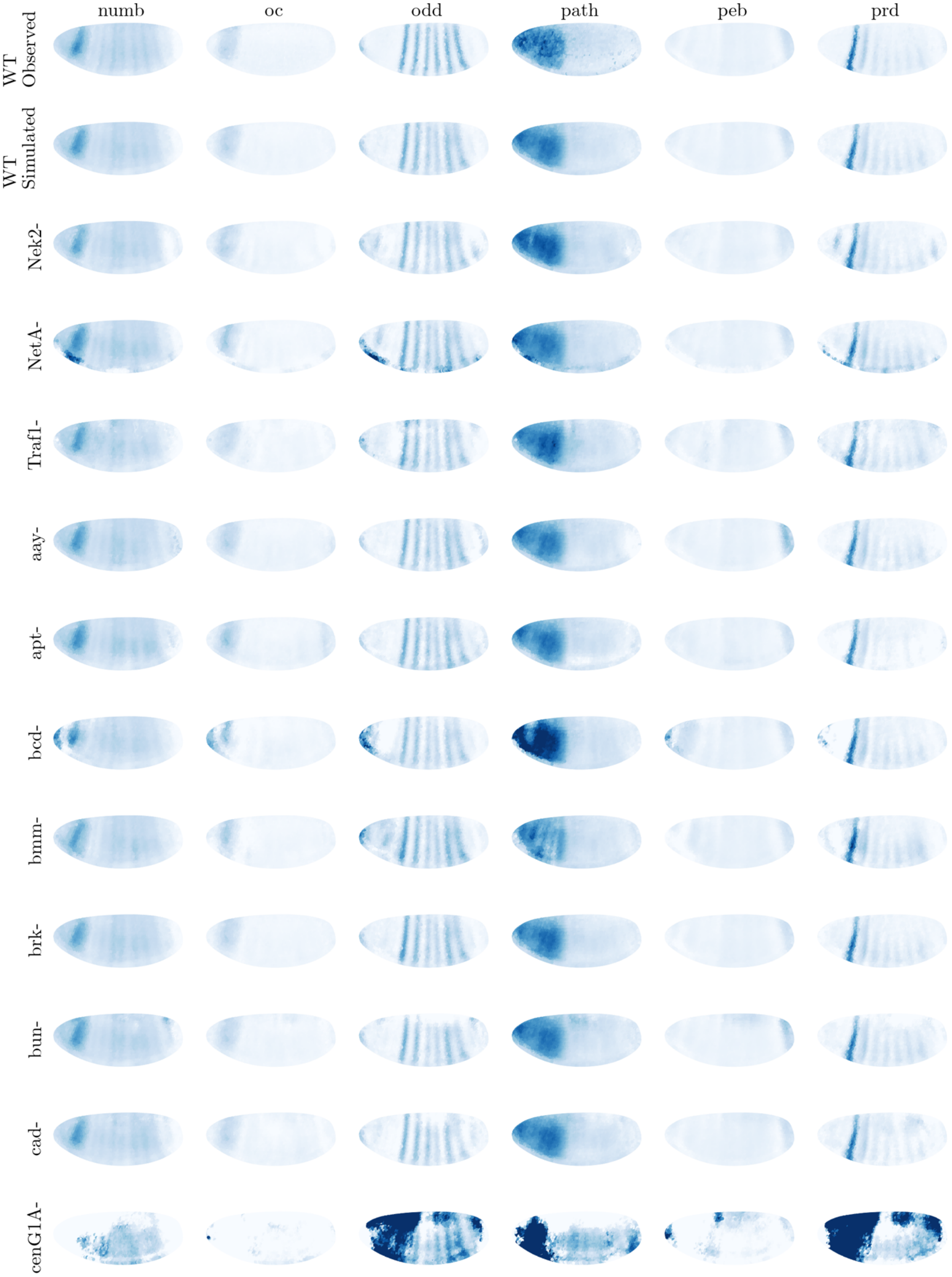
numb
oc
odd
path
peb
prd
WT Observed
WT Simulated
Nek2-
NetA-
Traf1-
aay-
apt-
bcd-
bmm-
brk-
bun-
cad-
cenG1A-

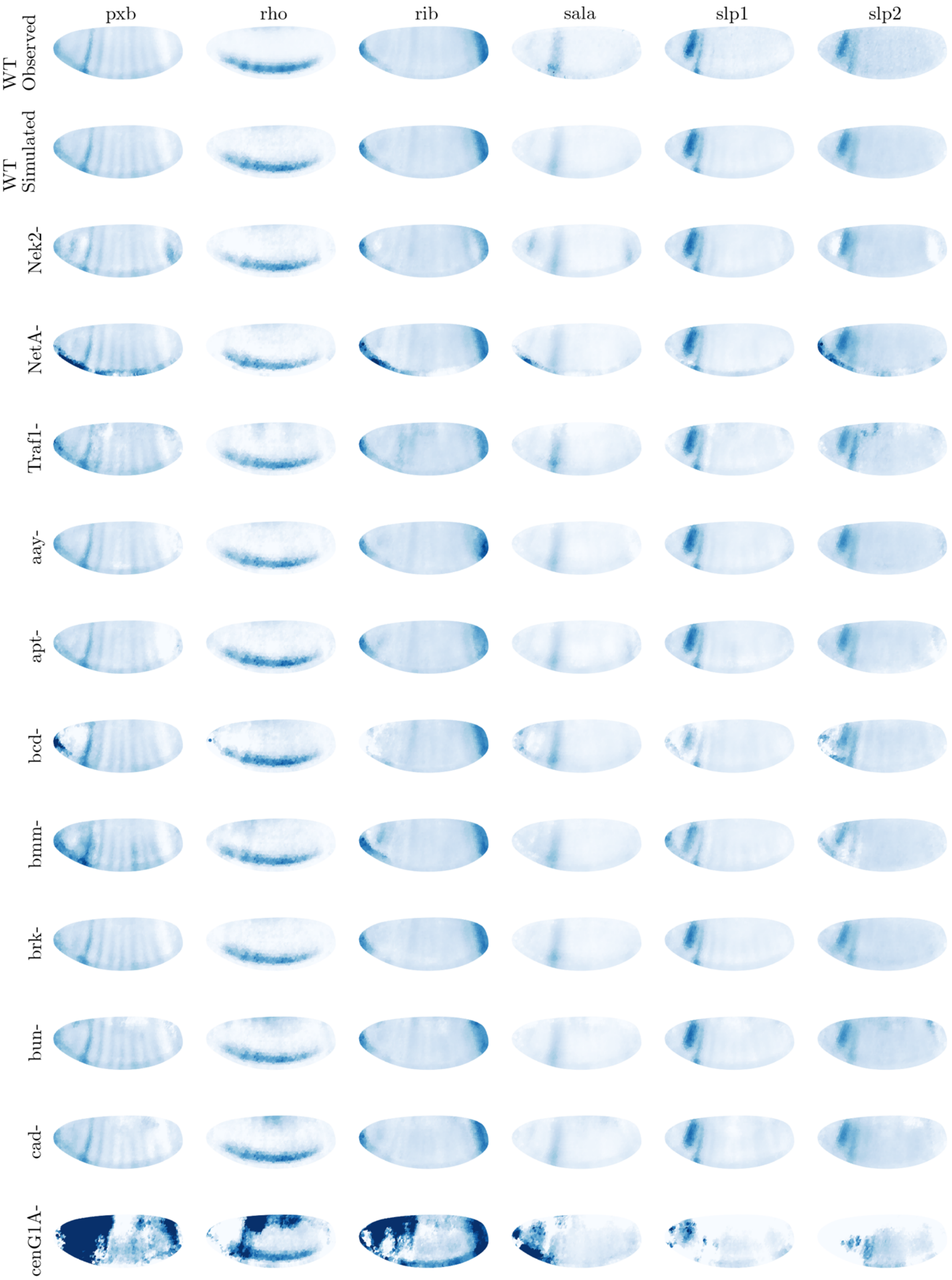

pxb
rho
rib
sala
slp1
slp2
WT Observed
WT Simulated
Nek2-
NetA-
Traf1-
aay-
apt-
bcd-
bmm-
brk-
bun-
cad-
cenG1A-

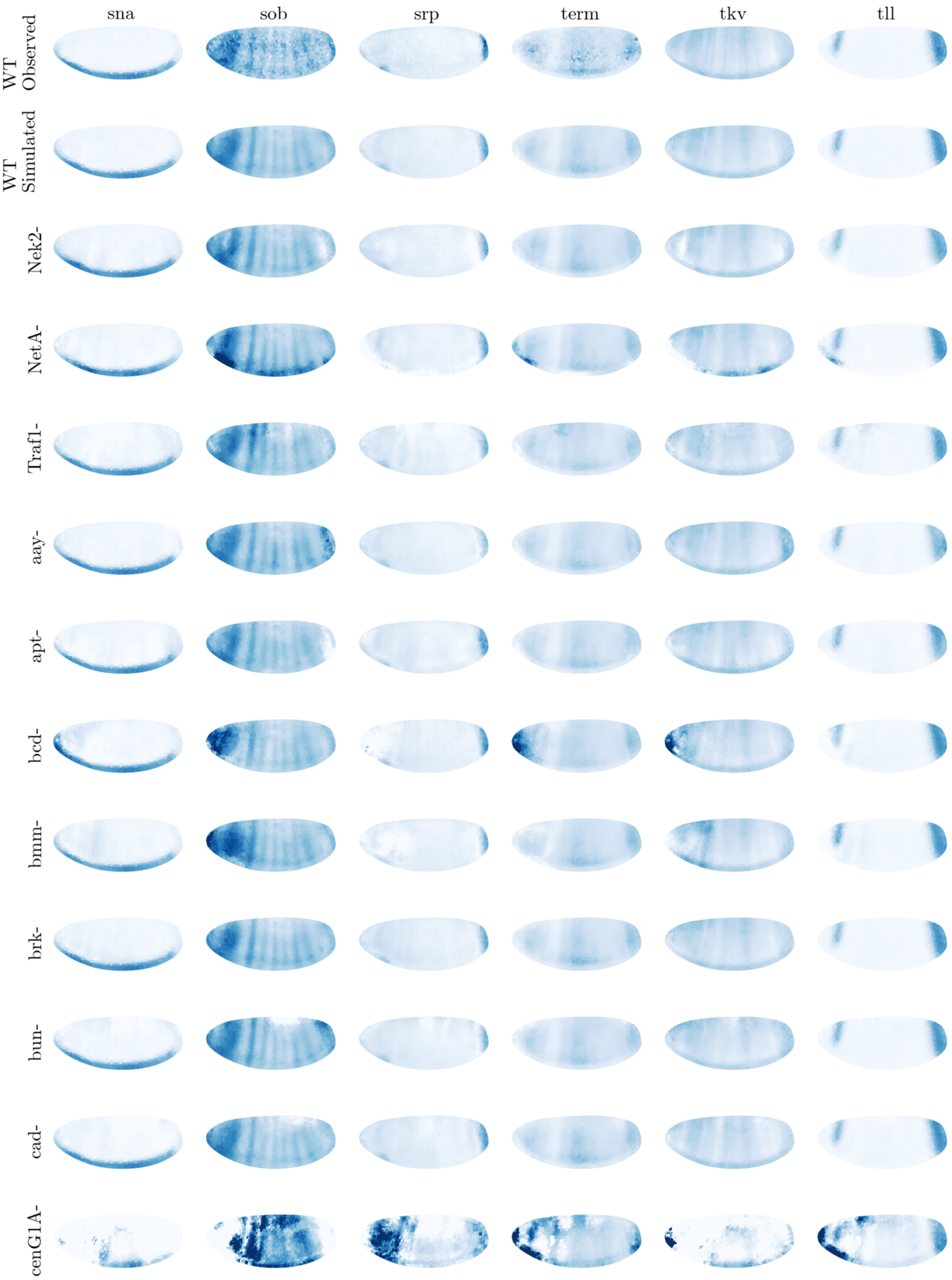
sna
sob
srp
term
tkv
tll
WT Observed
WT Simulated
Nek2-
NetA-
Traf1-
aay-
apt-
bcd-
bmm-
brk-
bun-
cad-
cenG1A-

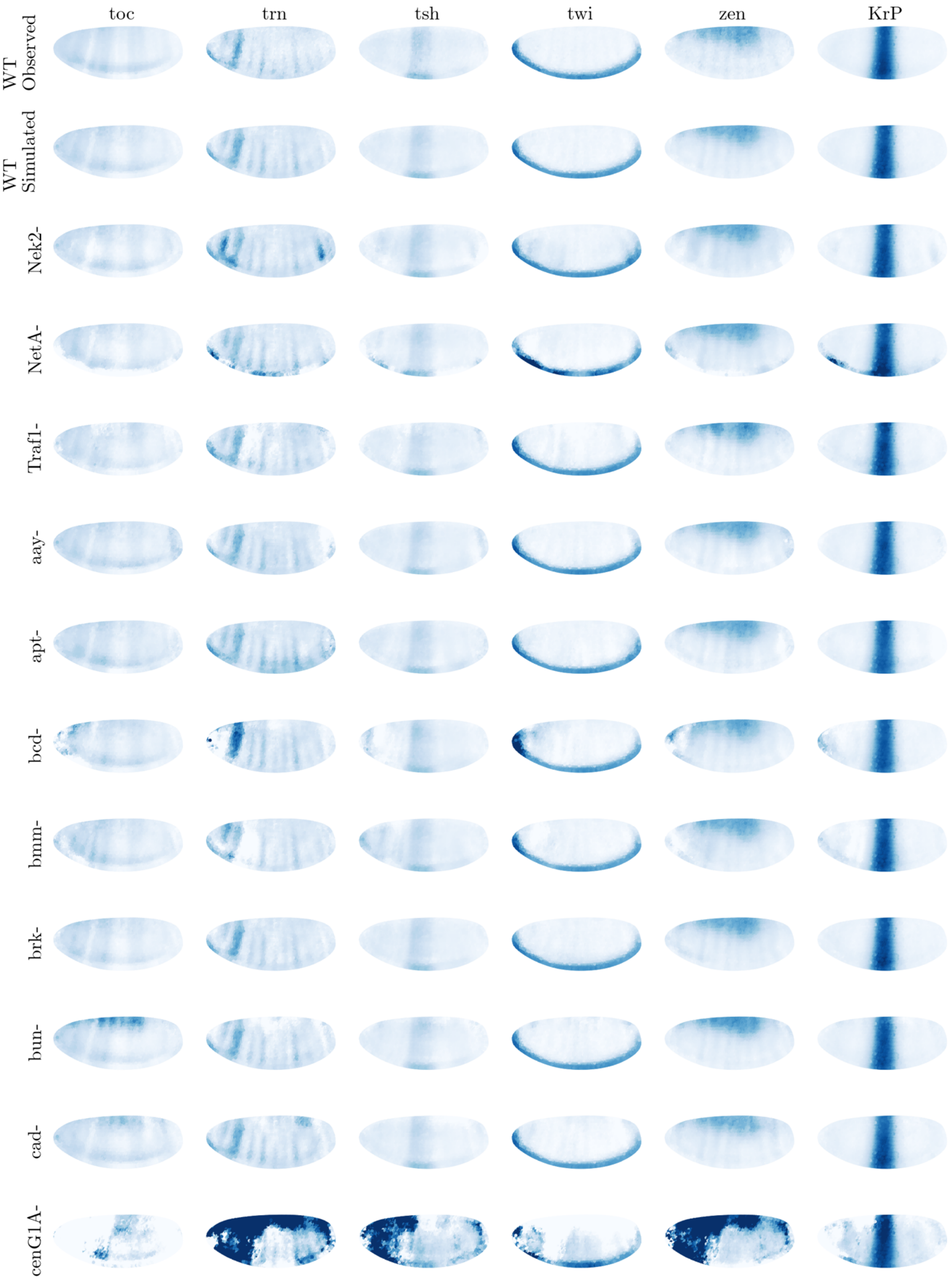

toc
trn
tsh
twi
zen
KrP
WT Observed
WT Simulated
Nek2-
NetA-
Traf1-
aay-
apt-
bcd-
bmm-
brk-
bun-
cad-
cenG1A-

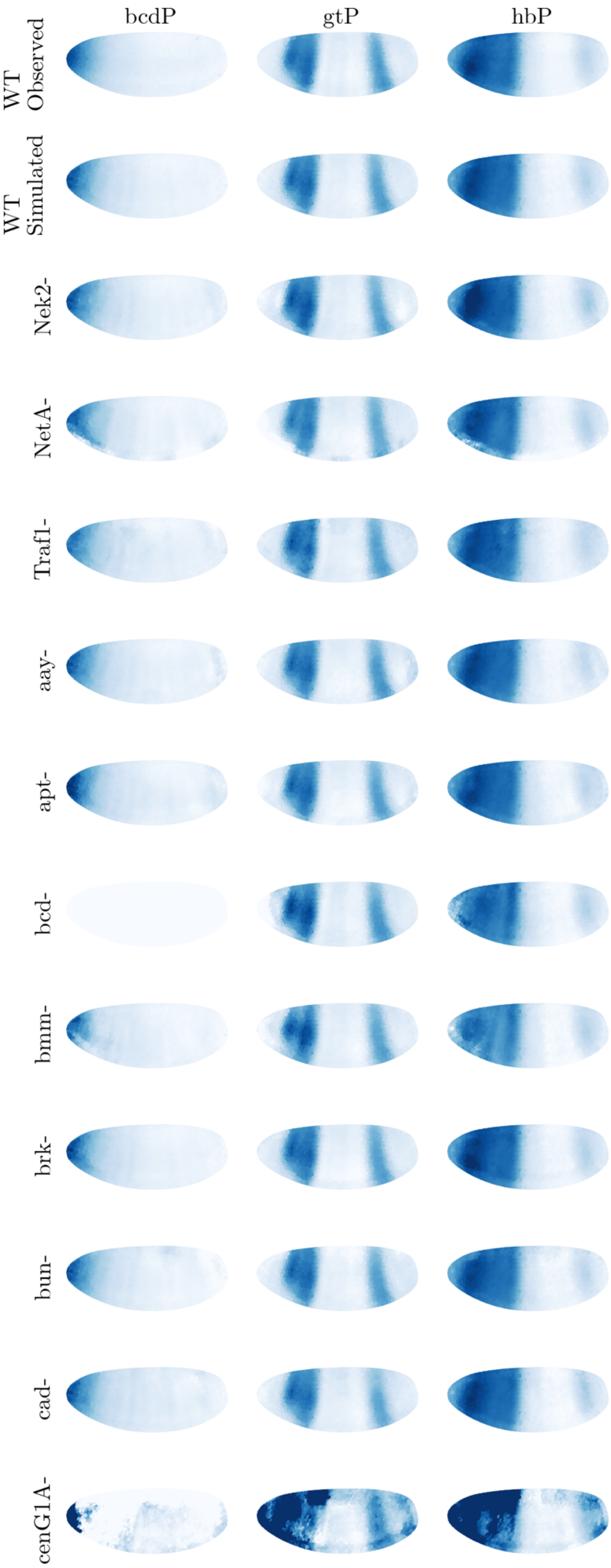
bcdP
gtP
hbP
WT Observed
WT Simulated
Nek2-
NetA-
Traf1-
aay-
apt-
bcd-
bmm-
brk-
bun-
cad-
cenG1A-

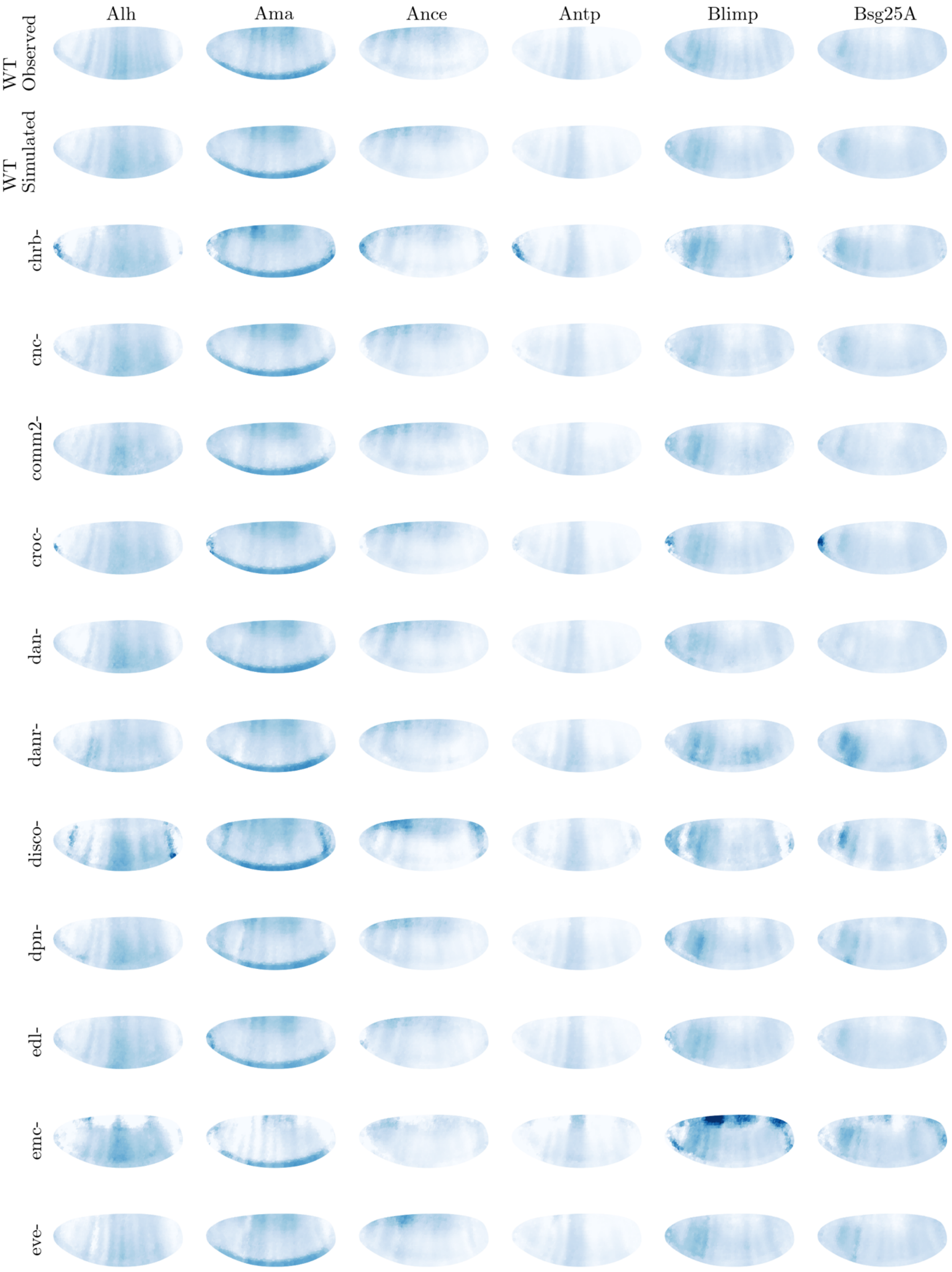

Alh
Ama
Ance
Antp
Blimp
Bsg25A
WT Observed
WT Simulated
chrb-
cnc-
comm2-
croc-
dan-
danr-
disco-
dpn-
edl-
emc-
eve-

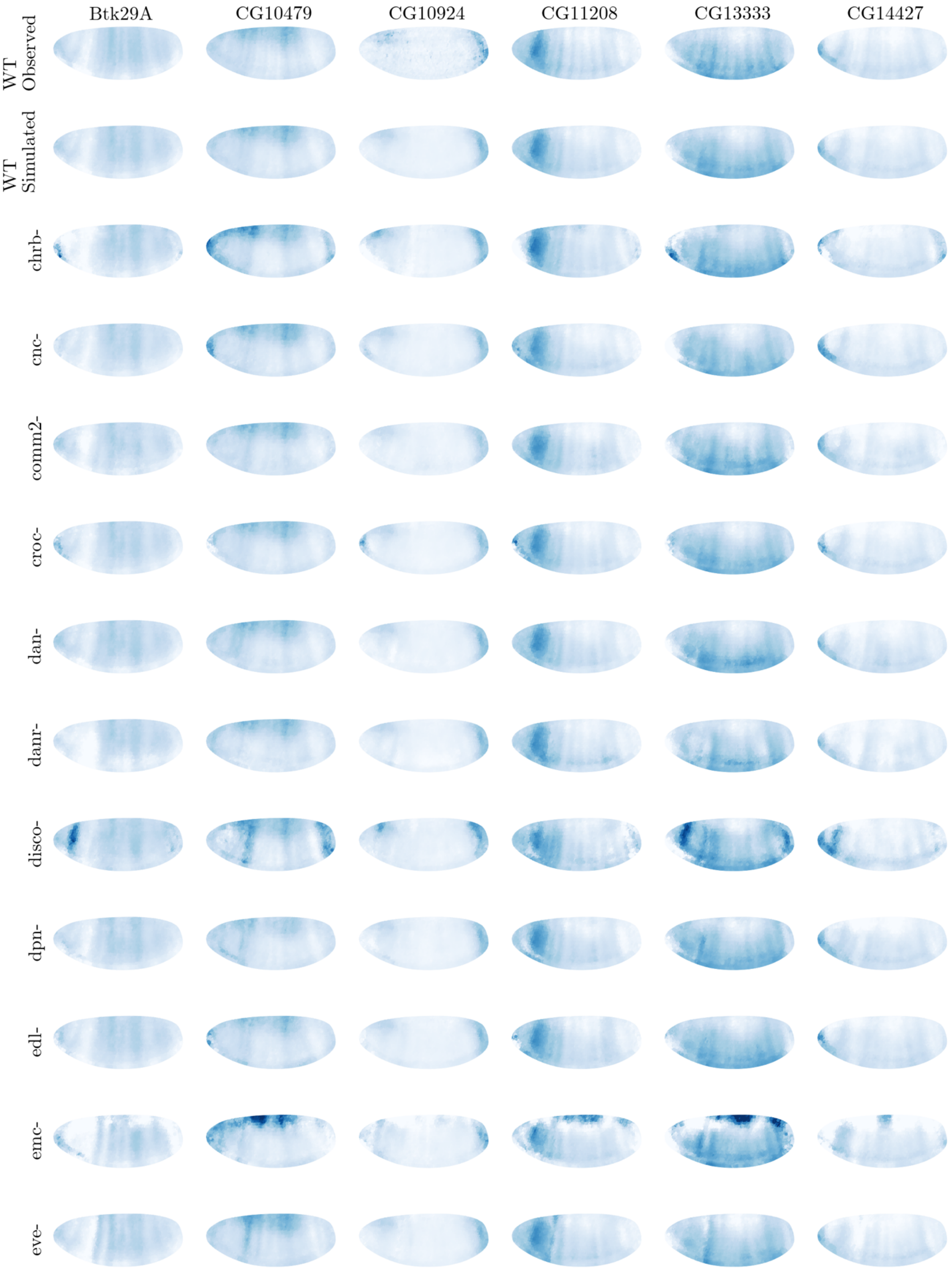

Btk29A
CG10479
CG10924
CG11208
CG13333
CG14427
WT Observed
WT Simulated
chrb-
cnc-
comm2-
croc-
dan-
danr-
disco-
dpn-
edl-
emc-
eve-

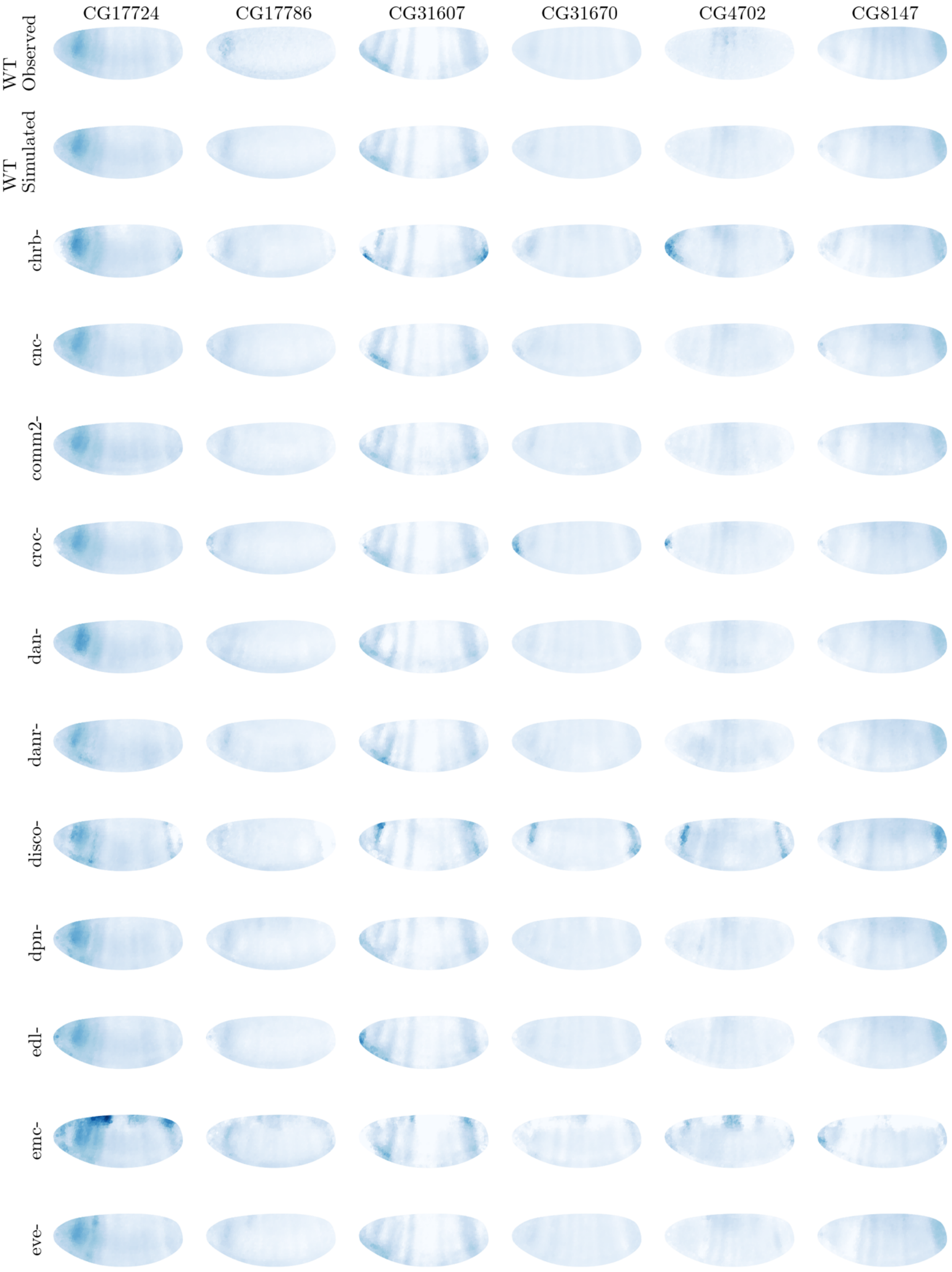
CG17724
CG17786
CG31607
CG31670
CG4702
CG8147
WT Observed
WT Simulated
chrb-
cnc-
comm2-
croc-
dan-
danr-
disco-
dpn-
edl-
emc-
eve-

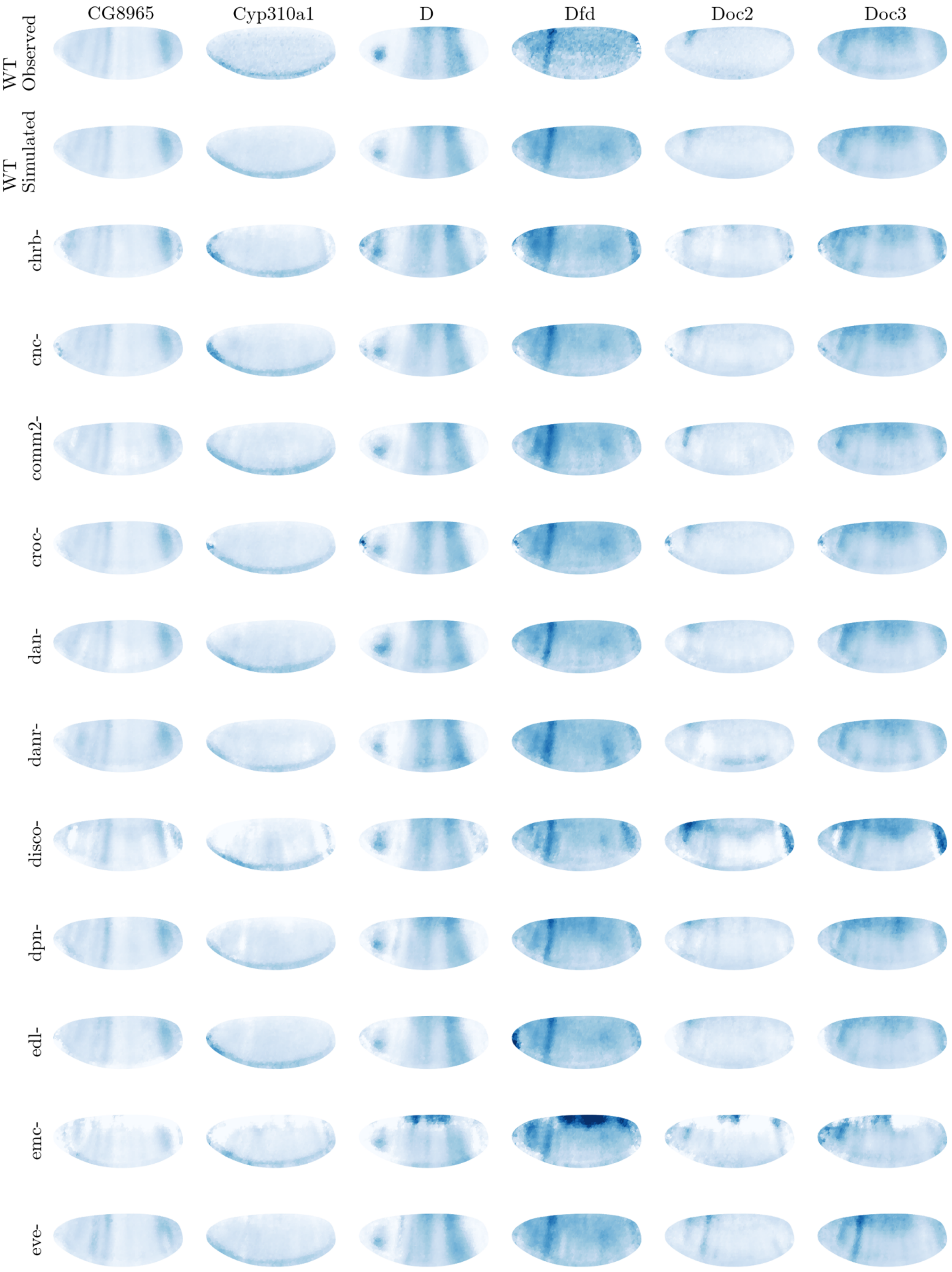

CG8965
Cyp310a1
D
Dfd
Doc2
Doc3
WT Observed
WT Simulated
chrb-
cnc-
comm2-
croc-
dan-
danr-
disco-
dpn-
edl-
emc-
eve-

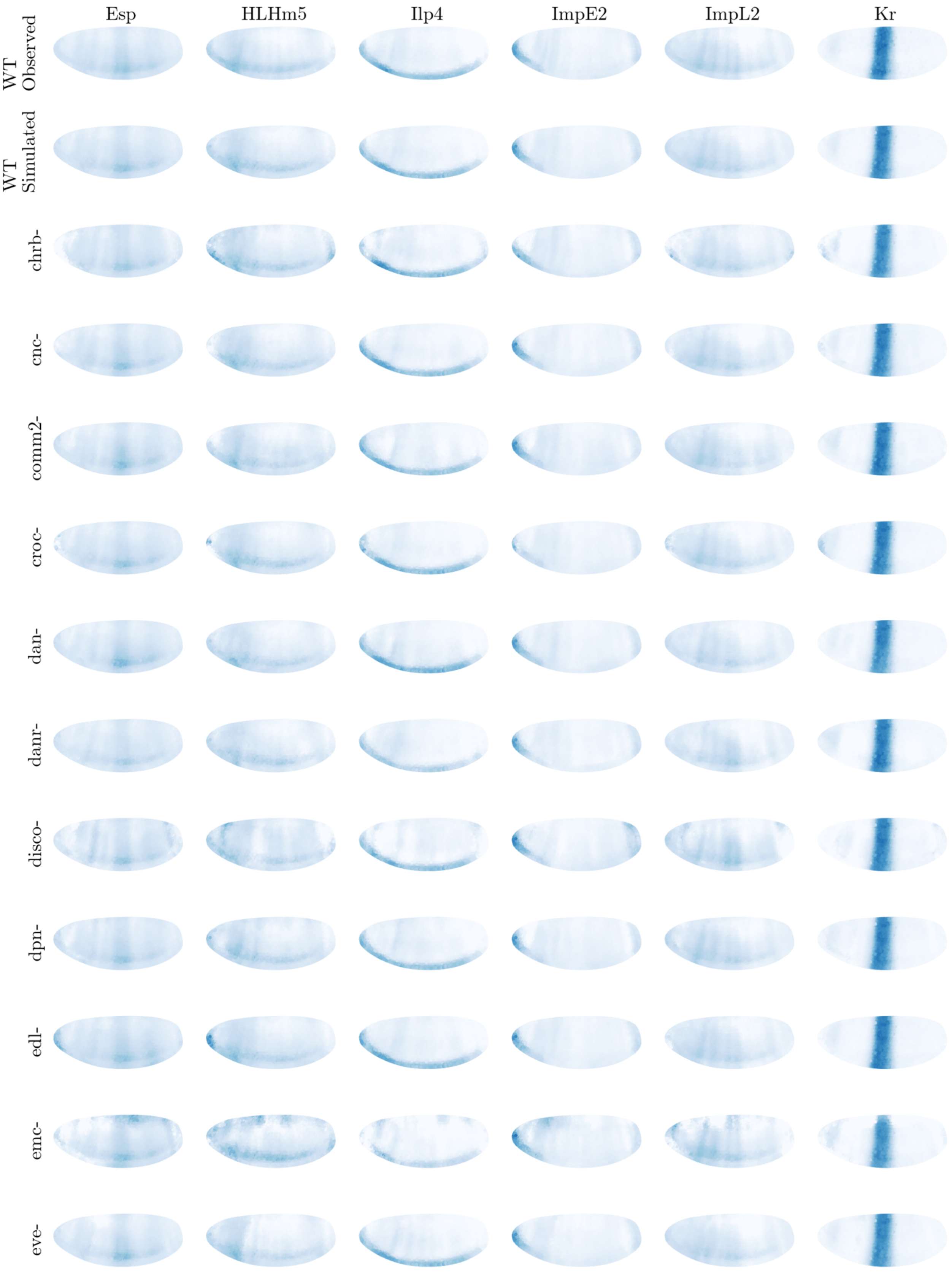
Esp
HLHm5
Ilp4
ImpE2
ImpL2
Kr
WT Observed
WT Simulated
chrb-
cnc-
comm2-
croc-
dan-
danr-
disco-
dpn-
edl-
emc-
eve-

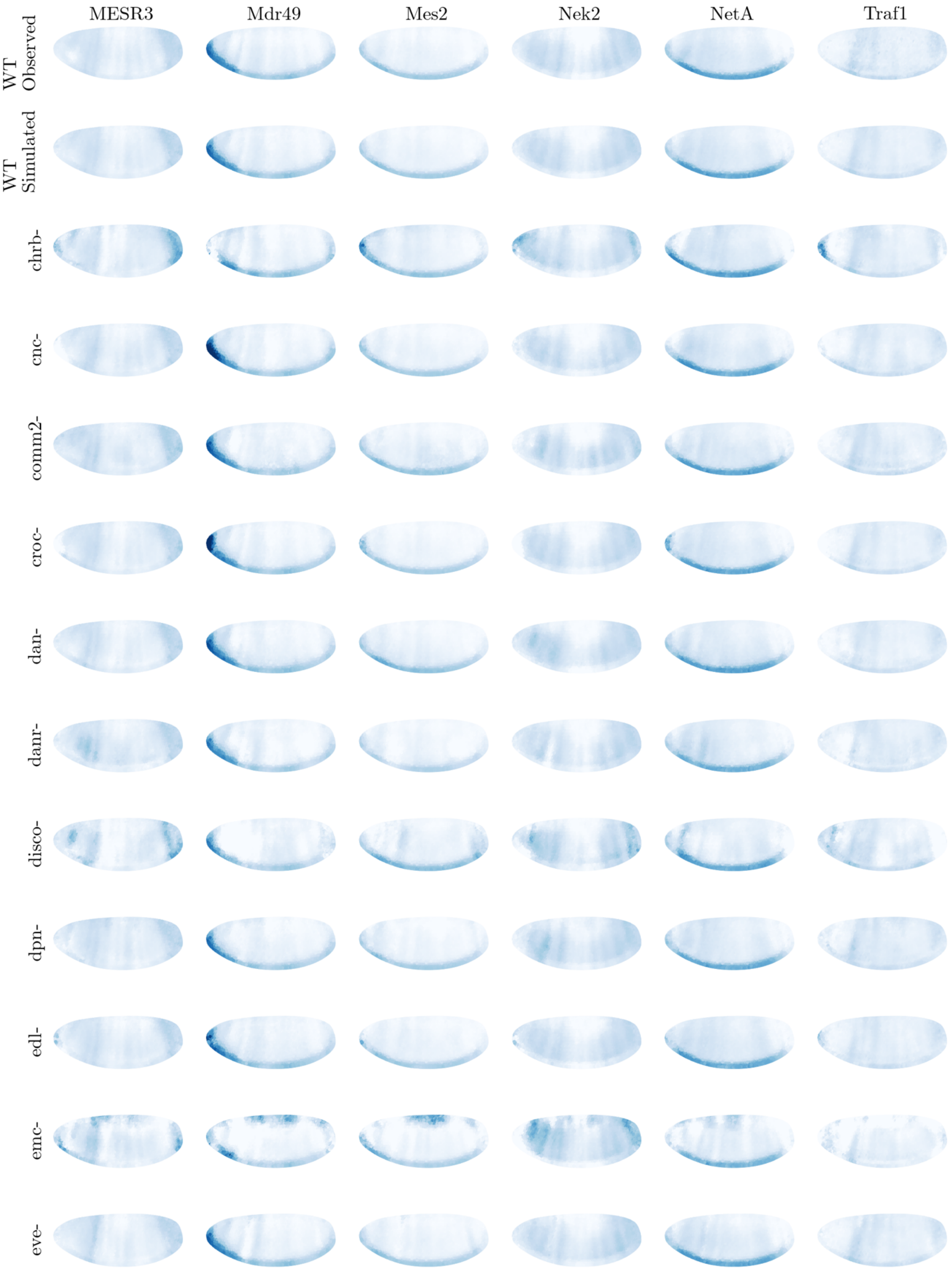
MESR3
Mdr49
Mes2
Nek2
NetA
Traf1
WT Observed
WT Simulated
chrb-
cnc-
comm2-
croc-
dan-
danr-
disco-
dpn-
edl-
emc-
eve-

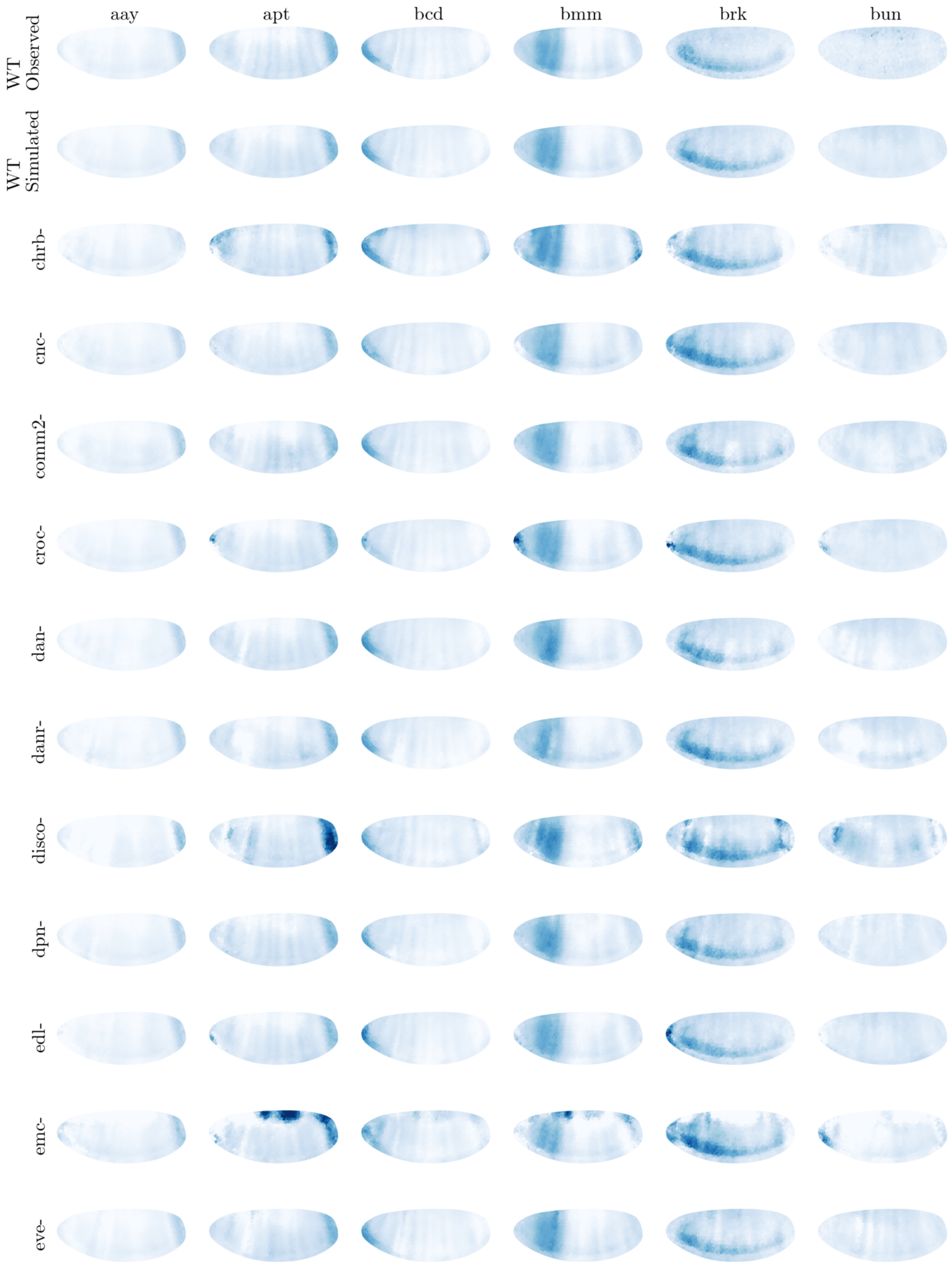
aay
apt
bcd
bmm
brk
bun
WT Observed
WT Simulated
chrb-
cnc-
comm2-
croc-
dan-
danr-
disco-
dpn-
edl-
emc-
eve-

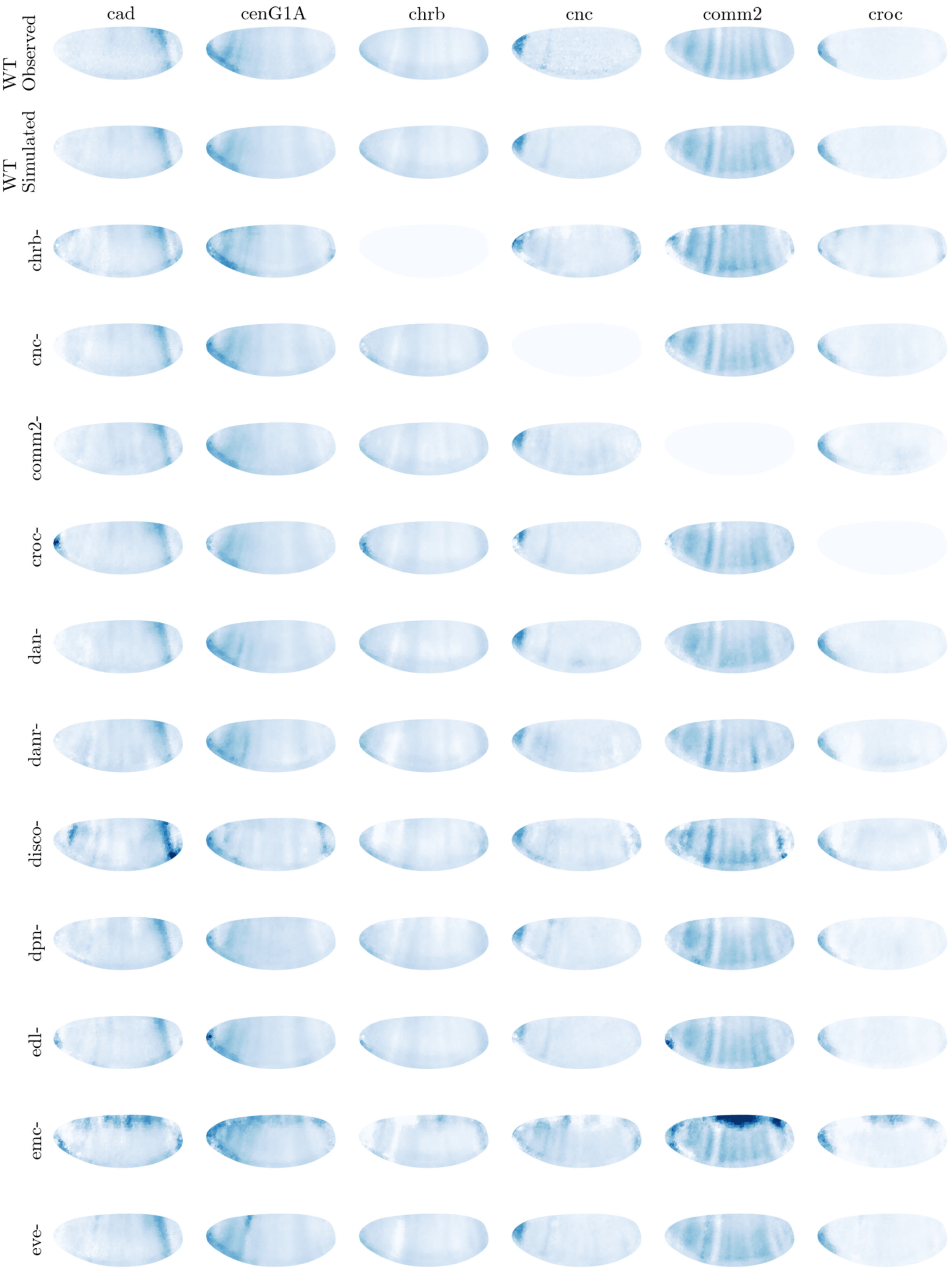
cad
cenG1A
chrb
cnc
comm2
croc
WT Observed
WT Simulated
chrb-
cnc-
comm2-
croc-
dan-
danr-
disco-
dpn-
edl-
emc-
eve-

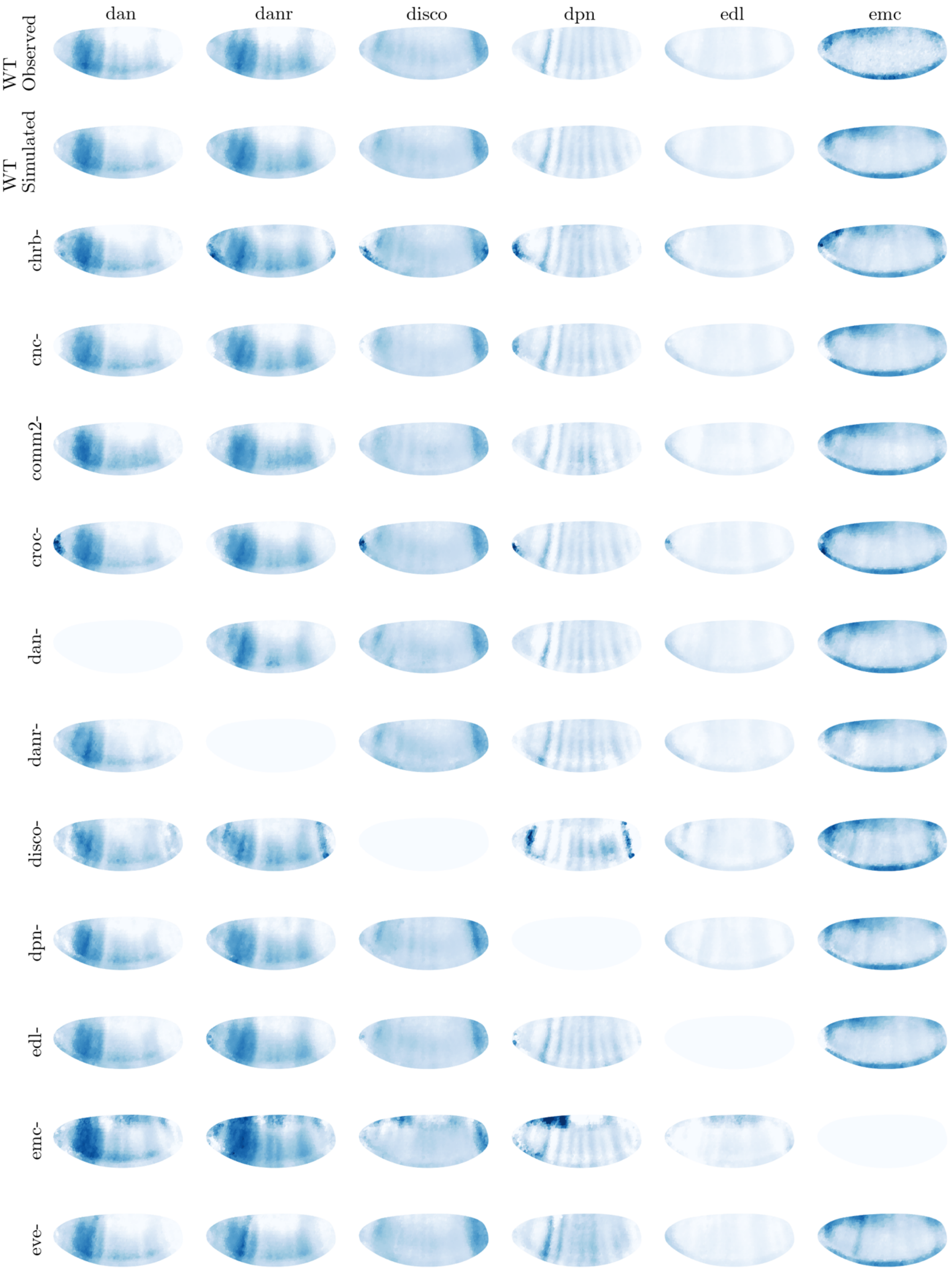

dan
danr
disco
dpn
edl
emc
WT Observed
WT Simulated
chrb-
cnc-
comm2-
croc-
dan-
danr-
disco-
dpn-
edl-
emc-
eve-

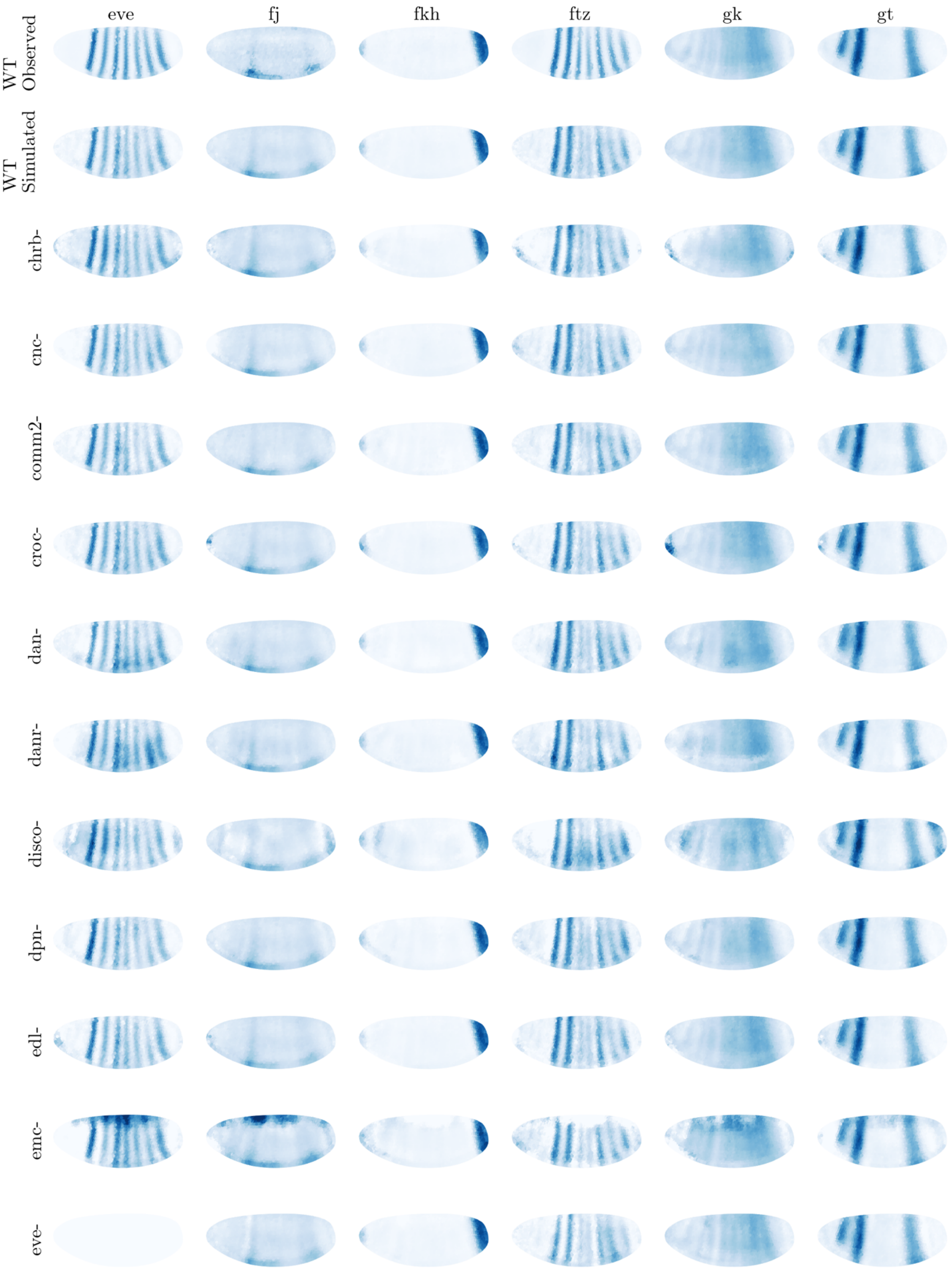
eve
fj
fkh
ftz
gk
gt
WT Observed
WT Simulated
chrb-
cnc-
comm2-
croc-
dan-
danr-
disco-
dpn-
edl-
emc-
eve-

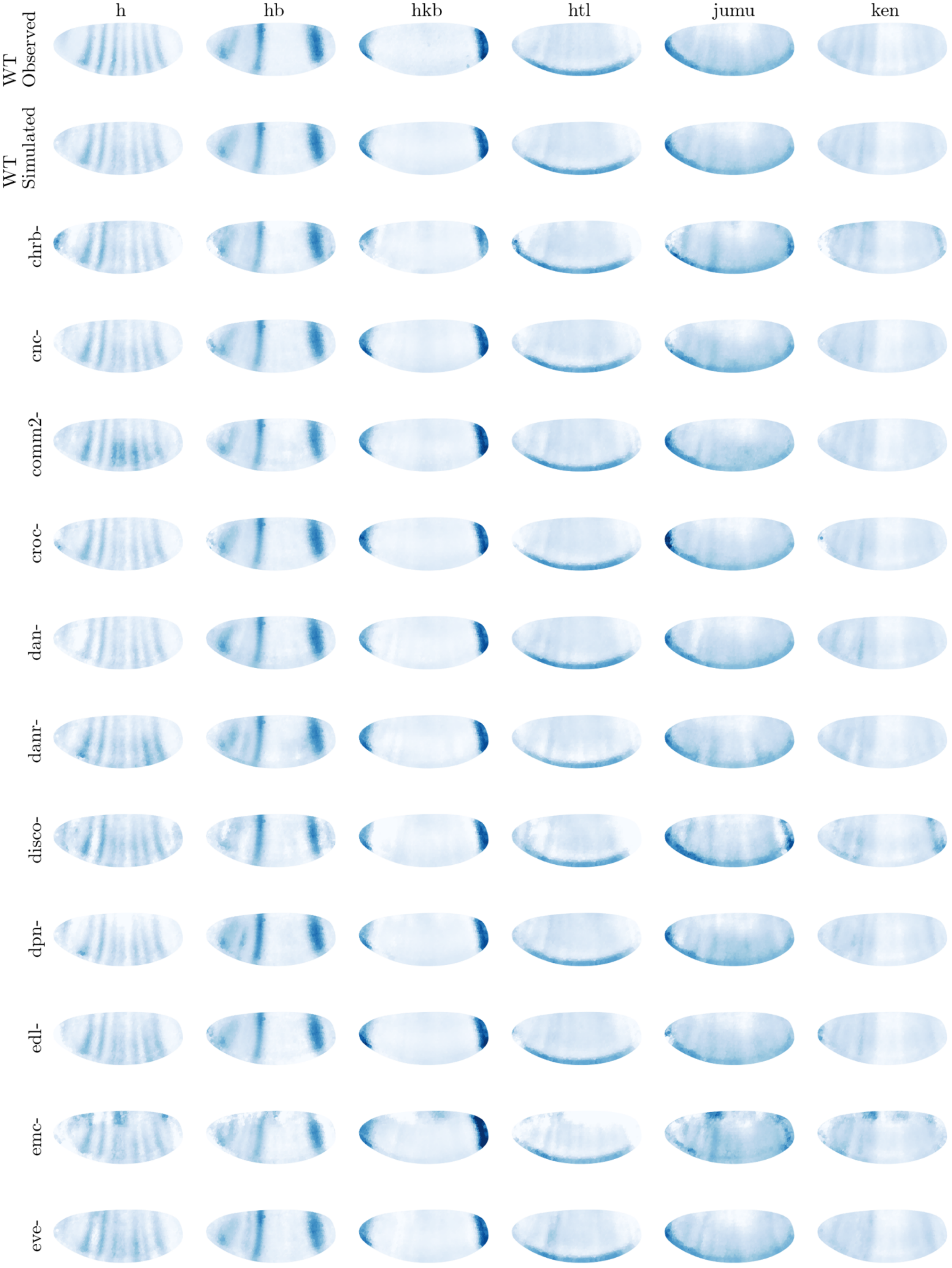
h
hb
hkb
htl
jumu
ken
WT Observed
WT Simulated
chrb-
cnc-
comm2-
croc-
dan-
danr-
disco-
dpn-
edl-
emc-
eve-

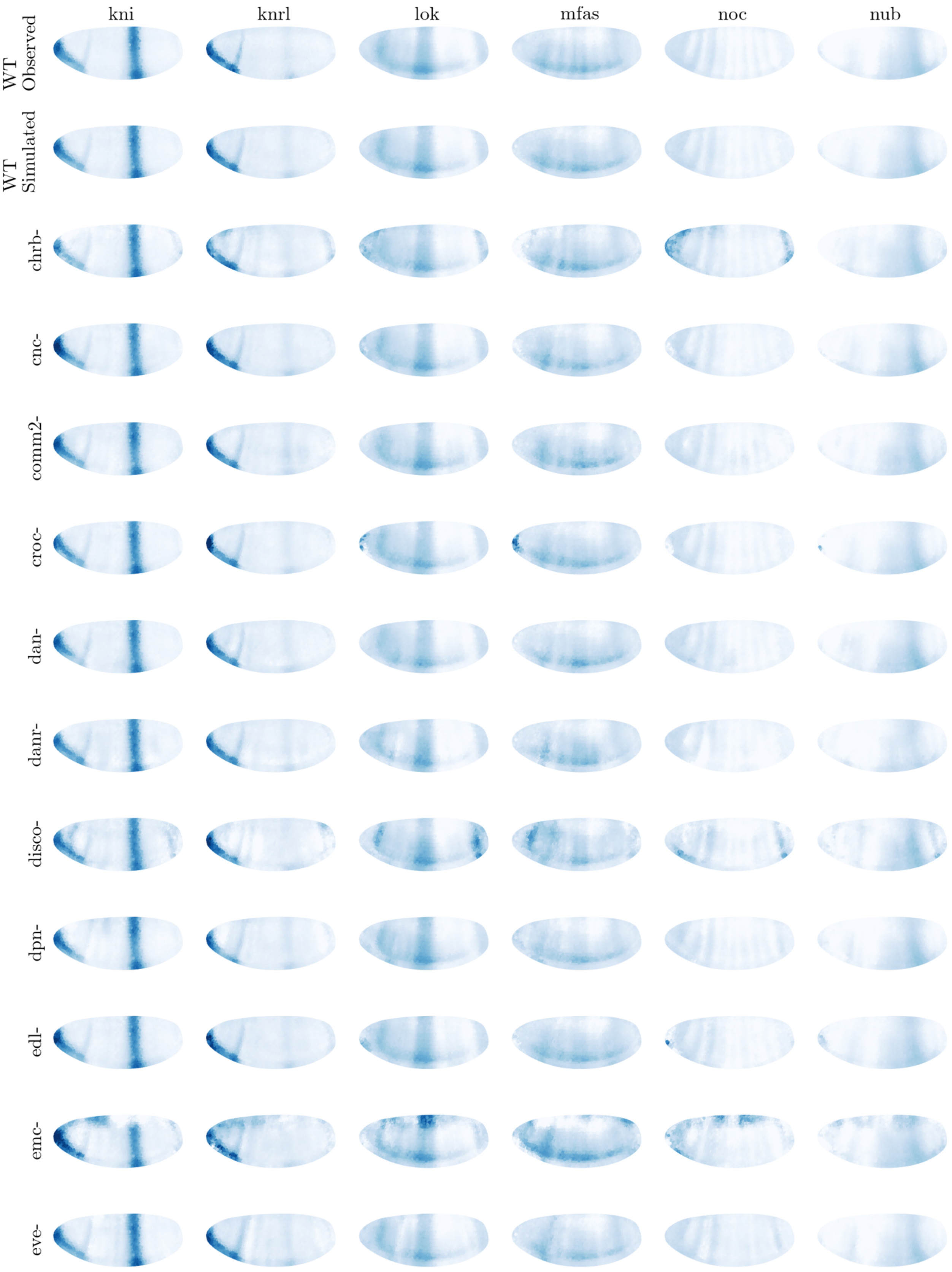
kni
knrl
lok
mfas
noc
nub
WT Observed
WT Simulated
chrb-
cnc-
comm2-
croc-
dan-
danr-
disco-
dpn-
edl-
emc-
eve-

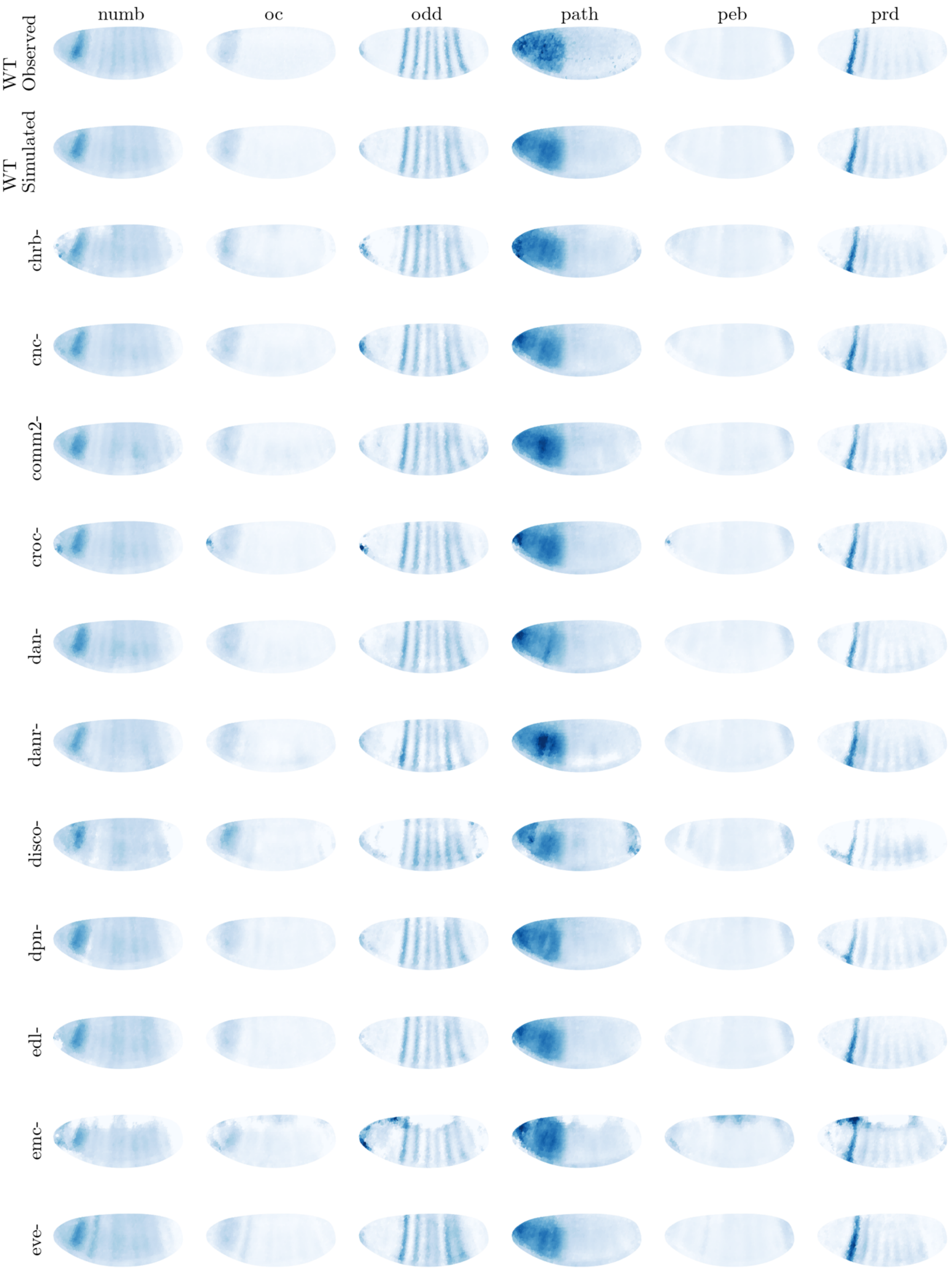
numb
oc
odd
path
peb
prd
WT Observed
WT Simulated
chrb-
cnc-
comm2-
croc-
dan-
danr-
disco-
dpn-
edl-
emc-
eve-

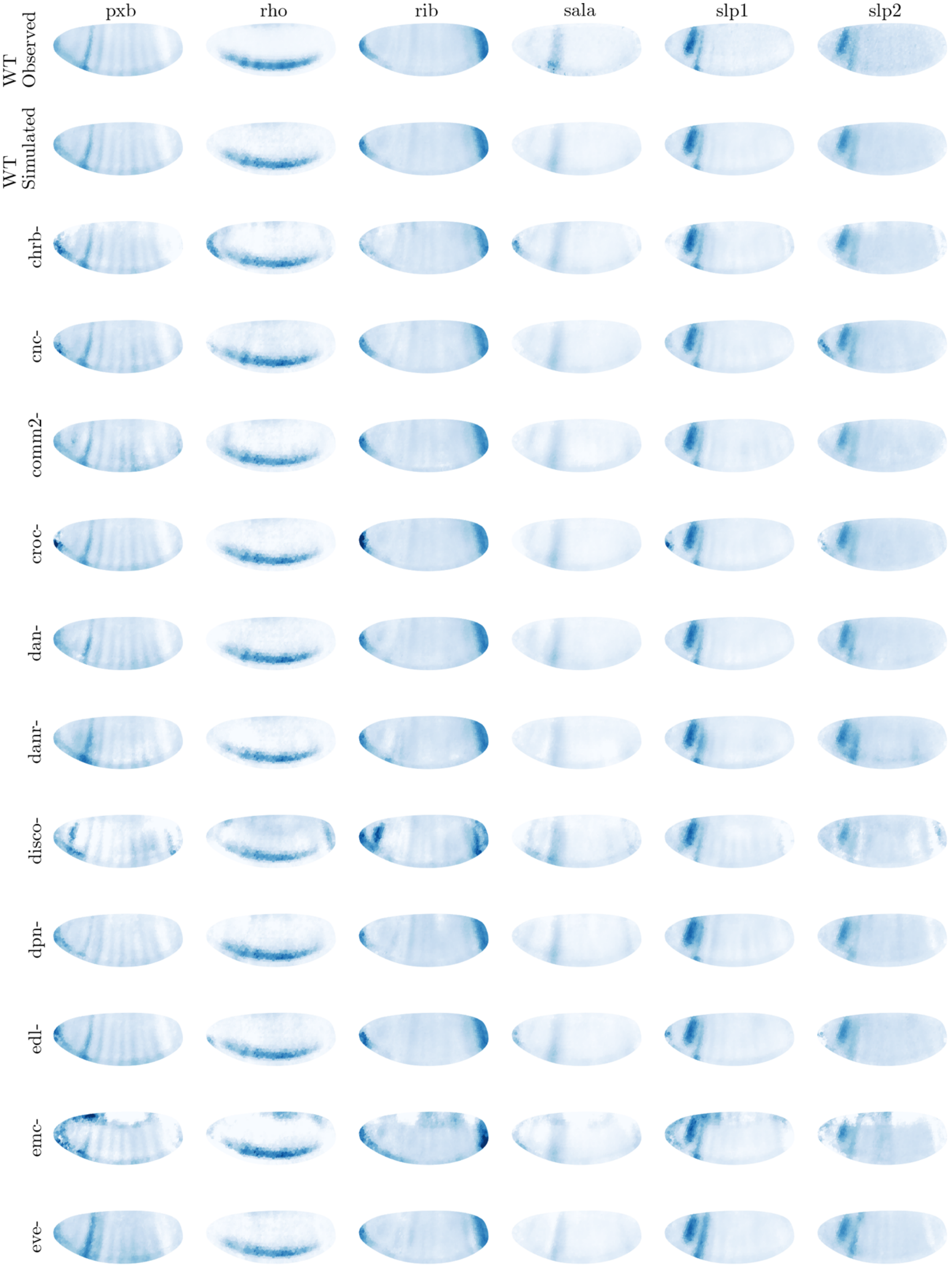
pxb
rho
rib
sala
slp1
slp2
WT Observed
WT Simulated
chrb-
cnc-
comm2-
croc-
dan-
danr-
disco-
dpn-
edl-
emc-
eve-

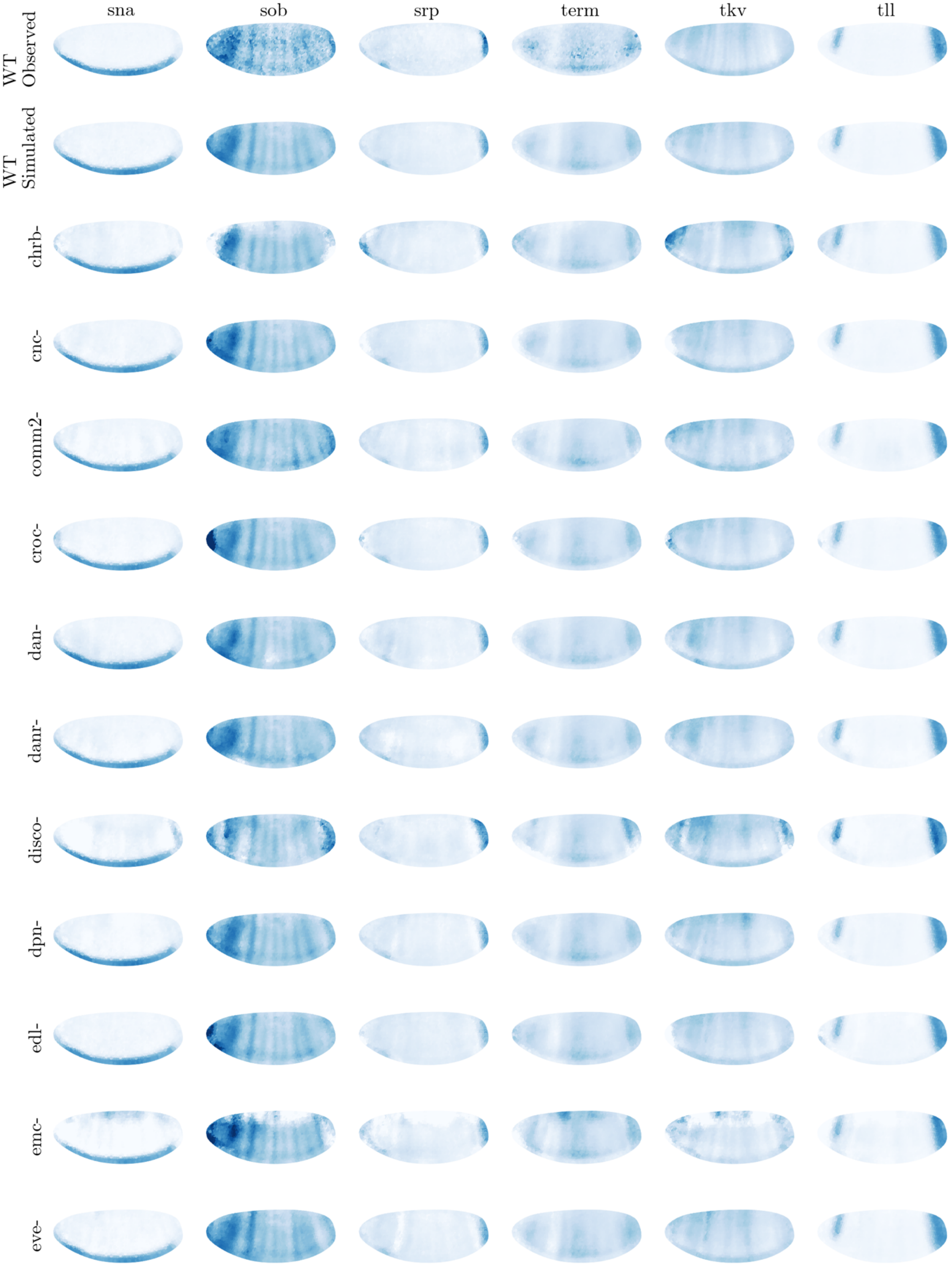
sna
sob
srp
term
tkv
tll
WT Observed
WT Simulated
chrb-
cnc-
comm2-
croc-
dan-
danr-
disco-
dpn-
edl-
emc-
eve-

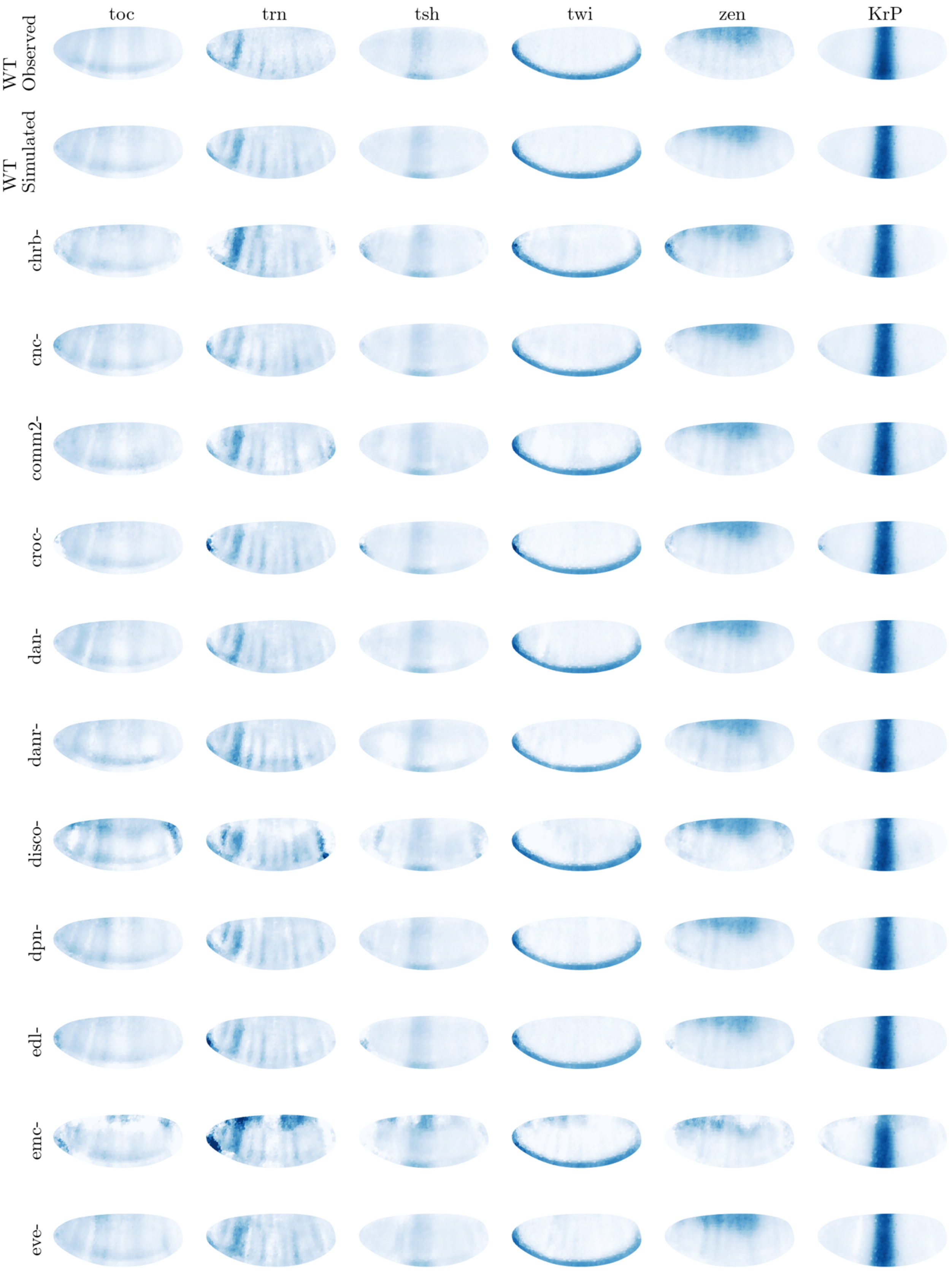
toc
trn
tsh
twi
zen
KrP
WT Observed
WT Simulated
chrb-
cnc-
comm2-
croc-
dan-
danr-
disco-
dpn-
edl-
emc-
eve-

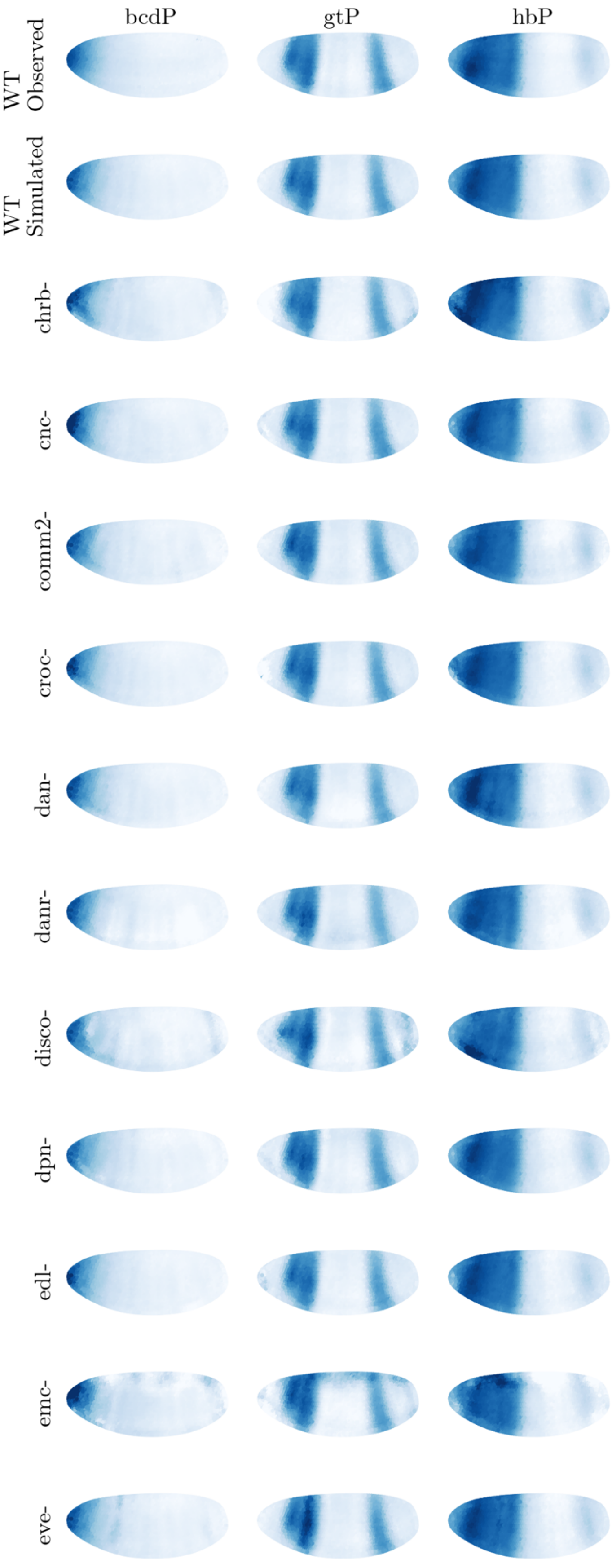
bcdP
gtP
hbP
WT Observed
WT Simulated
chrb-
cnc-
comm2-
croc-
dan-
danr-
disco-
dpn-
edl-
emc-
eve-

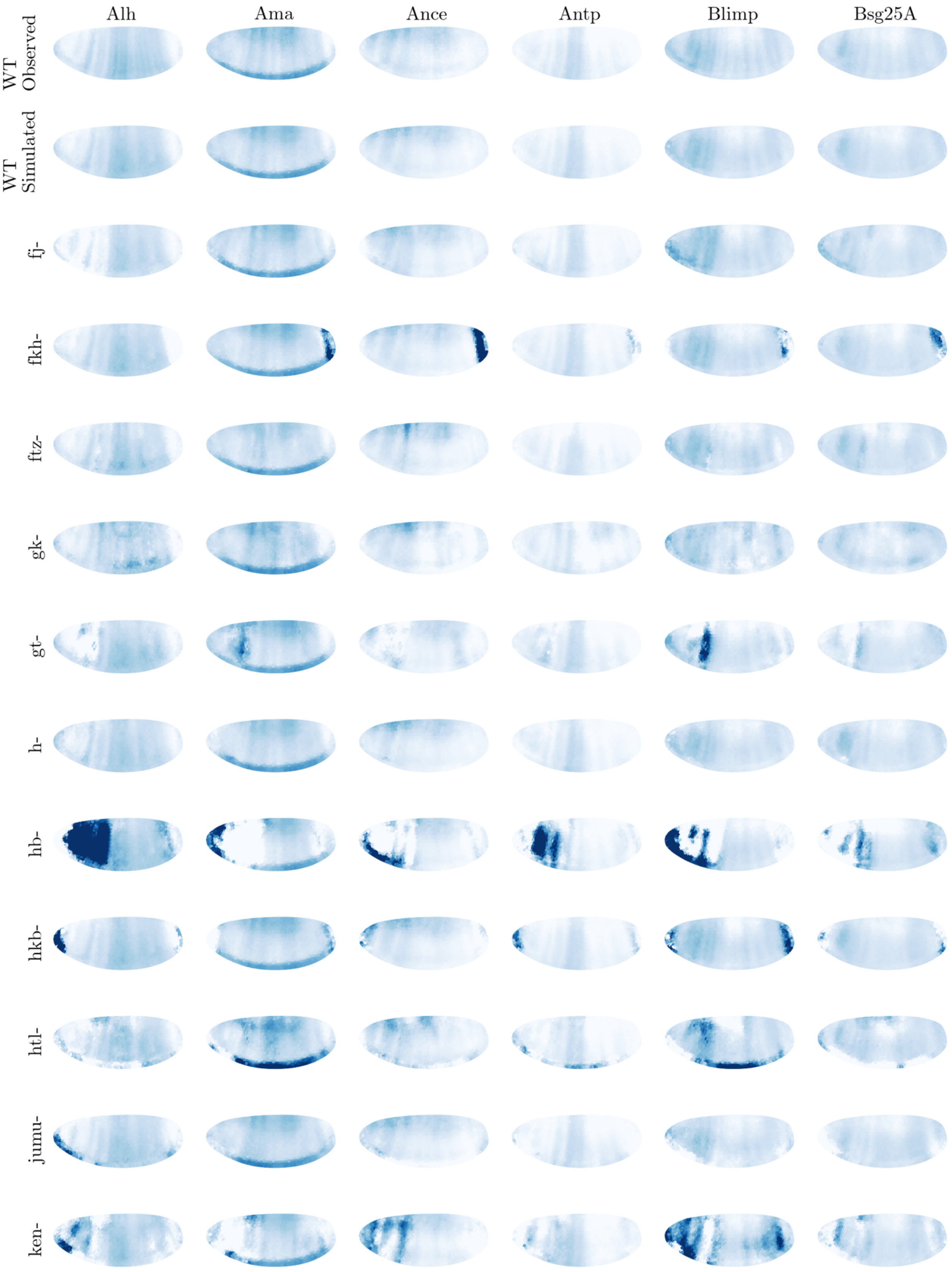
Alh
Ama
Ance
Antp
Blimp
Bsg25A
WT Observed
WT Simulated
fj-
fkh-
ftz-
gk-
gt-
h-
hb-
hkb-
htl-
jumu-
ken-

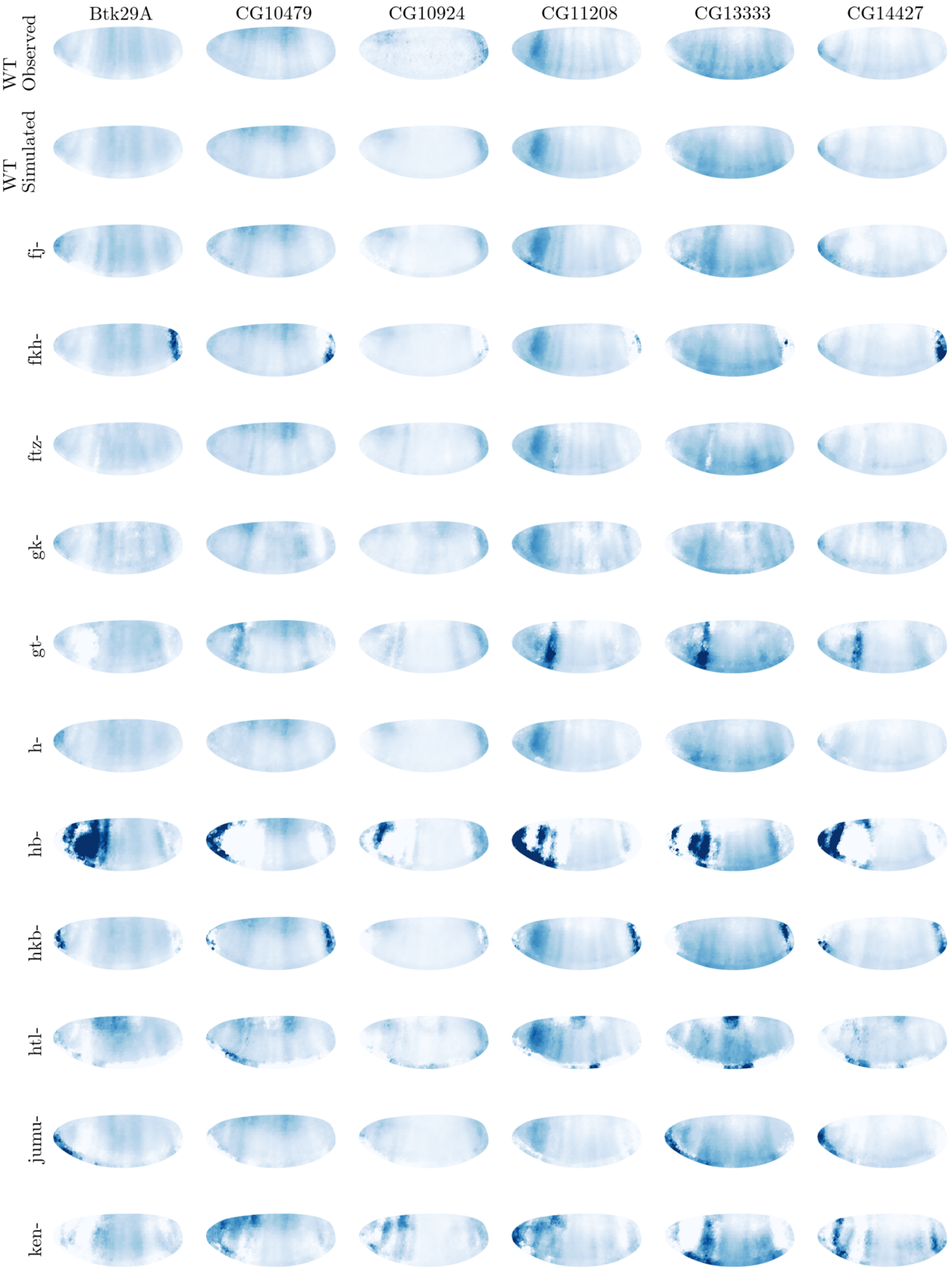

Btk29A
CG10479
CG10924
CG11208
CG13333
CG14427
WT Observed
WT Simulated
fj-
fkh-
ftz-
gk-
gt-
h-
hb-
hkb-
htl-
jumu-
ken-

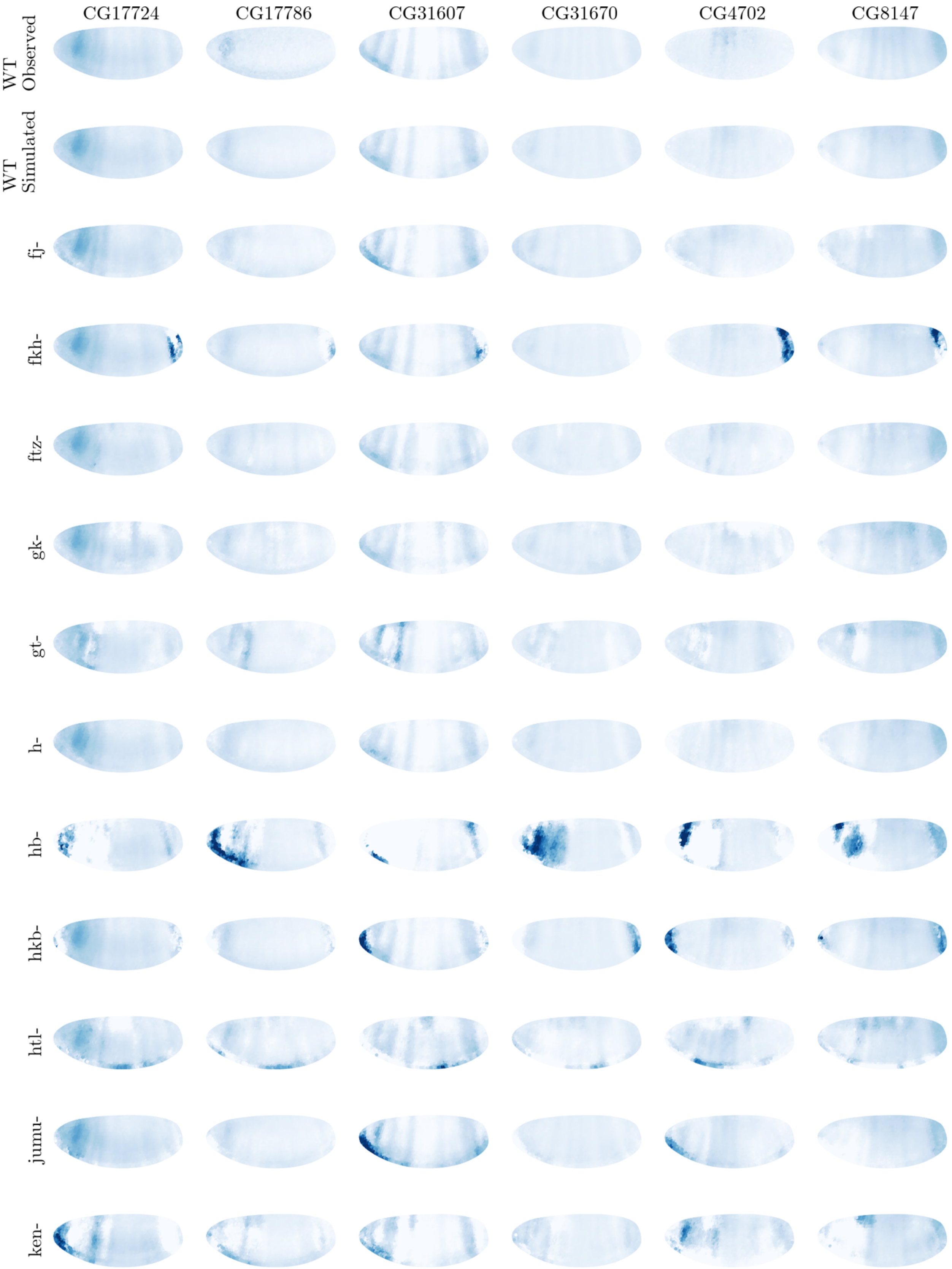
CG17724
CG17786
CG31607
CG31670
CG4702
CG8147
WT Observed
WT Simulated
fj-
fkh-
ftz-
gk-
gt-
h-
hb-
hkb-
htl-
jumu-
ken-

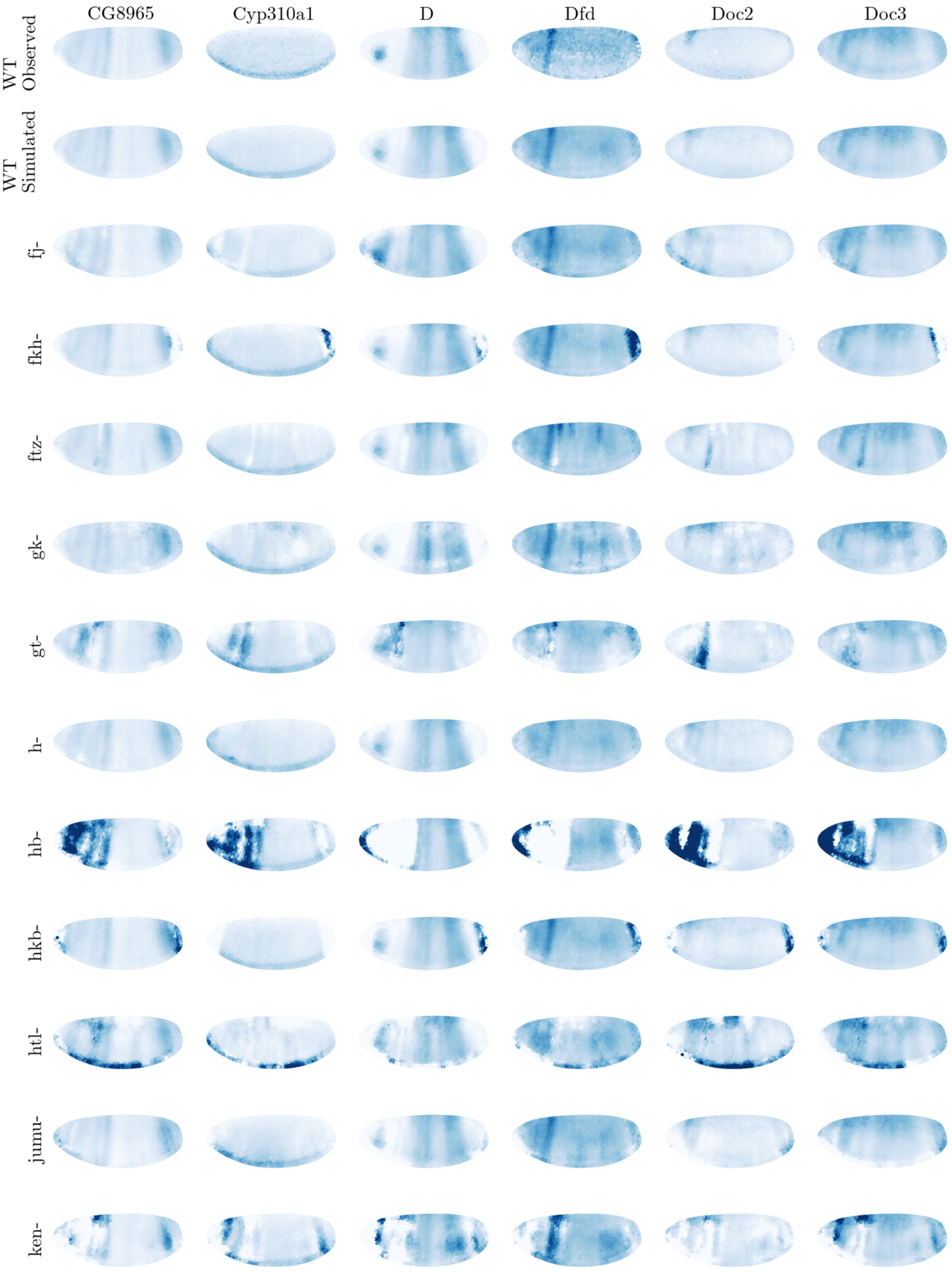
CG8965
Cyp310a1
D
Dfd
Doc2
Doc3
WT Observed
WT Simulated
fj-
fkh-
ftz-
gk-
gt-
h-
hb-
hkb-
htl-
jumu-
ken-

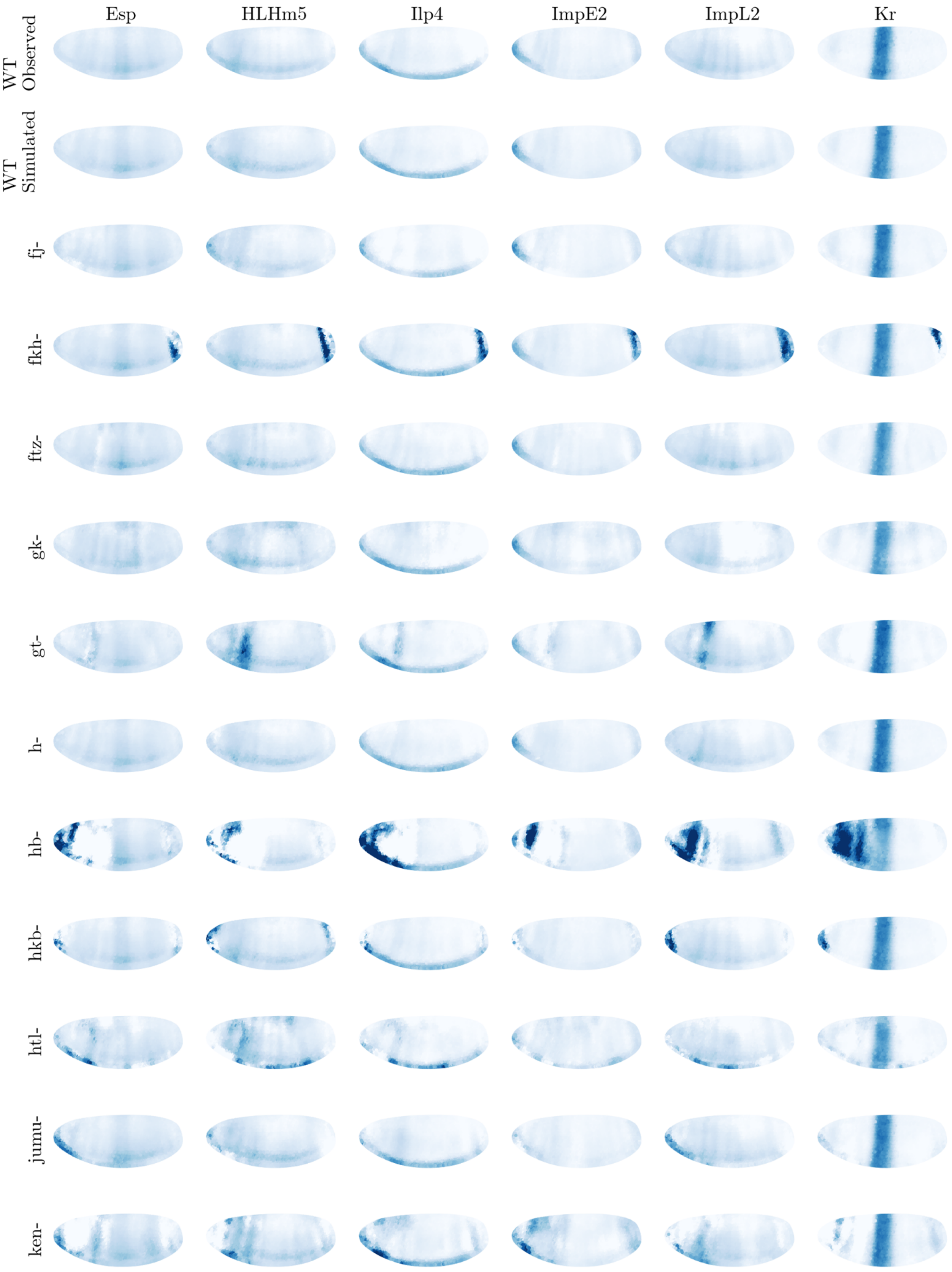
Esp
HLHm5
Ilp4
ImpE2
ImpL2
Kr
WT Observed
WT Simulated
fj-
fkh-
ftz-
gk-
gt-
h-
hb-
hkb-
htl-
jumu-
ken-

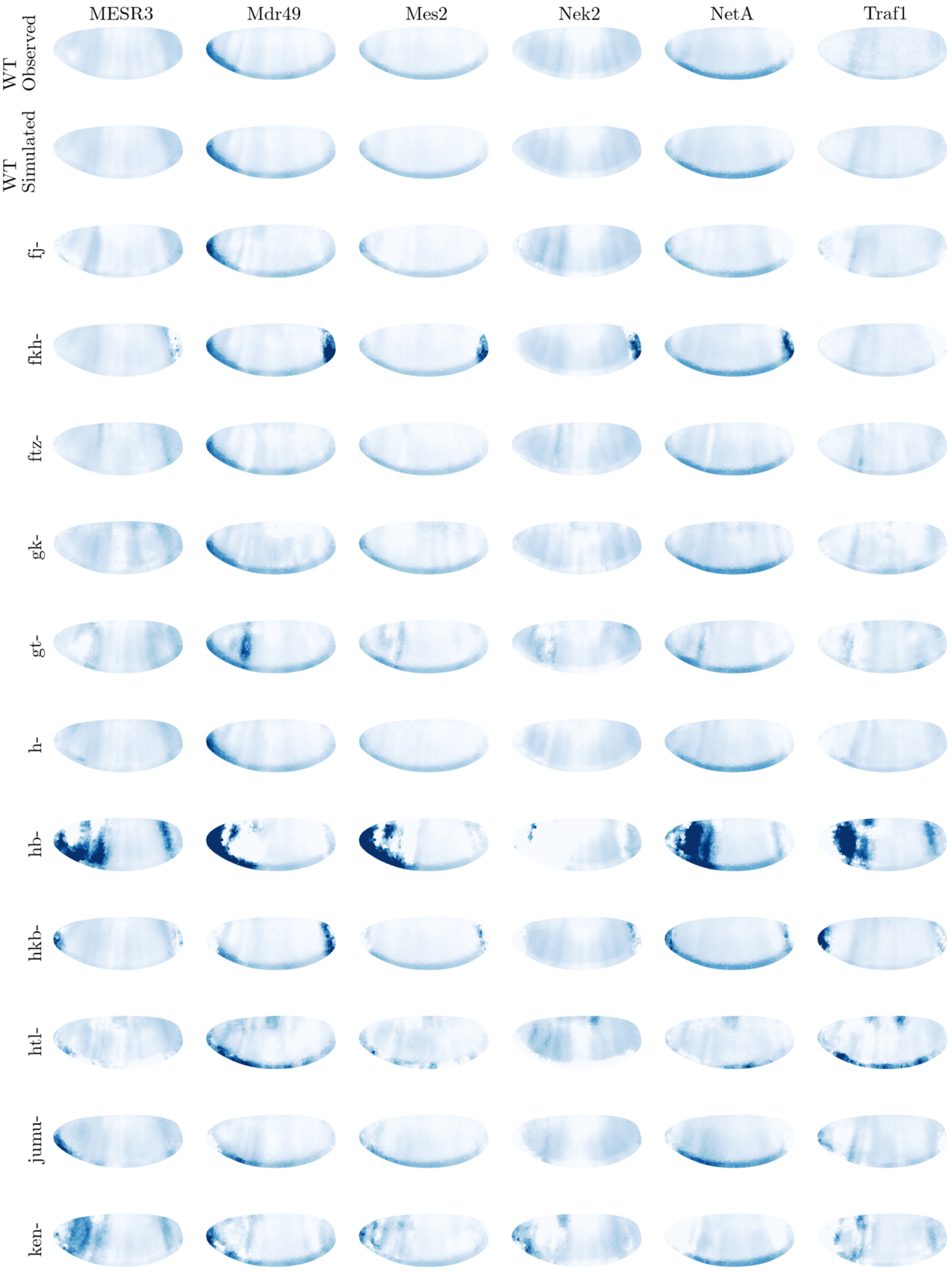
MESR3
Mdr49
Mes2
Nek2
NetA
Traf1
WT Observed
WT Simulated
fj-
fkh-
ftz-
gk-
gt-
h-
hb-
hkb-
htl-
jumu-
ken-

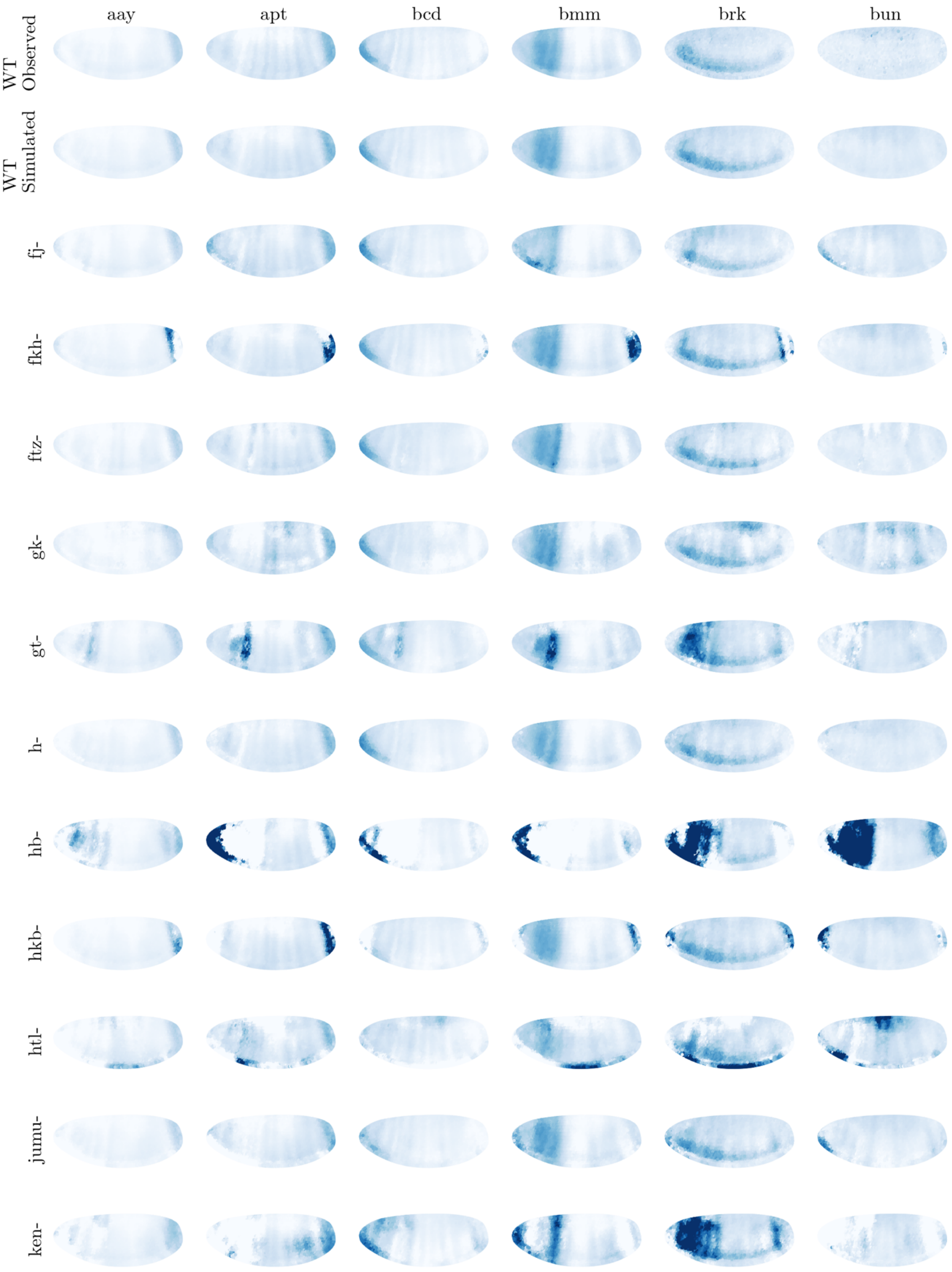
aay
apt
bcd
bmm
brk
bun
WT Observed
WT Simulated
fj-
fkh-
ftz-
gk-
gt-
h-
hb-
hkb-
htl-
jumu-
ken-

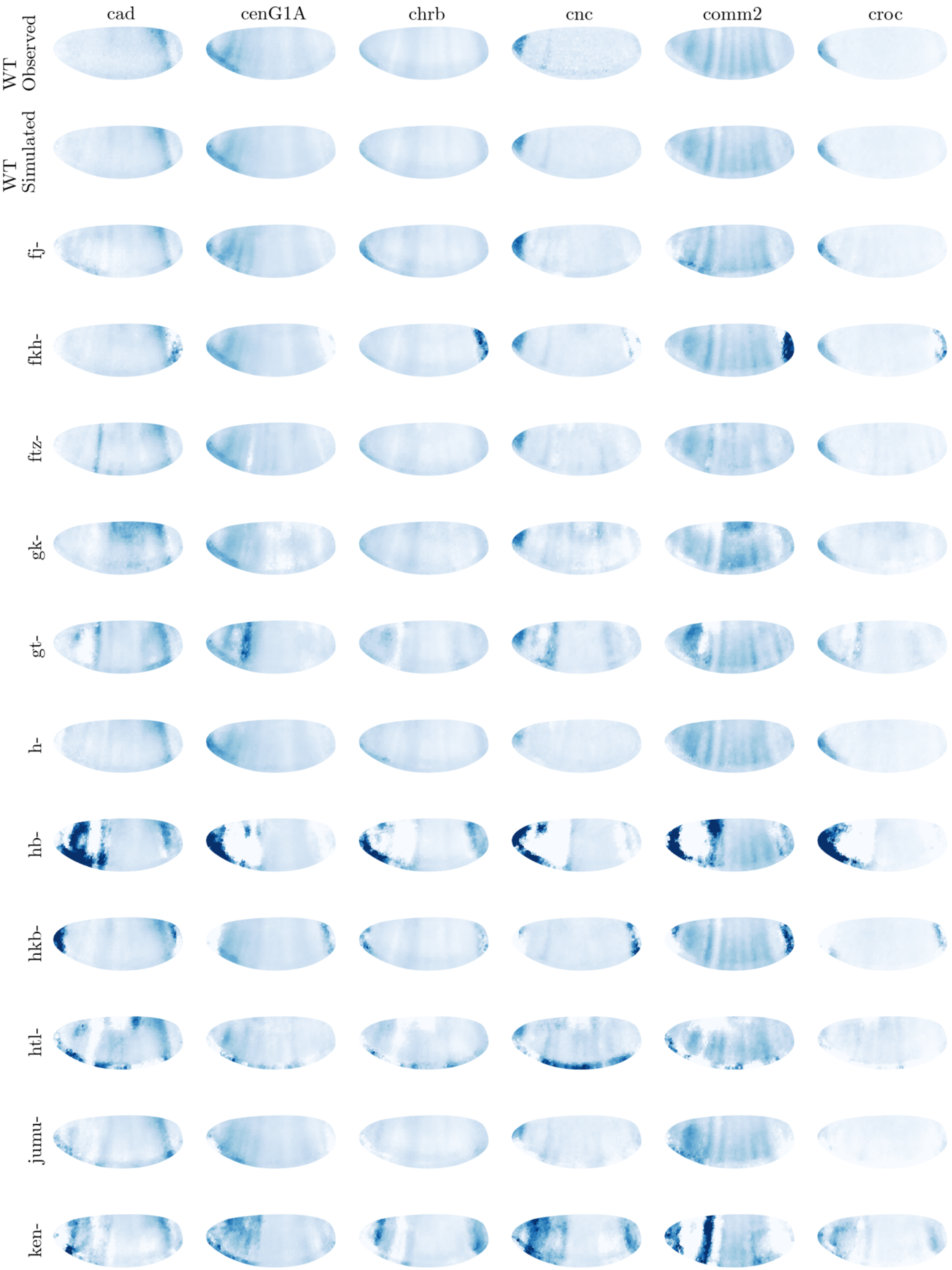

cad
cenG1A
chrb
cnc
comm2
croc
WT Observed
WT Simulated
fj-
fkh-
ftz-
gk-
gt-
h-
hb-
hkb-
htl-
jumu-
ken-

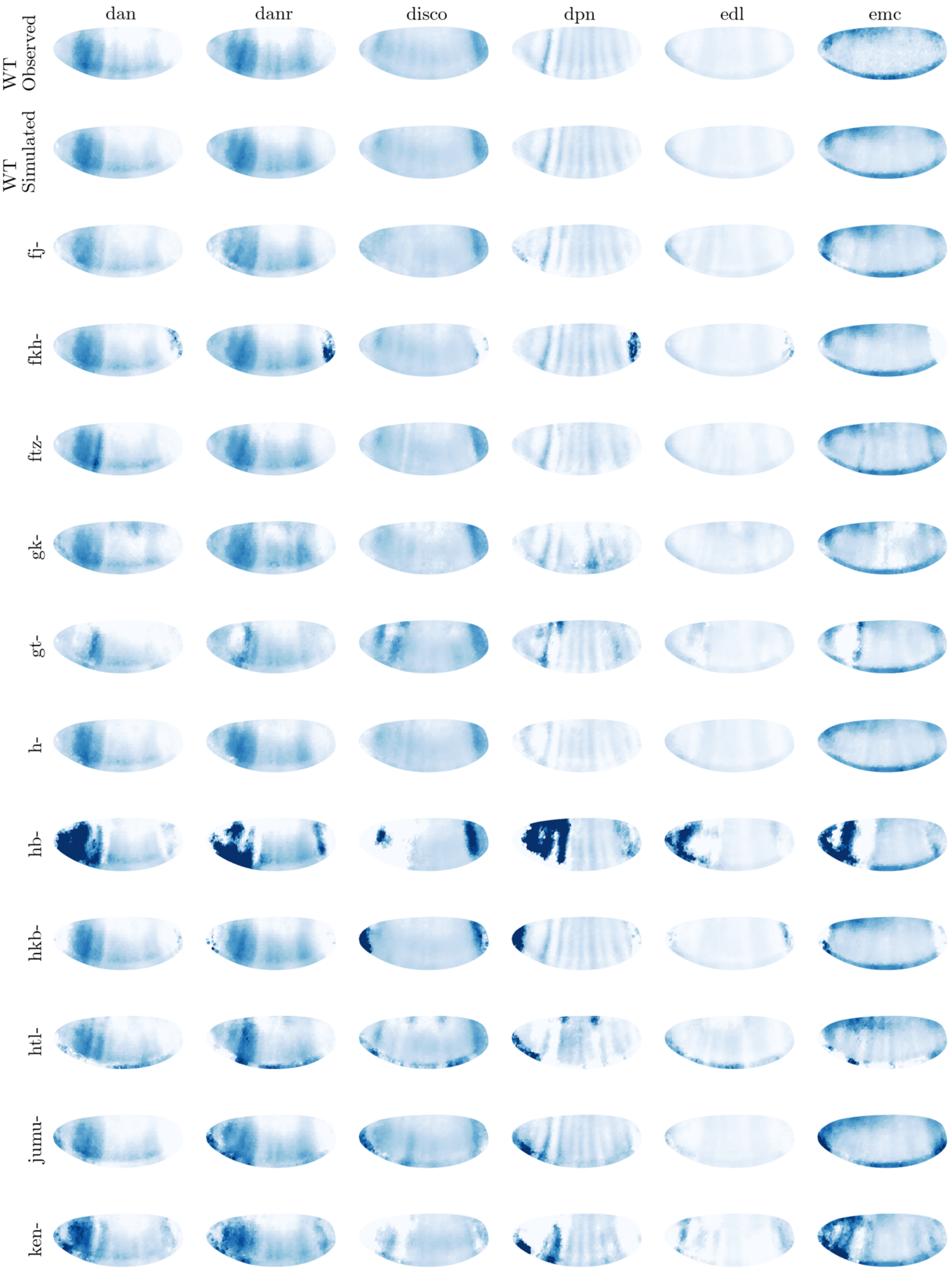
dan
danr
disco
dpn
edl
emc
WT Observed
WT Simulated
fj-
fkh-
ftz-
gk-
gt-
h-
hb-
hkb-
htl-
jumu-
ken-

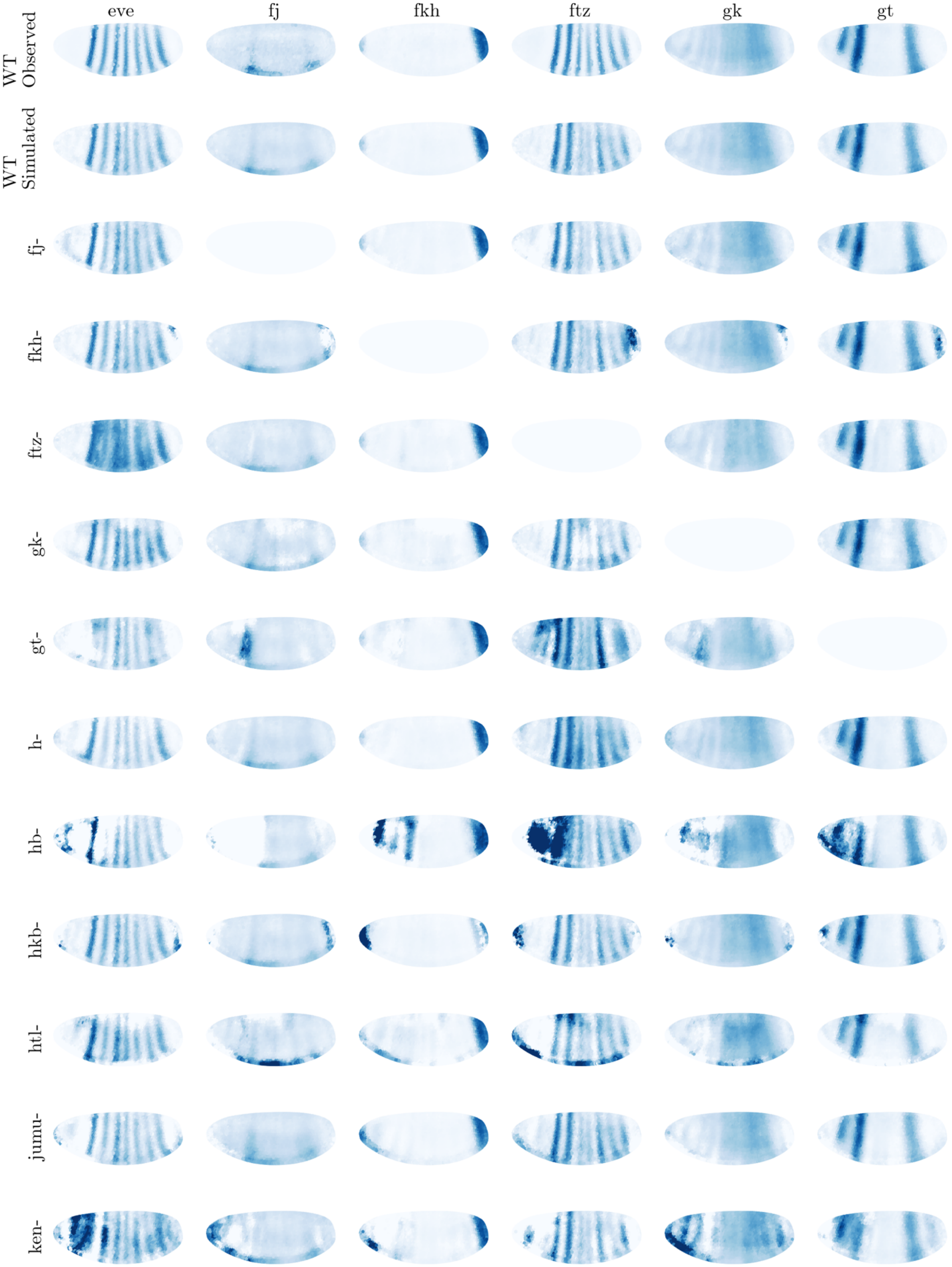

eve
fj
fkh
ftz
gk
gt
WT Observed
WT Simulated
fj-
fkh-
ftz-
gk-
gt-
h-
hb-
hkb-
htl-
jumu-
ken-

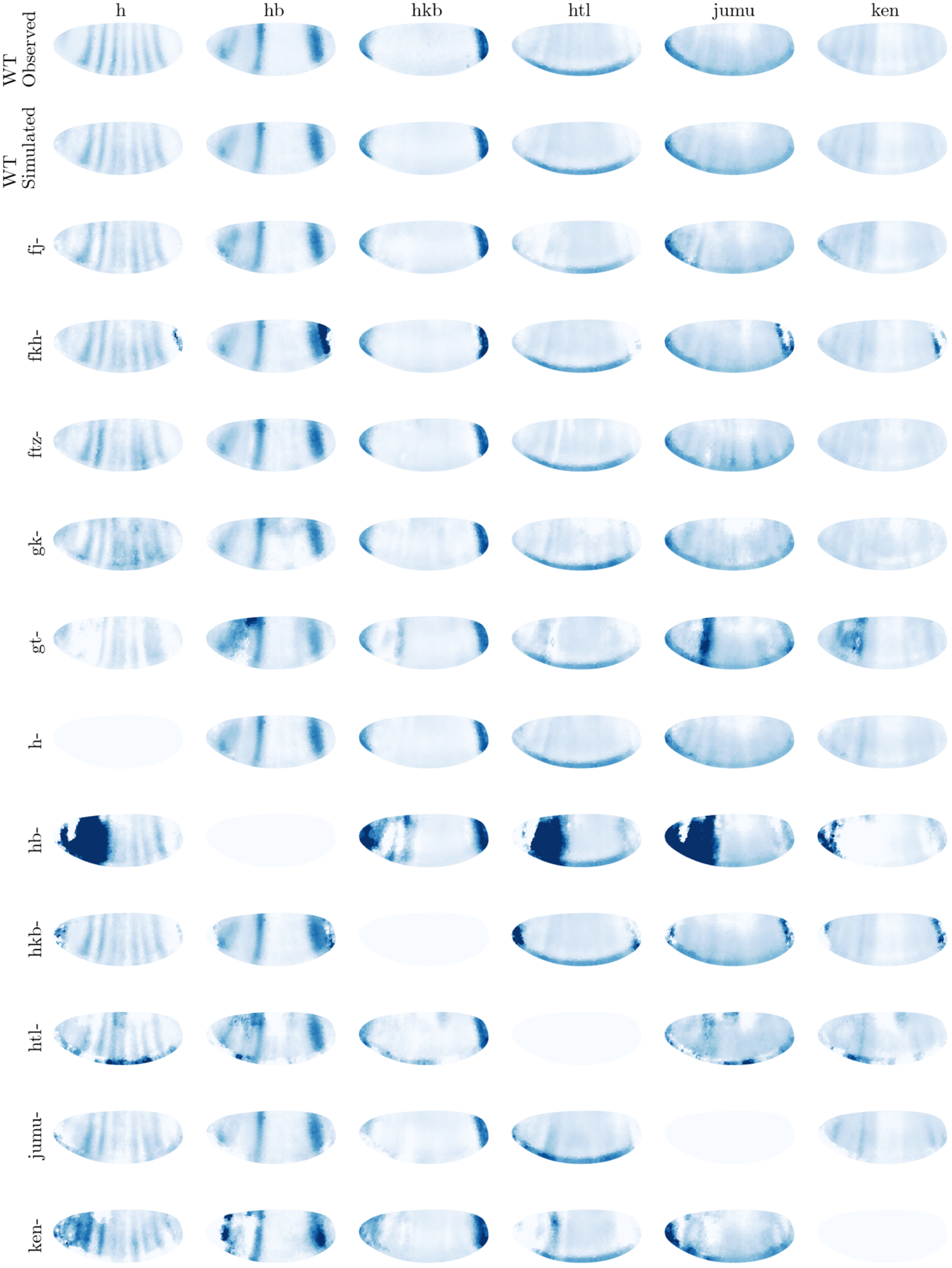
h
hb
hkb
htl
jumu
ken
WT Observed
WT Simulated
fj-
fkh-
ftz-
gk-
gt-
h-
hb-
hkb-
htl-
jumu-
ken-

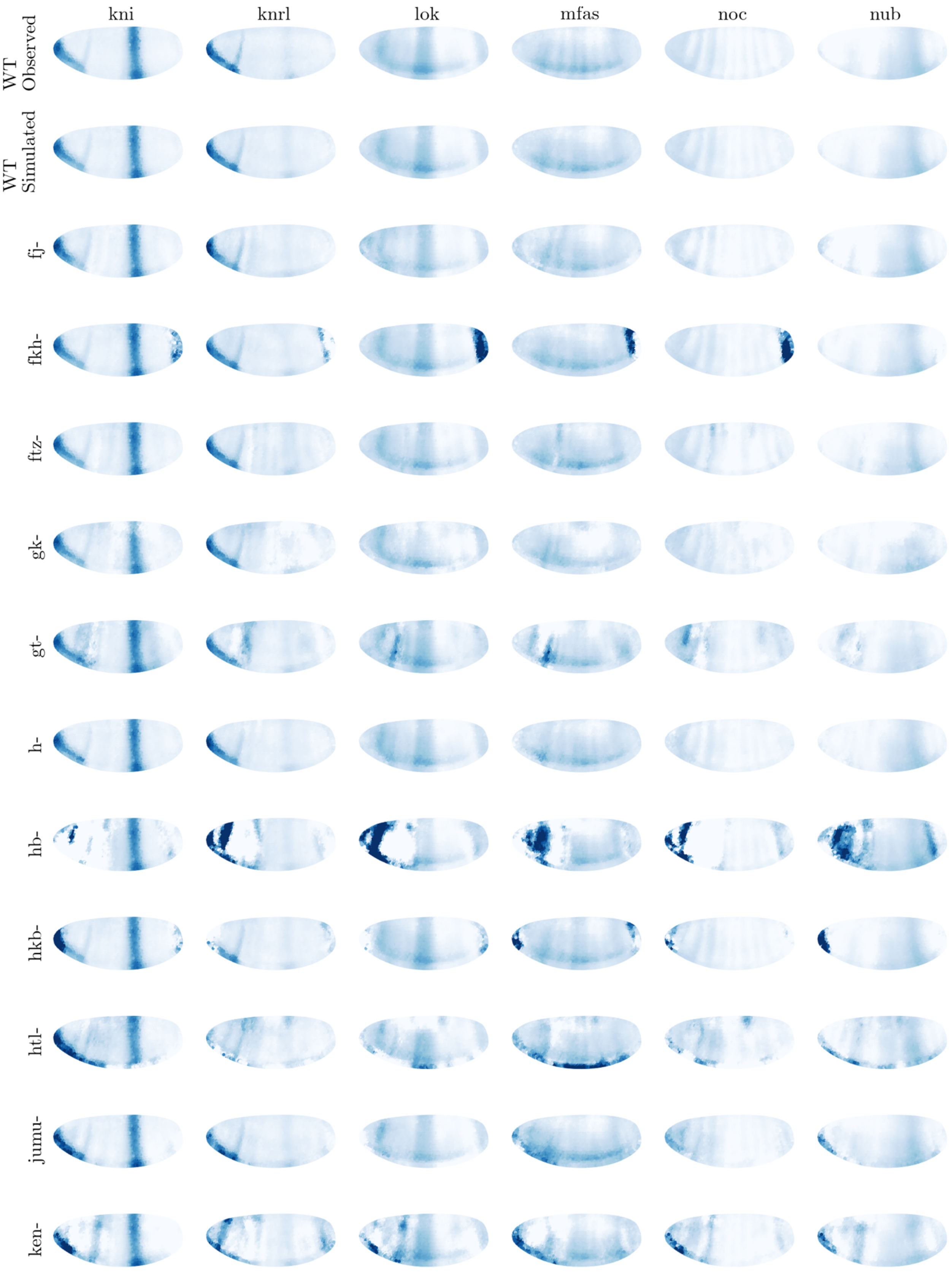
kni
knrl
lok
mfas
noc
nub
WT Observed
WT Simulated
fj-
fkh-
ftz-
gk-
gt-
h-
hb-
hkb-
htl-
jumu-
ken-

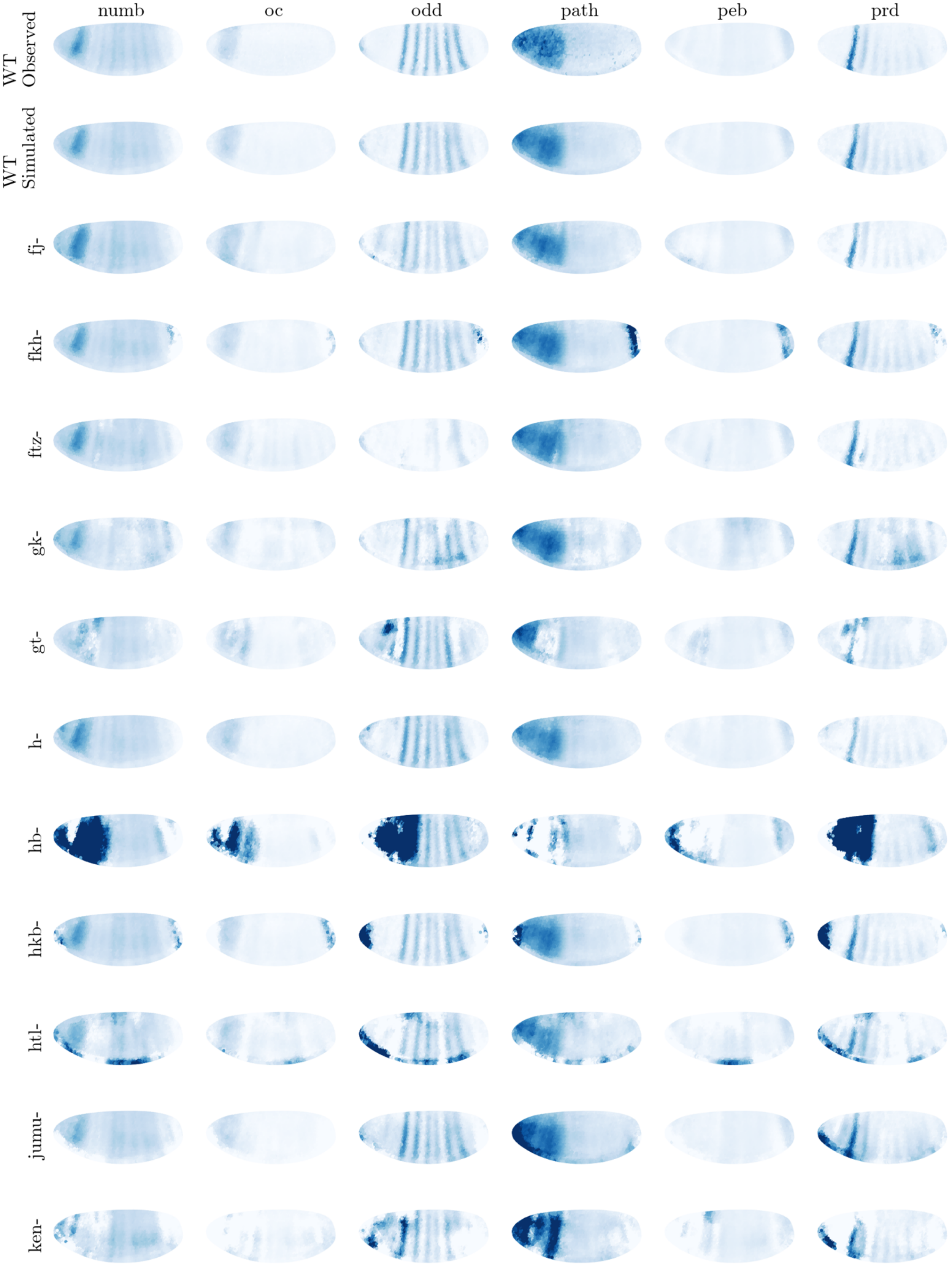
numb
oc
odd
path
peb
prd
WT Observed
WT Simulated
fj-
fkh-
ftz-
gk-
gt-
h-
hb-
hkb-
htl-
jumu-
ken-

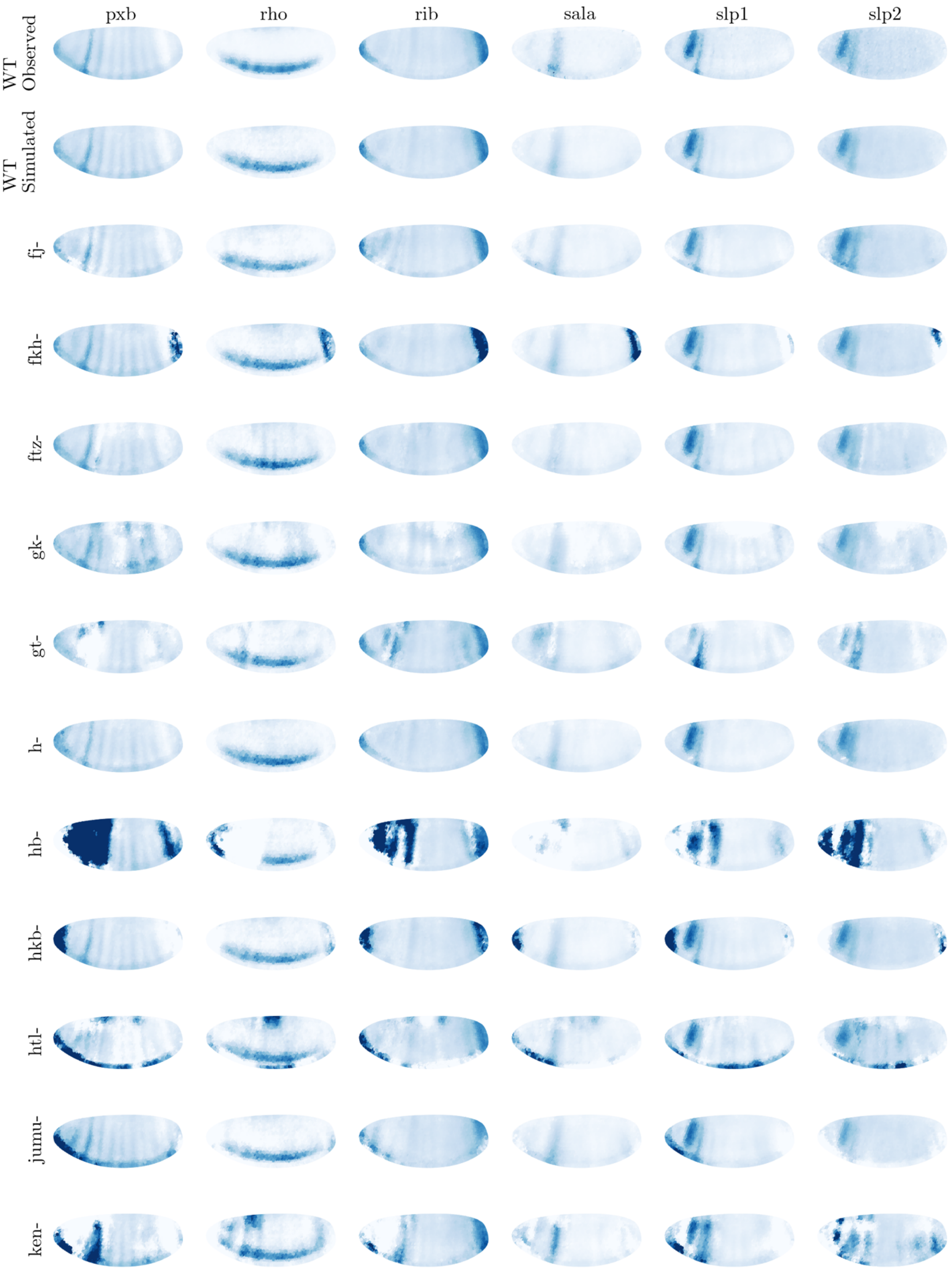

pxb
rho
rib
sala
slp1
slp2
WT Observed
WT Simulated
fj-
fkh-
ftz-
gk-
gt-
h-
hb-
hkb-
htl-
jumu-
ken-

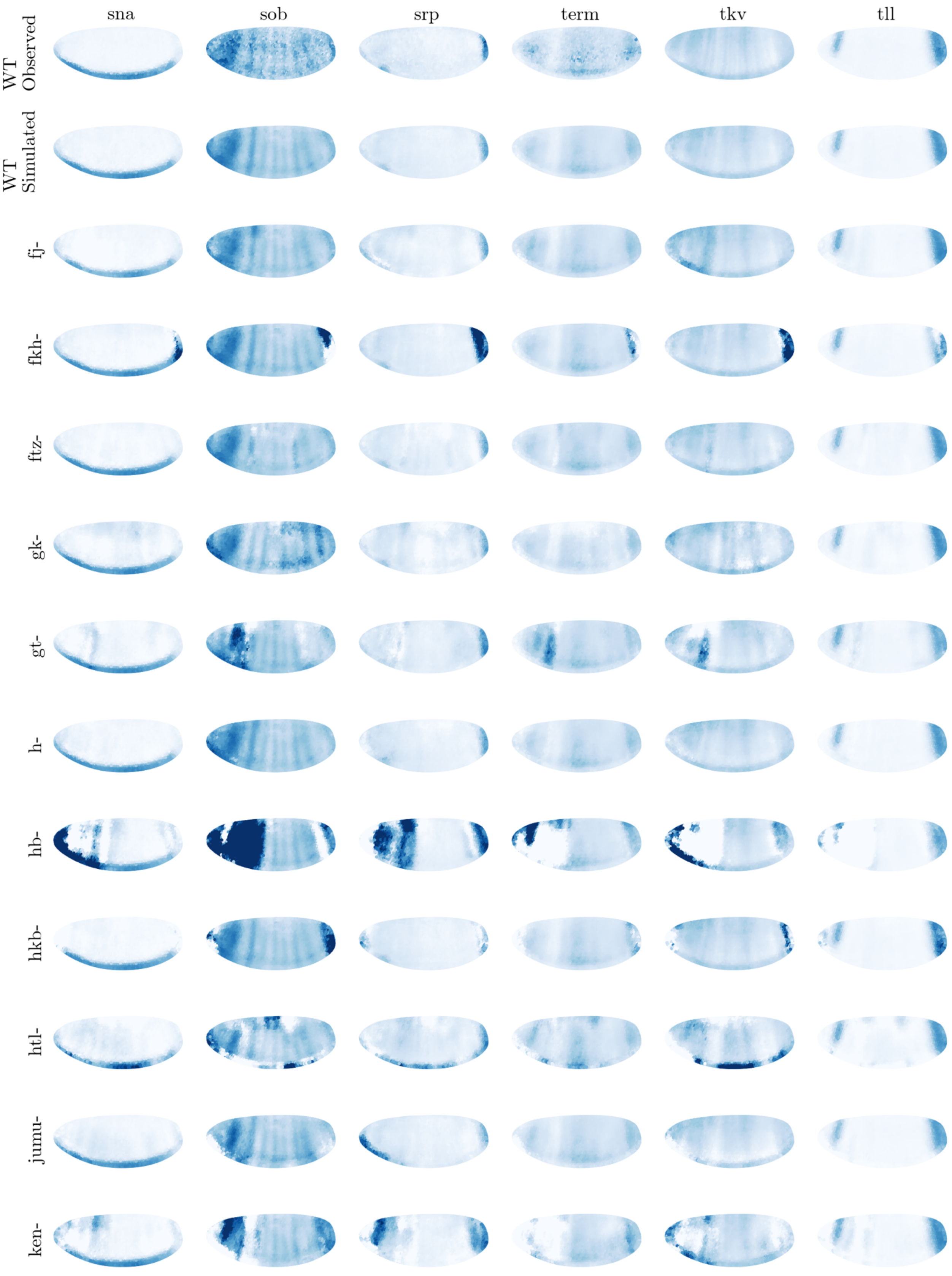
sna
sob
srp
term
tkv
tll
WT Observed
WT Simulated
fj-
fkh-
ftz-
gk-
gt-
h-
hb-
hkb-
htl-
jumu-
ken-

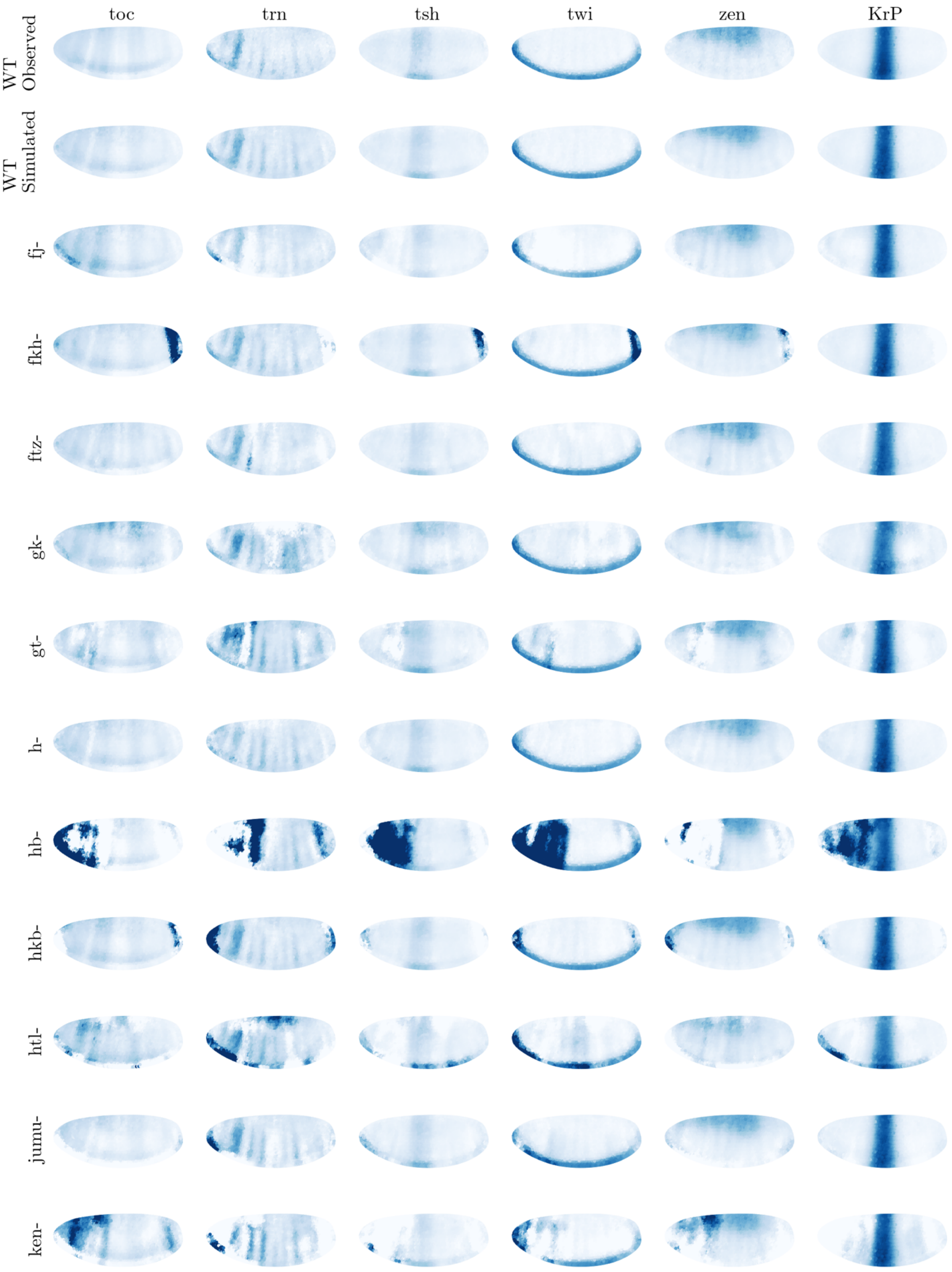
toc
trn
tsh
twi
zen
KrP
WT Observed
WT Simulated
fj-
fkh-
ftz-
gk-
gt-
h-
hb-
hkb-
htl-
jumu-
ken-

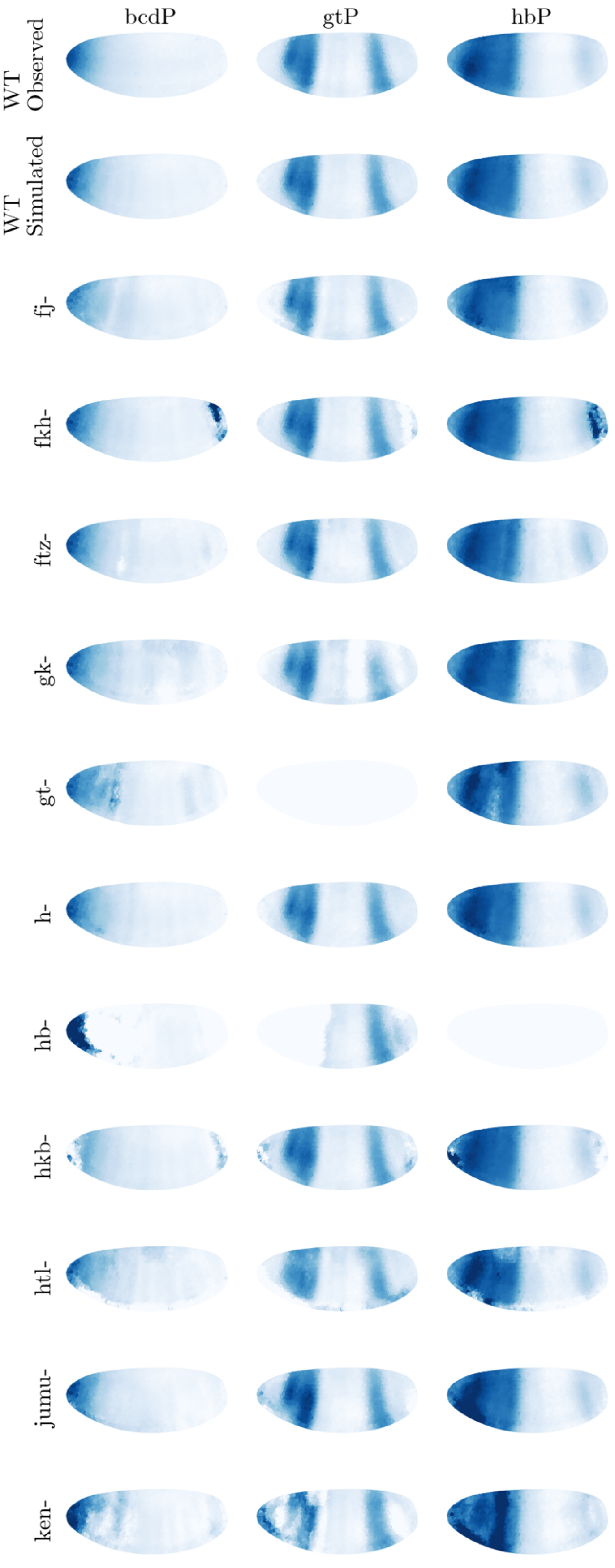
bcdP
gtP
hbP
WT Observed
WT Simulated
fj-
fkh-
ftz-
gk-
gt-
h-
hb-
hkb-
htl-
jumu-
ken-

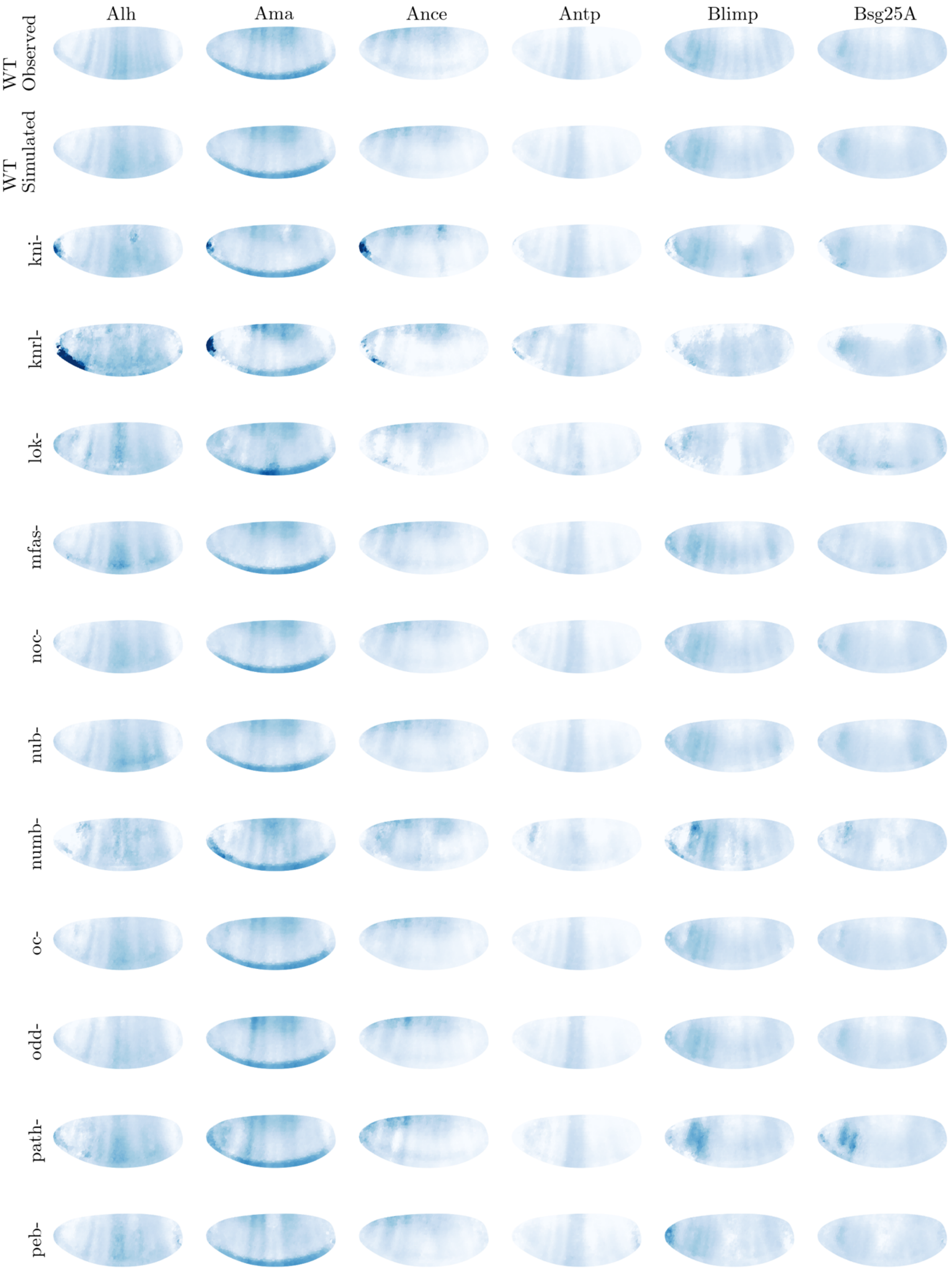
Alh
Ama
Ance
Antp
Blimp
Bsg25A
WT Observed
WT Simulated
kni-
knrl-
lok-
mfas-
noc-
nub-
numb-
oc-
odd-
path-
peb-

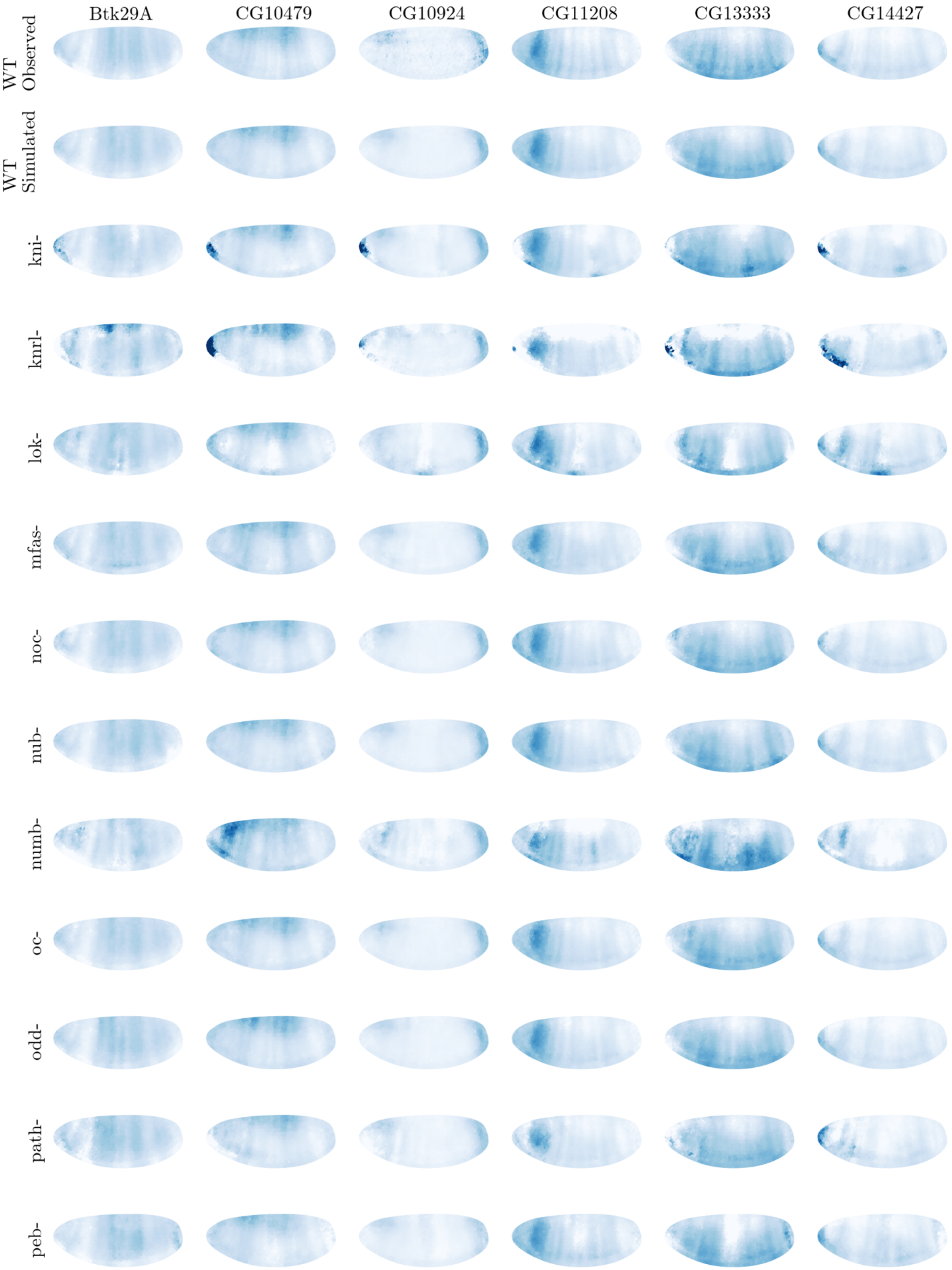
Btk29A
CG10479
CG10924
CG11208
CG13333
CG14427
WT Observed
WT Simulated
kni-
knrl-
lok-
mfas-
noc-
nub-
numb-
oc-
odd-
path-
peb-

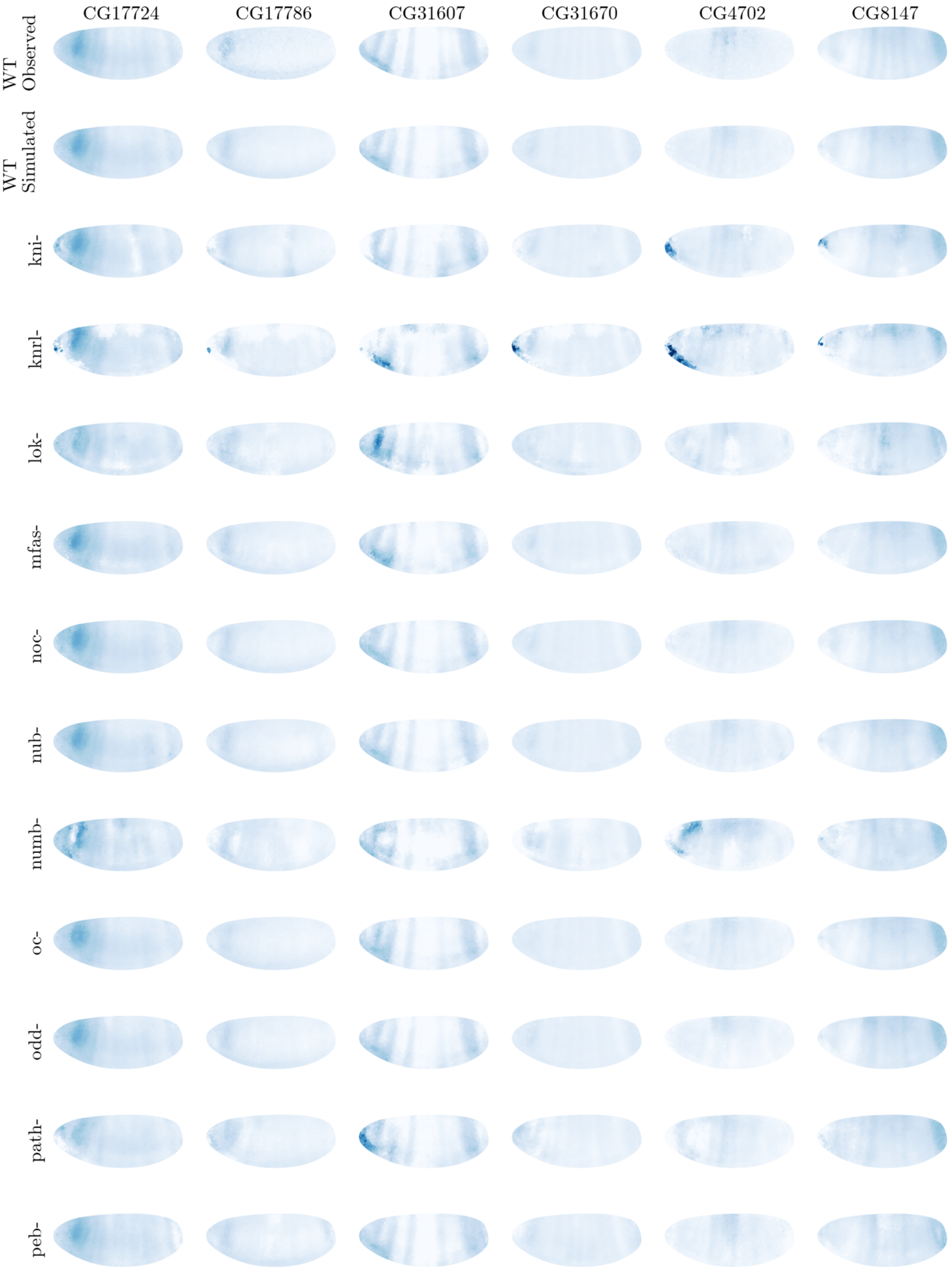
CG17724
CG17786
CG31607
CG31670
CG4702
CG8147
WT Observed
WT Simulated
kni-
knrl-
lok-
mfas-
noc-
nub-
numb-
oc-
odd-
path-
peb-

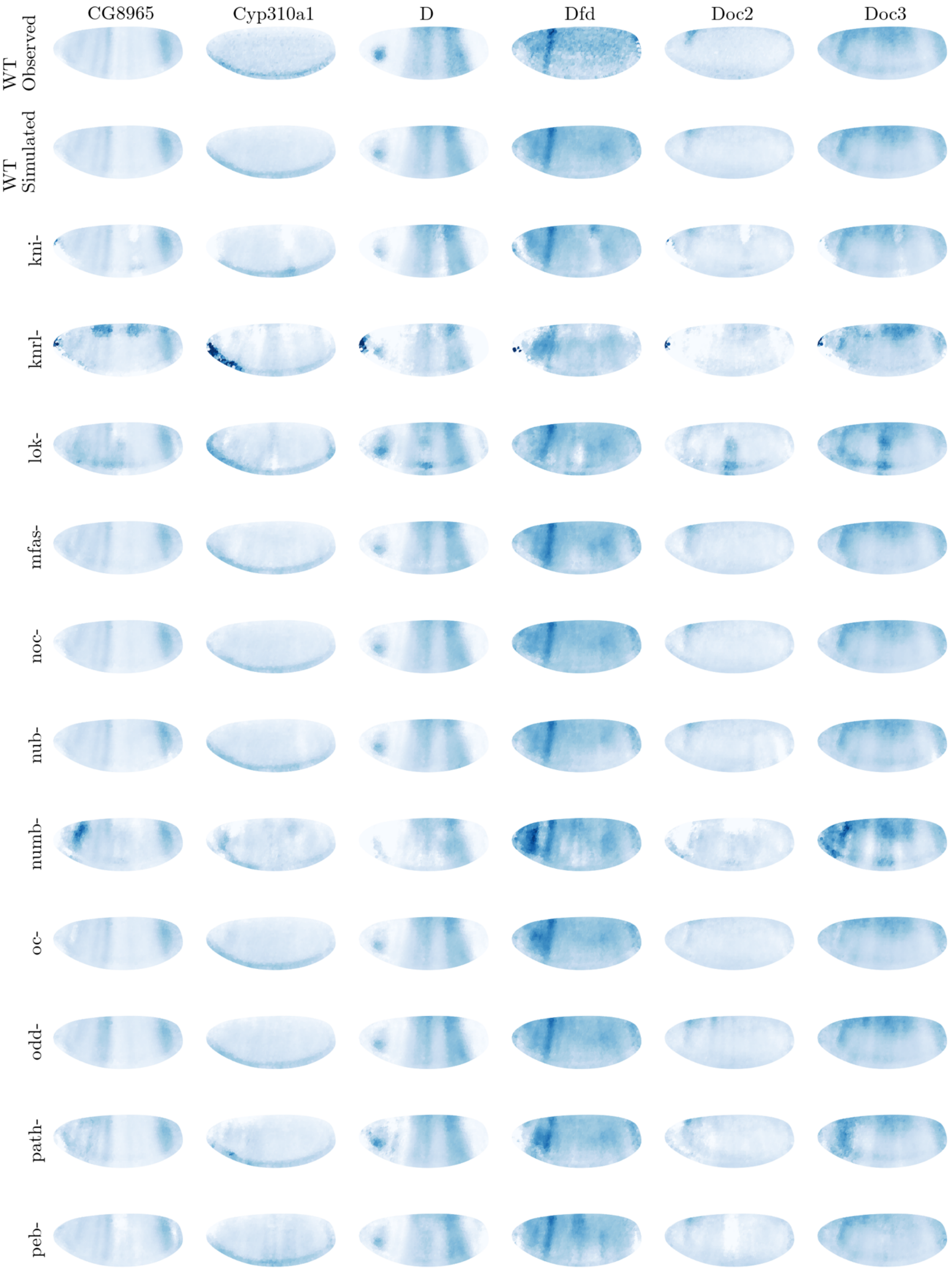
CG8965
Cyp310a1
D
Dfd
Doc2
Doc3
WT Observed
WT Simulated
kni-
knrl-
lok-
mfas-
noc-
nub-
numb-
oc-
odd-
path-
peb-

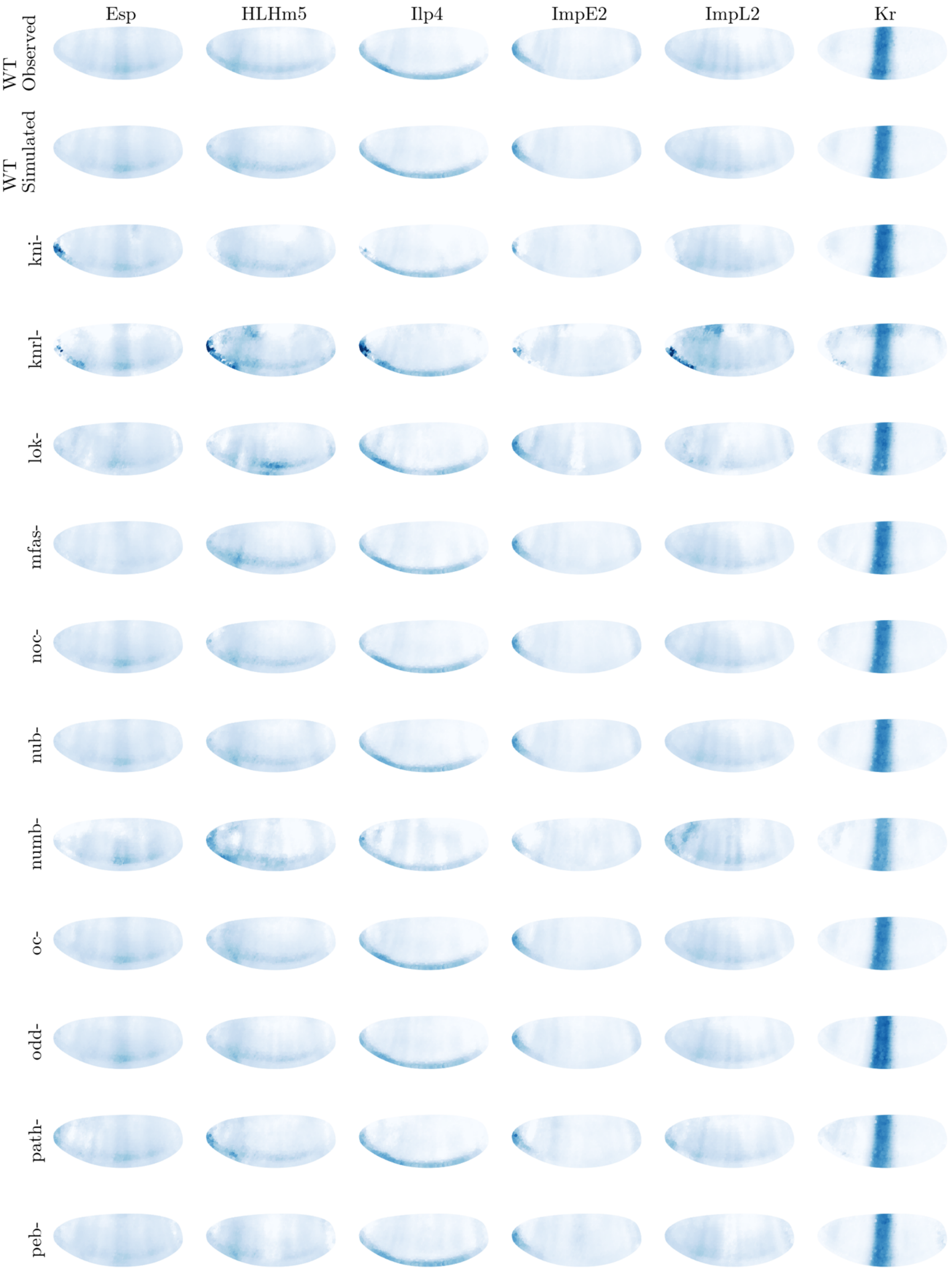
Esp
HLHm5
Ilp4
ImpE2
ImpL2
Kr
WT Observed
WT Simulated
kni-
knrl-
lok-
mfas-
noc-
nub-
numb-
oc-
odd-
path-
peb-

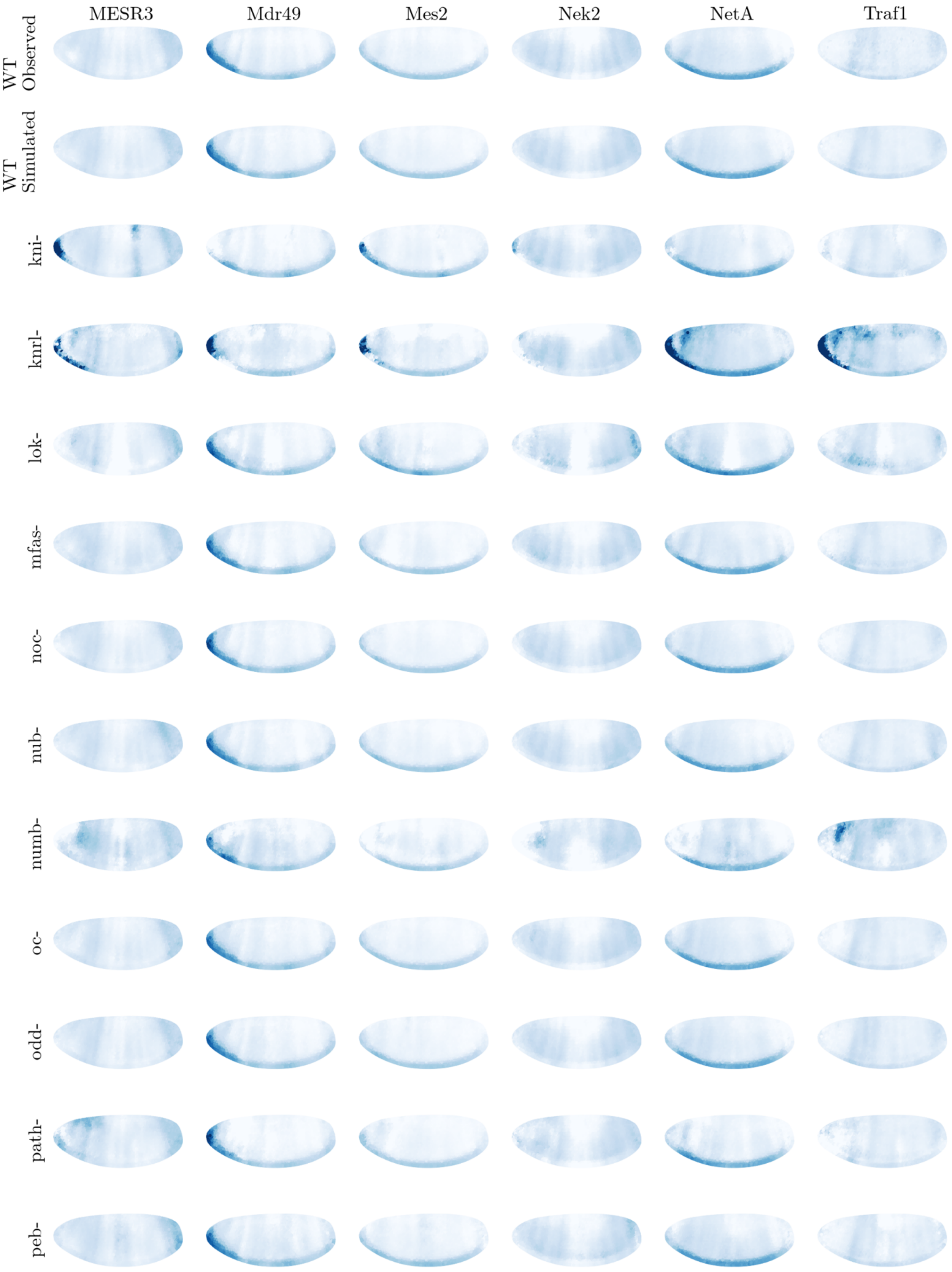
MESR3
Mdr49
Mes2
Nek2
NetA
Traf1
WT Observed
WT Simulated
kni-
knrl-
lok-
mfas-
noc-
nub-
numb-
oc-
odd-
path-
peb-

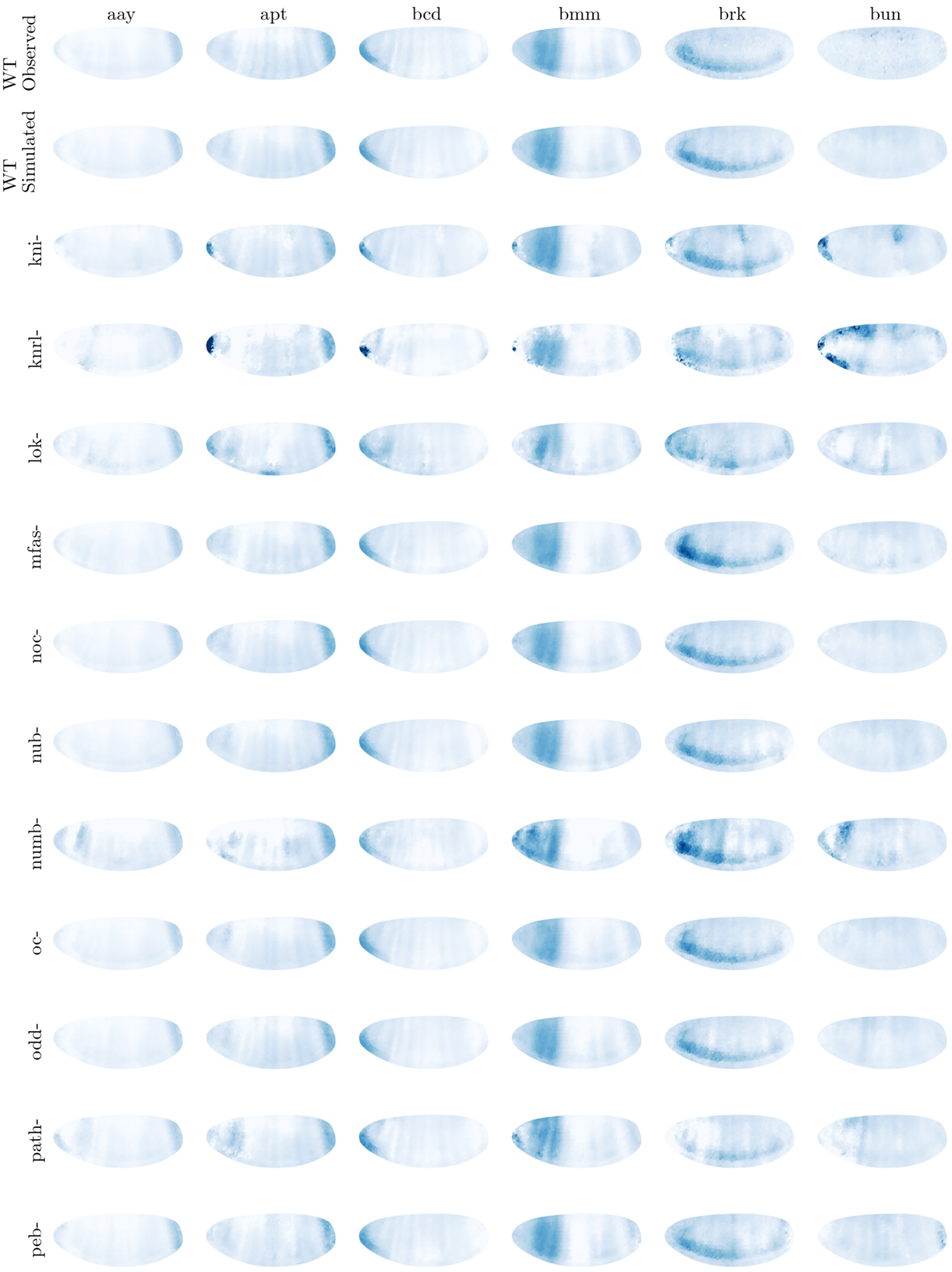
aay
apt
bcd
bmm
brk
bun
WT Observed
WT Simulated
kni-
knrl-
lok-
mfas-
noc-
nub-
numb-
oc-
odd-
path-
peb-

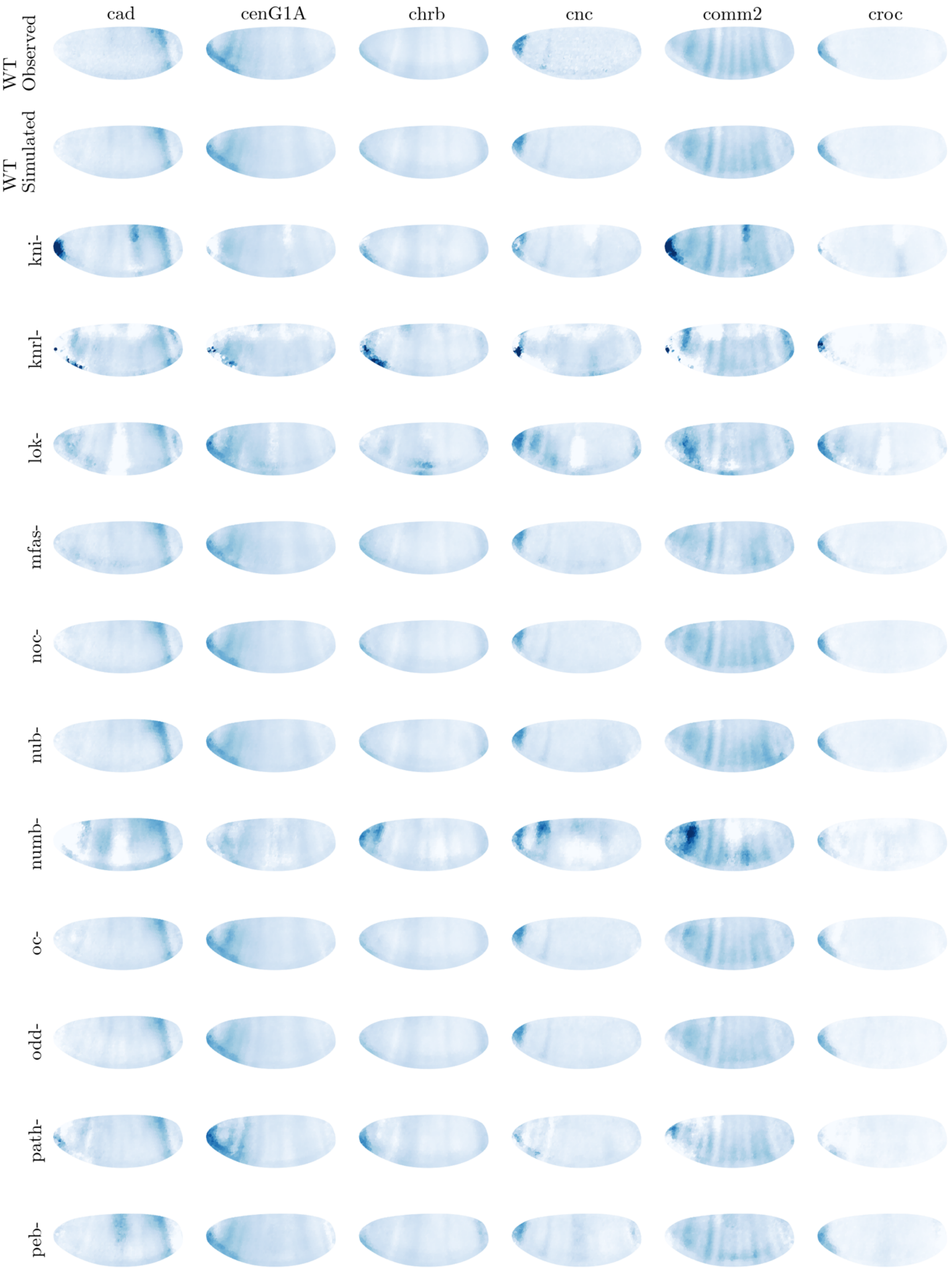

cad
cenG1A
chrb
cnc
comm2
croc
WT Observed
WT Simulated
kni-
knrl-
lok-
mfas-
noc-
nub-
numb-
oc-
odd-
path-
peb-

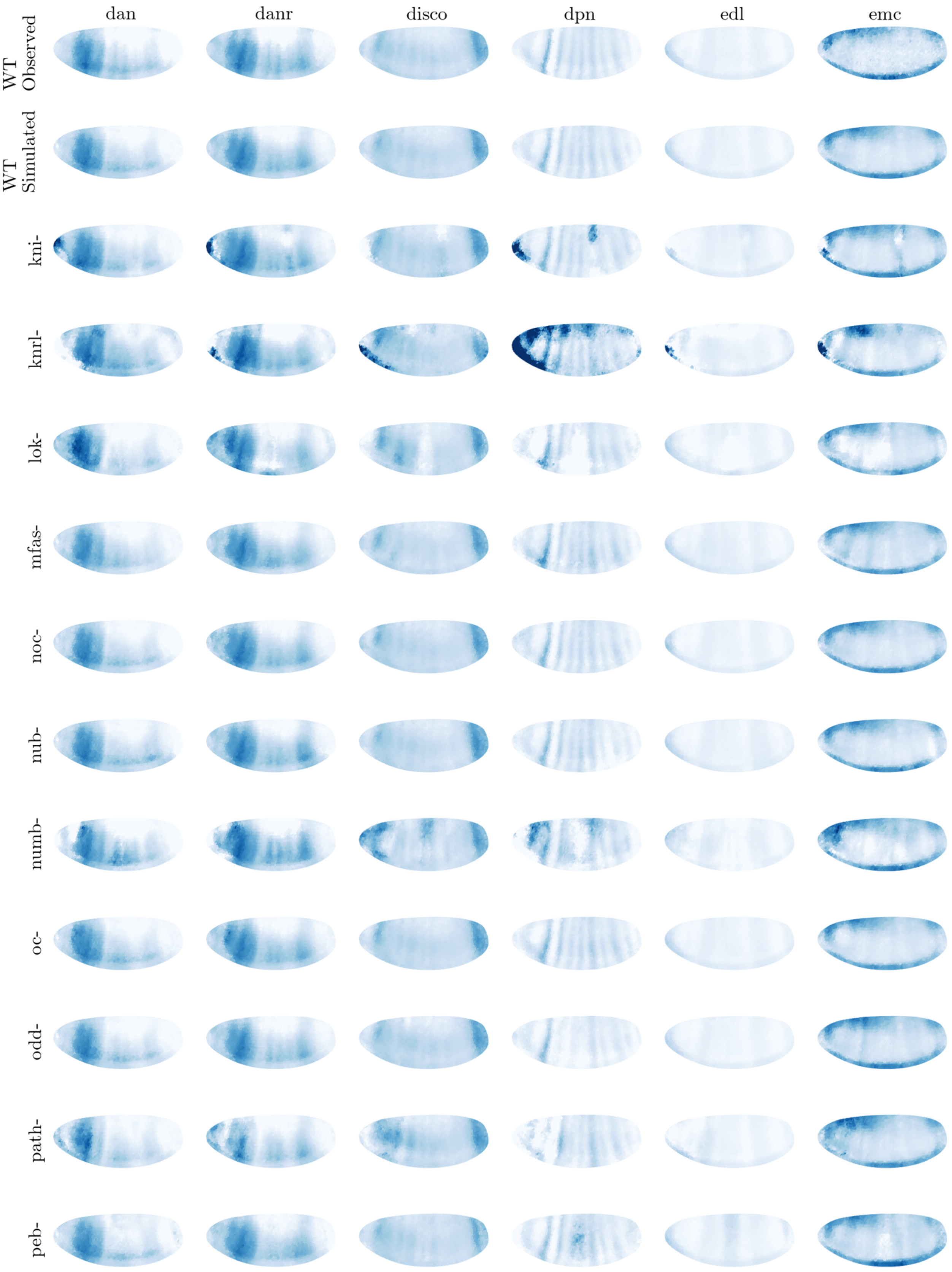
dan
danr
disco
dpn
edl
emc
WT Observed
WT Simulated
kni-
knrl-
lok-
mfas-
noc-
nub-
numb-
oc-
odd-
path-
peb-

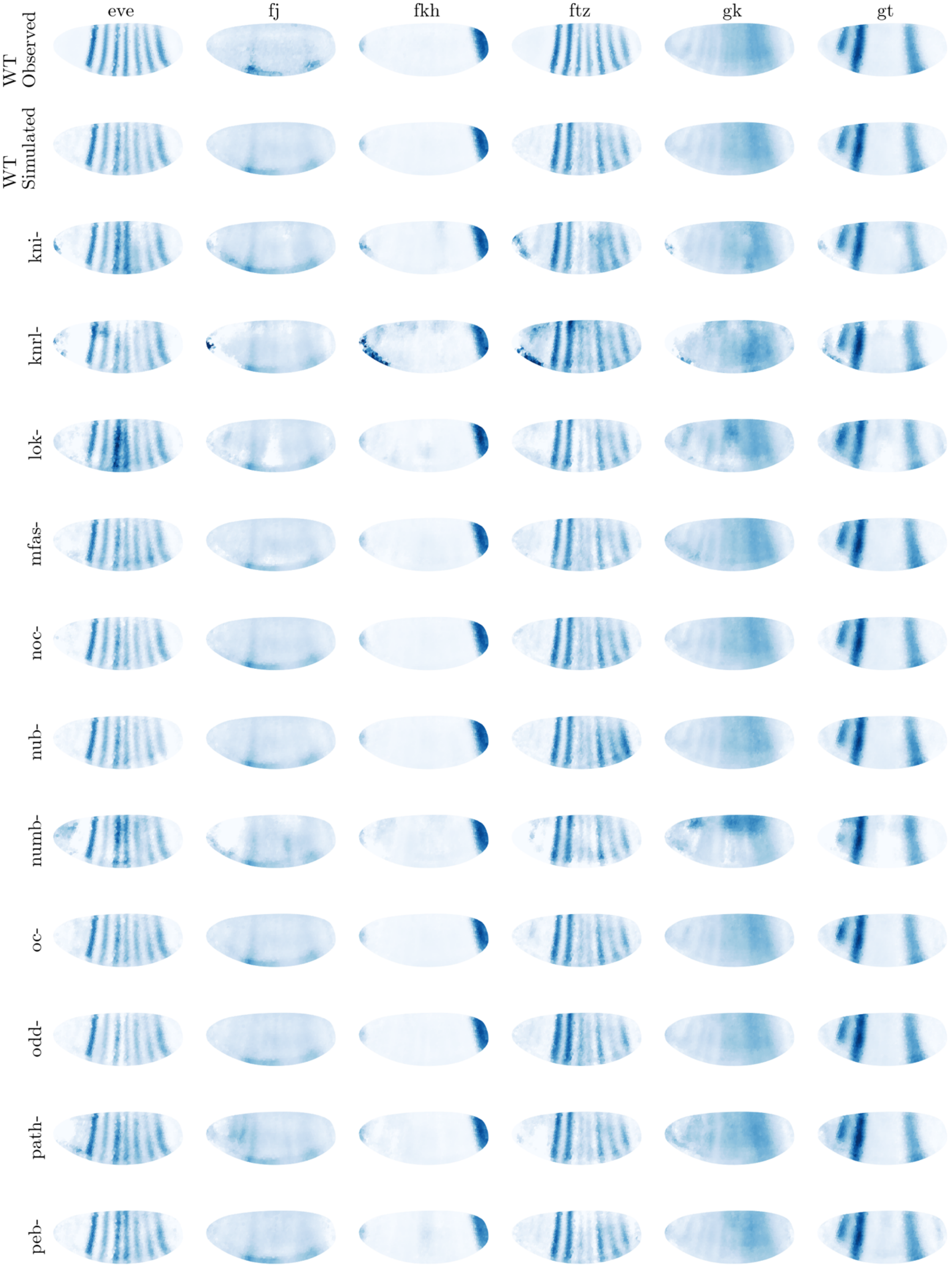

eve
fj
fkh
ftz
gk
gt
WT Observed
WT Simulated
kni-
knrl-
lok-
mfas-
noc-
nub-
numb-
oc-
odd-
path-
peb-

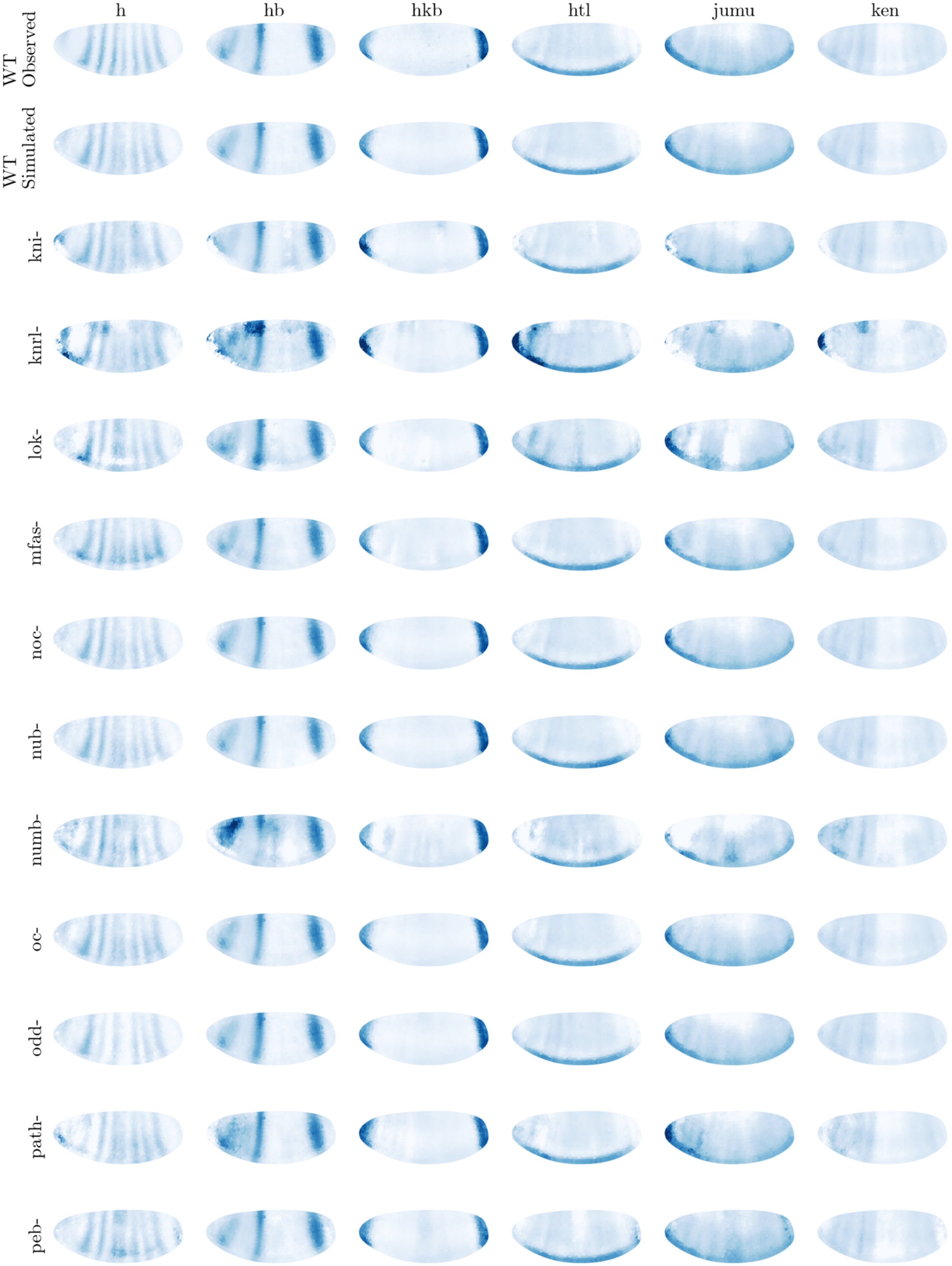
h
hb
hkb
htl
jumu
ken
WT Observed
WT Simulated
kni-
knrl-
lok-
mfas-
noc-
nub-
numb-
oc-
odd-
path-
peb-

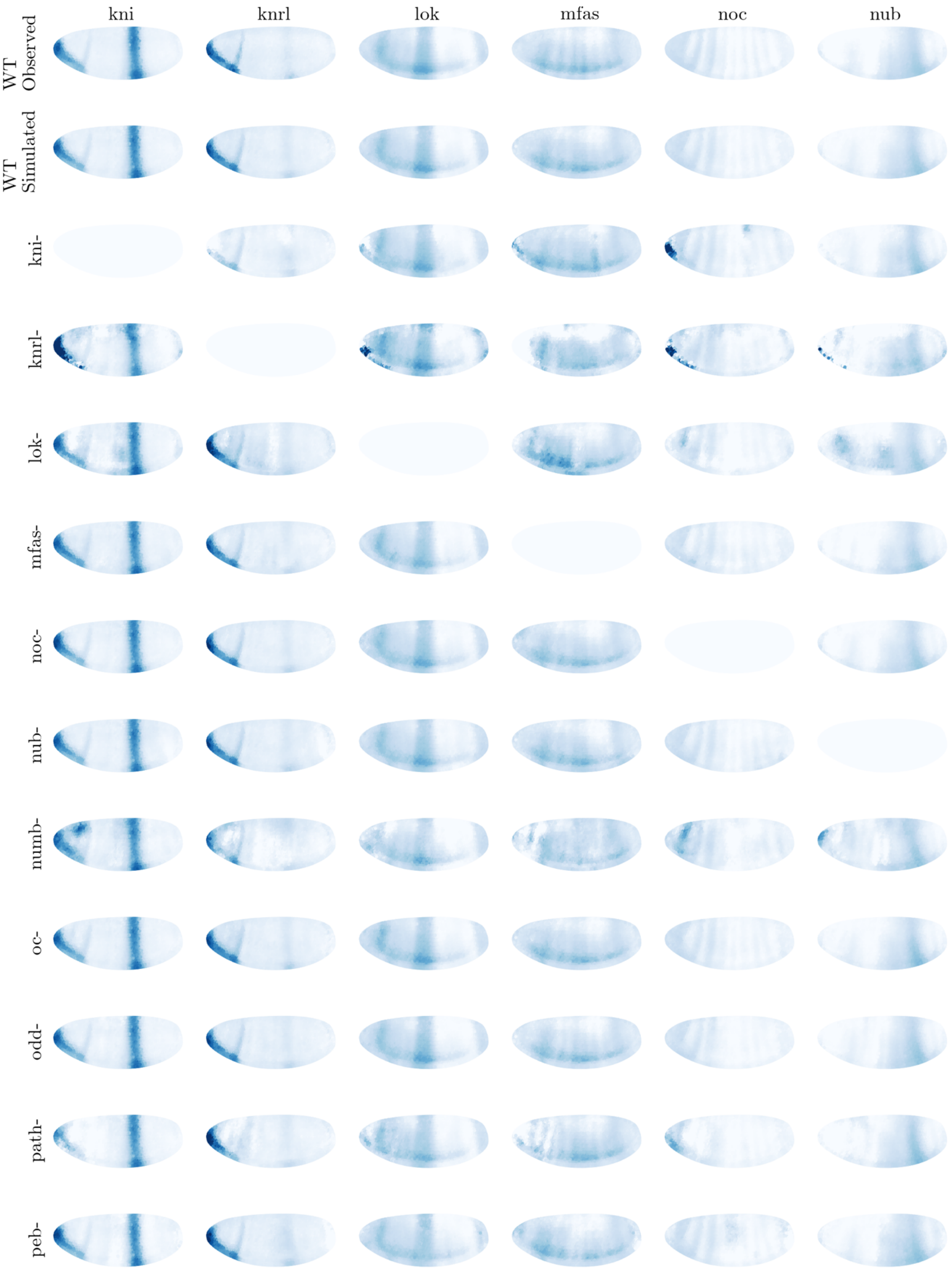
kni
knrl
lok
mfas
noc
nub
WT Observed
WT Simulated
kni-
knrl-
lok-
mfas-
noc-
nub-
numb-
oc-
odd-
path-
peb-

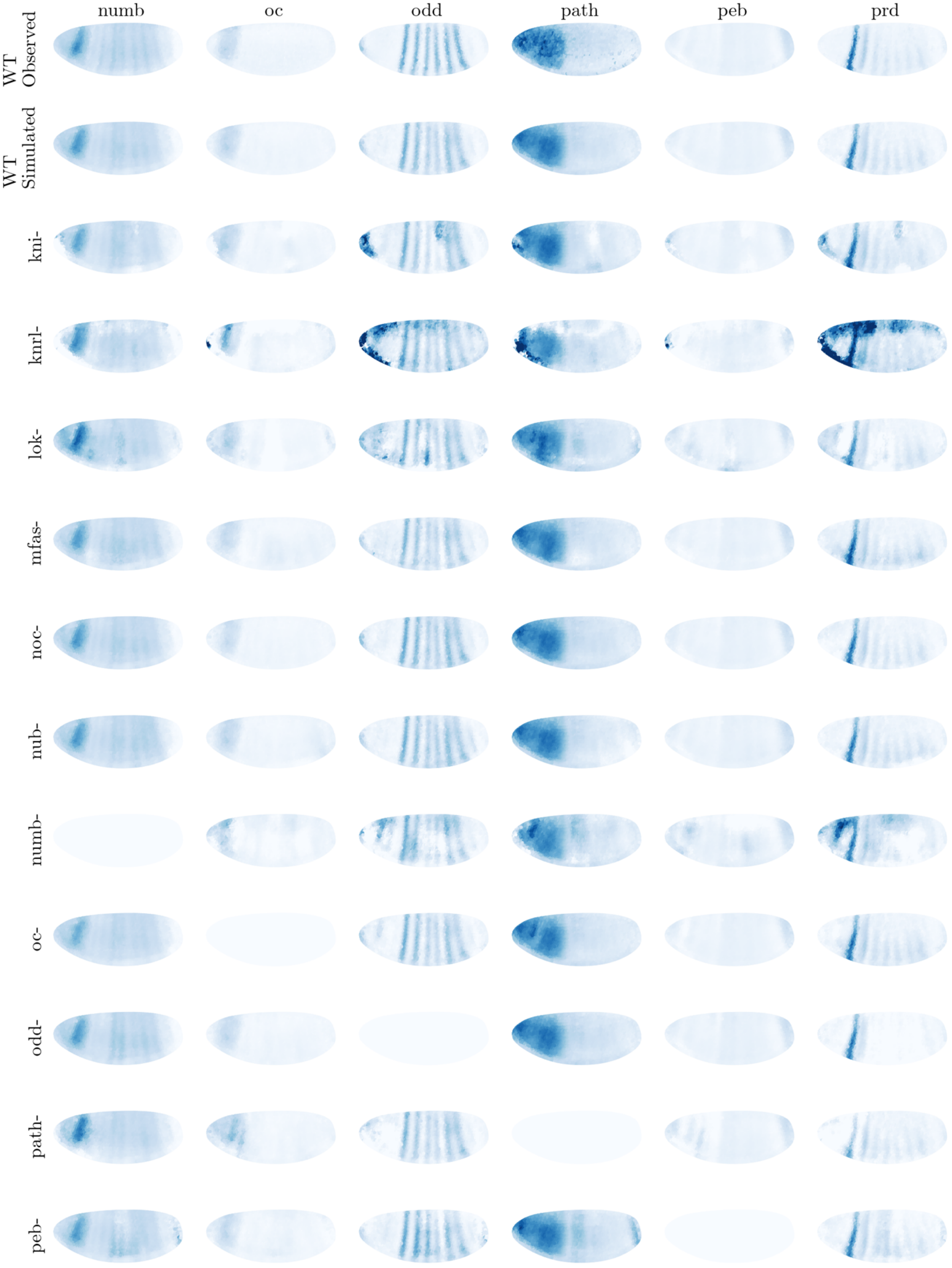
numb
oc
odd
path
peb
prd
WT Observed
WT Simulated
kni-
knrl-
lok-
mfas-
noc-
nub-
numb-
oc-
odd-
path-
peb-

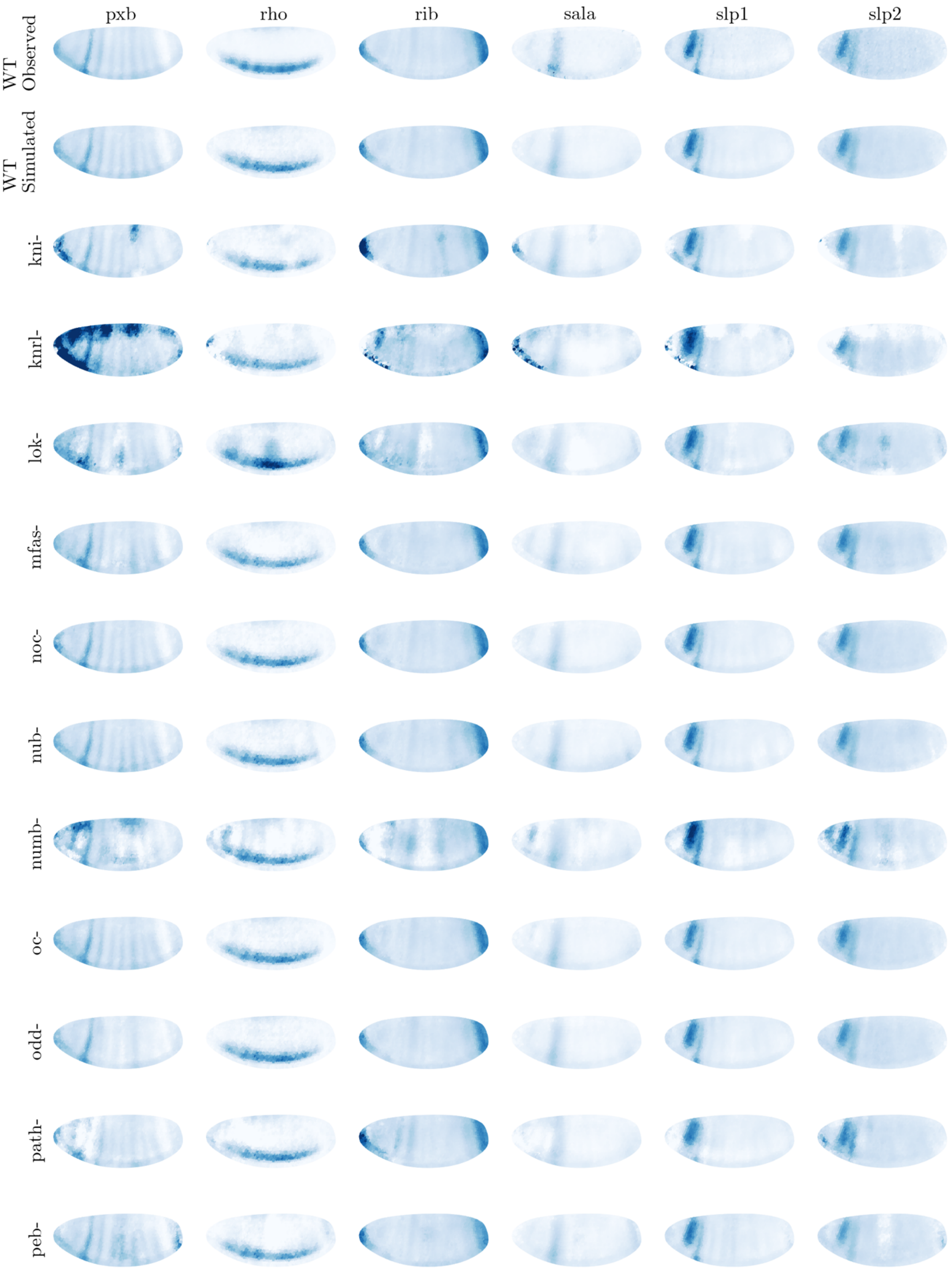
pxb
rho
rib
sala
slp1
slp2
WT Observed
WT Simulated
kni-
knrl-
lok-
mfas-
noc-
nub-
numb-
oc-
odd-
path-
peb-

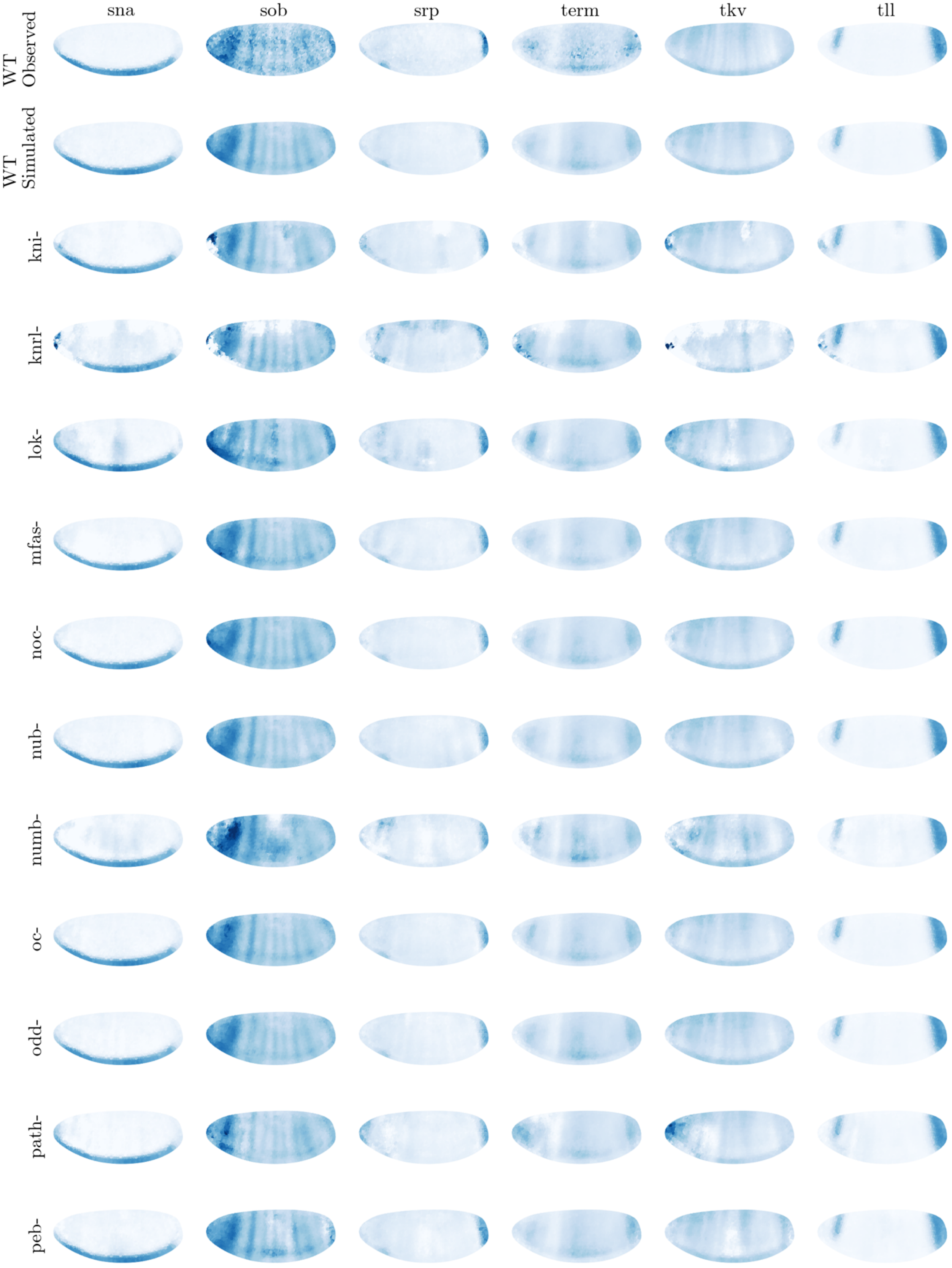
sna
sob
srp
term
tkv
tll
WT Observed
WT Simulated
kni-
knrl-
lok-
mfas-
noc-
nub-
numb-
oc-
odd-
path-
peb-

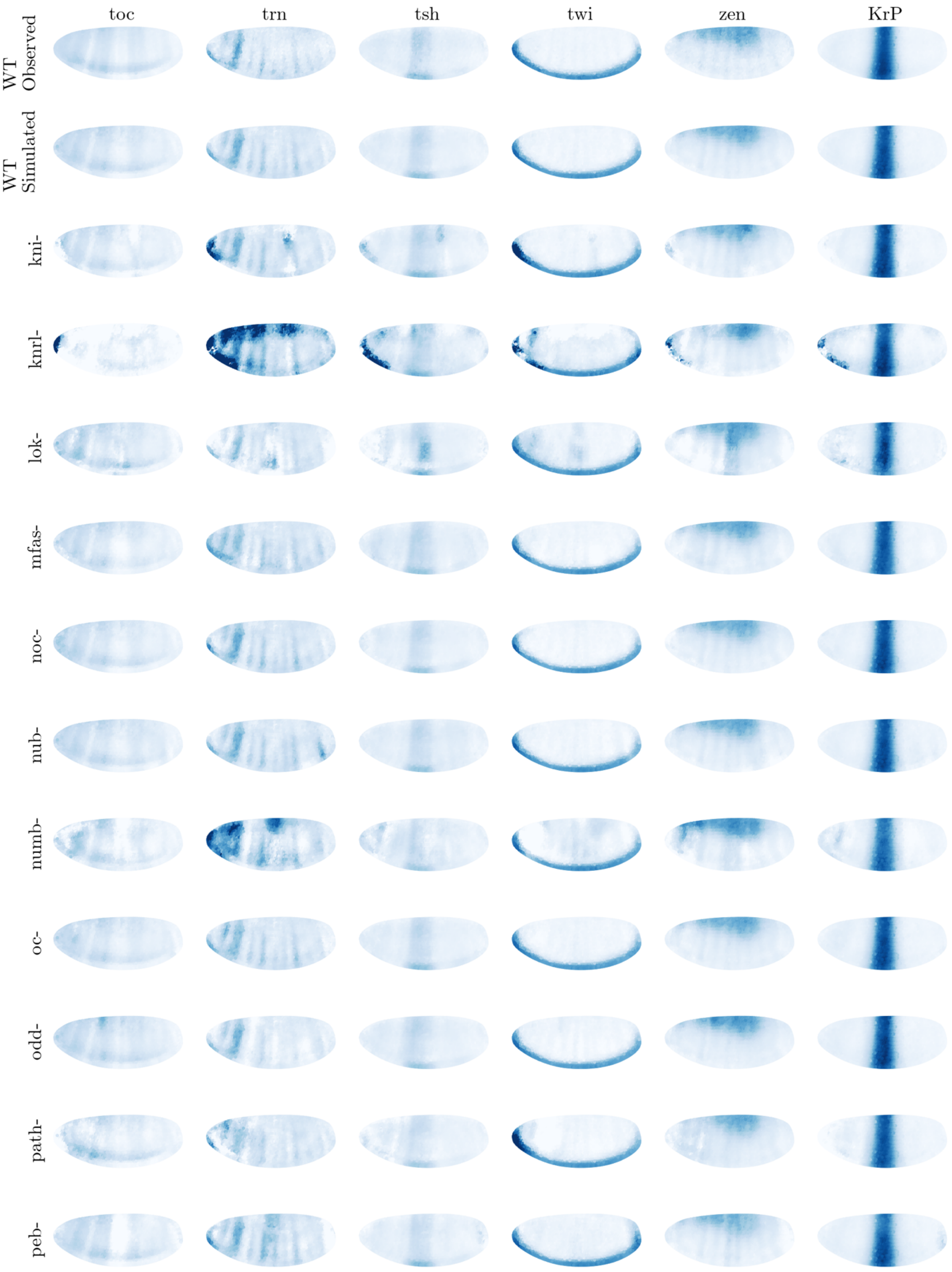
toc
trn
tsh
twi
zen
KrP
WT Observed
WT Simulated
kni-
knrl-
lok-
mfas-
noc-
nub-
numb-
oc-
odd-
path-
peb-

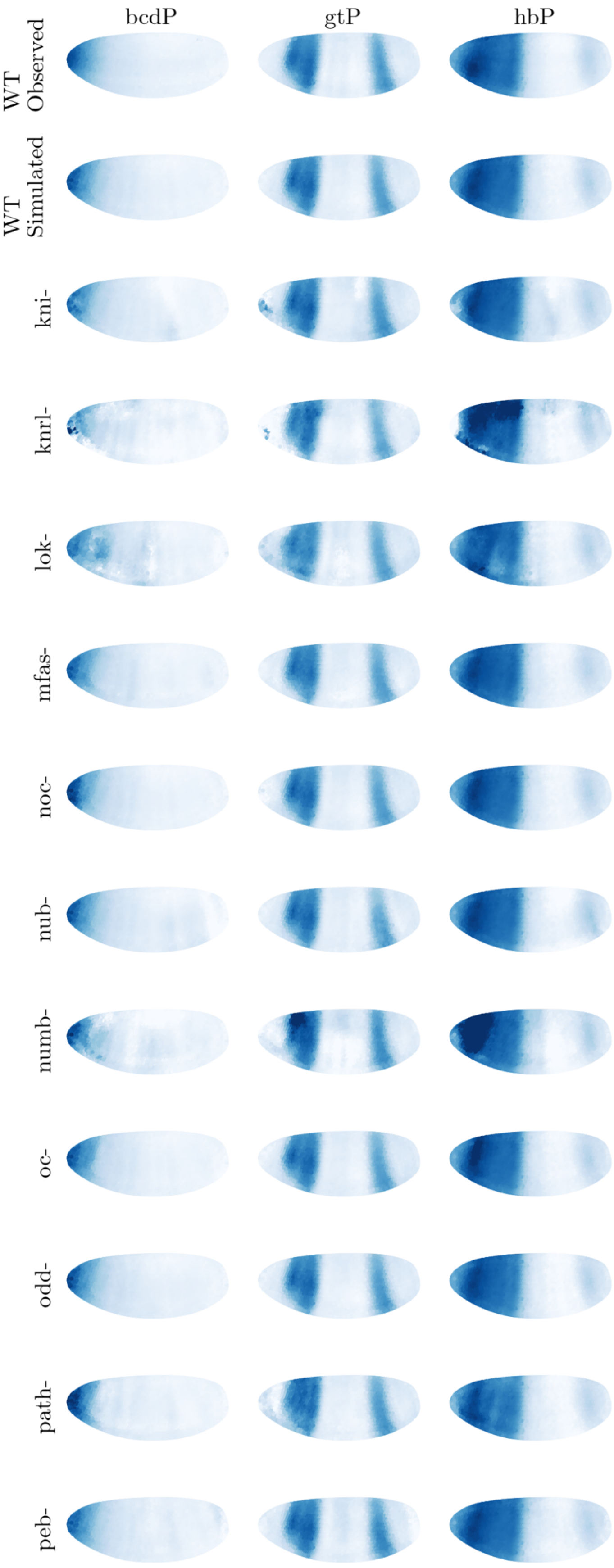
bcdP
gtP
hbP
WT Observed
WT Simulated
kni-
knrl-
lok-
mfas-
noc-
nub-
numb-
oc-
odd-
path-
peb-

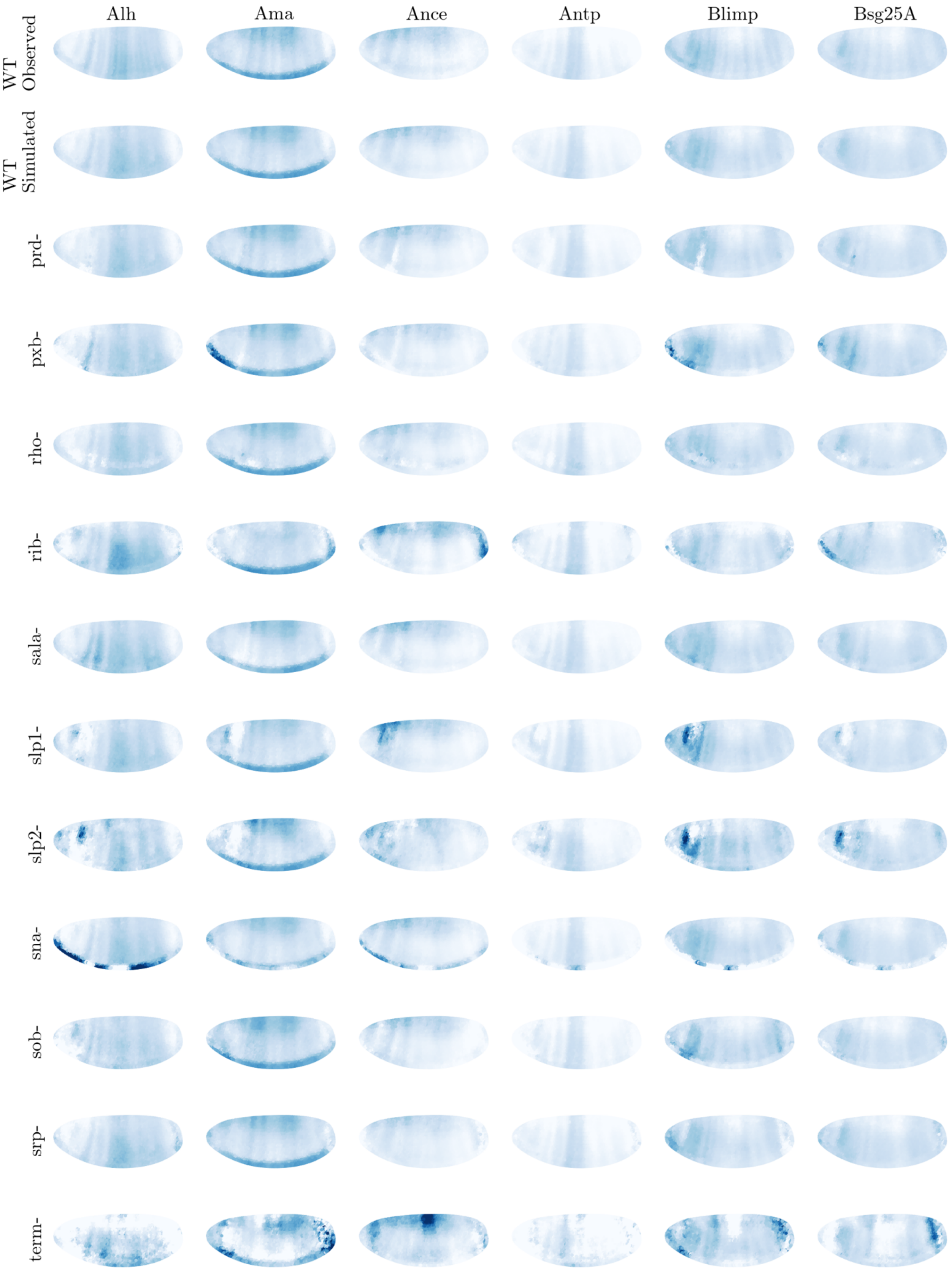
Alh
Ama
Ance
Antp
Blimp
Bsg25A
WT Observed
WT Simulated
prd-
pxb-
rho-
rib-
sala-
slp1-
slp2-
sna-
sob-
srp-
term-

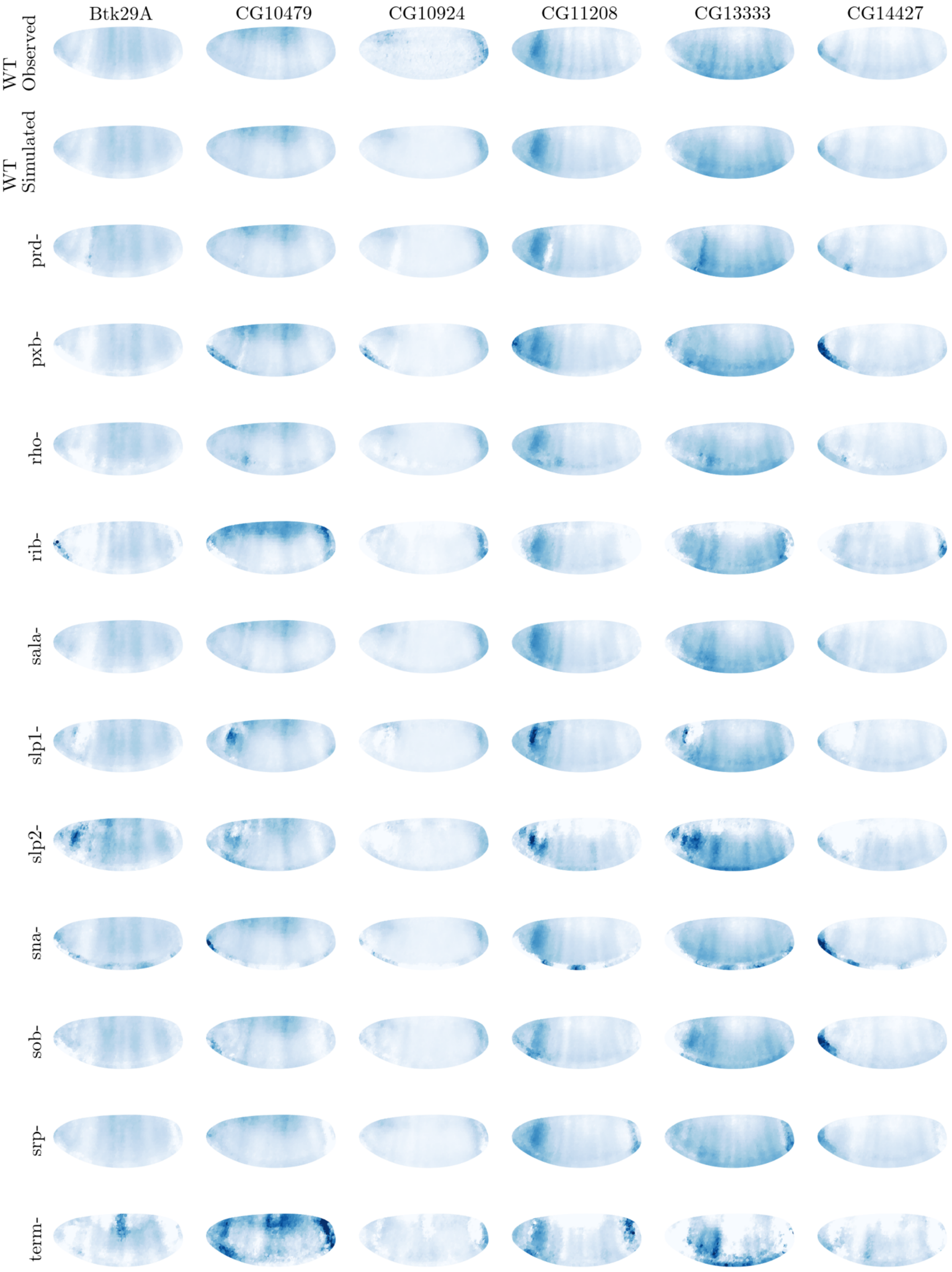
Btk29A
CG10479
CG10924
CG11208
CG13333
CG14427
WT Observed
WT Simulated
prd-
pxb-
rho-
rib-
sala-
slp1-
slp2-
sna-
sob-
srp-
term-

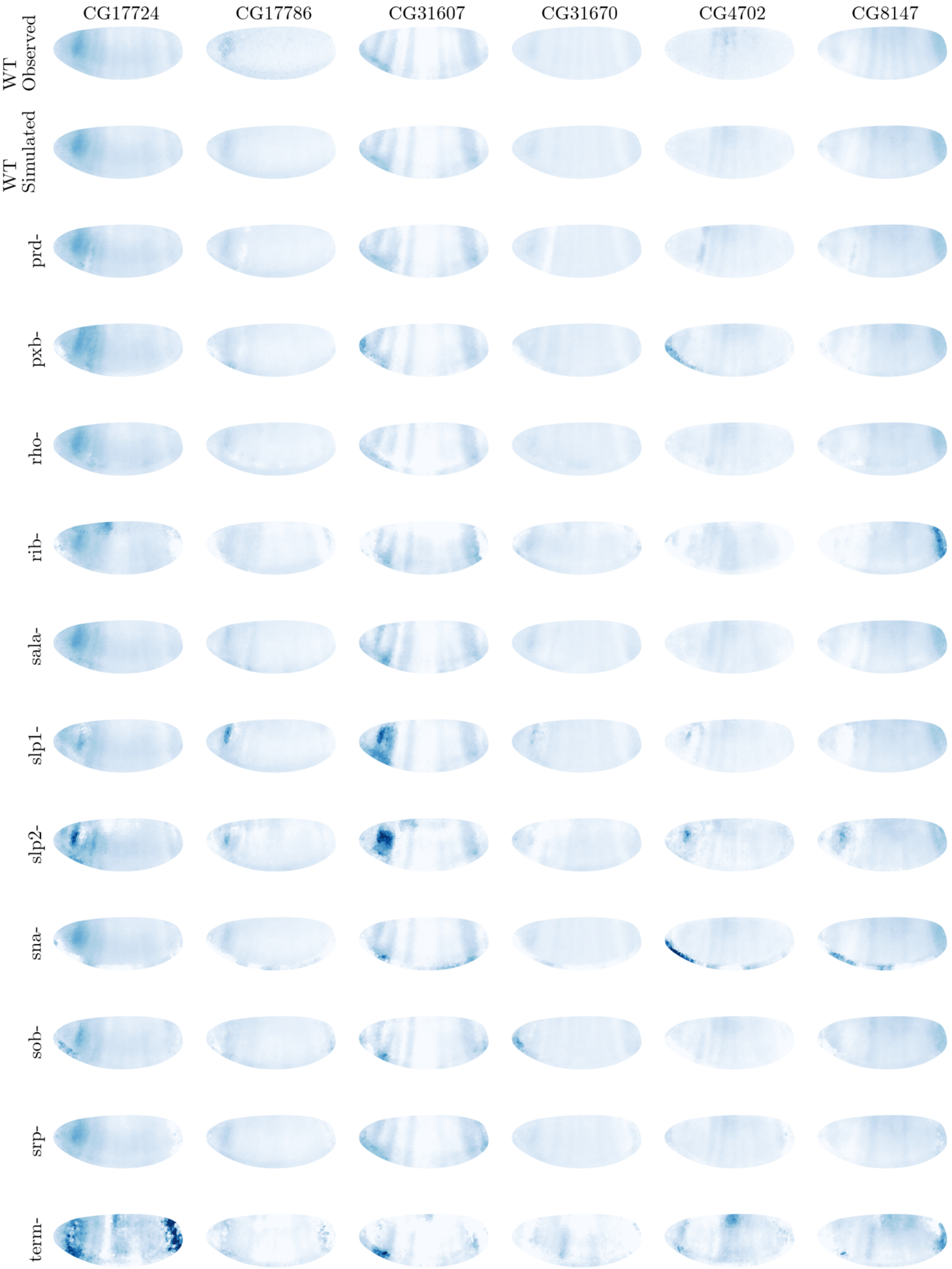
CG17724
CG17786
CG31607
CG31670
CG4702
CG8147
WT Observed
WT Simulated
prd-
pxb-
rho-
rib-
sala-
slp1-
slp2-
sna-
sob-
srp-
term-

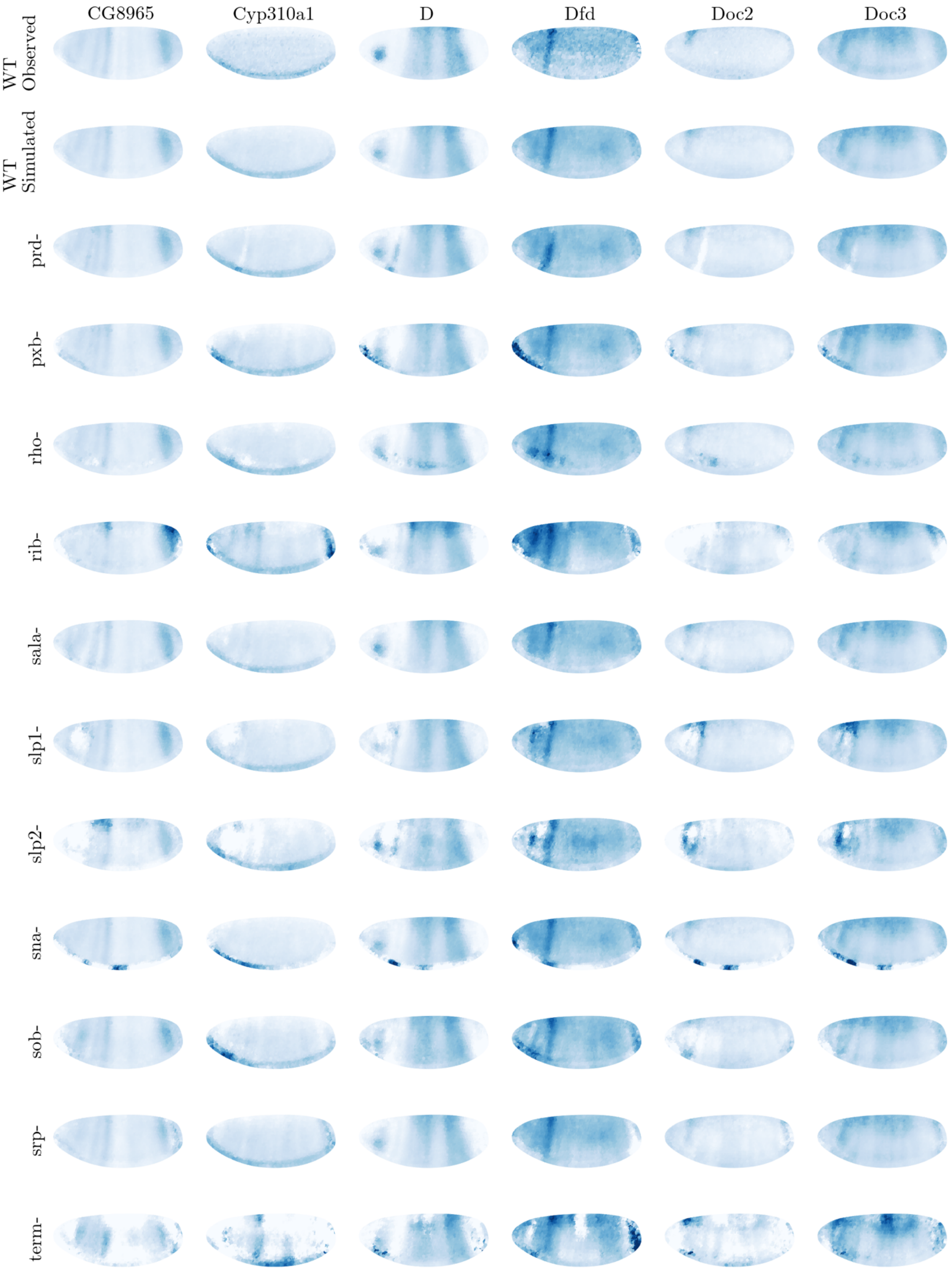
CG8965
Cyp310a1
D
Dfd
Doc2
Doc3
WT Observed
WT Simulated
prd-
pxb-
rho-
rib-
sala-
slp1-
slp2-
sna-
sob-
srp-
term-

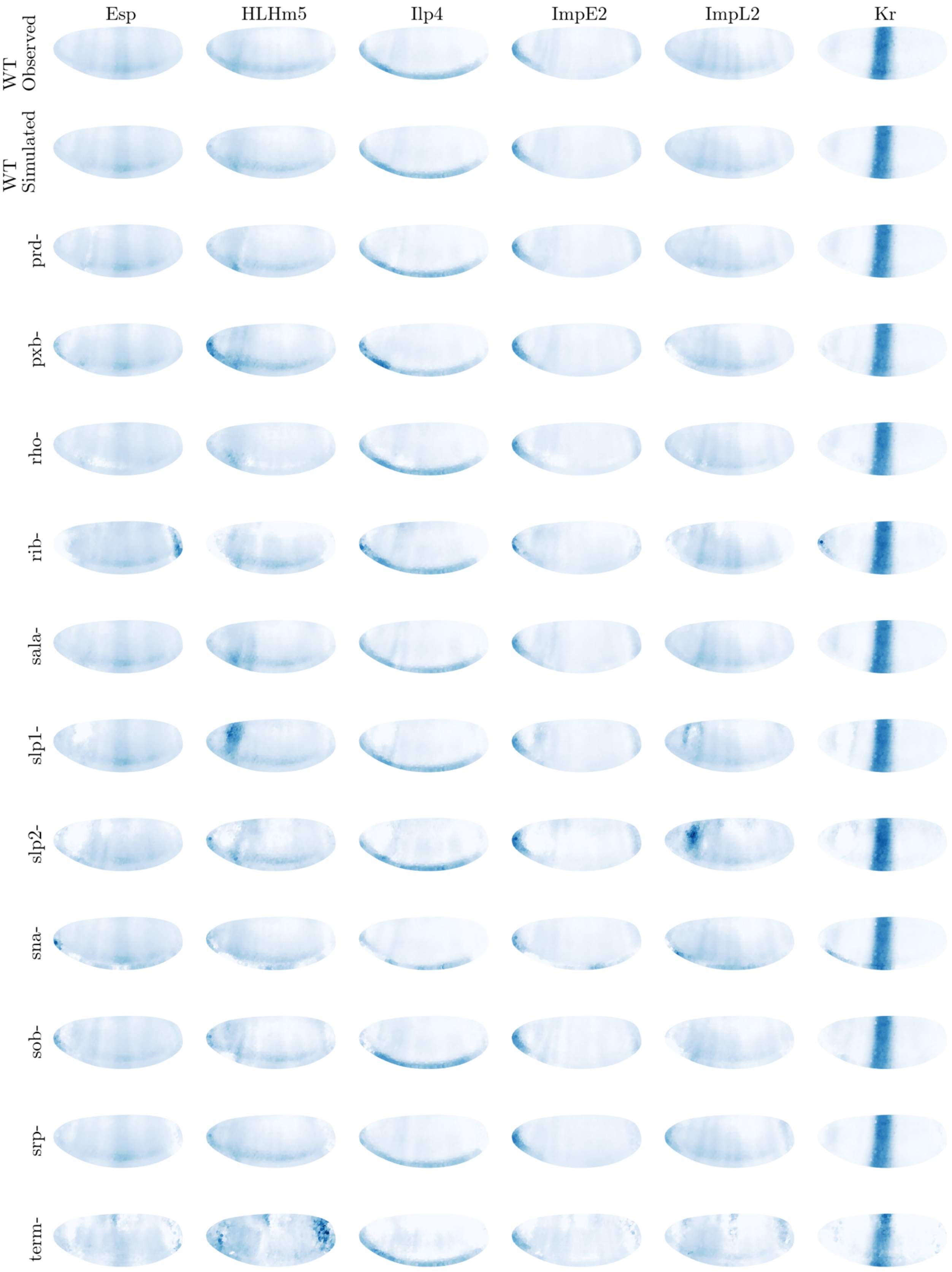
Esp
HLHm5
Ilp4
ImpE2
ImpL2
Kr
WT Observed
WT Simulated
prd-
pxb-
rho-
rib-
sala-
slp1-
slp2-
sna-
sob-
srp-
term-

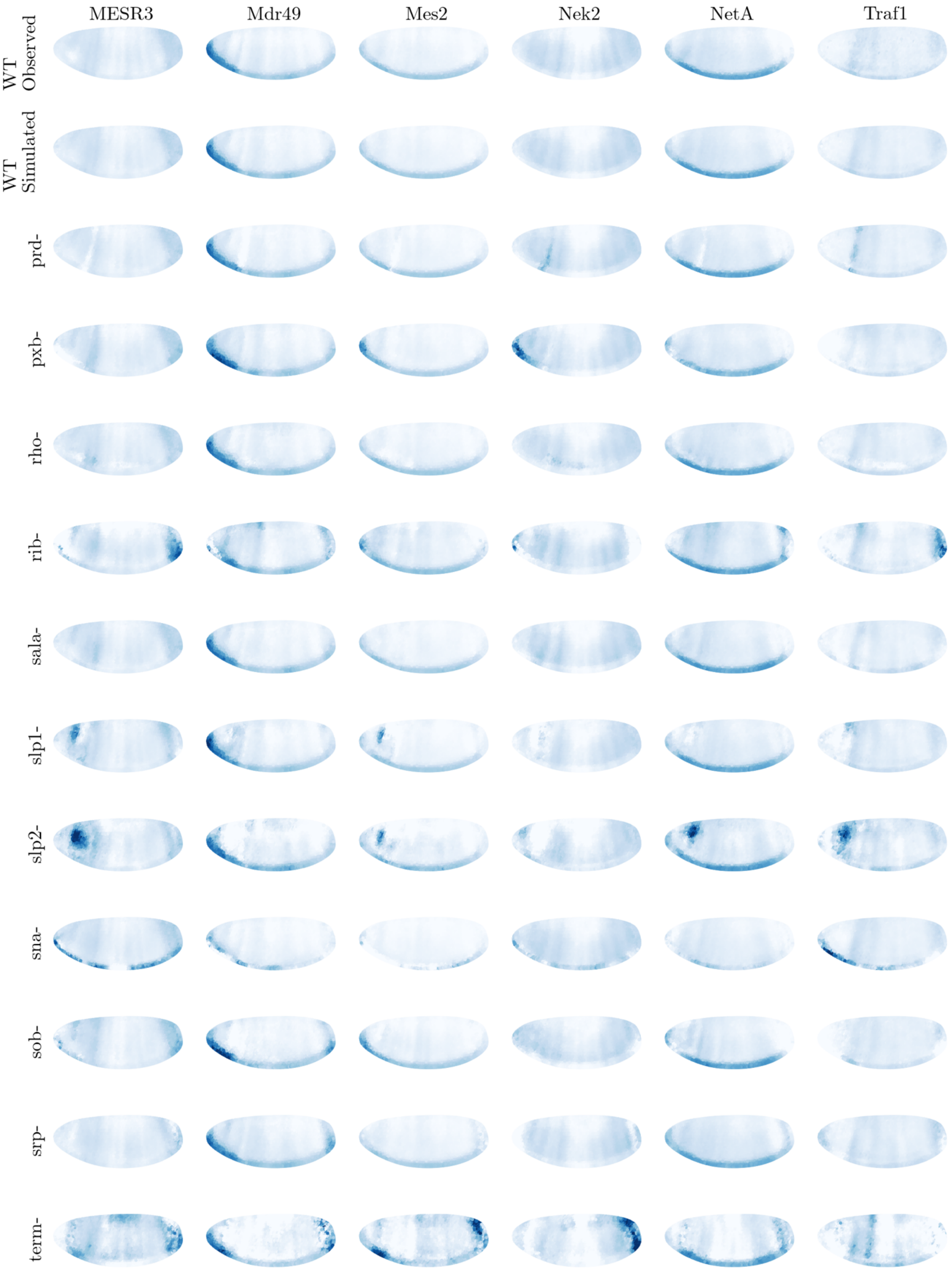

MESR3
Mdr49
Mes2
Nek2
NetA
Traf1
WT Observed
WT Simulated
prd-
pxb-
rho-
rib-
sala-
slp1-
slp2-
sna-
sob-
srp-
term-

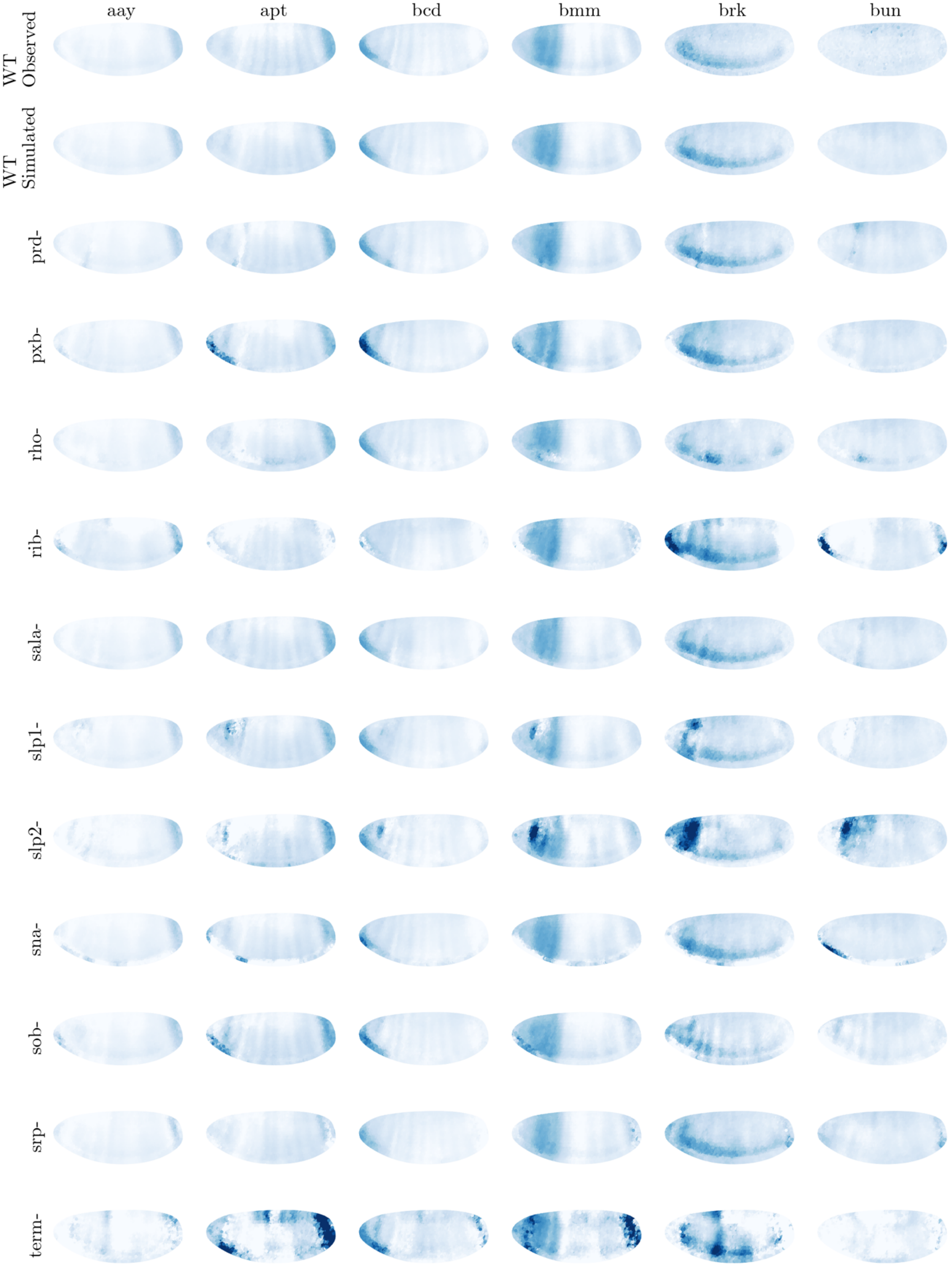

aay
apt
bcd
bmm
brk
bun
WT Observed
WT Simulated
prd-
pxb-
rho-
rib-
sala-
slp1-
slp2-
sna-
sob-
srp-
term-

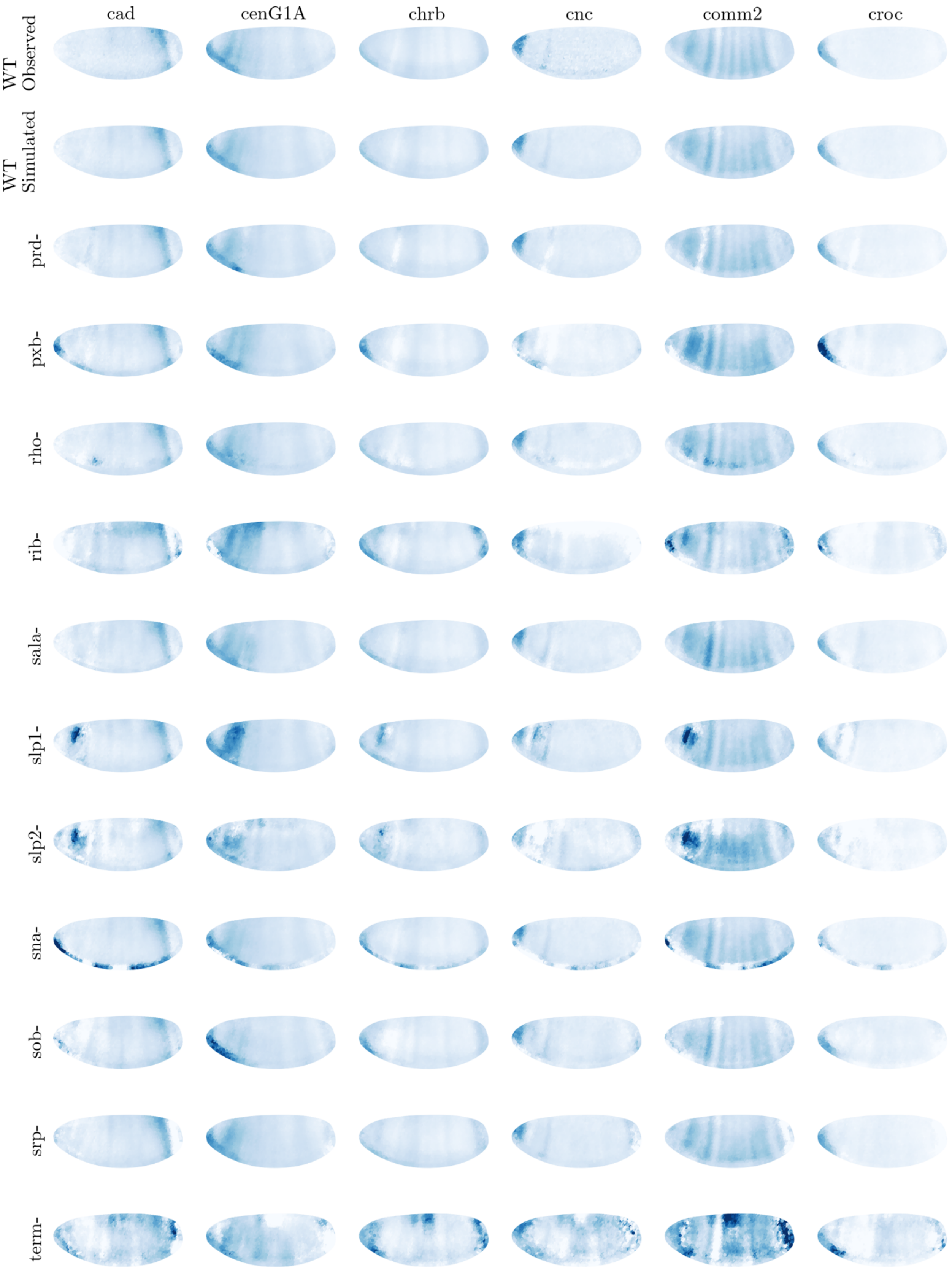
cad
cenG1A
chrb
cnc
comm2
croc
WT Observed
WT Simulated
prd-
pxb-
rho-
rib-
sala-
slp1-
slp2-
sna-
sob-
srp-
term-

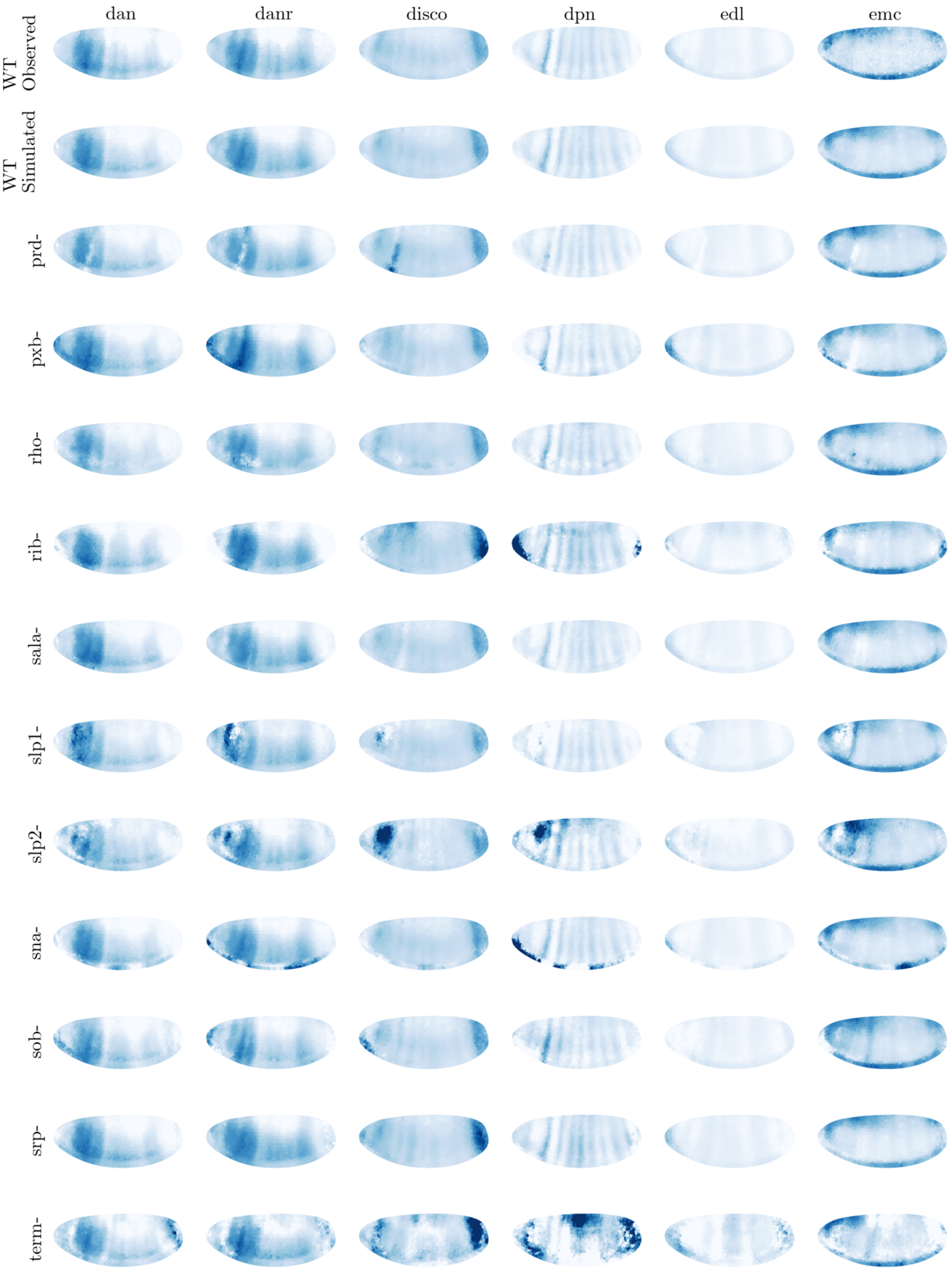

dan
danr
disco
dpn
edl
emc
WT Observed
WT Simulated
prd-
pxb-
rho-
rib-
sala-
slp1-
slp2-
sna-
sob-
srp-
term-

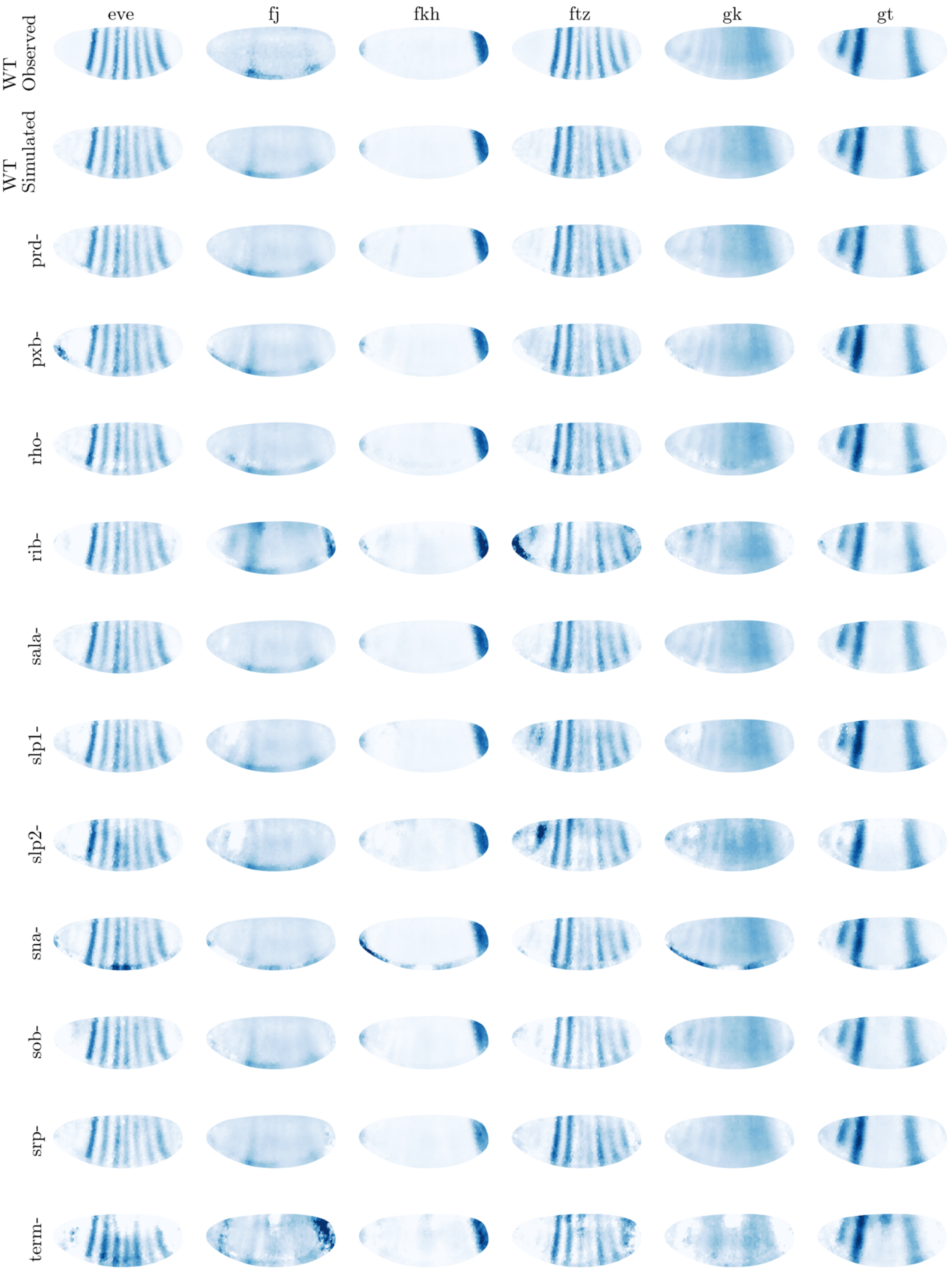
eve
fj
fkh
ftz
gk
gt
WT Observed
WT Simulated
prd-
pxb-
rho-
rib-
sala-
slp1-
slp2-
sna-
sob-
srp-
term-

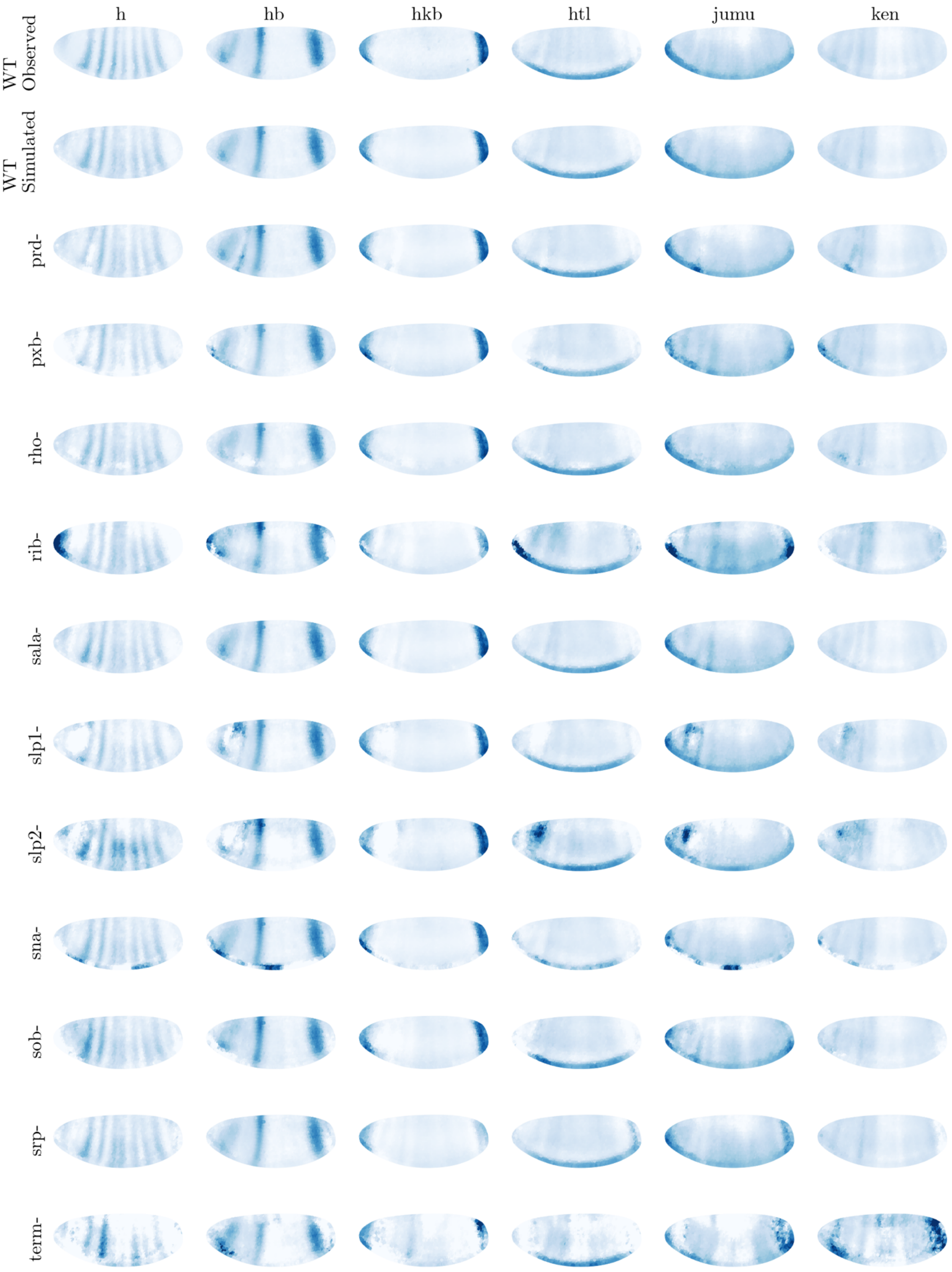
h
hb
hkb
htl
jumu
ken
WT Observed
WT Simulated
prd-
pxb-
rho-
rib-
sala-
slp1-
slp2-
sna-
sob-
srp-
term-

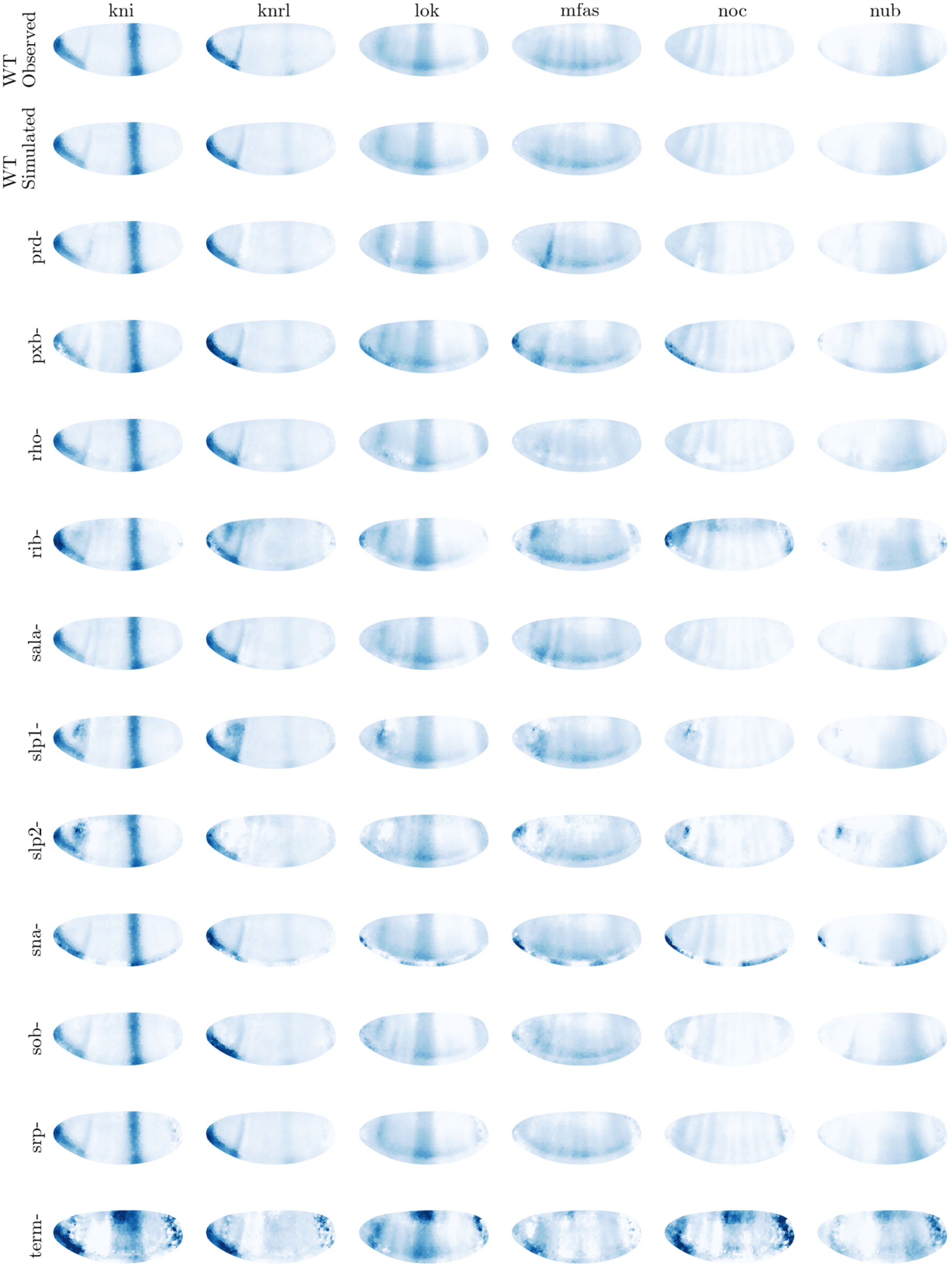
kni
knrl
lok
mfas
noc
nub
WT Observed
WT Simulated
prd-
pxb-
rho-
rib-
sala-
slp1-
slp2-
sna-
sob-
srp-
term-

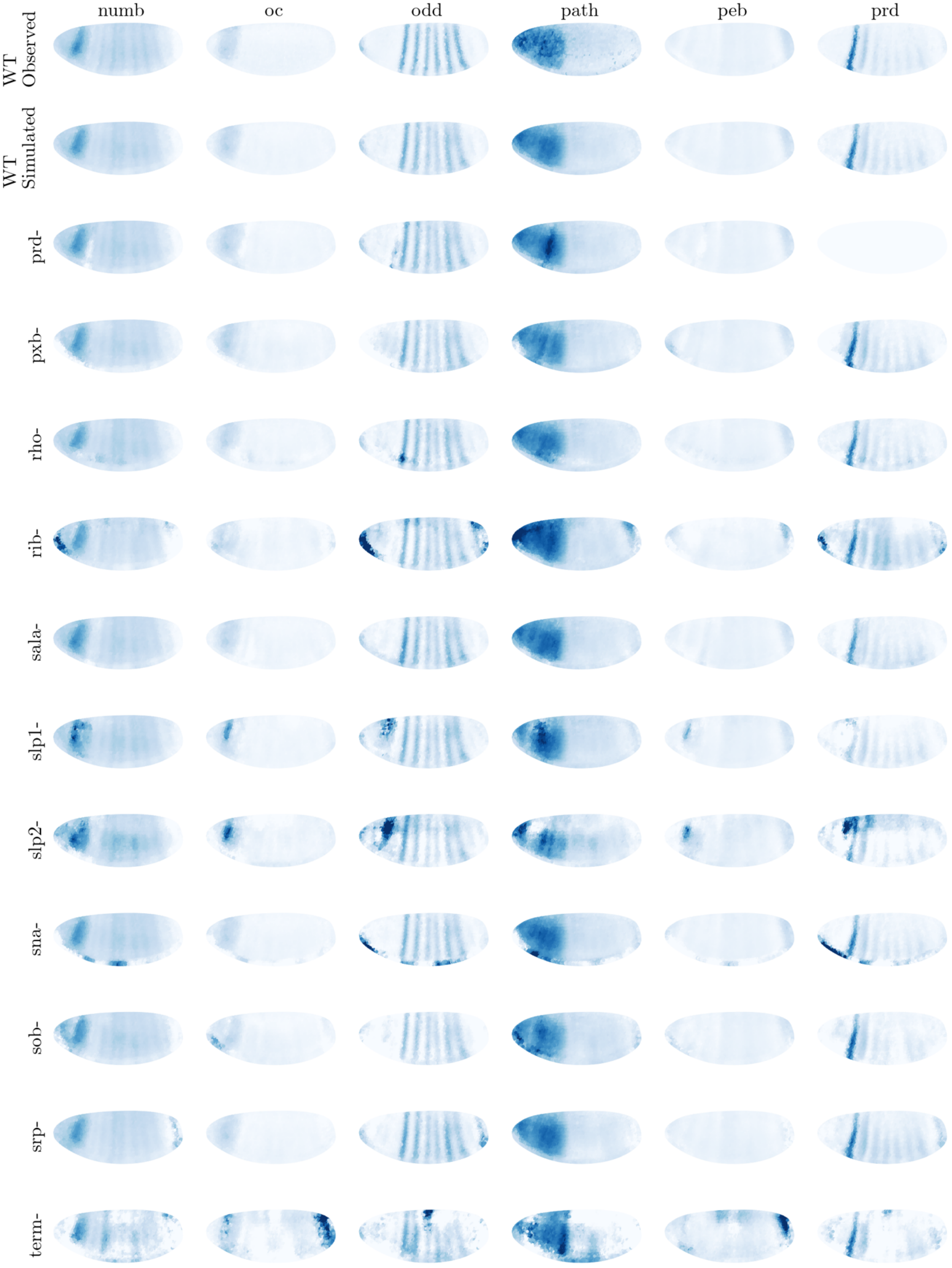
numb
oc
odd
path
peb
prd
WT Observed
WT Simulated
prd-
pxb-
rho-
rib-
sala-
slp1-
slp2-
sna-
sob-
srp-
term-

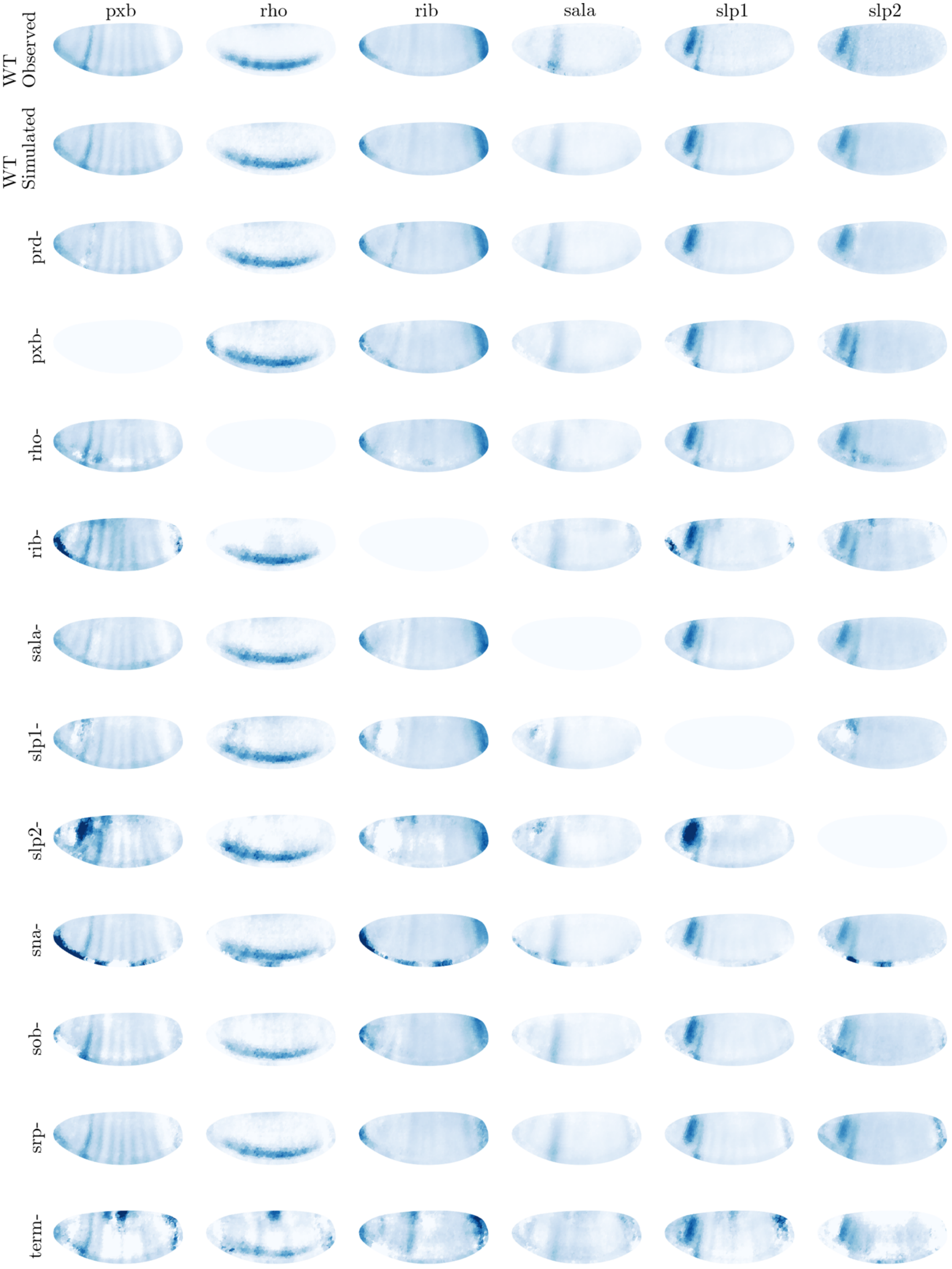
pxb
rho
rib
sala
slp1
slp2
WT Observed
WT Simulated
prd-
pxb-
rho-
rib-
sala-
slp1-
slp2-
sna-
sob-
srp-
term-

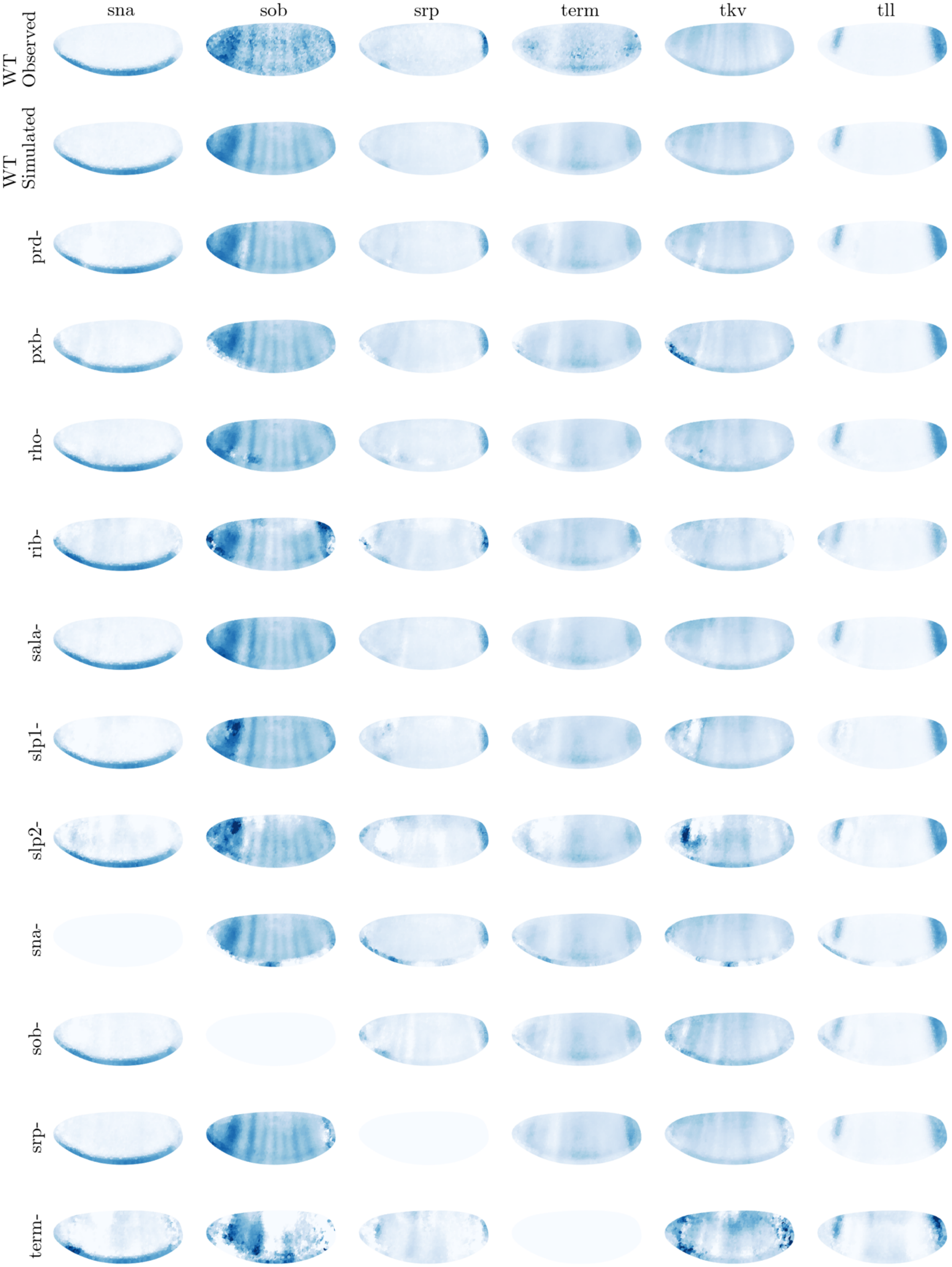
sna
sob
srp
term
tkv
tll
WT Observed
WT Simulated
prd-
pxb-
rho-
rib-
sala-
slp1-
slp2-
sna-
sob-
srp-
term-

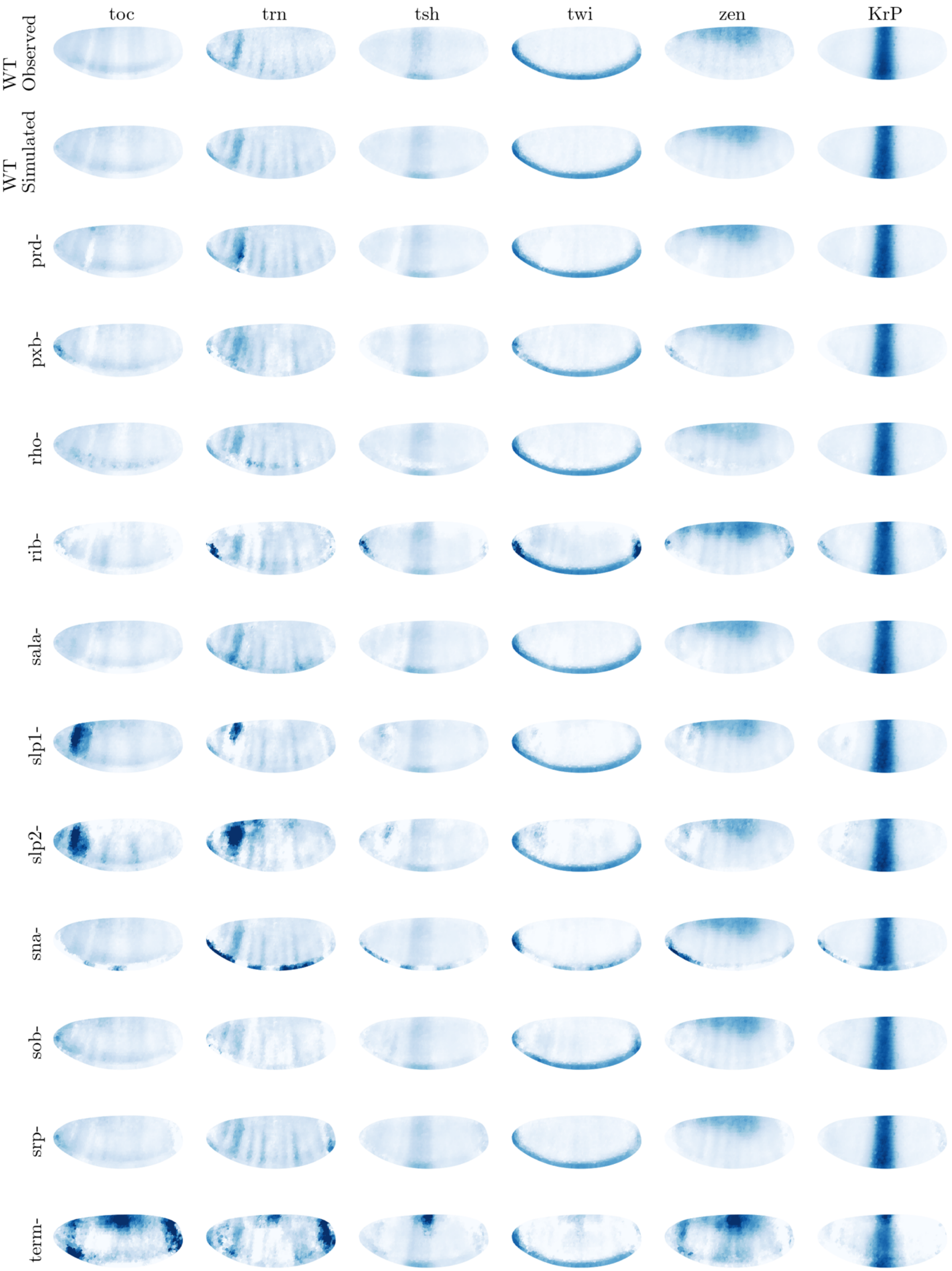
toc
trn
tsh
twi
zen
KrP
WT Observed
WT Simulated
prd-
pxb-
rho-
rib-
sala-
slp1-
slp2-
sna-
sob-
srp-
term-

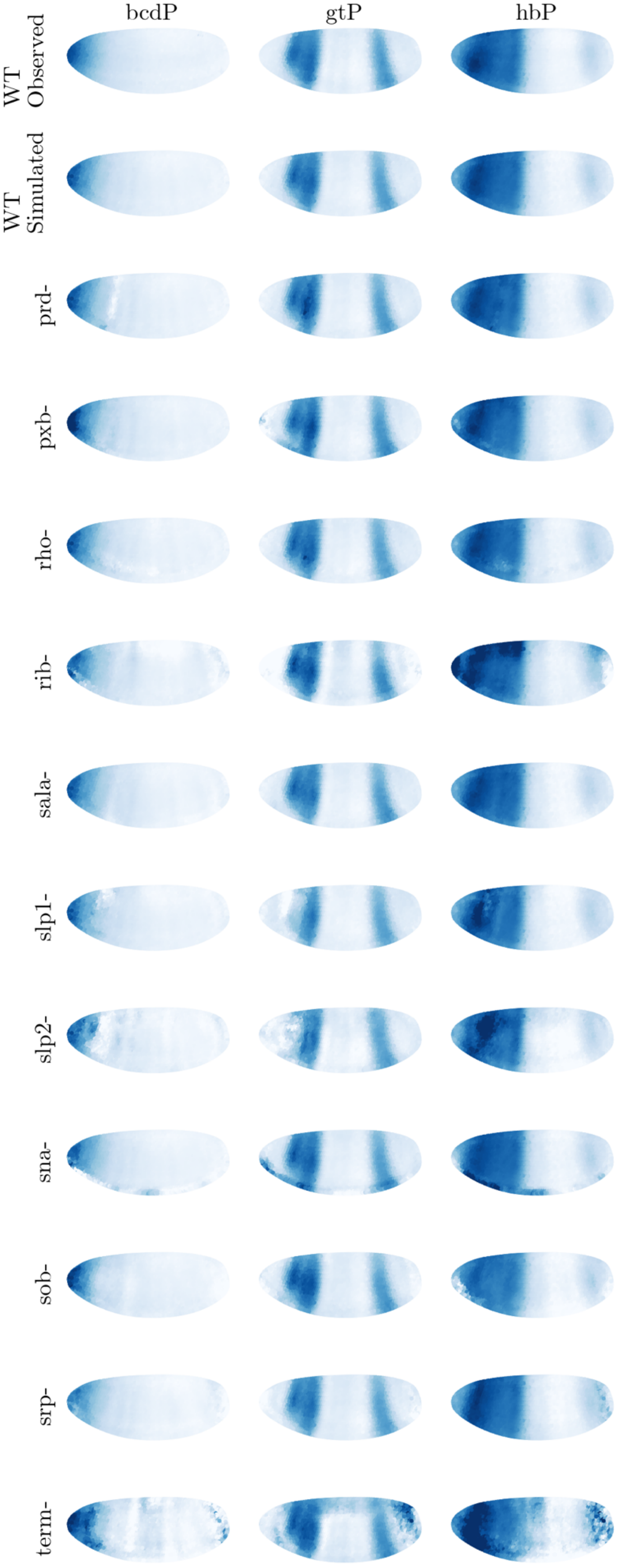
bcdP
gtP
hbP
WT Observed
WT Simulated
prd-
pxb-
rho-
rib-
sala-
slp1-
slp2-
sna-
sob-
srp-
term-

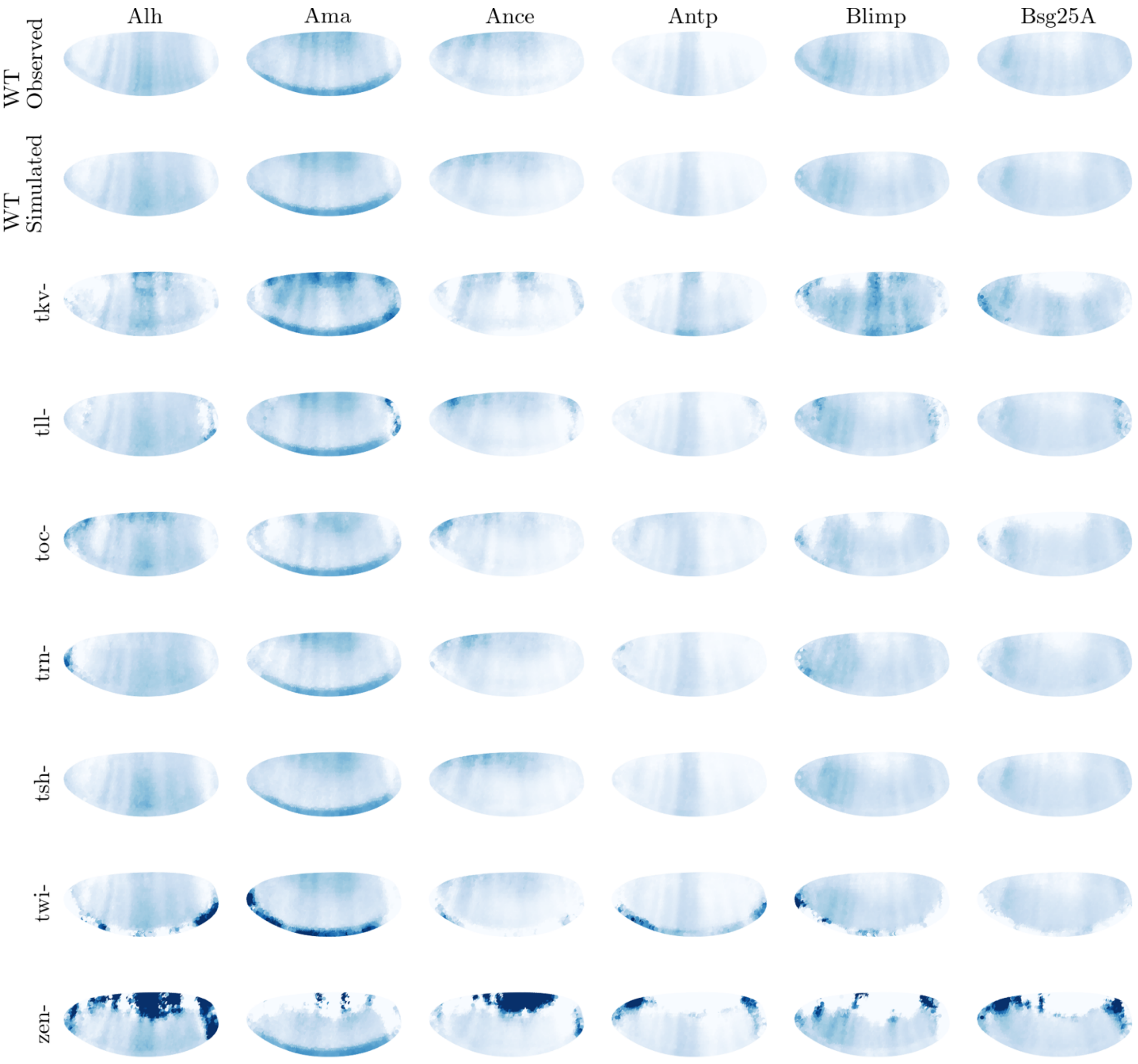
Alh
Ama
Ance
Antp
Blimp
Bsg25A
WT Observed
WT Simulated
tkv-
tll-
toc-
trn-
tsh-
twi-
zen-

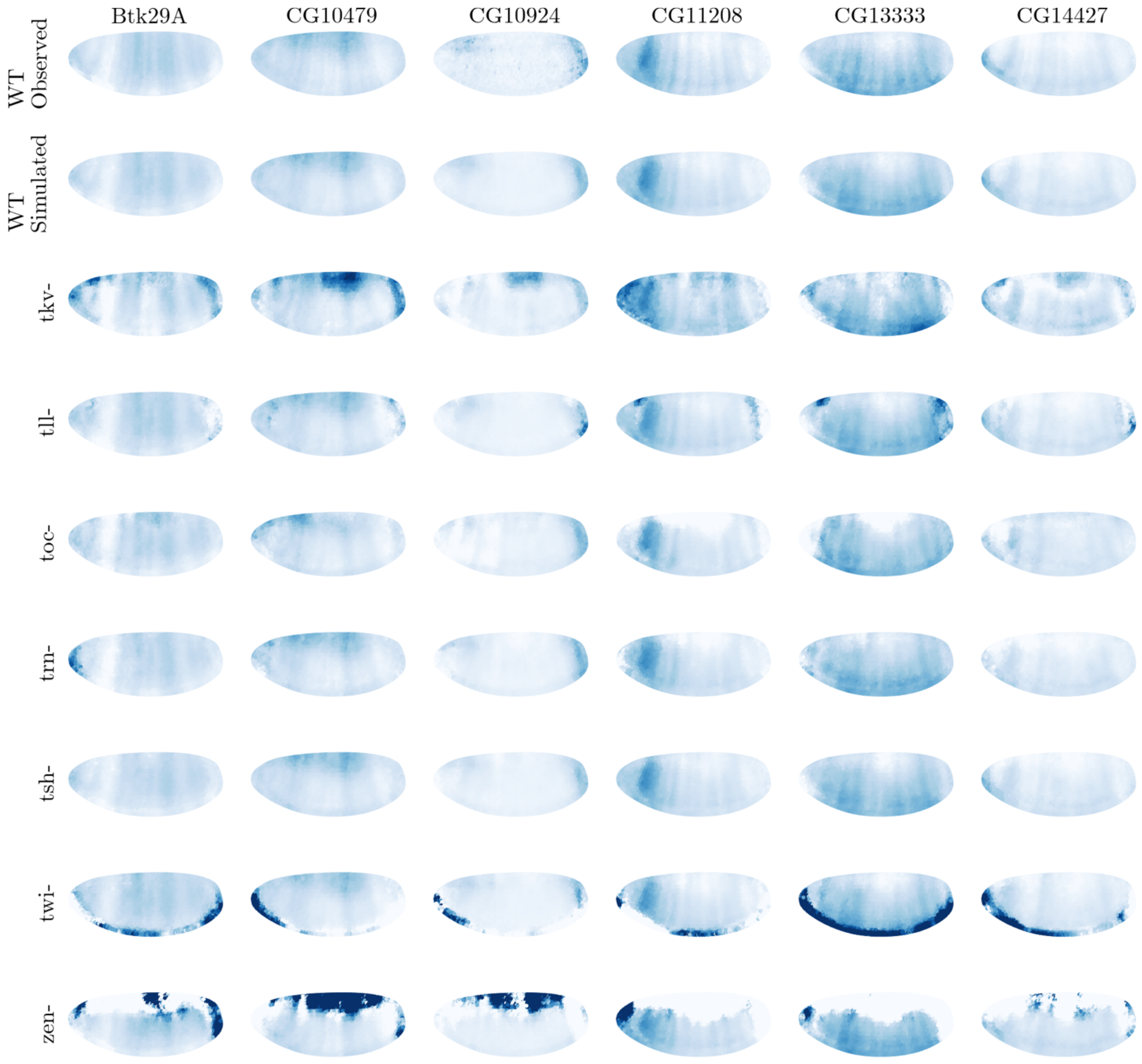
Btk29A
CG10479
CG10924
CG11208
CG13333
CG14427
WT Observed
WT Simulated
tkv-
tll-
toc-
trn-
tsh-
twi-
zen-

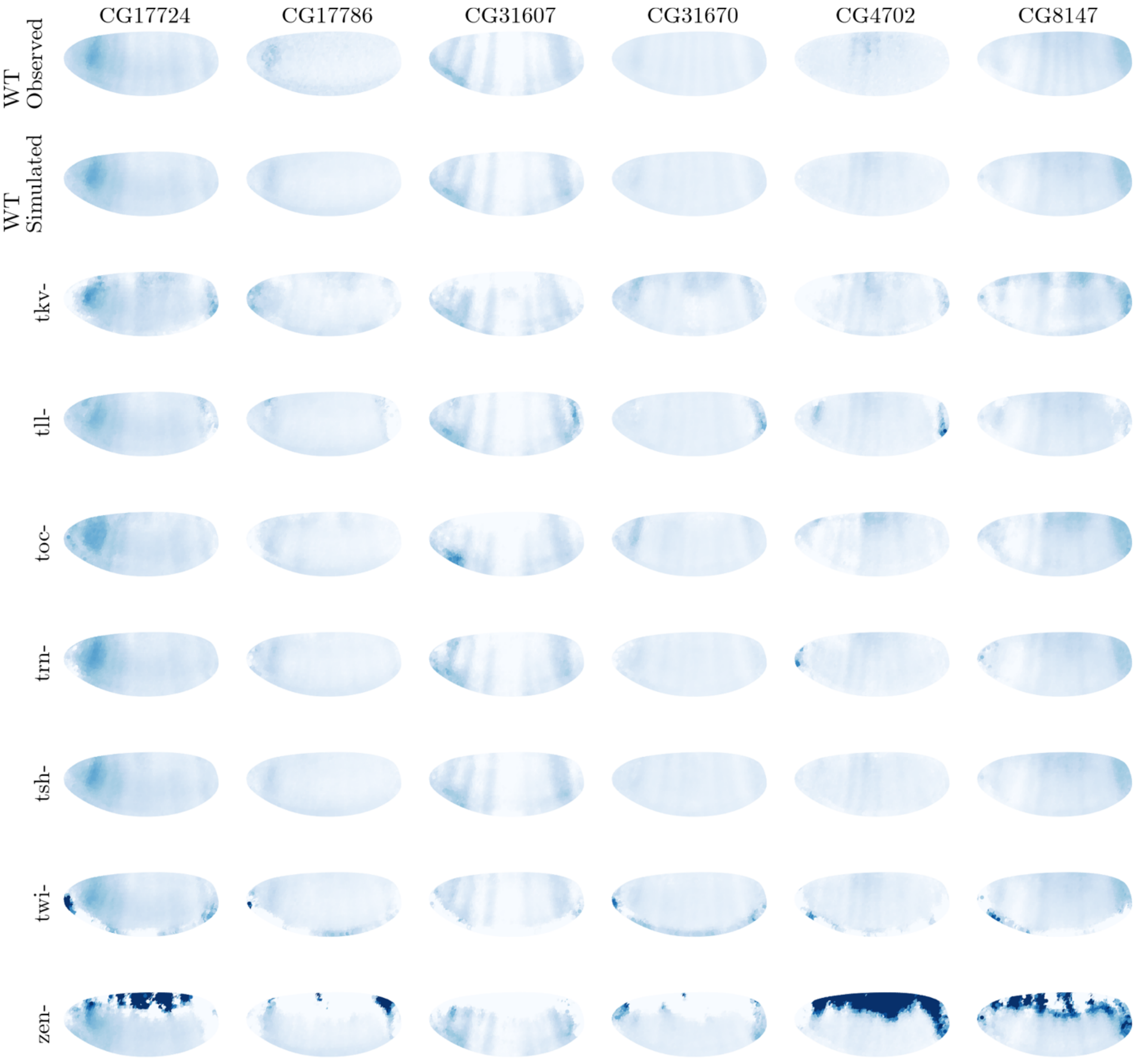
CG17724
CG17786
CG31607
CG31670
CG4702
CG8147
WT Observed
WT Simulated
tkv-
tll-
toc-
trn-
tsh-
twi-
zen-

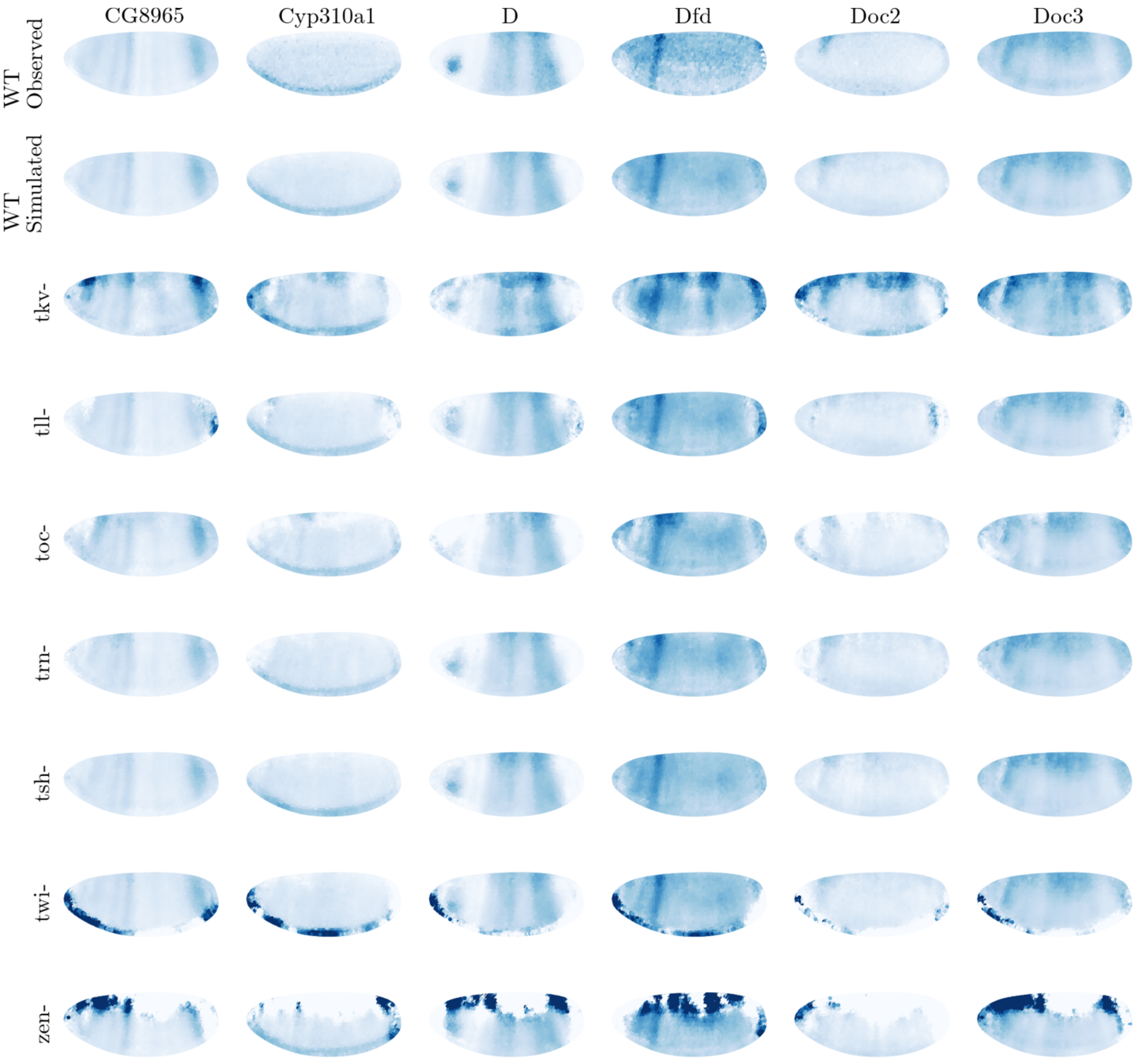

CG8965
Cyp310a1
D
Dfd
Doc2
Doc3
WT Observed
WT Simulated
tkv-
tll-
toc-
trn-
tsh-
twi-
zen-

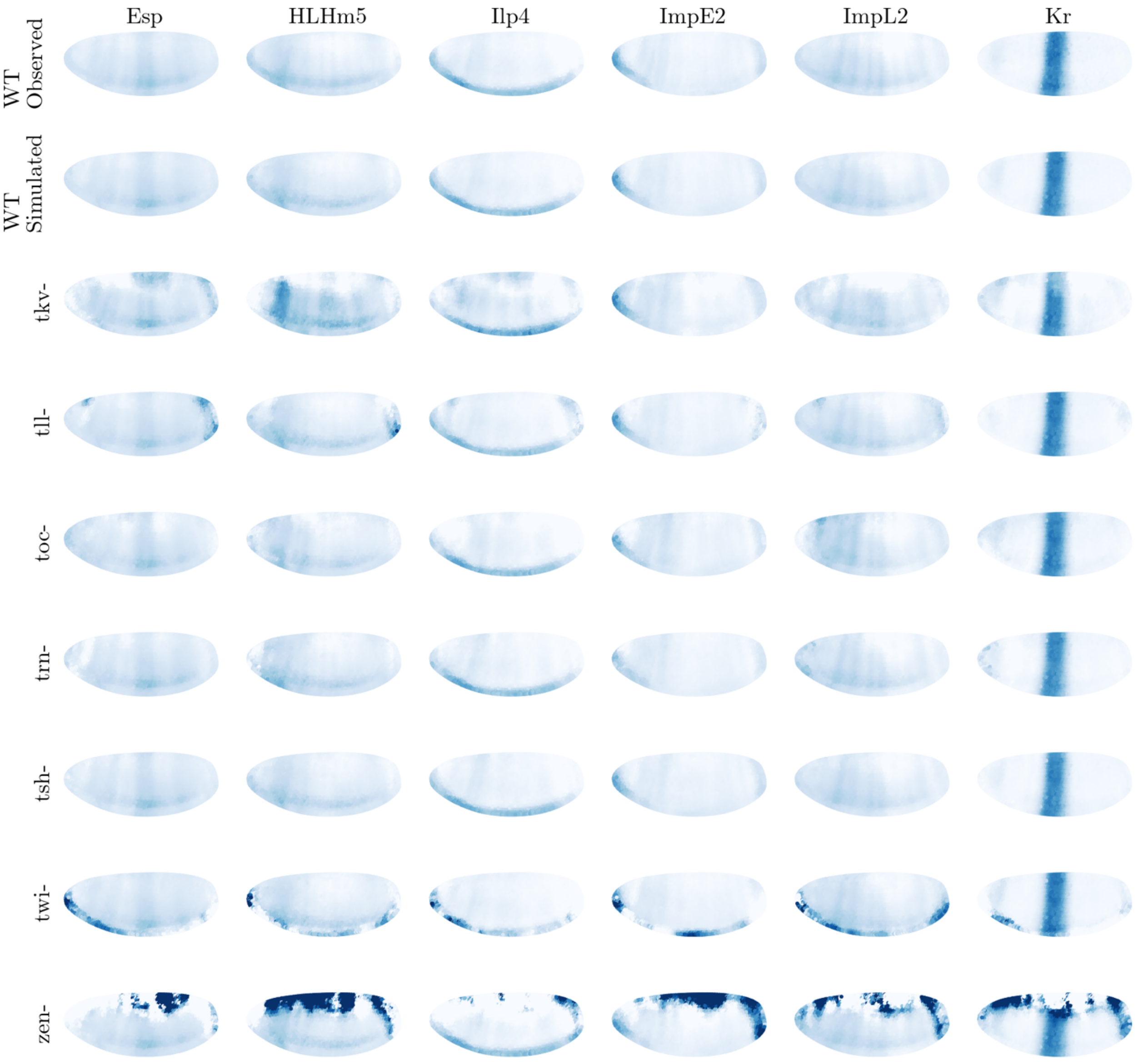
Esp
HLHm5
Ilp4
ImpE2
ImpL2
Kr
WT Observed
WT Simulated
tkv-
tll-
toc-
trn-
tsh-
twi-
zen-

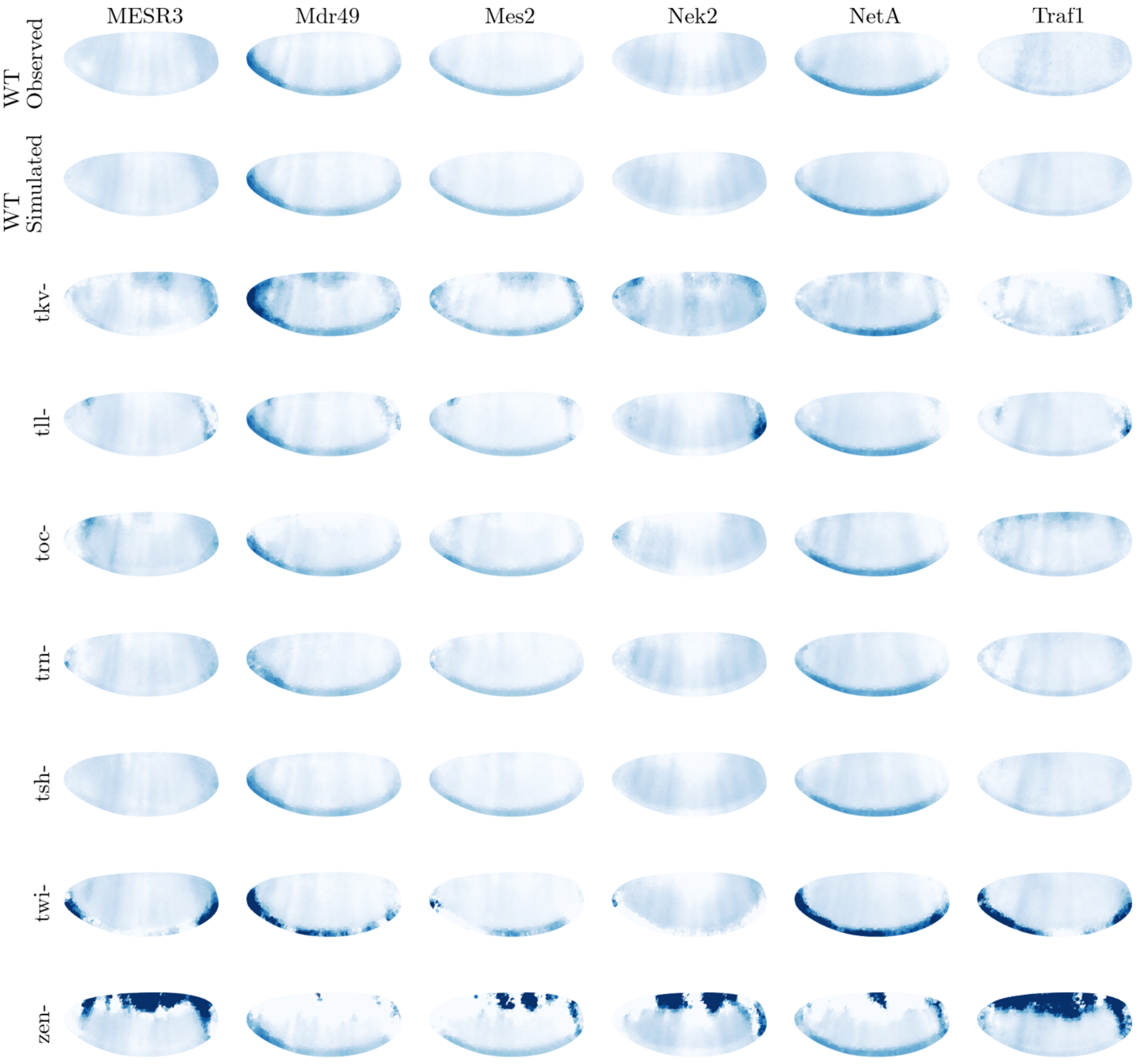
MESR3
Mdr49
Mes2
Nek2
NetA
Traf1
WT Observed
WT Simulated
tkv-
tll-
toc-
trn-
tsh-
twi-
zen-

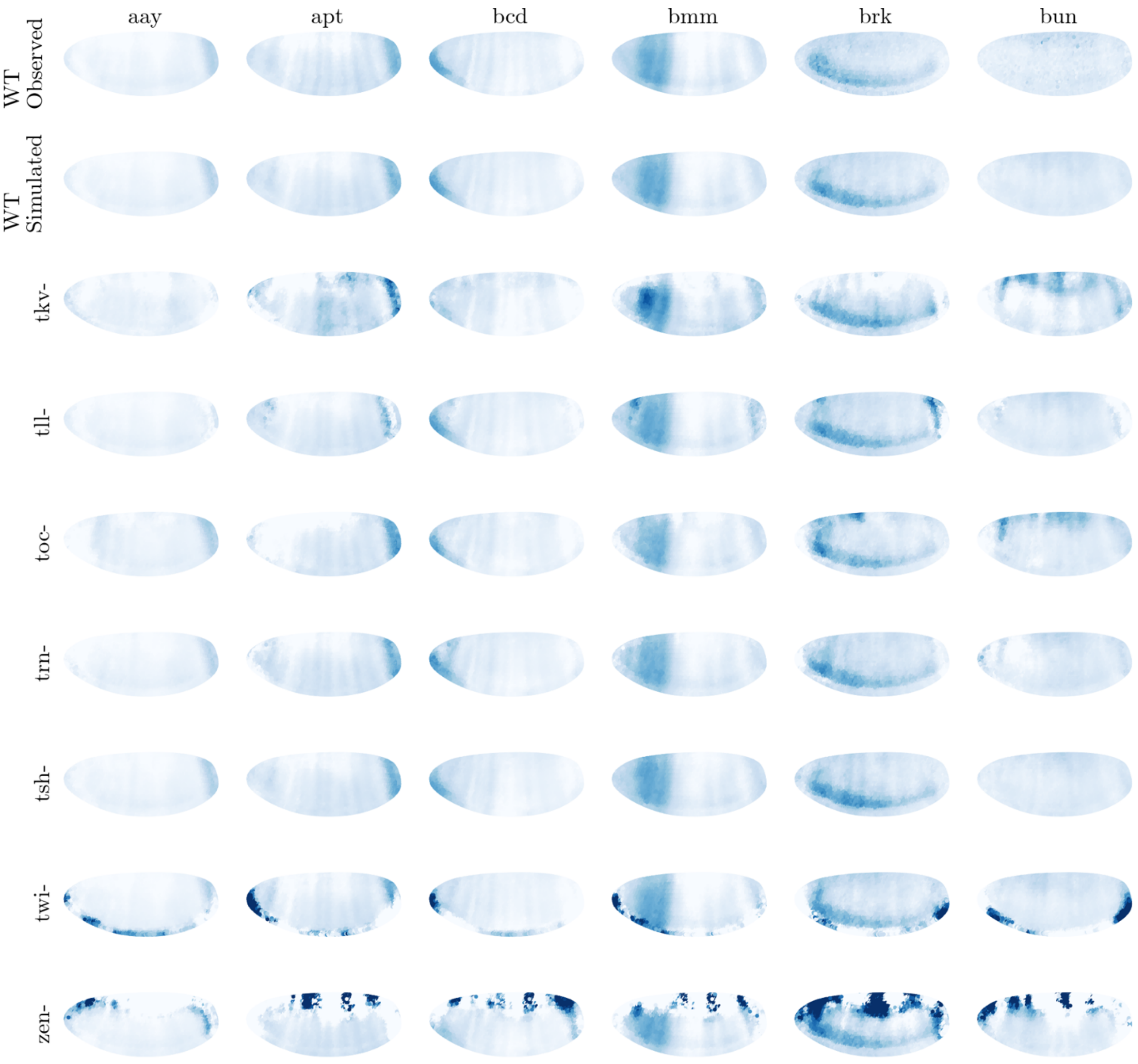
aay
apt
bcd
bmm
brk
bun
WT Observed
WT Simulated
tkv-
tll-
toc-
trn-
tsh-
twi-
zen-

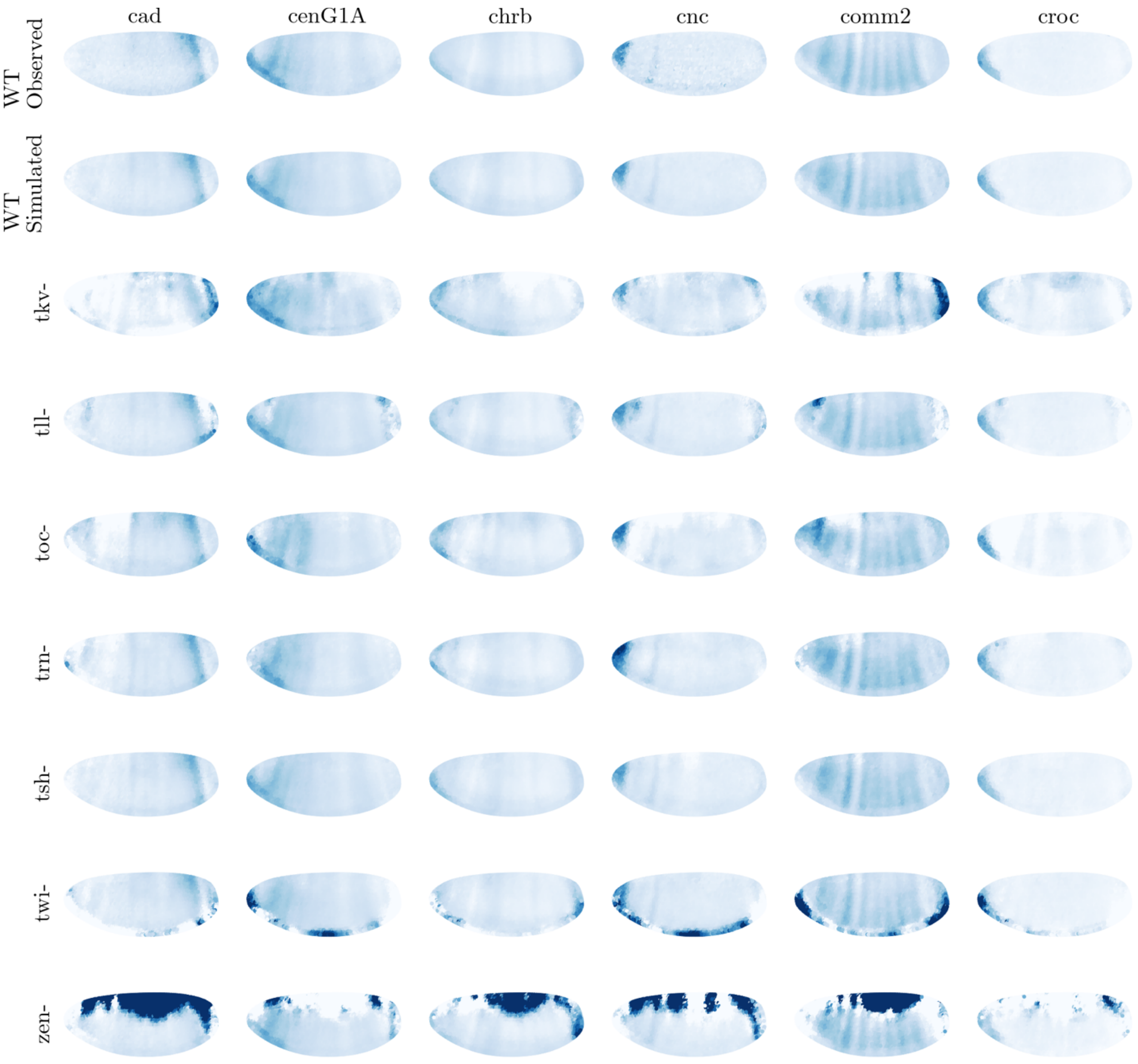
cad
cenG1A
chrb
cnc
comm2
croc
WT Observed
WT Simulated
tkv-
tll-
toc-
trn-
tsh-
twi-
zen-

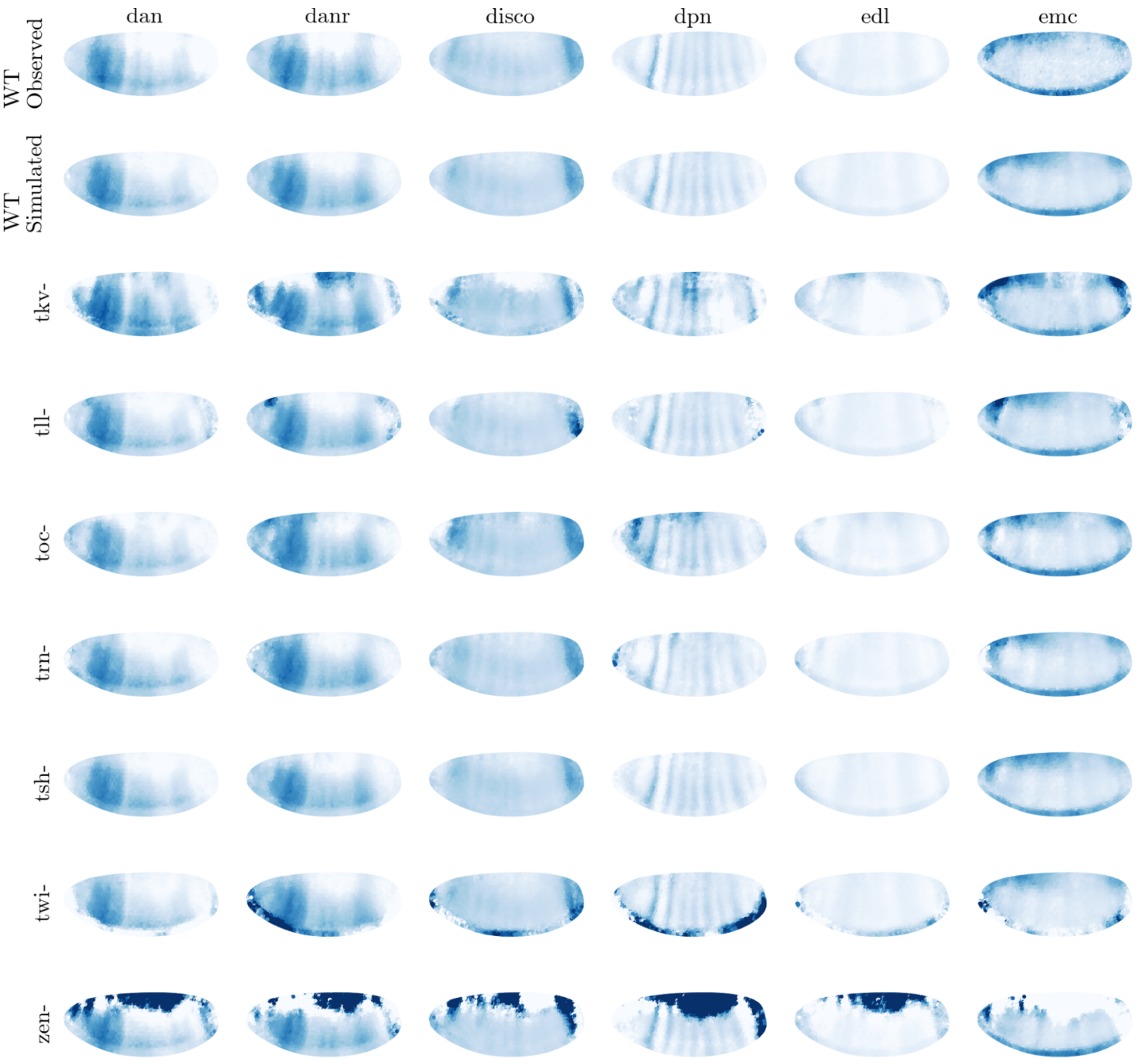

dan
danr
disco
dpn
edl
emc
WT Observed
WT Simulated
tkv-
tll-
toc-
trn-
tsh-
twi-
zen-

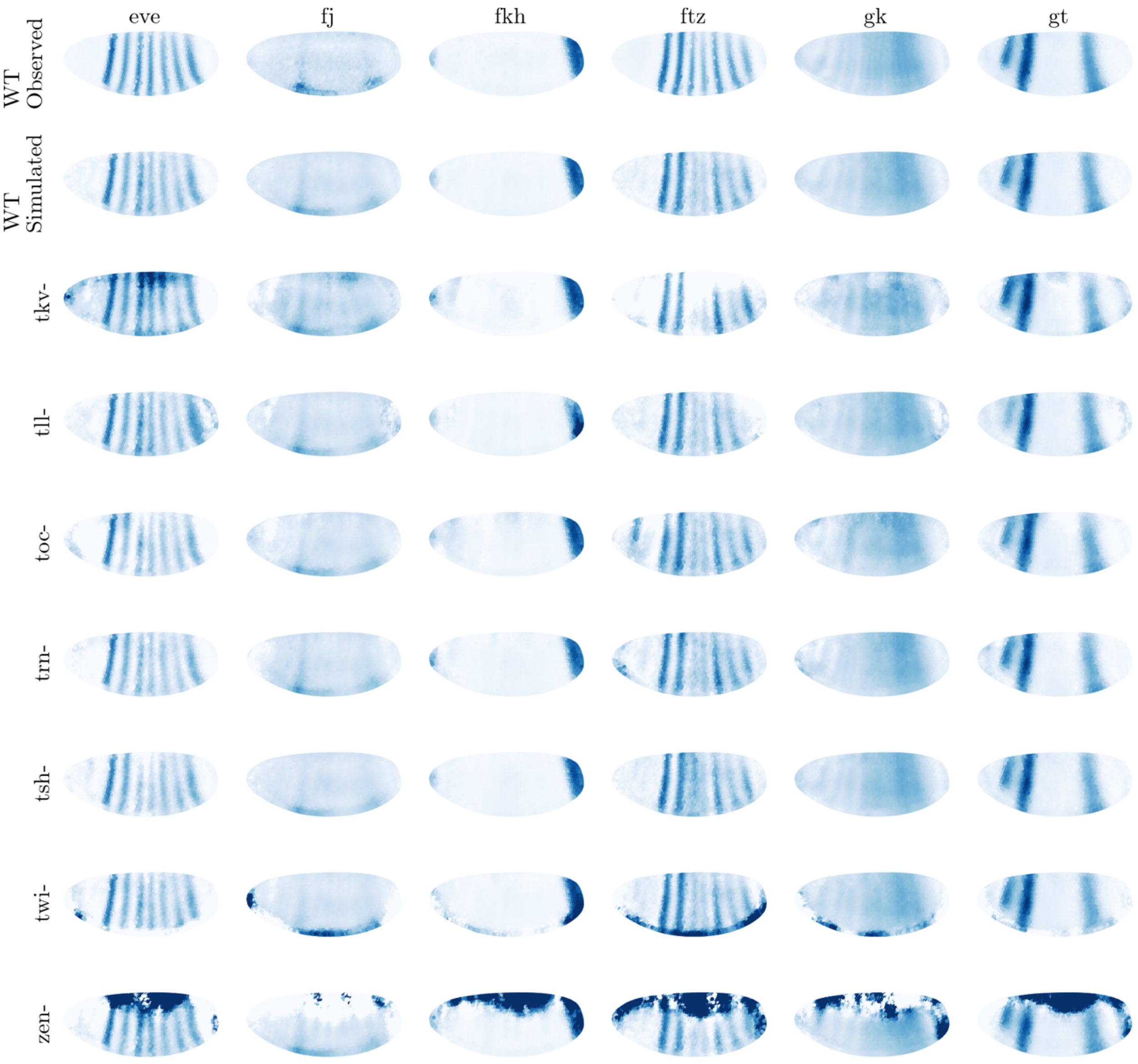
eve
fj
fkh
ftz
gk
gt
WT Observed
WT Simulated
tkv-
tll-
toc-
trn-
tsh-
twi-
zen-

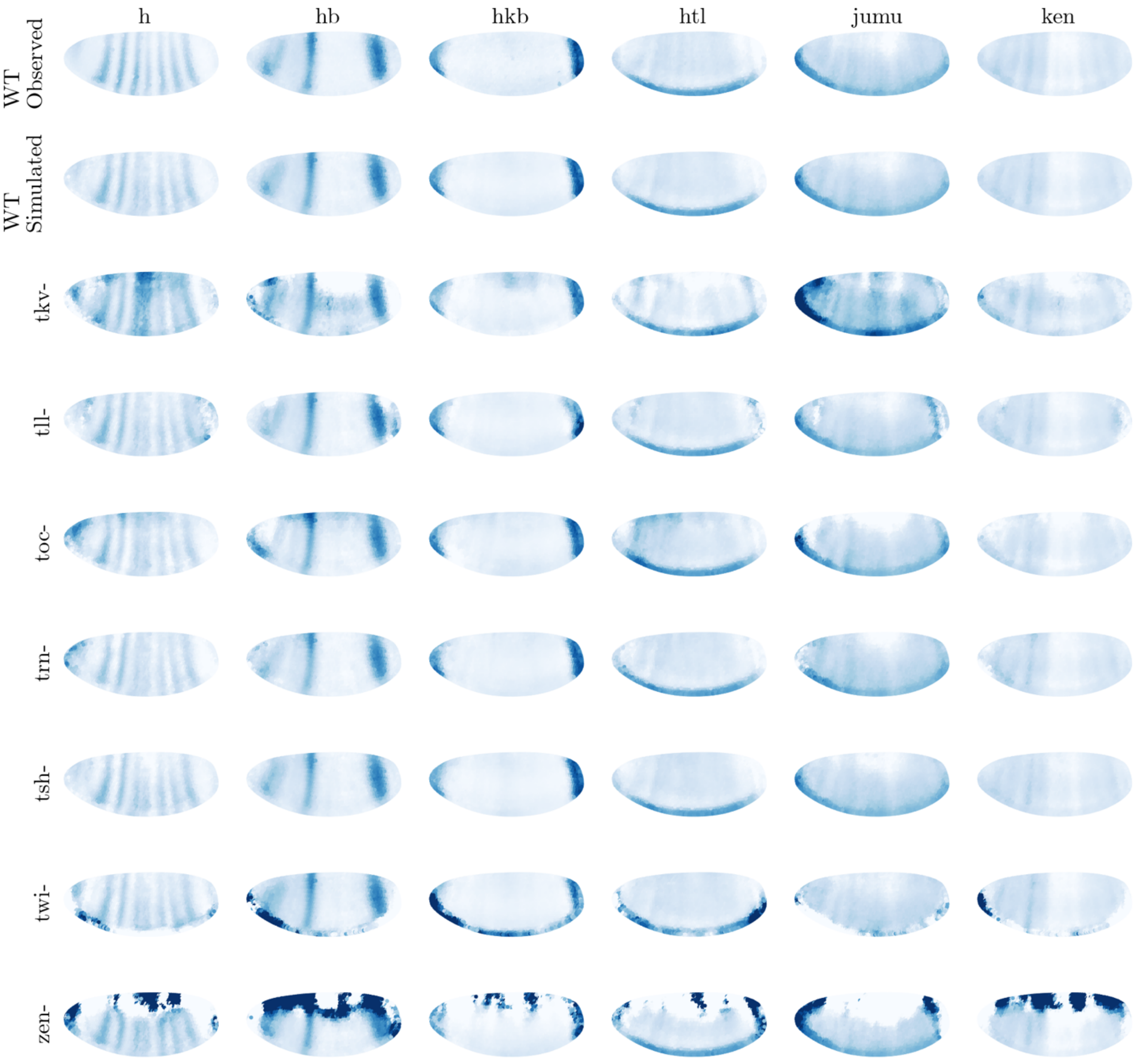

h
hb
hkb
htl
jumu
ken
WT Observed
WT Simulated
tkv-
tll-
toc-
trn-
tsh-
twi-
zen-

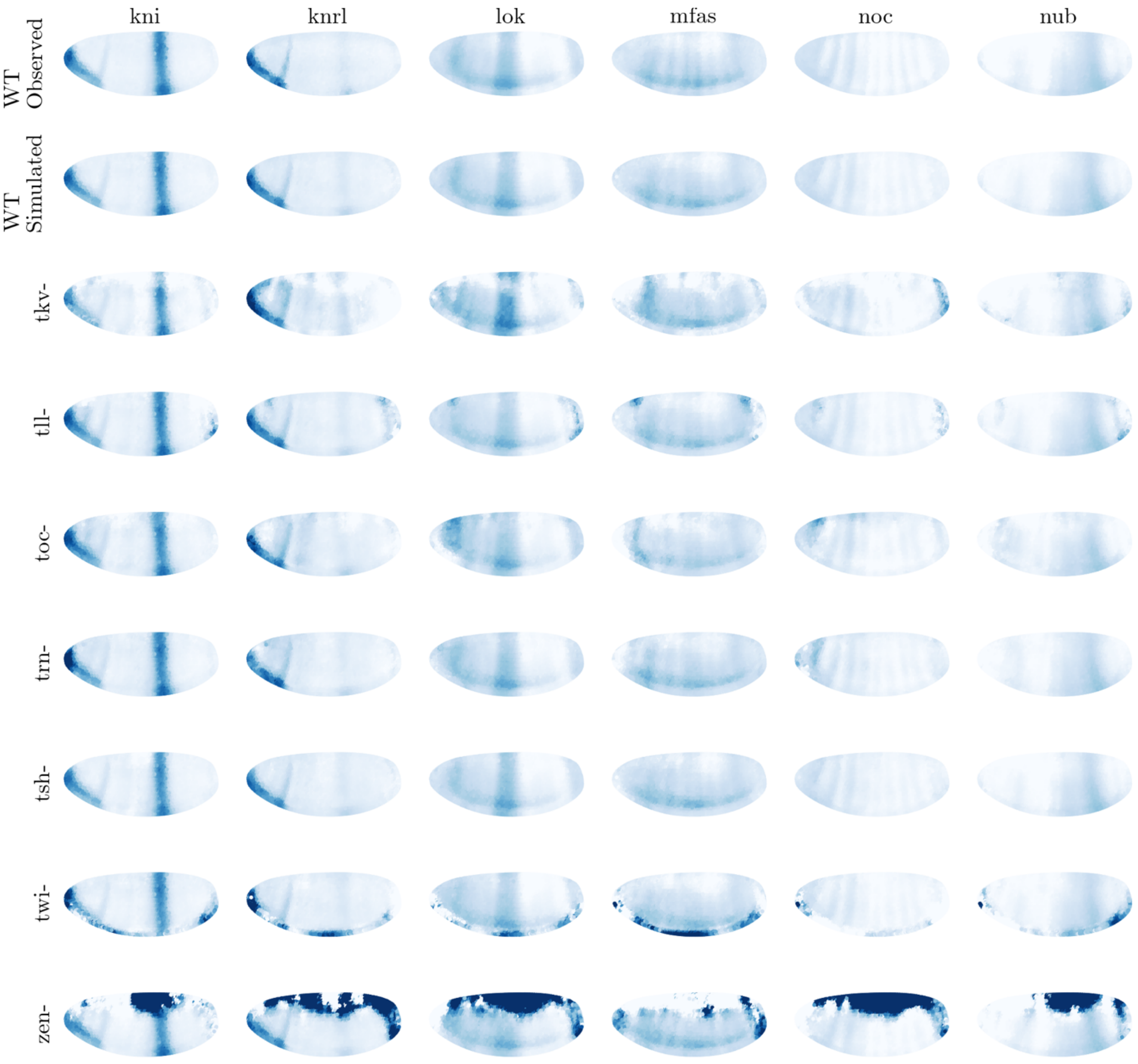
kni
knrl
lok
mfas
noc
nub
WT Observed
WT Simulated
tkv-
tll-
toc-
trn-
tsh-
twi-
zen-

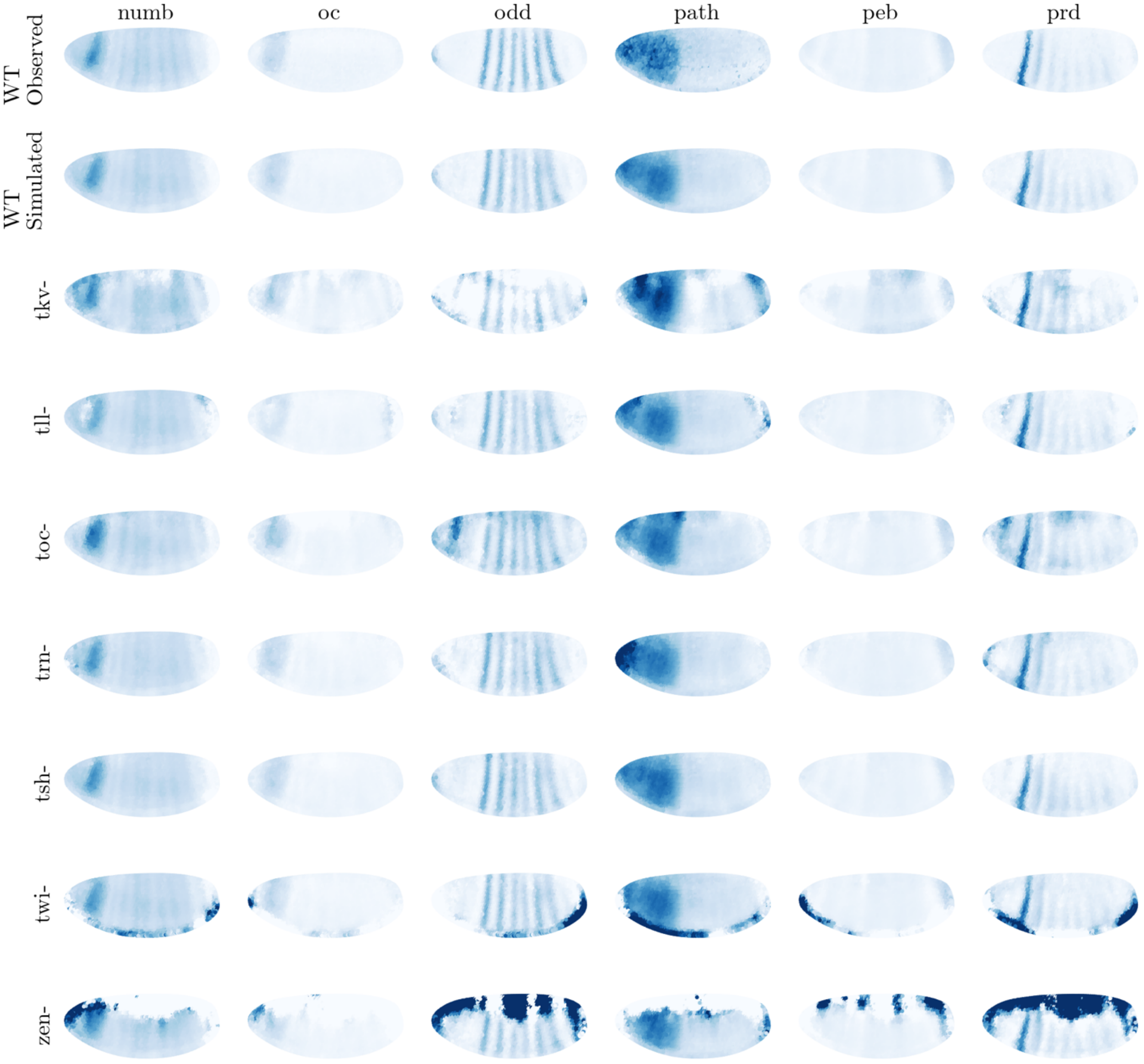
numb
oc
odd
path
peb
prd
WT Observed
WT Simulated
tkv-
tll-
toc-
trn-
tsh-
twi-
zen-

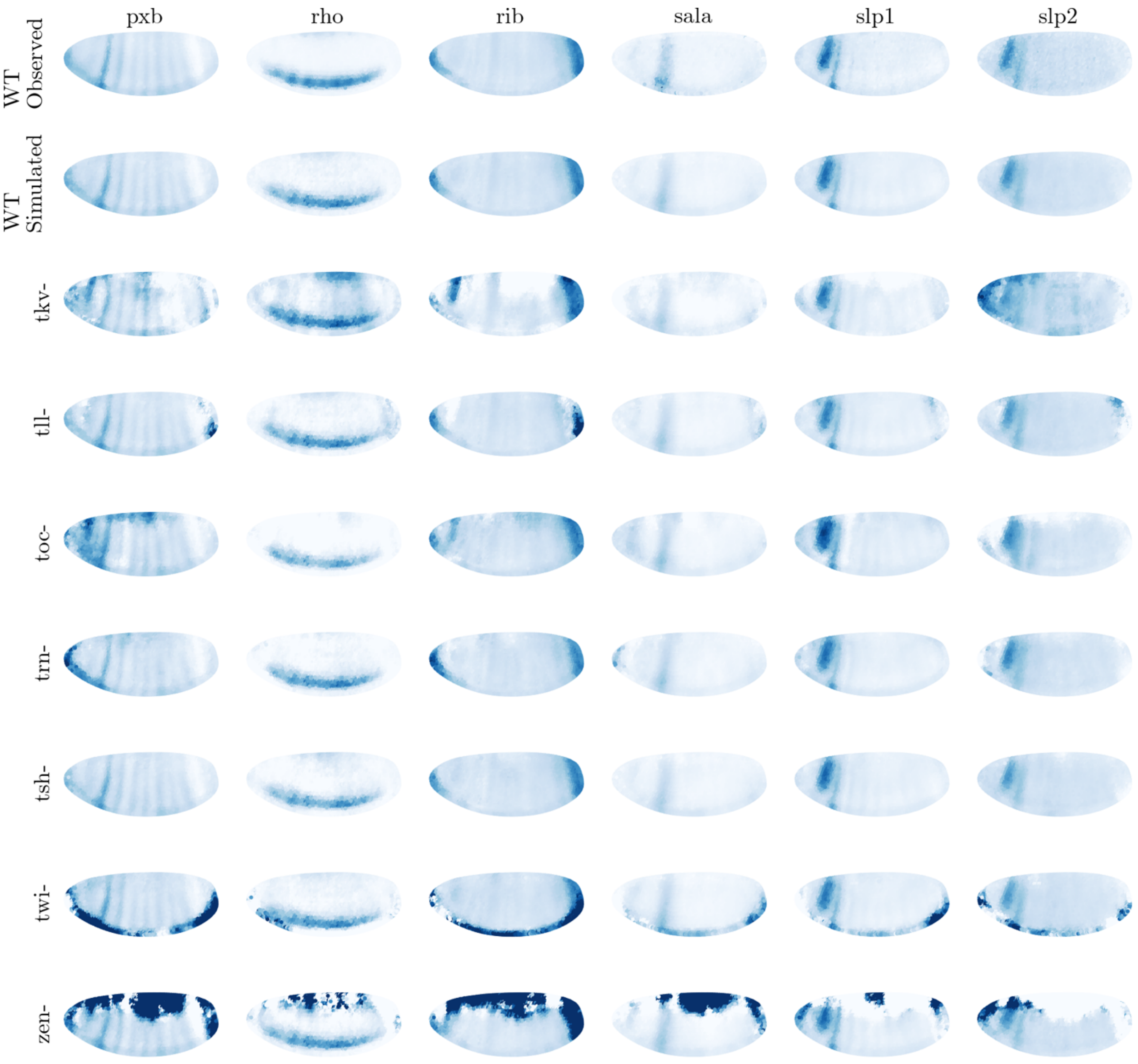
pxb
rho
rib
sala
slp1
slp2
WT Observed
WT Simulated
tkv-
tll-
toc-
trn-
tsh-
twi-
zen-

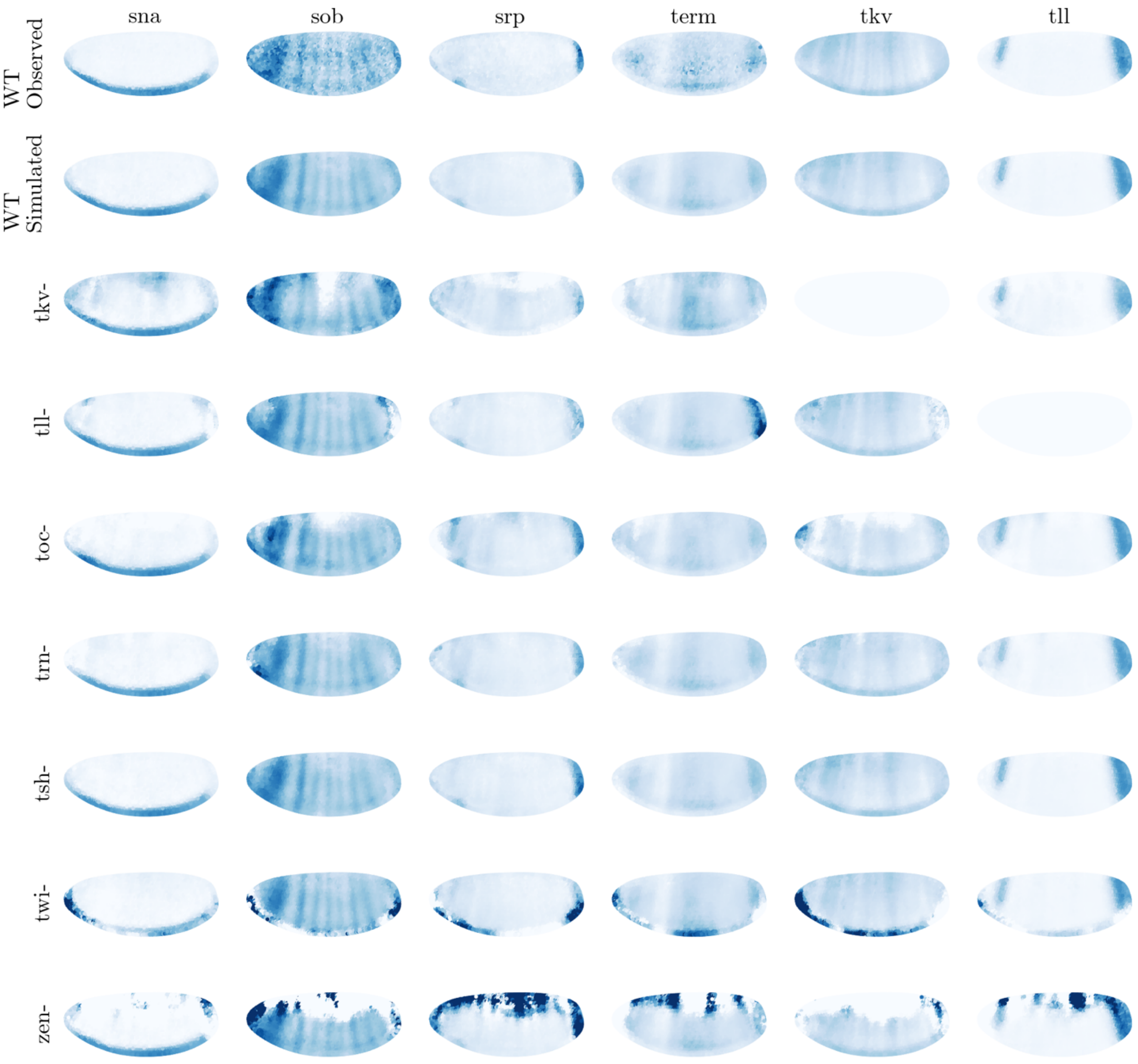
sna
sob
srp
term
tkv
tll
WT Observed
WT Simulated
tkv-
tll-
toc-
trn-
tsh-
twi-
zen-

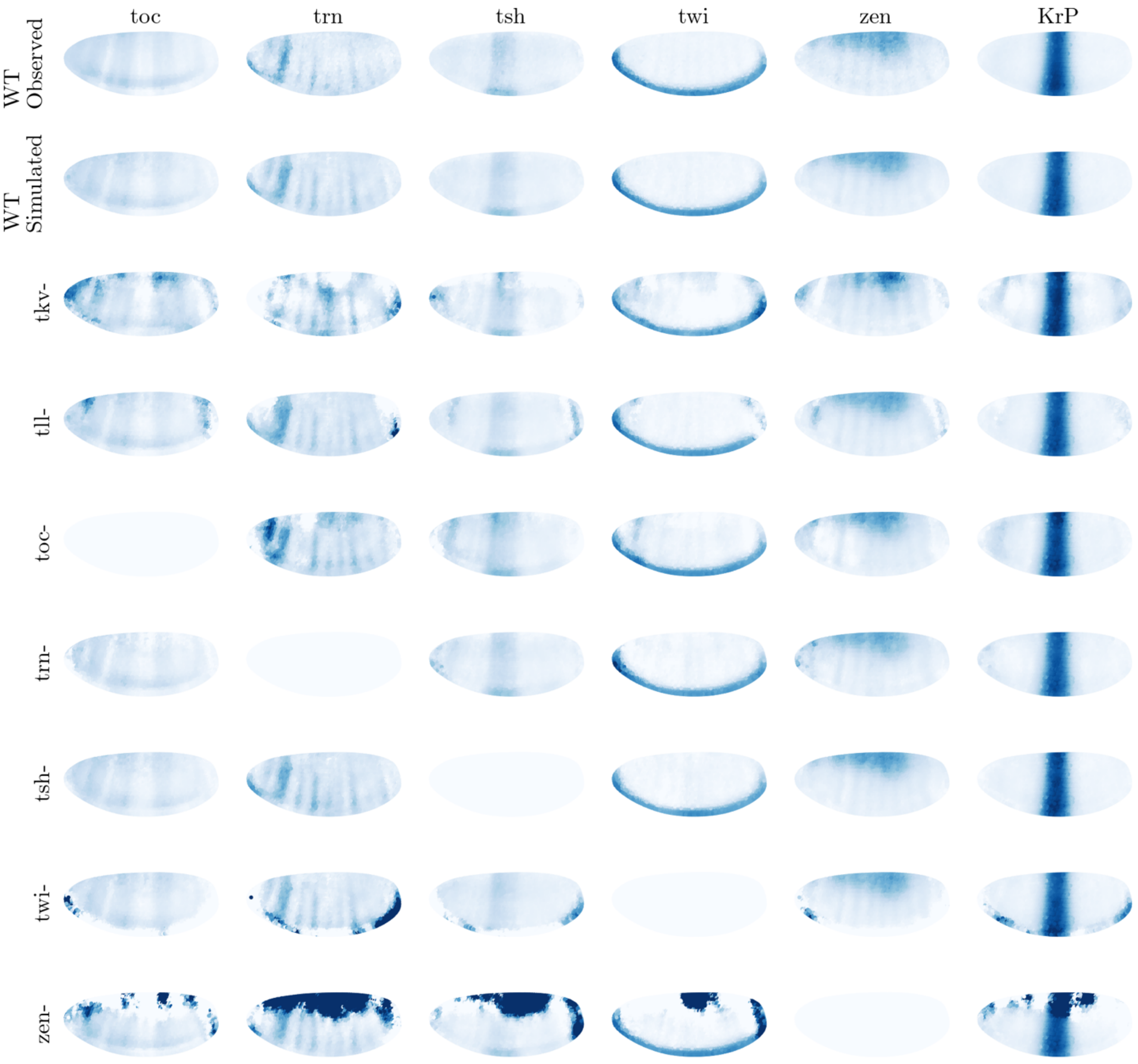
toc
trn
tsh
twi
zen
KrP
WT Observed
WT Simulated
tkv-
tll-
toc-
trn-
tsh-
twi-
zen-

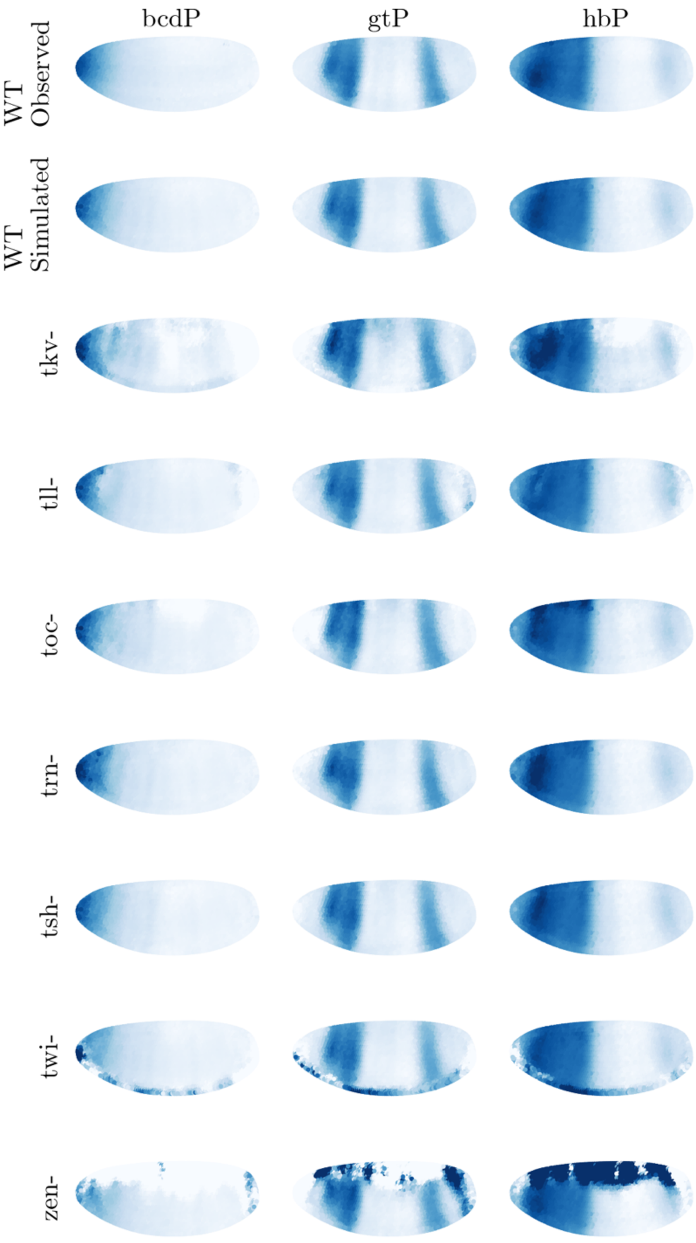

bcdP
gtP
hbP
WT Observed
WT Simulated
tkv-
tll-
toc-
trn-
tsh-
twi-
zen-
